# Identifying Candidate Genes for Sugar Accumulation in Sugarcane Cultivars: From a Syntenic Genomic Region to a Gene Coexpression Network

**DOI:** 10.1101/2024.05.08.593213

**Authors:** Mônica Letícia Turibio Martins, Danilo Augusto Sforça, Luís Paulo dos Santos, Ricardo José Gonzaga Pimenta, Melina Cristina Mancini, Alexandre Hild Aono, Cláudio Benício Cardoso da Silva, Sonia Vautrin, Arnaud Bellec, Renato Vicentini, Helene Bérgès, Carla Cristina da Silva, Anete Pereira de Souza

## Abstract

Elucidating the intricacies of the sugarcane genome is essential for breeding superior cultivars. This economically important crop originates from hybridizations of highly polyploid *Saccharum* species. However, the large size (10 Gb), high polyploidy, and aneuploidy of the sugarcane genome pose significant challenges to complete genome sequencing, assembly, and annotation. One successful strategy for identifying candidate genes linked to agronomic traits, particularly those associated with sugar accumulation, leverages synteny and potential collinearity with related species. In this study, we explored synteny between sorghum and sugarcane. Genes from a sorghum Brix QTL were used to screen bacterial artificial chromosome (BAC) libraries from two Brazilian sugarcane varieties (IACSP93-3046 and SP80-3280). The entire region was successfully recovered, confirming synteny and collinearity between the species. Manual annotation identified 51 genes in the hybrid varieties that were subsequently confirmed to be present in *Saccharum spontaneum*. To identify candidate genes for sugar accumulation, this study employed a multifaceted approach, including retrieving the genomic region of interest, performing gene-by-gene analysis, analyzing RNA-seq data of internodes from *Saccharum officinarum* and *S. spontaneum* accessions, constructing a coexpression network to examine the expression patterns of genes within the studied region and their neighbors, and finally identifying differentially expressed genes (DEGs). This comprehensive approach led to the discovery of three candidate genes potentially involved in sugar accumulation: an ethylene-responsive transcription factor (ERF), an ABA 8’-hydroxylase, and a prolyl oligopeptidase (POP). These findings could be valuable for identifying additional candidate genes for other important agricultural traits and directly targeting candidate genes for further work in molecular breeding.

## Introduction

In the 1880s, sugarcane (*Saccharum* spp.) farmers crossed *Saccharum spontaneum* (2n=5x=40 to 16x=128, x=8), which is resistant to biotic and abiotic stress, with *Saccharum officinarum* (2n=8x=80, x=10), which is considered a noble sugarcane due its high amount of sugar. To maintain a high sugar content in hybrids, successive backcrosses with *S. officinarum* were performed (Bremer, 1961, D’Hont et al., 1996; Garsmeur et al., 2018; Babu et al., 2022 and Healey et al., 2024). The crosses between both species generated modern sugarcane cultivars: plants with large genomes (10 Gb) that are highly polyploid and aneuploid with at least 50% repetitive regions (Cuadrado, 2004; Piperidis et al., 2010; Garcia et al., 2013; Thirugnanasambandam et al., 2018, Healey et al., 2024). The hybrid genome is a mixture of chromosomes originating from *S. officinarum* (70-80% of all chromosomes of the hybrids) and *S. spontaneum* (10-20%) and recombinant chromosomes (5-10%) (D’Hont et al., 1996; Cuadrado, 2004; Piperidis et al., 2010; Garsmeur et al., 2018 and Healey et al., 2024). The variable ploidy intrinsic to each genotype creates a unique genomic structure with chromosome numbers varying between 40 and 128 (D’Hont et al., 1996; Zhang et al., 2012; Garsmeur et al., 2018; Thirugnanasambandam et al., 2018), which renders the study of the sugarcane genome a challenge (Thirugnanasambandam et al., 2018, Babu et al., 2022).

Grasses with a reference genome, such as rice (International Rice Genome Sequencing Project and Sasaki, 2005), maize (Schnable et al., 2009), wheat (IWGSC et al., 2018), and miscanthus *(Miscanthus sinensis)* (Kim et al., 2014; Tsuruta et al., 2017), even sugarcane with an allele-defined genome of *Saccharum spontaneum* (Zhang et al., 2018) or a monoploid sequence reference for sugarcane (Garsmeur et al., 2018) and sorghum (McCormick et al., 2018), are commonly used as references for studies on the sugarcane genome (Thirugnanasambandam et al, 2018; Wang et al., 2021; Babu et al., 2022, Mancini et al., 2018; Garsmeur et al., 2018; Zhang et al., 2018, Zhang et al., 2022 and Healey et al., 2024). Recently, a highly representative genome of the R570 hybrid variety was presented to the scientific community (Healey et al., 2024). The *Miscanthus* genome has sorghum as an important reference for its assembly and annotation (Mitros et al., 2020), and sorghum is an ancestor of the *Saccharum* and *Miscanthus* groups. Sorghum has an assembled and annotated diploid genome that is one-tenth of the sugarcane genome size and diverged approximately 8 million years ago; nevertheless, its genome has maintained strong synteny and collinearity (Ming et al., 1998; Paterson et al., 2009; Wang et al., 2010; Garsmeur et al., 2018, Thirugnanasambandam et al., 2018; Babu et al., 2022). The R570 has a high inbreeding coefficient, with approximately half being identical by descent. Therefore, it is expected that its genome will have a small portion missing and another collapsed, totaling 8.72 Gb, approaching the estimated size of 10 Gb (Healey et al., 2024).

Therefore, choosing a sorghum quantitative trait locus (QTL) for a trait of interest and recovering its orthologous region in sugarcane can be an efficient strategy to retrieve a potentially target sugarcane genomic region (Mancini et al., 2018). Recent studies with the R570 variety confirmed that a significant portion of the alleles originated from *S. officinarum,* so the sugar-accumulating origin is identical and thus largely inaccessible to QTL mapping efforts (Healey et al., 2024). However, the genomic sequence of a cultivar does not fully reflect the genetic information about the species (Montenegro et al., 2017), where a cultivar may not be representative of the entire genomic content of the species (Garsmeur et al., 2018; Thirugnanasambandam et al., 2018 and Healey et al., 2024). The development of new commercial sugarcane varieties with higher yields is the main goal of most breeding programs. To increase yield, they seek genotypes that can tolerate biotic and abiotic stress but also have increased sugar accumulation (CURSI et al., 2021 and Healey et al., 2024). The exploration of candidate genes can be aided by the use of other omic technologies, such as RNA sequencing (RNA-seq), which offers a differential assessment not only between specific tissues but also between notably different varieties and related species (Stark et al., 2019). The transcriptome allows for the identification of differentially expressed genes (DEGs) and can be leveraged to construct coexpression networks, aiming to identify coexpressed genes and expand the evidence leading to candidate genes.

Many genes involved in the synthesis and transport of sucrose have been identified in sugarcane (Zhu et al. 2000; Carson and Botha 2002; Grivet and Arruda 2002; Casu et al. 2003; Vasantha et al., 2022). Although sucrose is synthesized in the cytosol of mesophyll cells in most plants, sugarcane requires the involvement of two cell types: the bundle sheath and mesophyll. Sucrose synthesis occurs predominantly in the mesophyll, utilizing glucose phosphates, which are then translocated through the conducting strands of sheath to the vascular compartments of internodal tissues, where they finally accumulate (Vasantha et al., 2022). Moreover, during the plant maturation phase, the sucrose concentration in culms increases, while the proportion of glucose and fructose decreases (Chandra et al., 2012). However, many processes related to sugar accumulation in sugarcane internodes are not fully understood, and possible pathways and related genes have yet to be identified.

The accumulation of proline in plant cells is associated with various physiological processes, such as cellular homeostasis, aiding in water absorption, and adaptation to abiotic stresses, enhancing the plant’s adaptive response (Rejeb et al., 2014; Kazemi-Shahandashti & Maali-Amiri, 2018; Sharma et al., 2014; Kazemi-Shahandashti & Maali-Amiri, 2018; Sharma et al., 2014; Kazemi-Shahandashti & Maali-Amiri, 2018; Sharma et al., 2019; Ghosh et al., 2021). The interaction of proline accumulation with sucrose during salt stress has been reported in sugarcane (Ghosh et al., 2019). The enzyme prolyl oligopeptidase (POP—serine protease family clan SC, family S9) is a cytoplasmic enzyme that hydrolyzes oligopeptides up to 30 residues and occurs at the C-terminal side of proline residues (Gutierrez et al., 2008; Baharin et al., 2022). In plants, POP has been associated with responses to biotic and abiotic stresses (Gutierrez et al., 2008; Singh et al., 2011; Tan et al., 2013).

In this context, a sorghum QTL for Brix (Shiringani et al., 2010) in sorghum was chosen as a target because its orthologous region was recovered in two Brazilian cultivars (SP80-3280 and IACSP93-3046), in R570 (Garsmeur et al., 2018) and in the *S. spontaneum* genome (Zhang et al. 2018). These genomic regions were compared to understand the level of genomic structural variation and genetic differences among sugarcane and sorghum. The genes found in this region were used to search for candidate genes for sugar accumulation through gene annotation evaluation, differential expression analysis in sugarcane stem transcriptomes (Aono et al., 2021) and a coexpression network. The combination of such strategies provides a more comprehensive and robust perspective in the search for candidate genes related to sugar accumulation in sugarcane. In exploring the region’s genes, a set of evidence combining genetic/genomic factors revealed three candidate genes related to sugar accumulation characteristics.

## Materials and methods

### Sorghum region of interest

A partial QTL that was mapped in sorghum for Brix, which is genetically located on chromosome SBI-02 between the EST-SSR markers Xtxp56 and Stgnhsbm36 (Shiringani et al., 2010), was selected. The marker sequences were used to define the physical chromosomal position using the v3.1 version of the S*orghum bicolor* genome (Paterson et al., 2009) available in the Phytozome 13.0 database (https://phytozome-next.jgi.doe.gov/, Goodstein et al., 2012). The QTL has a phenotypic variation with 21.9% explained by genotype (R^2^) % and a logarithm of odds (LOD) value of 10.08 (Shiringani et al., 2010). It spans from 61,568 kb to 61,952 kb on the sorghum chromosome SBI-02, totaling an approximate length of 385 kb. The target region was defined between 61,500 kb and 62,000 kb.

### Plant material

Two Brazilian sugarcane cultivars were analyzed in the present study. SP80-3280 is known for its high production of sucrose and good tillering. It is resistant to smut, mosaic, and rust and tolerant to scald (Embrapa—Brazilian Agricultural Research Company, 2022). The SP80-3280 variety has been widely used in studies to understand sugarcane genomics and genetics. This variety has a collection of sugarcane expressed sequence tags (SUCEST, Vettore, 2003), transcriptomes (Cardoso-Silva et al., 2014; Nishiyama et al., 2014; Mattiello et al., 2015), mapped QTLs (Aitken et al., 2006; Costa et al., 2016), a draft genome (Riaño-Pachón and Mattiello et al., 2017), and bacterial artificial chromosome (BAC) libraries (Figueira et al., 2012; Sforça et al., 2019). The use of sorghum synteny and collinearity has also been the focus of an approach for restoring genomic regions of agronomic interest (Mancini et al., 2018). The economic importance of the IACSP93-3046 cultivar is due to its high sucrose content, good tillering, resistance to rust and suitability for mechanized harvesting (Mancini et al., 2012). This cultivar also has a transcriptome (Cardoso-Silva et al., 2014) and a BAC library (Sforça et al., 2019).

### Recovering the sorghum ortholog region in sugarcane

#### Primer design

The coding sequences (CDSs) of the genes within the target sorghum genomic region were recovered, as well as five genes before the delimited region and two genes after the delimited region, totaling 58 genes; the BLASTn algorithm (AltschuP et al., 1990) was used to align the CDS against sugarcane leaf transcripts (Cardoso-Silva et al., 2014) with a cutoff of E < 1e-10. Sorghum gene sequences that did not have similar transcripts in the sugarcane leaf transcriptome were compared to those in the SUCEST database (Vettore, 2003) and the NCBI database (AltschuP et al., 1990). Only genes that aligned with sugarcane leaf transcripts, were in the SUCEST database or NCBI database, had putative exons of 200 base pairs (bp) or larger and were not duplicated in the sorghum genome were used for primer development.

#### Identification of BAC clones, sequencing and assembly

To recover the sequences of interest in the equivalent target region of sorghum in sugarcane varieties, BAC libraries from the varieties SP80-3280 and IACSP93-3046 (Sforça et al., 2019) were used. Positive clone selection and preparation of BAC DNA for sequencing and pooling followed the steps described by Mancini et al. (2018). Sequencing was performed on the PacBio® Sequel platform (Pacific Biosciences) at the Arizona Genomics Institute (AGI—Tucson, USA). Vector and *Escherichia coli* genomic sequences were removed with the BBtools package (https://sourceforge.net/projects/bbmap/). Assembly was performed with the Canu v2.1 program (Koren et al., 2017) with default parameters, except for corOutCoverage = 200. The refinement of the final contig consensus sequence was performed by aligning the raw reads against the assembled contigs with the pbalign program, and error correction was performed with the Arrow program. Both programs are present in the SMRTLink v7.0 package (Pacific Biosciences).

#### Annotation of contig sequences

The annotation of BACs for repetitive elements was performed using the LTR FINDER retrotransposon predictor (Xu and Wang, 2007) and the giriREPBASE database (Kohany et al., 2006). Gene annotation was performed with the NCBI (Altschul et al., 1990) and Phytozome v12.0 (Goodstein et al., 2012) databases. The Artemis program of the Sanger Institute (Rutherford et al., 2000) was used to visualize genes and repetitive elements. The sorghum CDSs and the manually annotated sugarcane variety CDSs were used to perform similarity searches using BLASTn tools (Altschul et al., 1990) against the following databases: NCBI (Sayers et al., 2022), UniProt (The UniProt Consortium, 2023) and Pfam (Protein Families) (Mistry et al., 2021). Genes were considered similar if they exhibited a sequence identity of 80% or greater. Contigs that did not have genes, had only one gene or were smaller than 25 kb in size were discarded.

*Manual curation of orthologous regions in* S. spontaneum: Manual homology curation of the orthologous regions in *S. spontaneum* was performed. The CDS of each QTL sorghum gene was aligned against the four alleles of the Sspon02 chromosomal set using BLASTn tools (Altschul et al., 1990). This allowed for enhanced accuracy of the automated annotation performed by Zhang et al., 2018, including the identification of pseudogenes, thereby providing a more precise definition of the genomic architecture in this specific region in *S. spontaneum*.

The information obtained was used for a detailed literature review of each gene. This review describes the proteins and their functions, the biological pathways in which they are supposedly implicated, and their potential role in sugar accumulation in plants, particularly in grasses and sugarcane.

#### Comparative genomic analyses

Comparative analyses were performed between the genes present in the target region in both varieties. In addition, the genes of the orthologous region in the *S. bicolor, S. spontaneum* (Zhang et al., 2018) and the sugarcane hybrid variety R570 (Garsmeur et al., 2018) genomes (Phytozome 12) were also used for comparative analysis. The analyses were performed to determine the synteny, collinearity and genomic structure of the region.

The homologous region in *S. spontaneum* was located using the BLASTn tool (Altschul et al. 1990) against the four homologous chromosome sequences of the *S. spontaneum* homologous chromosome 02 group (Sspon02), thus called Sspon2A, Sspon2B, Sspon2C and Sspon2D (Zhang et al., 2018). Genes were manually curated only in the sorghum orthologous region using the Artemis program from the Sanger Institute (Rutherford et al., 2000) for visualization. Manual curation was performed with NCBI databases (Sayers et al., 2022) and Phytozome 12.0 (Goodstein et al., 2012) databases with visualization through the Artemis program of the Sanger Institute (Rutherford et al., 2000).

### Differential gene expression analysis

The expression of genes within the QTL region was analyzed in internode tissues using sugarcane RNA-Seq data. Gene expression data of the top (3) and bottom (8) internodes of the IACSP93-3046 and SP80-3280 varieties, as well as of the parental species *S. officinarum* (Badila de Java) and *S. spontaneum* (Krakatau), were obtained as described by Aono et al. (2021). Briefly, RNA-Seq reads were trimmed, and gene expression was quantified with Salmon (Patro et al., 2015) using the longest isoforms of *S. spontaneum* CDSs as a reference and automatic annotations by Zhang et al. (2018). A heatmap depicting the expression of all genes within the QTL was generated using the pheatmap R package (Kolde et al., 2012) in R software (R Core Team, 2011).

Differentially expressed genes (DEGs) were identified using the edgeR package version 3.38.4 (Robinson et al., 2010). The raw count data first underwent normalization using the counts per million (CPM) method. Genes with a CPM value ≥ 1 in all samples of at least one biological condition were retained. To identify DEGs, counts were subsequently normalized using the trimmed mean of M-values (TMM) method. Statistical comparisons were conducted between *S. spontaneum* samples and all other samples. DEGs were determined using a false discovery rate (FDR) threshold of p ≤ 0.05 and a log_2_ fold change (FC) cutoff of ≥ 1.

### Gene coexpression network analyses

To further investigate the biological processes associated with the genes within the QTL, a gene coexpression network was constructed with R software employing the highest reciprocal rank (HRR) methodology (Mutwill et al., 2010). Raw count data were normalized using the transcripts per million (TPM) method, and genes with a TPM > 0 in all samples of at least one biological condition were retained. Pairwise Pearson R correlation coefficients were calculated for pairs of filtered genes. To ensure robust associations, a minimum absolute correlation coefficient threshold of 0.8 was used to consider two genes to be connected.

## Results

### QTL gene identification, BAC clone selection, sequencing, assembly and annotation

In the QTL for Brix in sorghum, 51 genes were identified, and seven genes in the expanded region were also identified; of these genes, 21 aligned with sugarcane leaf transcripts and presented exons with sizes equal to or greater than 200 bp. Primer pairs were developed for these 21 genes, and one pair failed to produce amplicons. The 20 primer pairs developed were used for screening clones of interest in the BAC libraries of the SP80-3280 and IACSP93-3046 varieties. Of the remaining 38 genes, 14 were found to be duplicated in the sorghum genome, and 24 did not meet the other selection criteria. For each gene, a number was assigned, except for two tandemly duplicated genes, which were given a single number (25), as shown in Supplementary Table 3, for a total of 57 genes.

In the screening of the BAC library of IACSP93-3046, 37 clones were positive for at least two genes, and 30 clones were sequenced. Among these contigs, 28 were assembled and manually annotated, representing 26 BACs. In the screening of the BAC library of SP80-3280, 56 clones were positive for at least two genes, and 31 clones were sequenced. Of these, genes from the region were found in 16 assembled contigs, and these were manually annotated, representing 16 BACs. The size of all contigs varied between 3,960 bp (pool 25) and 192,924 bp (pool 17), and the total length of the contigs was 5,850,46 bp (Supplementary Table 1).

From the 71 contigs that were generated, 43 carried the target region’s genes. Each contig was related to a BAC, and some BACs were represented by two contigs (Supplementary Table 2).

The sorghum orthologous region was recovered in the variety IACSP93-3046 (Figure 2), which has 50 annotated genes. Seven sorghum genes were not found in the recovered sequence (Supplementary Figure 5). Between genes 35 and 36, there was a gap. In one of the haplotypes, one annotated gene did not belong to this region in sorghum, although it is located in another region of sorghum chromosome SBI-2. In the SP80-3280 variety (Figure 1), the region was recovered almost in its entirety, with 44 annotated genes. Of the seven genes that were not found in the IACSP93-3046 contig sequences, six were not found in this variety, and one was annotated as a pseudogene. It is possible to observe two gaps, one between genes 24 and 26 and the other between genes 43 and 47.

**Figure 1:**
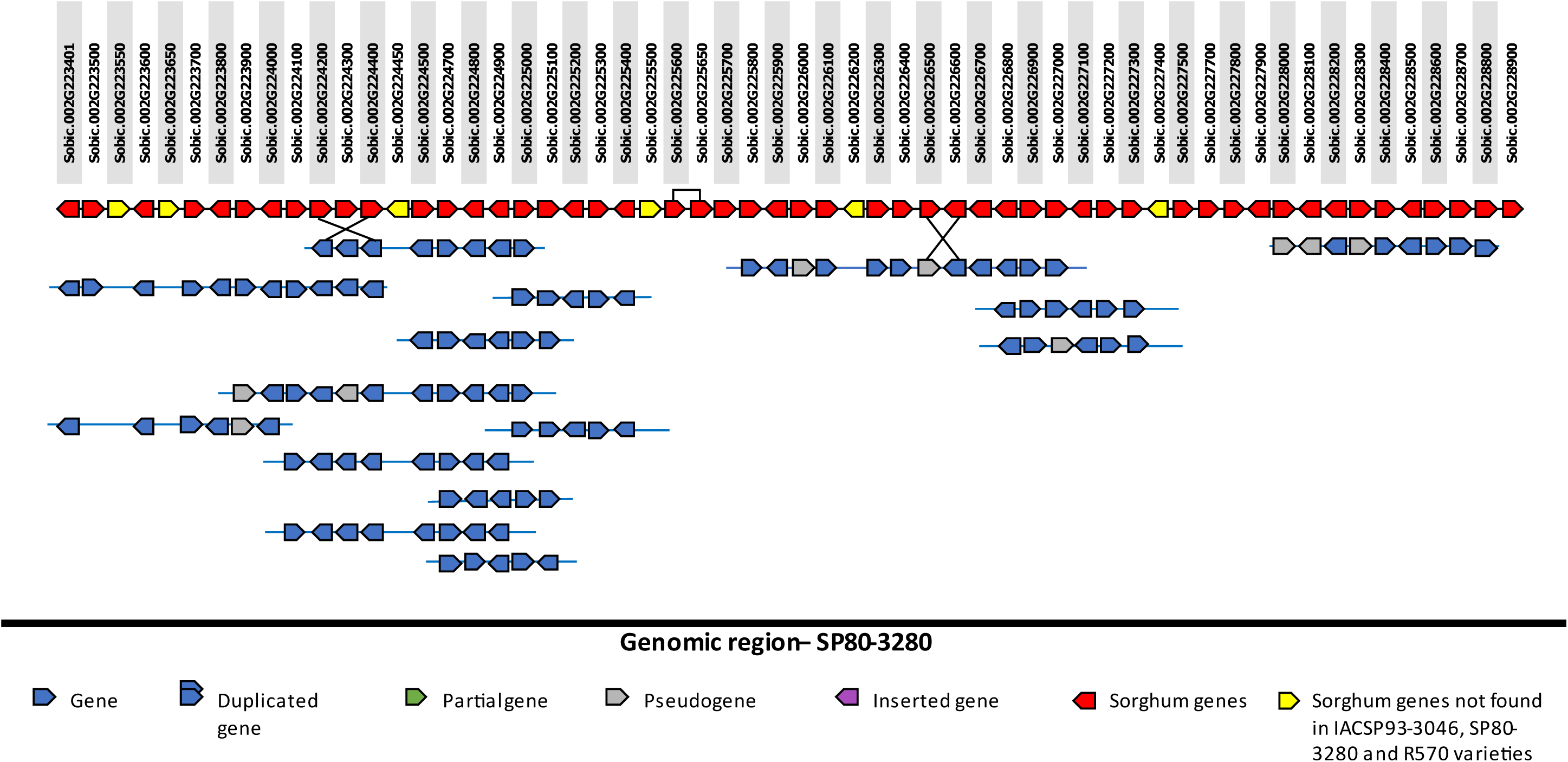
Genomic region of sorghum×Genomic region of SP80-3280. Each square represents a gene and shows the direction of the gene in the genome, where right-facing arrows indicate the forward direction, and left-facing arrows indicate the reverse direction. The solid lines with squares in red and yellow represent the 57 sorghum QTLs. The genes in red are orthologous sorghum genes, while genes in yellow lack orthologs in the hybrid varieties. For each gene, a number has been designated, and above each number, the name allocated to it in Phytozome v.13 is indicated. Each solid line below the genomic region of sorghum represents a BAC. Genes shown with solid lines represent those successfully recovered within a BAC; together, the BACs reconstruct the genomic region of SP80-3280.

**Figure 2:**
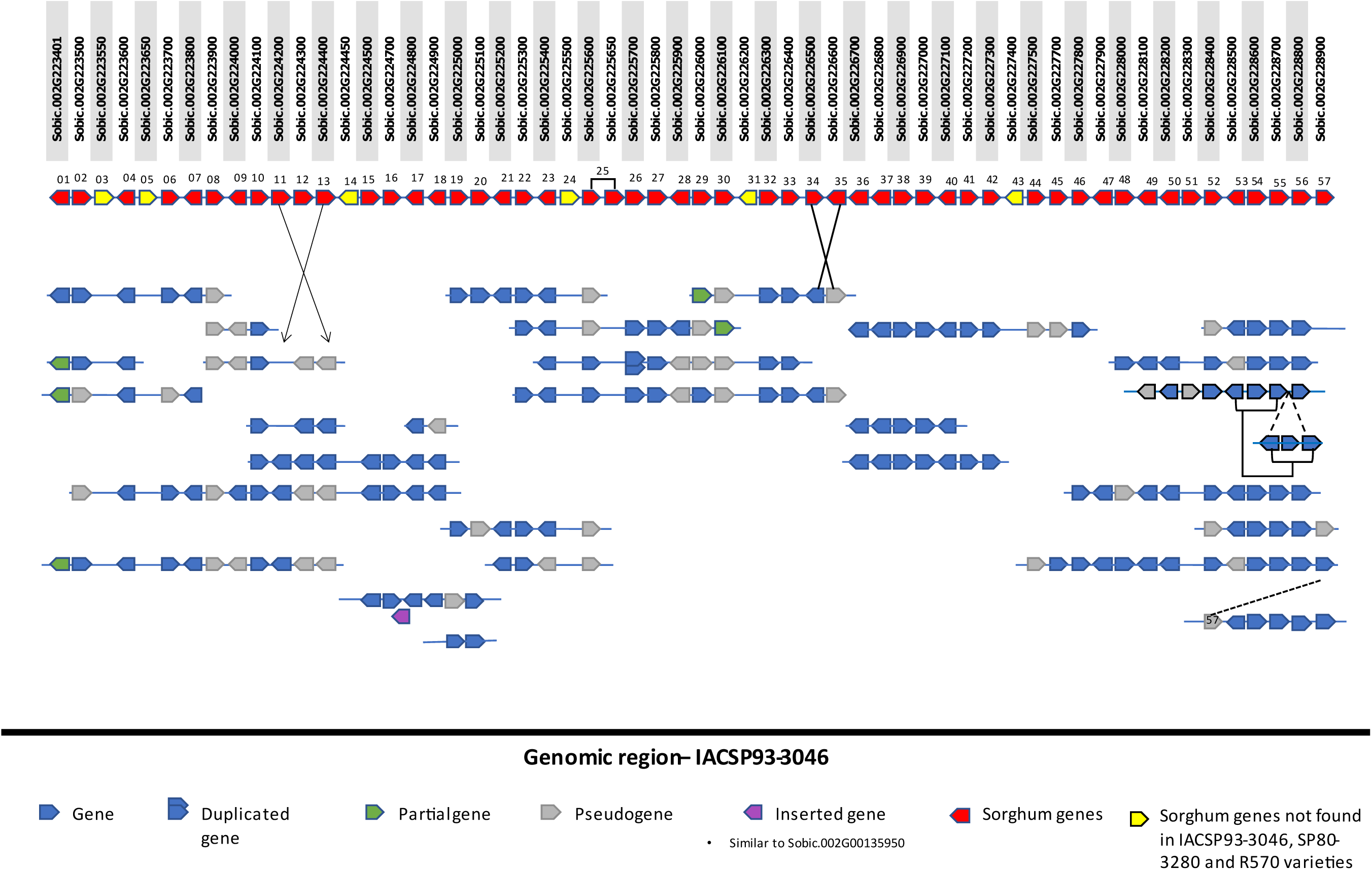
Sorghum×IACSP93-3046 genomic region. Each square represents a gene and shows the direction of the gene in the genome, where right-facing arrows indicate the forward direction, and left-facing arrows indicate the reverse direction. The solid lines with squares in red and yellow represent the 57 sorghum QTLs. The genes in red are orthologous sorghum genes, while genes in yellow lack orthologs in the hybrid varieties. For each gene, a number has been designated, and above each one, the name allocated to it in Phytozome v.13 is indicated. Each solid line below the genomic region of sorghum represents a BAC. Genes shown with solid lines represent those successfully recovered within a BAC; together, the BACs reconstruct the genomic region of IACSP93-3046. The gene depicted in purple is an exclusive finding within this BAC. This gene is not one of the 57 sorghum QTL genes; however, it is similar to the sorghum gene Sobic.002G00135950 (Phytozome v.13). A pseudogene orthologous to this gene was observed on chromosome Sspon.2B (Supplementary Figure 2).

There were 45 pseudogenes among the homo(e)logous genes in the variety IACSP93-3046 and nine probable pseudogenes in the variety SP80-3280. In the variety IACSP93-3046, genes with insertions of transposons in intronic regions (6-13.3%), insertions/deletions of one or more nucleotides (36–80%) and partial gene sequences (3-6.7%) were considered pseudogenes. Among nine homo(e)logous genes considered probable pseudogenes in SP80-3280, four (44.5%) exhibited an insertion/deletion of one or more nucleotides, three (33.5%) exhibited a transposon insertion in intronic regions, and in two (22.2%) of these genes, the pseudogene was a fragment of the gene.

#### Main differences in sorghum-sugarcane synteny and collinearity in the target region

##### Chromosome Sspon2A (Supplementary Figure 1)

The orthologous region on chromosome Sspon2A is 794,054 bp long and is the closest in size to chromosome SbI-02 of sorghum. It is located between bases 35,019,101 and 35,813,155. Among the 57 genes present in sorghum, 50 orthologs were found in Sspon2A. The seven missing orthologous genes (03, 14, 24, 40, 46, 55 and 56) were not detected throughout the chromosome and not only in the delimited region; they were not detected in the IACSP93-3046, SP80-3280 and R570 varieties. In the region delimited in Sspon2A, the gene Sspon.02G0013290 was found, and it is orthologous to a sorghum gene from chromosome Sb10 (Sobic.010G093001). Three additional genes were not detected in the IACSP93-3046 and R570 varieties, 31 and 43; these genes were detected on chromosome Sspon2A but as pseudogenes. Gene 51 was also detected as a pseudogene in the SP80-3280 variety but was the IACSP93-3046 and R570 varieties. Eight inversions were observed, two of which were common to the varieties IACSP93-3046 and SP80-3280, and they involved from two to eight genes. Duplications, some *in tandem*, were also observed. Therefore, there is synteny, as almost all the genes are present, but there are many breaks in collinearity.

##### Chromosome Sspon2B (Supplementary Figure 2)

On this chromosome, between the first and last genes of the studied region, there are 1,134,230 bp, more than double the region in sorghum, located between bases 32,312,667 and 33,446,897. In this chromosome, it was possible to observe many collinearity breaks, with inversions and an insertion within a cluster of 12 genes. In this orthologous region, it was possible to observe rearrangements and reorganizations, but most of the genes were present, guaranteeing synteny. Of the 57 genes present in sorghum, six were absent from the entire chromosome: 07, 26, 27 and 28. Genes 14 and 31 were also missing and were also not found in the IACSP93-3046, SP80-3280 and R570 varieties. An insertion with an eight-gene cluster, similar to a region immediately posterior to the one studied, in sorghum is present in this allele.

##### Chromosome Sspon2C (Supplementary Figure 3)

The chromosome Sspon2C region is the region that most resembles the sorghum chromosome SbI-02. Considering synteny, although between the first and the last gene, it is almost twice the size of the region, reaching 969,275 bp, and is located between bases 37,413,201 and 38,382,476. As in the other alleles, Sspon2C also has collinearity breaks with inversions and insertions, and there are gene sequences from the region that are displaced and inserted in other stretches. Among the 57 sorghum genes in the region, there are two that are absent on this chromosome, and these genes are also absent in the hybrid varieties IACSP93-3046, SP80-3280 and R570: genes 14 and 31. Although the region is quite large, compared to sorghum, there are no insertions with genes similar to those of other chromosomes in *S. spontaneum*.

##### Chromosome Sspon2D (Supplementary Figure 4)

This chromosome also maintains synteny with sorghum. Of the 57 genes in the region, 51 remained. Among the six missing genes, four were not detected in the IACSP93-3046, SP80-3280 or R570 varieties. The region was divided into two subregions. The first subregion is between bases 28,425,371 and 29,097,118 (671,747 bp), and the second is between bases 49,915,069 and 50,120,906 (205,837 bp). These two regions are approximately 21 Mb in length.

##### SP80-3280 variety (Figure 1)

Of the sequenced 31 clones, 16 were recognized as part of the target region using BLASTn. The orthologous region was partially recovered using region-belonging BACs. Among the recovered genes, synteny and collinearity may be presumed. Although there are breaks in collinearity with the variety IACSP93-3046, some of those observed are the same in both varieties, such as an inversion between genes 11 and 13 and another between genes 34 and 35. Out of 15 sequenced and annotated BACs, 11 were positive for the Sobic.002G223900 gene (gene 08), and of these, eight were also positive for one of the last 20 genes in the target region (genes 38 to 57). Some genes were not observed in the annotations of the varieties IACSP93-3046 and R570; this also occurred with the variety SP80-3280, except for one gene (51) that was observed as a possible pseudogene. Two pronounced gaps were detected: the absence of BACs containing genes 24 to 26 and the absence of BACs containing genes 43 to 47. The last gene flanking the region, 57, was also not recovered.

##### IACSP93-3046 (Figure 2)

This region was recovered with 29 annotated BACs. The synteny between sorghum and sugarcane in this specific region was confirmed, but some collinearity breaks were detected. Two inversions were observed, one between genes 11 and 13 and the other between genes 34 and 35. These inversions are observed in all haplotypes where these genes could be present. Gene 25, which is duplicated in sorghum, appeared in a single copy in the annotated haplotypes; on the other hand, gene 26 was duplicated in one of the three haplotypes observed. A sequence of three genes (53, 54 and 55) was found to be duplicated exactly in this sequence, resulting in a collinearity break; however, this finding appears in only one of the seven haplotypes that could have these genes. In one annotated BAC, an insertion of gene 57 between genes 52 and 53 was observed. Another interesting insertion was found between genes 16 and 17; it was a gene similar to Sobic.002G135950 from sorghum, and chromosome SbI-02 at positions 20,517,004-20,519,005, and Sobic.002G195033 from sorghum was located at sites Sb02 58,315,718-58,317,644; in other words, this gene was in another region but on the same chromosome.

##### R570 (Supplementary Figure 5)

Upon comparing the findings of Brazilian varieties with those of R570, some commonalities were observed. The inversion between genes 11 and 13 is present in all three hybrid cultivars, indicating that this observation is a characteristic of the *Saccharum* genus, as is also observed in *S. spontaneum*. Tandem duplications, such as that of gene 25, were noted, mirroring observations in sorghum. Interestingly, IACSP93-3046 lacks this duplication, and due to a gap in the sequencing of this region, this duplication could not be detected in SP80-3280. In R570, genes 48 and 49 are duplicated in tandem, a feature not observed in sorghum. Similar findings were not observed in *S. spontaneum* or in the varieties SP80-3280 and IACSP93-3046.

### Expression analysis and search for candidate genes related to sugar accumulation: Investigation of selected genes

A summary of genes 01 to 57, their orthologs in *S. spontaneum* and *S. bicolor,* as well as their proteins, is provided in Supplementary Table 03. Based on this analysis, 10 candidate genes for sugar accumulation were selected: 02, 06, 09, 10, 15, 19, 20, 23, 25, and 43. These genes are possibly involved directly, indirectly, or in fundamental upstream steps involved in some phase of the process of sugar accumulation, which begins with carbon fixation from the atmosphere (photosynthesis), sucrose biosynthesis and transport to the stems, and subsequent accumulation (Supplementary Table 03).

### DEG analyses

The expression of the genes within the QTL was evaluated using RNA-Seq data from internodes 3 (younger) and 8 (more mature) of IACSP93-3046 and SP80-3280, as well as data from accessions of the two species considered the main ancestors of modern cultivars, namely, *S. spontaneum* and *S. officinarum*. Seven of the 51 sorghum genes under analysis had no orthologs in the *S. spontaneum* genome, which was used for the gene quantification procedures; therefore, they are not represented in the expression data.

A heatmap (Figure 3) depicting the expression of the remaining 44 genes normalized by TPM is shown in Figure 3. This allowed us to observe internode gene expression patterns in two commercial sugarcane varieties, IACSP93-3046 and SP80-3280, and the parental species *S. officinarum* and *S. spontaneum*. For ten genes (10, 11, 12, 13, 15, 22, 25, 34, 45, 46, and 56), no expression was detected in any biological replicate, or there was minimal expression in up to three biological replicates. These genes may play crucial roles in other plant organs, such as leaves or roots, or could also be relevant in other stages of plant maturation. However, due to a lack of evidence of expression in the organ/tissue and maturation stages under analysis, these genes were not considered candidates for involvement in sugar production.

**Figure 3:**
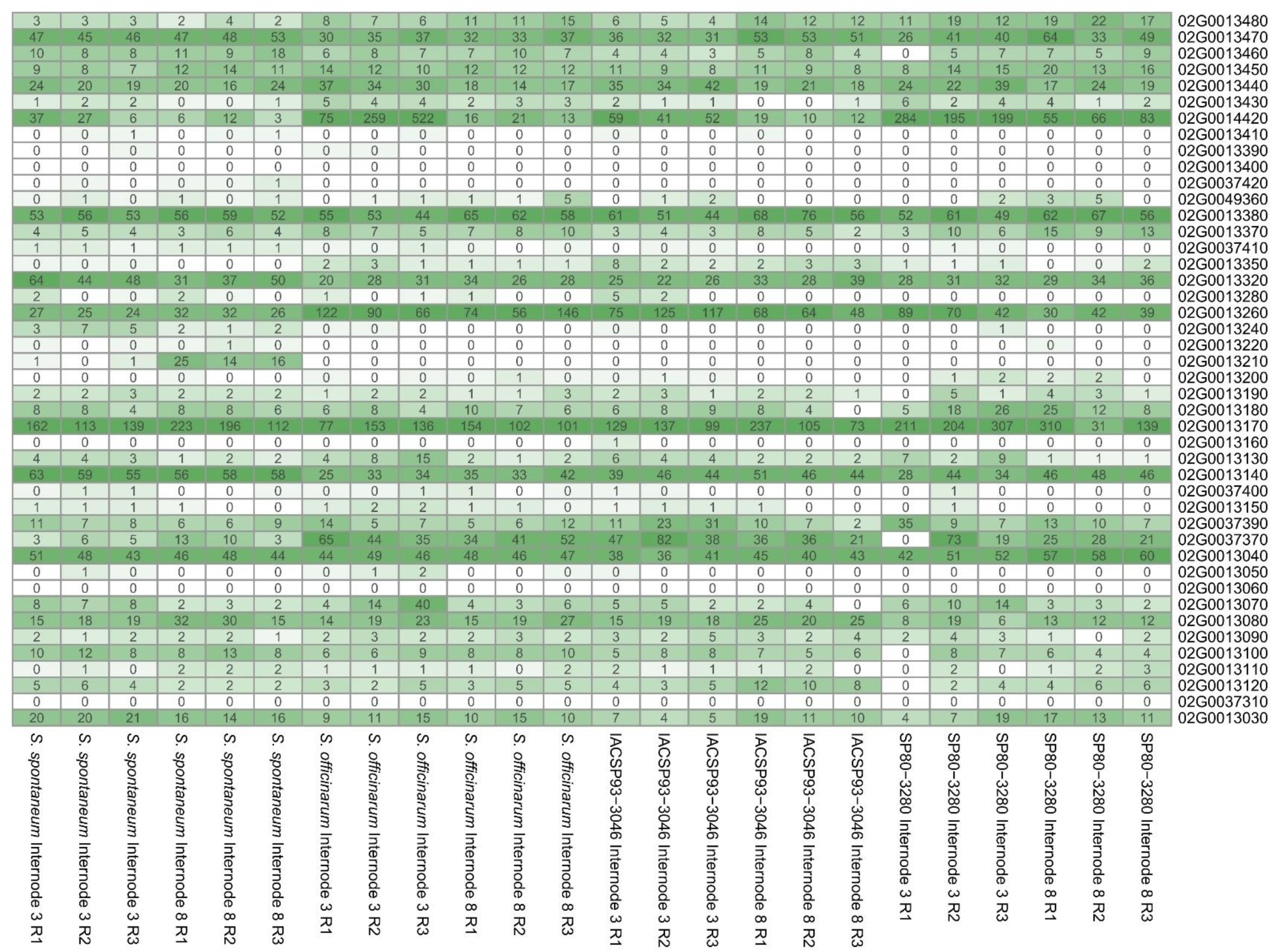
Heatmap representing the level of gene expression in the internodes of SP80-3280, IACSP93-3046, *S. officinarum* and *S. spontaneum.* A heatmap depicting gene expression levels across tissues, internodes 3 (top) and 8 (bottom) of sugarcane plants (varieties SP80-3280 and IACSP93-3046; *S. officinarum* and *S. spontaneum*) in triplicate. The darker the shade of green is, the higher the expression level.

After filtering, 22,859 of the 35,471 genes present in the *S. spontaneum* CDSs were stably expressed under at least one biological condition and were thus retained for DEG analyses. By comparing varieties with high (*S. officinarum*, IACSP93-3046, and SP80-3280) and low (*S. spontaneum*) sugar contents, 6,264 DEGs were identified (Supplementary Table 4). Seven genes within the QTL region were DEGs; their log_2_(FC) values, FDR-corrected p values and annotations are available in Table 1.

**Table 1:**
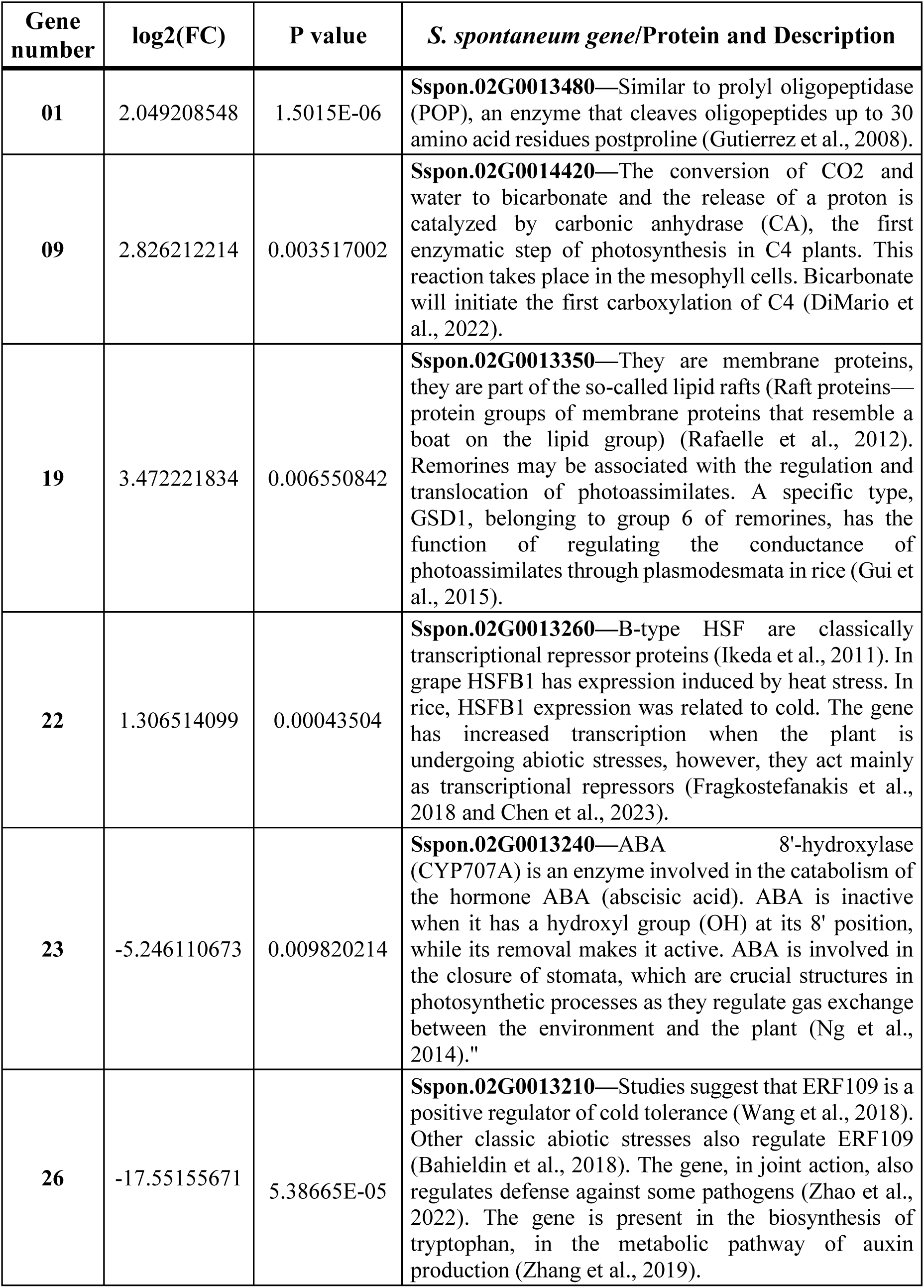
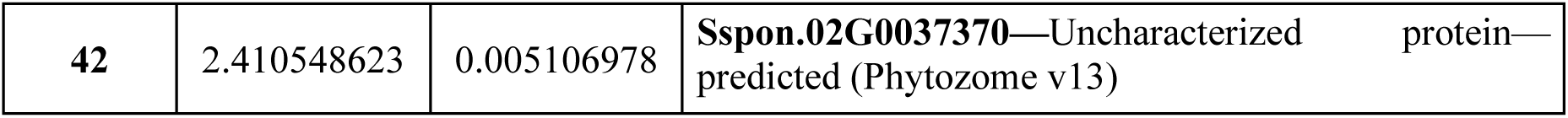
Differentially expressed genes (DEGs) located within the QTL under analysis.

### Gene coexpression network analyses

Based on the expression data, an HRR coexpression network was constructed to explore new evidence that could contribute to the search for candidate genes. During filtering procedures, 7,565 genes were excluded, and the remaining 27,906 genes were used as input to construct the network. The final network had 6,809 connected nodes (genes) and an average of 17 neighbors per node. Among these genes, 3,397 genes were identified as DEGs, and six genes were identified within the QTL. A first neighbor search was employed to identify genes related to potential sugar accumulation candidates and to assess whether these genes could support their role in this process. The first neighbors of the genes within the QTL represented in the network can be seen in Table 2. Three of these genes—01, 23, and 26—were also identified as DEGs (Table 1).

**Table 2:**
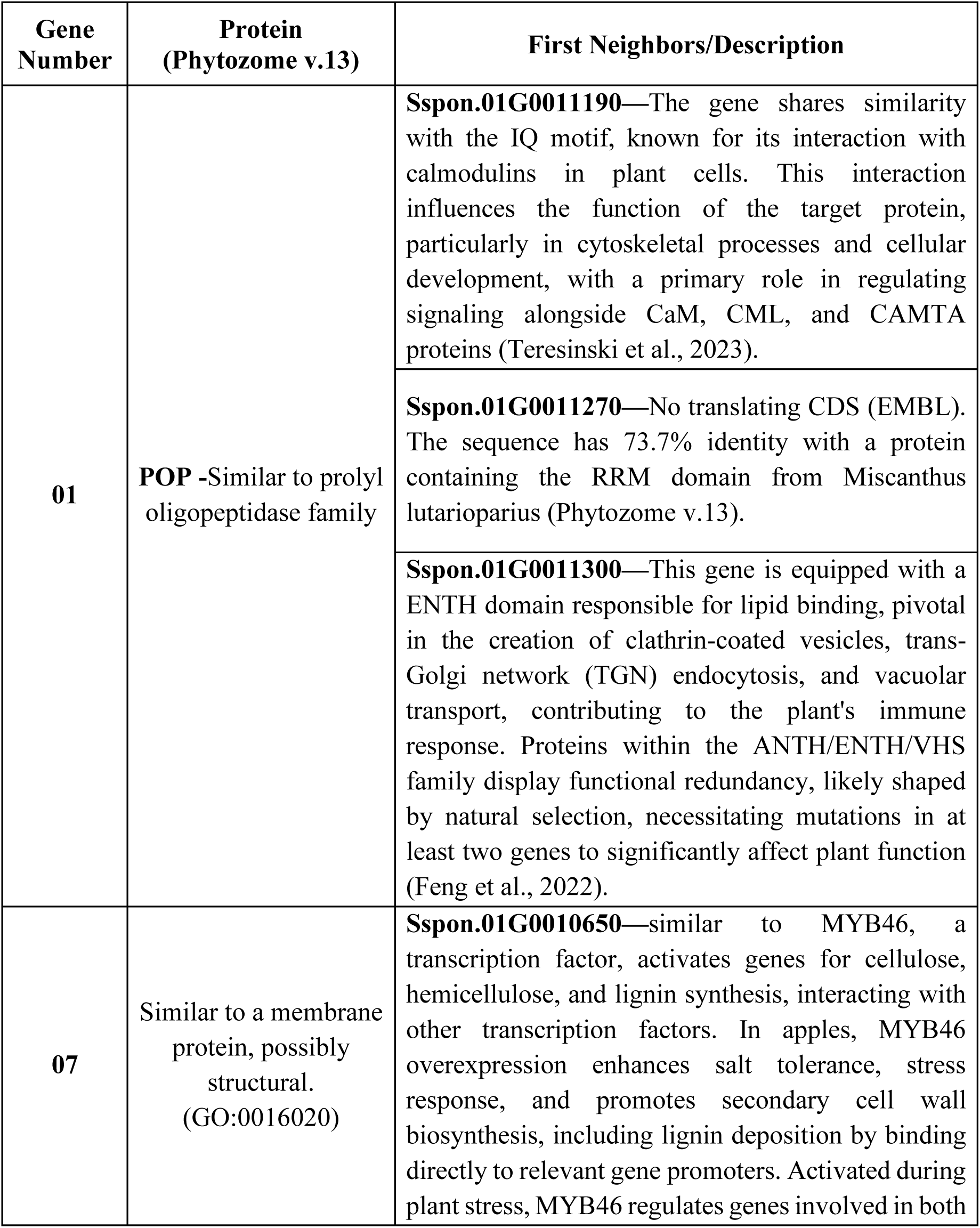
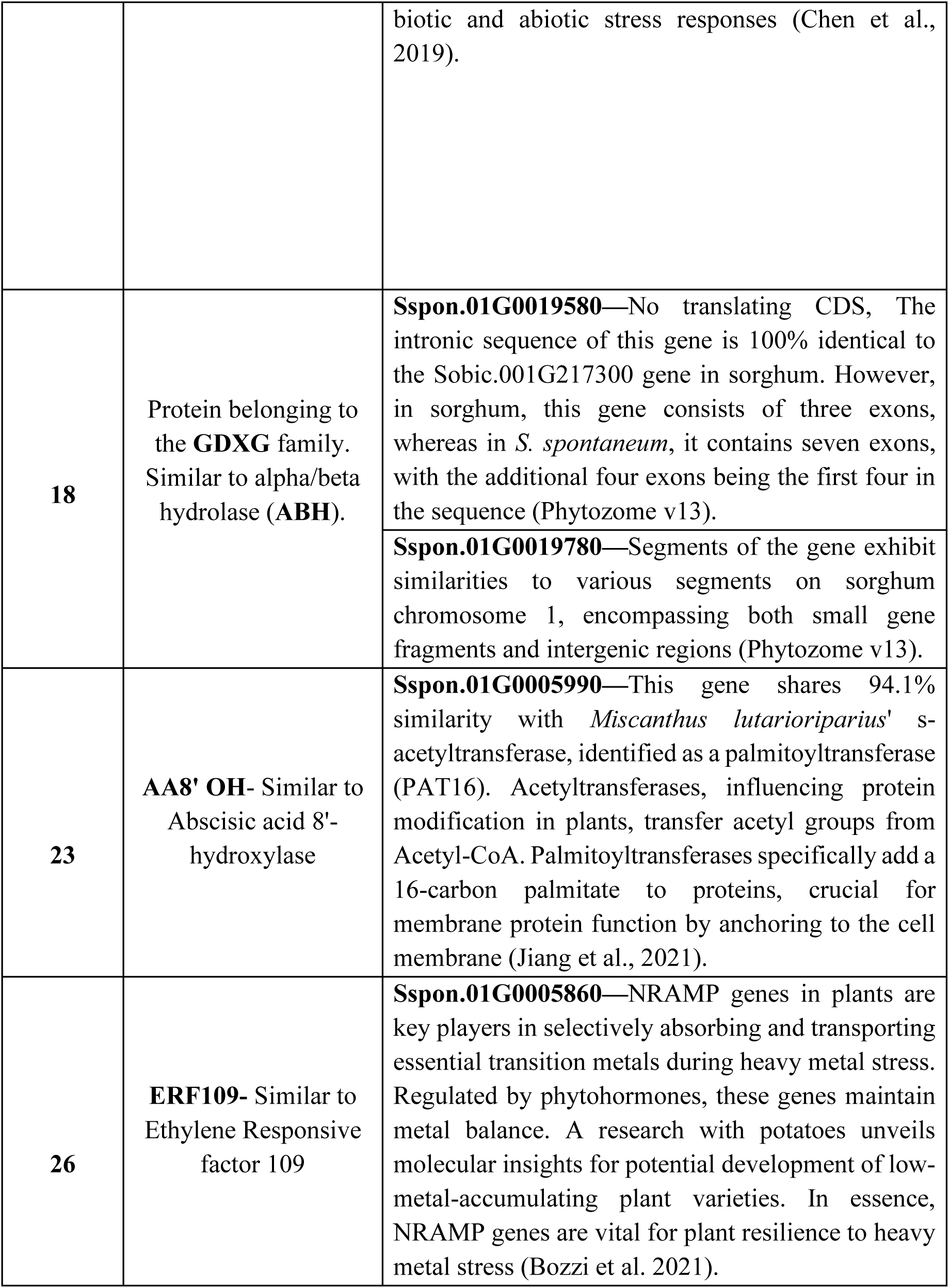
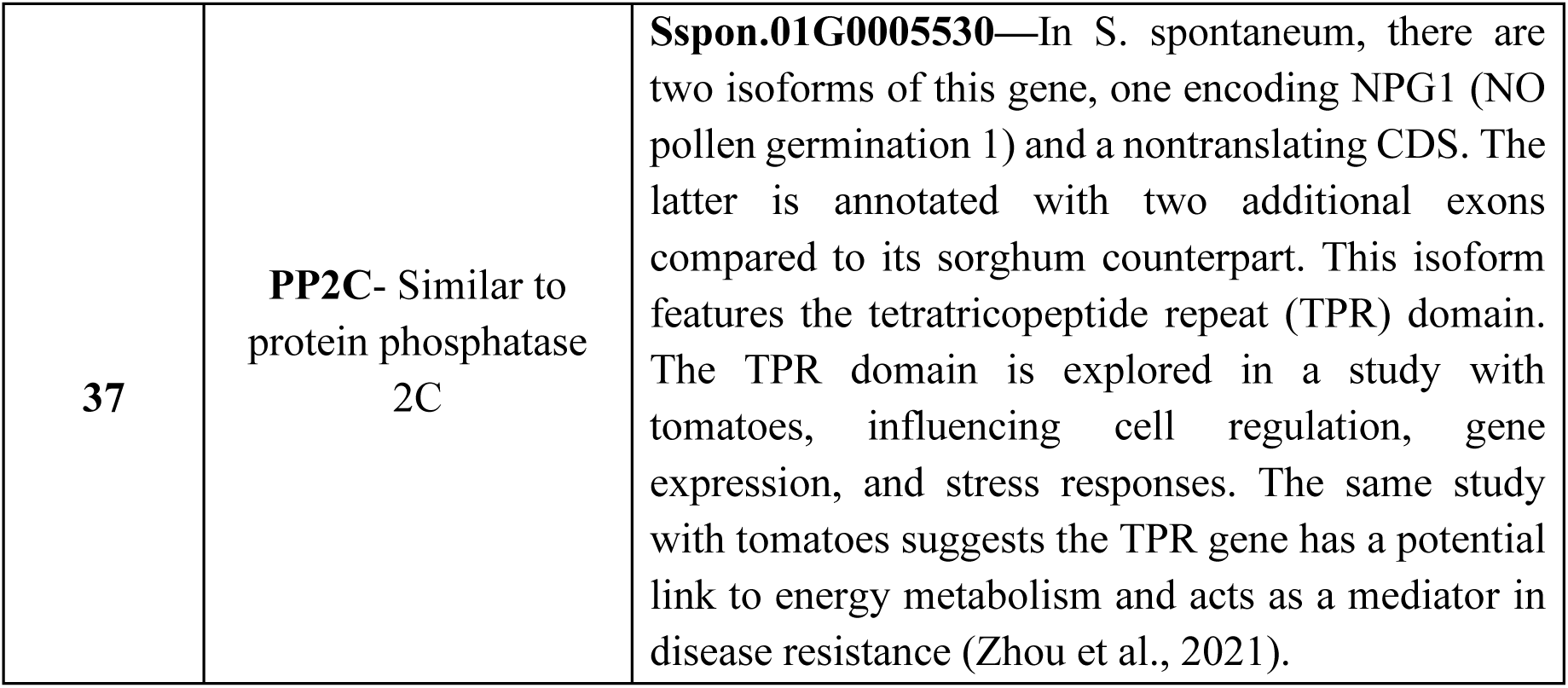
Sugarcane genes and their first neighbors. The first neighbor genes are defined by their names in *S. spontaneum*, and their descriptions are based on the annotations of the EMBL database.

Gene 01 (prolyl oligopeptidase—POP) exhibited relatively low expression in the stems of *S. spontaneum* and relatively high expression in samples from sugar-accumulating plants. Gene 23 (abscisic acid 8’-hydroxylase 3—ABA8’OH) has virtually no expression in the internodes of the sugar-accumulating plants sampled and is expressed at low levels in *S. spontaneum.* Gene 26 (ethylene responsive factor 109—ERF109) also has almost no expression in sugarcane plants, while it is expressed in *S. spontaneum*; however, in this case, there is significantly greater expression in the more mature internodes of *S. spontaneum* (I8) than in less mature internodes. This evidence led to the selection of genes 01, 23, and 26 as the primary candidate genes in the QTL for sugar accumulation.

## Discussion

### Main differences in genomic architecture

Synteny and collinearity have been used to compare and recover genomic regions of interest in sugarcane using sorghum (Ming et al., 1998; Paterson et al., 2009; Wang et al., 2010; Figueira et al., 2012; Mancini et al., 2018; Garsmeur et al., 2018; Thirugnanasambandam et al., 2018; Zhang et al., 2018; Sforça et al., 2019; Aono et al., 2021; Federico et al., 2022 and Healey et al., 2024) and miscanthus (Mitros et al., 2020 and Zhang et al., 2021) genomes as references, revealing high gene retention (Mancini et al., 2018; Garsmeur et al., 2018; Sforça et al., 2019 and Feng et al., 2021). The comparison of the same region between sugarcane varieties and their ancestral species can provide insight into the genomic complexity of sugarcane. The region evaluated in this work showed substantial differences among the genotypes studied, such as gene duplications, loss of gene exons, pseudogenization, gene inversions, gene deletions and insertions.

For example, gene 25 (similar to alpha-amylase—AMY—Supplementary Table 3) is duplicated in tandem in sorghum and in R570, but in four IACSP93-3046 haplotypes, it is in a single copy (there is a gap in the region SP80-3280). The sequences of genes 53, 54, and 55 (Supplementary Table 3) were duplicated in tandem in BAC Shy141H03 of IACSP93-3046 (Figure 2), but they were not duplicated in sorghum, neither in the recovered SP80-3280 haplotypes nor in any of the alleles of the *S. spontaneum* genome. In addition to gene duplications, gene inversions were detected in the orthologous region between sorghum and all the *Saccharum* accessions evaluated (Figure 1), which suggests that both inversions occurred after sorghum–sugarcane divergence. Overall, when inversions do not significantly disrupt the gene balance of an organism, the direct consequences tend to be minimal. Documented cases exist where inversion results in pseudogenization or even deletion of one of the genes (Jurka et al., 2001; Zhao et al., 2016; Redd et al., 2023).

In BAC Shy411A07 of the IACSP93-3046 variety, gene 11 (Supplementary Table 3) was absent, yet the remaining genes (12 and 13—Supplementary Table 3) indicated that an inversion occurred. Fragments of Harbinger-type (HARB) repetitive elements were found near pseudogenes 12 and 13 (Supplementary Table 3). HARB transposons are classified as class II transposable elements (TEs) that carry out the cleavage and transfer of single DNA strands mediated by transposases (Zhao et al., 2016; Redd et al., 2023). The presence of these HARB transposons suggests a possible relationship between these elements and these inversions, which were present in all examined varieties, especially with the probable pseudogenization of genes 12 and 13 (Supplementary Table 3). The process of cleavage followed by fusion may have led to the deletion of bases, resulting in the truncation of genes and, consequently, the loss of their functions. In the SP80-3280 variety, gene 13 (Supplementary Table 3) exhibited a single exon spanning 2,682 base pairs. On the other hand, gene 12 (Supplementary Table 3) maintains two introns, even in its pseudogenized state, and in this case, it is situated between two TEs, similar to the HARB type (BAC Shy260G24). In BAC Shy492F12, gene 12 (Supplementary Table 3) is also close to a HARB-type TE flanking the last exon. In this case, gene 12 (Supplementary Table 3) exhibited characteristics indicative of a functional gene. The gene had different CDS base pair compositions among the haplotypes but was always between 1347 bp and 1488 bp. Additionally, gene 11 (Supplementary Table 3) retained a single intron, with a length ranging between 1,200 and 1,209 bp. However, neither variation was detected in sorghum, suggesting that it might be a unique characteristic of the *Saccharum* genus. On chromosome Sspon2B of *S. spontaneum*, gene 11 (Supplementary Table 3) has a single exon.

In a haplotype of the variety IACSP93-3046, represented by BAC Shy112C03, a gene whose ortholog in sorghum is not found in the QTL studied was detected. Notably, this gene is similar to the two sorghum genes Sobic.002G135950 and Sobic.002G195033. These sorghum genes have 91% sequence identity, and both have a zinc finger domain. The probable orthologous gene in the IACSP93-3046 variety is inserted in a retrotransposon similar to Copia22-ZM_I/LTR. Interestingly, this gene exhibits all the characteristic features of being a functional gene, even though it is inserted in a TE. One possible explanation for this insertion is that the gene was cotransported with the retrotransposon. As Class II TEs, they can replicate a copy of themselves, which is subsequently inserted into different genomic regions. As such, there is a substantial likelihood that this haplotype is a copy of the gene. The presence of a TE within an expressed gene (CENP-C) in sugarcane has been previously described (Sforça et al., 2019), demonstrating that the proximity or overlap of TEs and genes does not hinder the function of the gene, at least in sugarcane. This gene, specific to the IACSP93-3046 haplotype (BAC Shy112C03), has a zinc finger domain; in plants, proteins featuring this domain are transcription factors (TFs) related to the control of cell division in totipotent tissues (petunias), histone-DNA binding (wheat), leaf budding (Chinese cabbage), soil salinity tolerance (Arabidopsis) and carbon metabolism (potato) (Takatsuji, 1999).

This gene has also been detected in the orthologous region of the *S. spontaneum* chromosome Sspon2B but as a gene fragment. In IACSP93-3046, the gene is located between genes 16 and 17 (Supplementary Table 3), in reverse orientation, and in Sspon2B, it is located between genes 39 and 41 (Supplementary Table 3), with gene 40 (Supplementary Table 3) being inserted into another fragment of the orthologous region, in strand orientation. It is possible that this gene could also have been transposed with a TE, as possibly occurred with the hybrid. Importantly, *S. spontaneum* is a wild species that has been evolving under the pressure of natural selection, without the same level of human interference that fully domesticated plants undergo, as is the case with modern sugarcane cultivars—commercial hybrids. Despite such variability, we can observe the presence of potential genomic structure characteristics of *S. spontaneum* in commercial hybrids, such as inversions 11-13 (Supplementary Table 3) and 34-35 (Supplementary Table 3), which are present in at least three of the four alleles and are also present in all recovered haplotypes of the SP80-3280 and IACSP94-3046 varieties, where these inversions could be observed, as well as in the R570 variety (Garsmeur et al., 2018).

The orthologous regions in the hybrid varieties appear to be more similar to those in sorghum than to those in *S. spontaneum*. Some fundamental characteristics are shared, such as synteny. However, differences such as inversions, duplications, insertions of orthologous genes from the same genomic region and even from sequences that are similar to genes from sorghum chromosomes other than SBI-02, possible pseudogenization and translocation were detected. However, this finding is not surprising considering that the chromosomes originating from *S. spontaneum* found in hybrids constitute only 10% to 20% of the chromosomes of modern hybrids (D’Hont et al., 1996; Cuadrado, 2004; Piperidis et al., 2010; Garsmeur et al., 2018). Furthermore, wild species such as *S. spontaneum* have been subjected to natural selection pressure, resulting in a high level of expected heterozygosity for wild plants. While wild species have evolved naturally, commercial varieties have been selected and improved over the past 120 years to meet human needs (Singh et al., 2020), which has led to substantial genomic differences between them.

### Investigation of genes involved in sugar accumulation

Of the 51 studied genes, 17 were annotated as genes associated with tolerance or response to stress (genes 01, 04, 11, 12, 13, 20, 22, 23, 24, 26, 32, 33, 34, 36, 37, 41, and 57—Supplementary Table 3). The period when sugarcane accumulates sucrose in its stems coincides with the dry and high luminosity period in a significant portion of the crops. During this period, leaves gradually fall, while sugar accumulates in culms (Garcia et al., 2019). Sucrose accumulation occurs in response to stressful conditions (Souza et al., 2018). Therefore, there may be a connection between sucrose accumulation in sugarcane and genes related to abiotic stress.

Considering the characteristics of the crop, the significant number of genes related to the abscisic acid (ABA) response were also observed (genes 20, 23, 32, 33 and 37— Supplementary Table 3). In addition to being a crucial hormone for the photosynthetic process by regulating stomatal closure and opening, ABA is highly responsive to stress, particularly water stress (Chen et al., 2019). Leaf water potential and stomatal conductance are crucial factors for sugarcane to be able to produce carbohydrates that are converted into sucrose, transported to stalks, and subsequently accumulate (Smit et al., 2006 and Aluko et al., 2021).

In addition to these genes, five genes (genes 17, 18, 45, 46, and 47— Supplementary Table 3) belonging to the lipolytic enzyme GDXG family were detected. This enzyme family is characterized by having two consensus sequences containing a histidine residue and a serine residue as putative active site residues (van der Vlugt-Bergmans et al., 2001). In some genes encoding these enzymes, the presence of the alpha/beta hydrolase (ABH) domain may occur, which is known to play a role in catalyzing the cleavage of carbon double bonds and decarboxylation. Additionally, six genes (8, 49, 50, 52, 53, and 54—Supplementary Table 3) containing the ABH domain but belonging to the carboxylesterase (CXE) family were identified, five of which are sequential. The specificities of genes within the same family characterized by shared domains may vary significantly, necessitating further exploration of the biological roles of each gene. Notably, a group of genes sharing closely related domains and families on the same chromosome, even sequentially, may suggest a potential origin through duplication events that diverged during evolution into distinct genes while maintaining some similarity (Zhang et al., 2003).

### Candidate genes for sugar accumulation

The ERF109 (gene 26—Supplementary Table 3), ABA 8’ OH (gene 23— Supplementary Table 3), and POP (gene 01—Supplementary Table 3) genes are candidates for sucrose accumulation in sugarcane, considering that sugarcane needs soil with low humidity, approximately 15%, for greater sugar accumulation (FAO, 2024).

Gene 26 (Supplementary Table 3) is similar to ERF109, an ethylene-responsive TF. The expression of ERF109 is related to anthocyanin accumulation in apples, as ERF109 directly binds to the promoters of anthocyanin synthesis genes (Ma et al., 2021). Jasmonic acid (JA) accumulation in plant wounds also activates the expression of ERF109. ERF109 induces the biosynthesis of the auxin protein ASA1, which aids in the process of secondary root formation mediated by JA-dependent ERF109 signaling (Guarneri et al., 2023). In sugarcane, more than 16,000 genes have been identified as potential targets regulated by ERF109, indicating that ERF109 has a broad influence on gene expression. Functions are diverse and include metabolic activities such as Rubisco activity, triggering hormone biosynthesis such as that of cytokinins, and gibberellin-mediated responses (Yu et al., 2024). ERF109 was not expressed in the internodes of any of the analyzed sugar-accumulating plants or in the younger internodes of *S. spontaneum*, but there was significant expression in its older internodes. In transgenic lemon, the overexpression of ERF109 causes global reprogramming of plant expression. ERF109 acts as a stress-responsive TF, but theorizing that this gene is a candidate gene for sugar accumulation requires further investigation.

### Proline-dependent genes

Genes 01 and 23 (Supplementary Table 3) are related to proline accumulation. Proline is an amino acid with a unique configuration, restricting its free rotation at the α-carbon because the nitrogen and α-carbon are combined in a pyrrolidine ring. This structure contributes to the rigidity of proteins containing proline residues and requires specific enzymes, including POP, for cleavage (Dong et al., 2017). Enzymes that specifically cleave proline are known to be involved in proline accumulation in the cytosol of plant cells. This accumulation is essential for the plant’s adaptive response to adverse situations (Ghiffari et al., 2022). Plants accumulate proline to maintain cellular homeostasis, aid in water absorption, and better adapt to abiotic stresses such as drought, salinity, and heavy metals. These adverse conditions lead to excess production of reactive oxygen species (ROS) and consequences such as lipid peroxidation, increased osmolyte levels, and activation of antioxidant systems (Rejeb et al., 2014; Kazemi-Shahandashti & Maali-Amiri, 2018; Sharma et al., 2019; Ghosh et al., 2021). As such, proline accumulation enhances adaptive responses in plants. Plants may increase proline biosynthesis in response to the above conditions or reuse presynthesized proline from proteins and peptides that are not essential (Ghifari et al., 2022).

The exogenous application of proline in maize has been reported to increase sugar, oil, moisture, and protein levels in seeds under drought conditions (Ali et al., 2013; Gosh et al., 2021). In sugarcane, the efficiency of photosynthesis and stomatal conductance is especially related to sucrose accumulation in stalks (Singels et al., 2021). In mature plants, the relationship between leaves as a sugar source and other organs, including internodes, is critical for the regulation of photosynthesis rates and sucrose accumulation in stalks (Souza et al., 2018). Proline interacts with other metabolites, including soluble sugars. The phenomenon of proline accumulation interacting with sucrose, for example, to adjust osmotic balance during salt stress, has been reported (Ghosh et al., 2019). However, it is unclear whether this interaction could be driven by other physiological changes in different plants with different stimuli. Although the connection between stress and plants has been clarified, there is still much to be elucidated. In sugarcane, the accumulation of sucrose and starch in leaves coincides with a reduction in photosynthetic rates, which occurs during low water availability (Garcia et al., 2019).

Gene 23 (Supplementary Table 3), which shares similarities with ABA 8’-hydroxylase enzymes from the cytochrome-P450 family, converts ABA into 8’-hydroxy ABA and then into phaseic acid (Kronchko et al., 1998), regulating ABA metabolism and influencing plant responses to environmental stress and development, including germination, root growth, and fruit maturation (Wang et al., 2023). Inhibition of this enzyme affects the balance of processes involving ABA (Wang et al., 2023), such as stomatal closure in response to water, salt, and thermal stresses. Studies in grapes have shown that inhibiting ABA 8’-hydroxylase results in reduced leaf water potential and stomatal conductance (Tomiyama et al., 2020), accompanied by proline accumulation in leaves and the growth of adventitious roots (Tomiyama et al., 2020).

In sugar-accumulating plants, ABA 8’-hydroxylase is expressed at low levels, suggesting a potential role for ABA regulation in sugar storage tissues (Figure 3— gene 13240). The dry climate during the sugar accumulation period in sugarcane, as observed during sample collection, indicates potential moderate water stress. Under water stress conditions, the sugarcane genotypes with the most efficient sugar accumulation tend to maintain greater stomatal conductance (Sajid et al., 2023). The lack of expression of the ABA catabolism gene suggested that the need for stomatal conductance regulation in these plants may be linked to maintaining open stomata. Additionally, in grapes, inhibition of the ABA 8’-hydroxylase gene improved tolerance to dehydration and promoted adventitious root formation, demonstrating an effective strategy for coping with water stress. Gene 23 was not expressed in the internodes (gene 13240—Figure 3), suggesting that the plant may use this strategy to improve its tolerance to potential water deficits.

Gene 01 (Supplementary Table 3) shares similarities with POP, which belongs to the serine protease family (clan SC, family S9) that includes various peptidases. POP is a cytoplasmic enzyme that hydrolyzes peptide bonds at the C-terminal side of proline residues (Gutierrez et al., 2008; Baharin et al., 2022). The enzyme’s three-dimensional structure allows for the postproline cleavage of peptides containing up to 30 amino acid residues (Gutierrez et al., 2008; Baharin et al., 2022). Post- or preproline cleavage enzymes can belong to different peptidase families, including aminopeptidases, endopeptidases, or oligopeptidases (PAP/PEP/POP). The most common domain in family S9 is a substrate-limiting β-propeller domain preventing unwanted digestion, while the α/β hydrolase domain catalyzes the reaction at the carboxy terminus of proline residues (Baharin et al., 2022). POP is a ubiquitous protein with a well-established structure and mechanism of action. However, its biological role in plants has not been fully elucidated. Increased expression in plants is known to be associated with tolerance to various types of abiotic stresses (Gutierrez et al., 2008; Singh et al., 2011; Tan et al., 2013). In flax, this phenomenon seems to be related to a fundamental mechanism for embryo growth in seeds (Gutierrez et al., 2008). In coffee, POP overexpression is linked to a significant increase in the number of branches in transgenic plants (Singh et al., 2011). Other peptidases that hydrolyze with proline specificity are related to plant development, such as pollen development (Ghifari et al., 2022), flowering, increased ABA activity, protection of photosynthetic activity during salt stress, elimination of reactive oxygen species, and overall osmotic potential adjustment (Ghosh et al., 2021).

Gene 01 is a DEG (Table 1) that is more highly expressed in the internodes of sugar-accumulating sampled plants and significantly less expressed in *S. spontaneum*, a sugar nonaccumulating sugarcane species known for its resistance to various stress types. Although POP is related to stress resistance and tolerance, there is more evidence suggesting that this gene is involved in this process. It has been observed that proline cleavage enzymes occur when a plant needs to accumulate proline, a phenomenon that usually occurs when the plant requires osmotic regulation due to stresses such as water and saline stress. It is also known that proline can bind to soluble sugars such as sucrose when there is a need to regulate the homeostasis of plant cells. Gene 01 has three first neighbors, one of which possesses the ENTH domain—a lipid-binding region crucial for clathrin-coated vesicle formation, endocytosis at the trans-Golgi network (TGN), and vacuolar transport. This gene could play a role in the transport of proline and sucrose, given its correlation with POP.

While sugarcane cannot accumulate sucrose under severe stress, previous studies have shown that mild water deficiency enhances photosynthetic rates and the accumulation of starch and sucrose in leaves (Garcia et al., 2018). The mechanism of proline accumulation is related to plants facing challenging situations, and the ability of proline to bind to sucrose adds another layer of complexity. Therefore, considering the paramount importance of comprehending sucrose accumulation processes in sugarcane and its connection with water deficit events during the period of peak sugar accumulation in the stems, a thorough examination of the role of POP is crucial. This includes exploring its potential association with proline accumulation and understanding how this accumulation might impact sugar storage.

Identifying genes that control or influence agronomic traits is one of the objectives of molecular breeding. Sugarcane, however, lags behind sorghum in terms of available genetic and genomic information. This study proposes a novel approach for transferring genetic knowledge from sorghum (donor) to sugarcane (recipient). Building upon existing methods (Shiringani et al., 2010; Garsmeur et al., 2011; Mancini et al., 2018; Sforça et al., 2019; Aono et al., 2021), we integrated genomic and coexpression network analysis to validate the relevance of sorghum-derived information in sugarcane. Furthermore, we analyzed the same genomic region in two Brazilian cultivars, revealing their differential genomic architecture and potential impact on sugar accumulation, using expression information to validate the results.

---

**Supplementary Figure 1.**
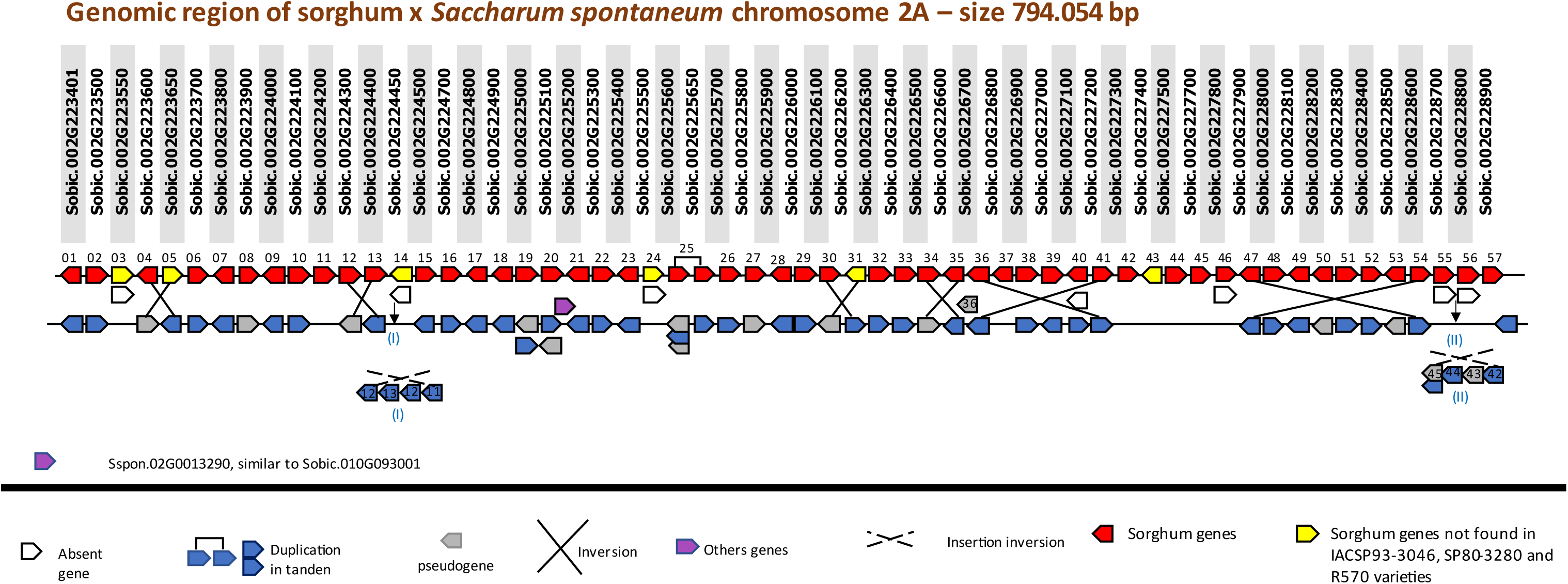
Sorghum×Sspon.2A genomic regions. *Genomic region of sorghum:* Each square represents a gene and shows the direction of the gene in the genome, where the right-facing arrow indicates the forward direction, and the left-facing arrow indicates the reverse direction. The solid lines with squares in red and yellow represent the 57 sorghum QTLs. *Genomic region of chromosome Sspon.2A:* Below the representation of the genomic region of sorghum is the orthologous genomic region in *S. spontaneum* for the chromosome Sspon.2A. Genes are shown in blue squares on a solid line, pseudogenes in gray, and genes that do not have orthologs in the sorghum QTL in purple. Each gene was numbered from 1 to 57 (Supplementary Table 3). The insertions of one or more gene clusters are shown by the Roman numerals indicated by the black arrows. In white, genes are depicted as absent not only in the represented chromosomal region but also throughout the entire chromosome. Inversions are indicated for crossed lines. The dotted crossed lines show an inversion inside an insertion. Synteny can be observed despite breaks in collinearity. An insertion represented by the purple square can be observed for this insertion in the gene Sspon.02G0013290, a duplicate of a gene from the same chromosome that is orthologous to sorghum, Sobic.010G093001, but in this case, the gene is observed on chromosome Sb01.

**Supplementary Figure 2.**
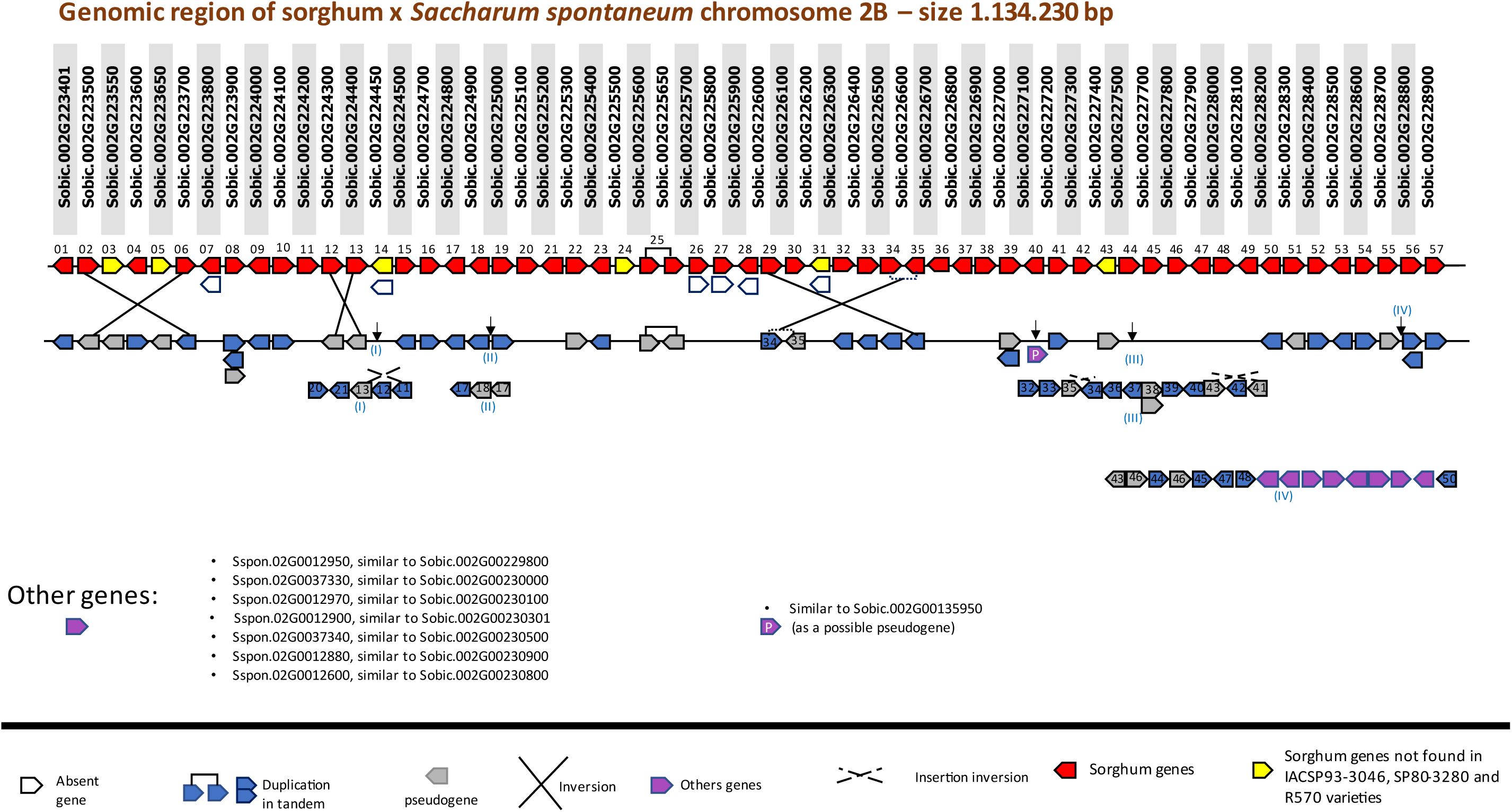
Sorghum×*S. spontaneum* Sspon.2B genomic regions. *Genomic region of sorghum:* Each square represents a gene and shows the direction of the gene in the genome, where the right-facing arrow indicates the forward direction, and the left-facing arrow indicates the reverse direction. The solid lines with squares in red and yellow represent the 57 sorghum QTLs. *Genomic region of Chromosome Sspon.2B:* Below the representation of the genomic region of sorghum is the orthologous genomic region in *S. spontaneum* for the chromosome Sspon.2B. Genes are shown in blue squares on a solid line, pseudogenes in gray, and genes that do not have orthologs in the sorghum QTL in purple. Each gene was numbered from 1 to 57 (Supplementary Table 3). The insertions of one or more gene clusters are shown by the Roman numerals indicated by the black arrows. In white, genes are depicted as absent not only in the represented chromosomal region but also throughout the entire chromosome. Inversions are indicated for crossed lines. The dotted cross lines show an inversion inside an insertion. Synteny can be observed despite breaks in collinearity. The cluster of genes depicted in purple in insertion IV is orthologous to sorghum, which has the same chromosome and the same sequence (Phytozome v.13). The absence of similar findings in the other *S. spontaneum* alleles, as well as in the IACSP93-3046, SP80-3280 and R570 haplotypes, suggests the possibility of a specific duplication in this particular allele.

**Supplementary Figure 3.**
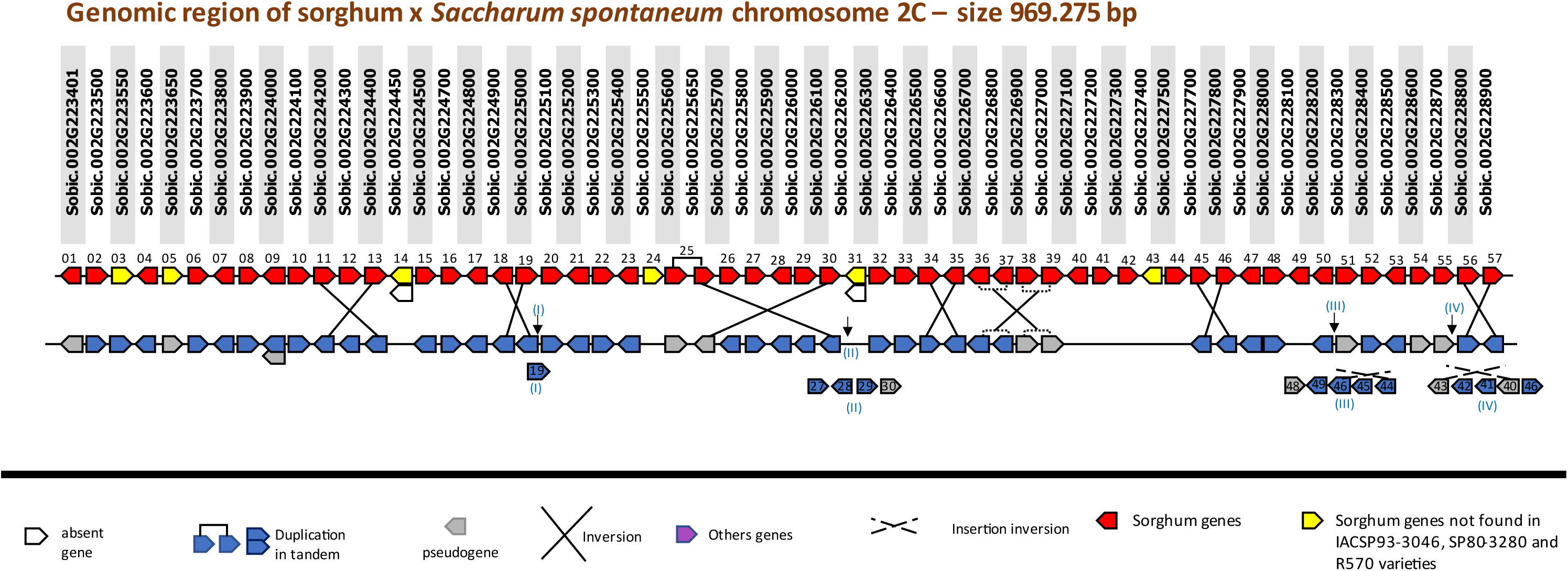
Sorghum×Sspon.2C. *Genomic region of sorghum:* Each square represents a gene and shows the direction of the gene in the genome, where the right-facing arrow indicates the forward direction, and the left-facing arrow indicates the reverse direction. The solid lines with squares in red and yellow represent the 57 sorghum QTLs. *Genomic region of chromosome Sspon2C:* Below the representation of the genomic region of sorghum are the orthologous genomic regions in *S. spontaneum* for each allele of chromosome Sspon.2C. Genes are shown in blue squares on a solid line, and pseudogenes are shown in gray. Each gene was numbered from 1 to 57 (Supplementary Table 3). The insertions of one or more gene clusters are shown by the Roman numerals indicated by the black arrows. In white, genes are depicted as absent not only in the represented chromosomal region but also throughout the entire chromosome. Inversions are indicated for crossed lines. The dotted cross lines show an inversion inside an insertion. Synteny can be observed despite breaks in collinearity.

**Supplementary Figure 4.**
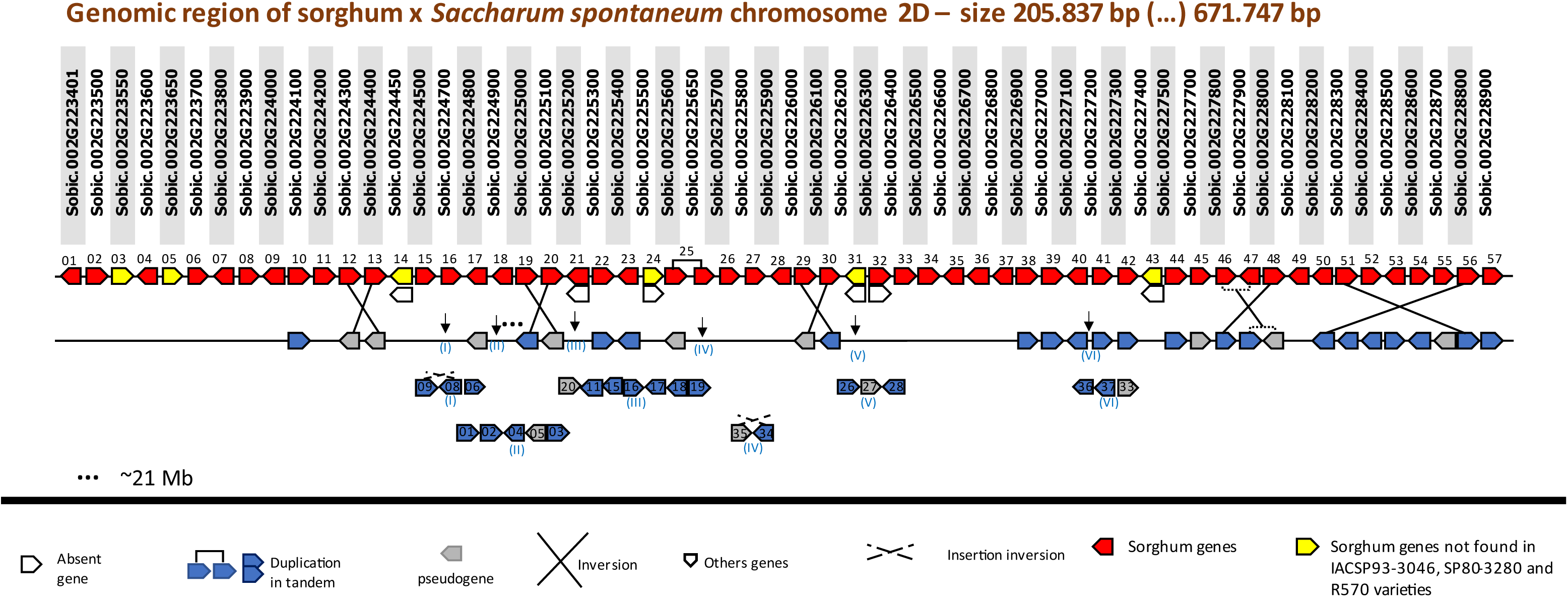
Sorghum×Sspon.2D. *Genomic region of sorghum:* Each square represents a gene and shows the direction of the gene in the genome, where the right-facing arrow indicates the forward direction, and the left-facing arrow indicates the reverse direction. The solid lines with squares in red and yellow represent the 57 sorghum QTLs. *Genomic region of chromosome Sspon.2D:* Below the representation of the genomic region of sorghum are the orthologous genomic regions in *S. spontaneum* for each allele of chromosome Sspon.2D. Genes are shown in blue squares on a solid line, and pseudogenes are shown in gray. Each gene was numbered from 1 to 57 (Supplementary Table 3). The insertions of one or more gene clusters are shown by the Roman numerals indicated by the black arrows. In white, genes are depicted as absent not only in the represented chromosomal region but also throughout the entire chromosome. Inversions are indicated for crossed lines. The dotted cross lines show an inversion inside an insertion. The three dots indicate a significant break in collinearity with sorghum, specifically on chromosome Sspon.2D, where the genomic region is separated by approximately 21 million base pairs. In addition to many other differences, many genes that seem to be absent in the allele are indeed present. Chromosomal absences are marked by white squares.

**Supplementary Figure 5.**
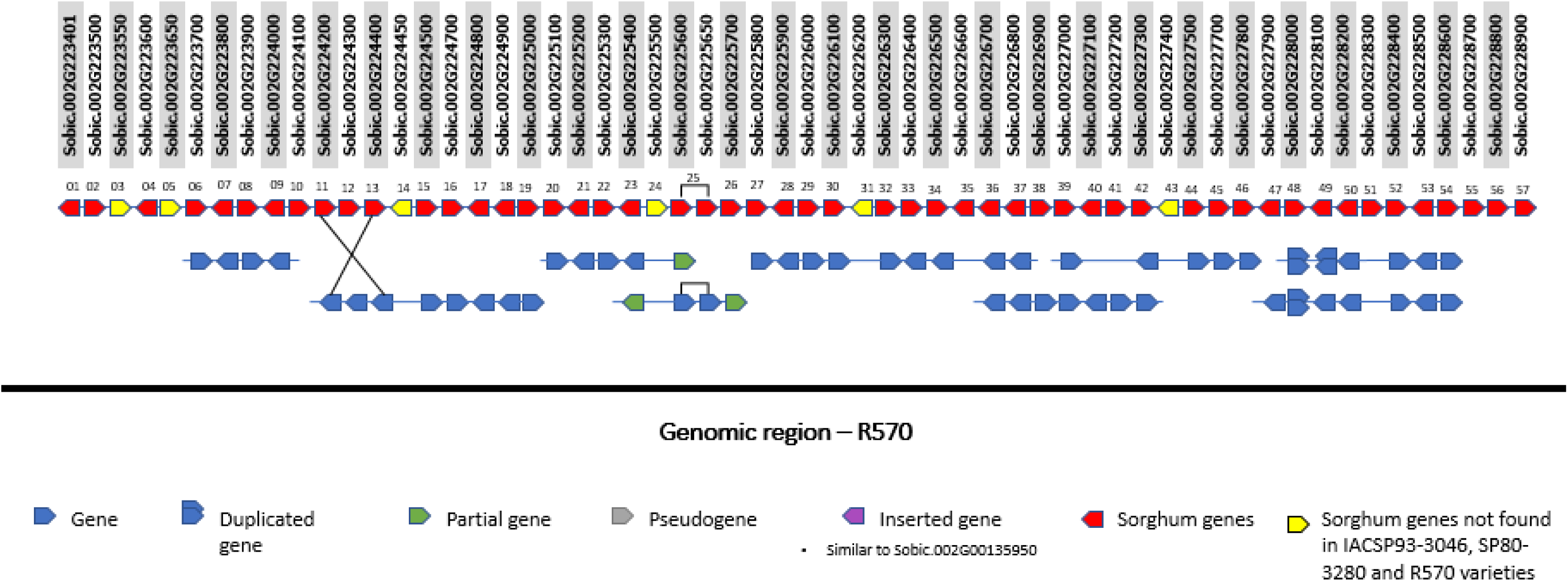
Sorghum×R570 genomic regions. *Genomic region of sorghum:* Each square represents a gene and shows the direction of the gene in the genome, where the right-facing arrow indicates the forward direction, and the left-facing arrow is the reverse direction. The solid lines with squares in red and yellow represent the 57 sorghum QTLs. *Genomic region of R570:* Below the representation of the sorghum genomic region, the orthologous genomic region in R570 is shown. Genes are depicted in blue. Possible pseudogenes are not represented (Garsmeur et al., 2018).

## Supplementary Tables

**Supplementary Table 1:**
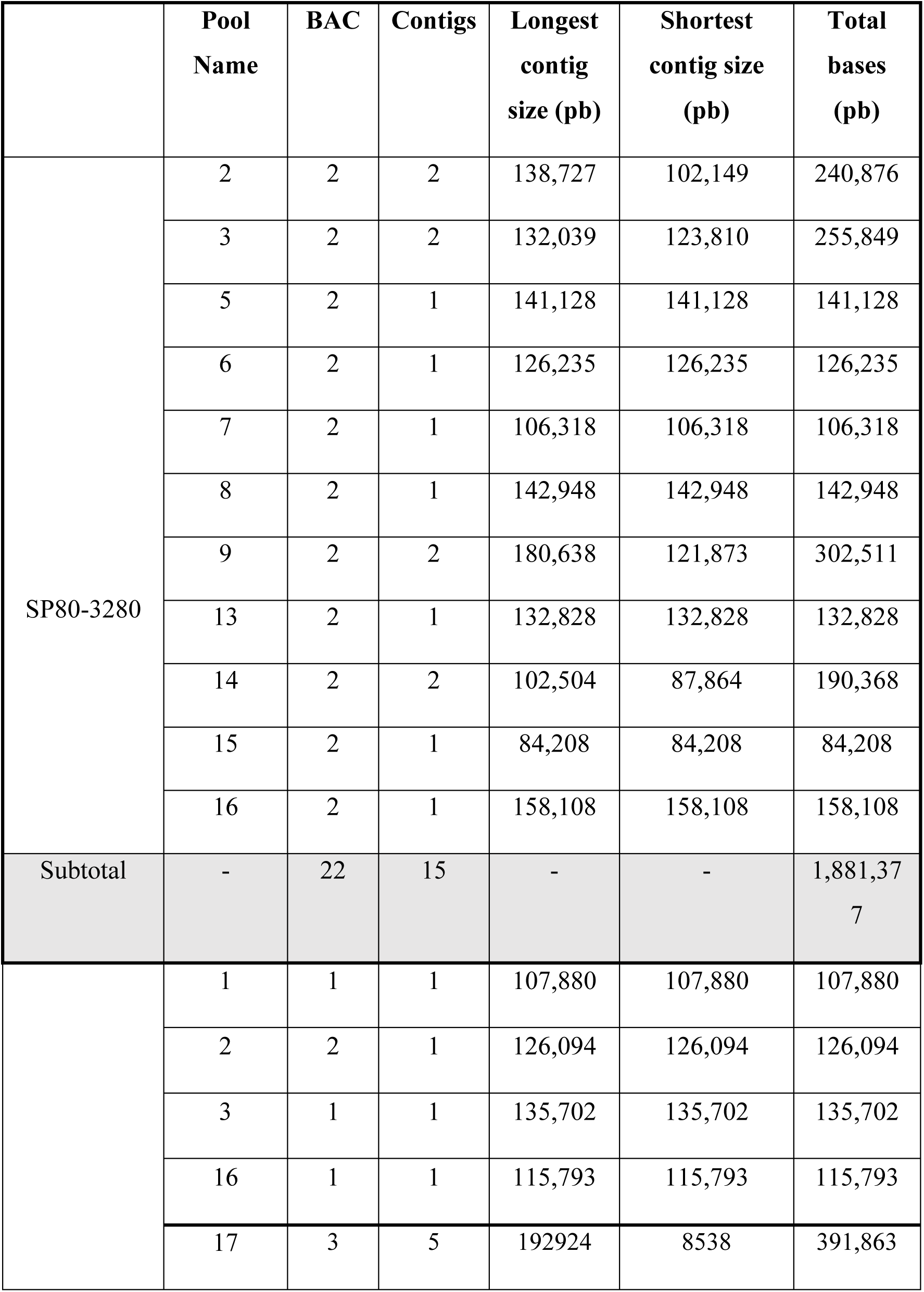
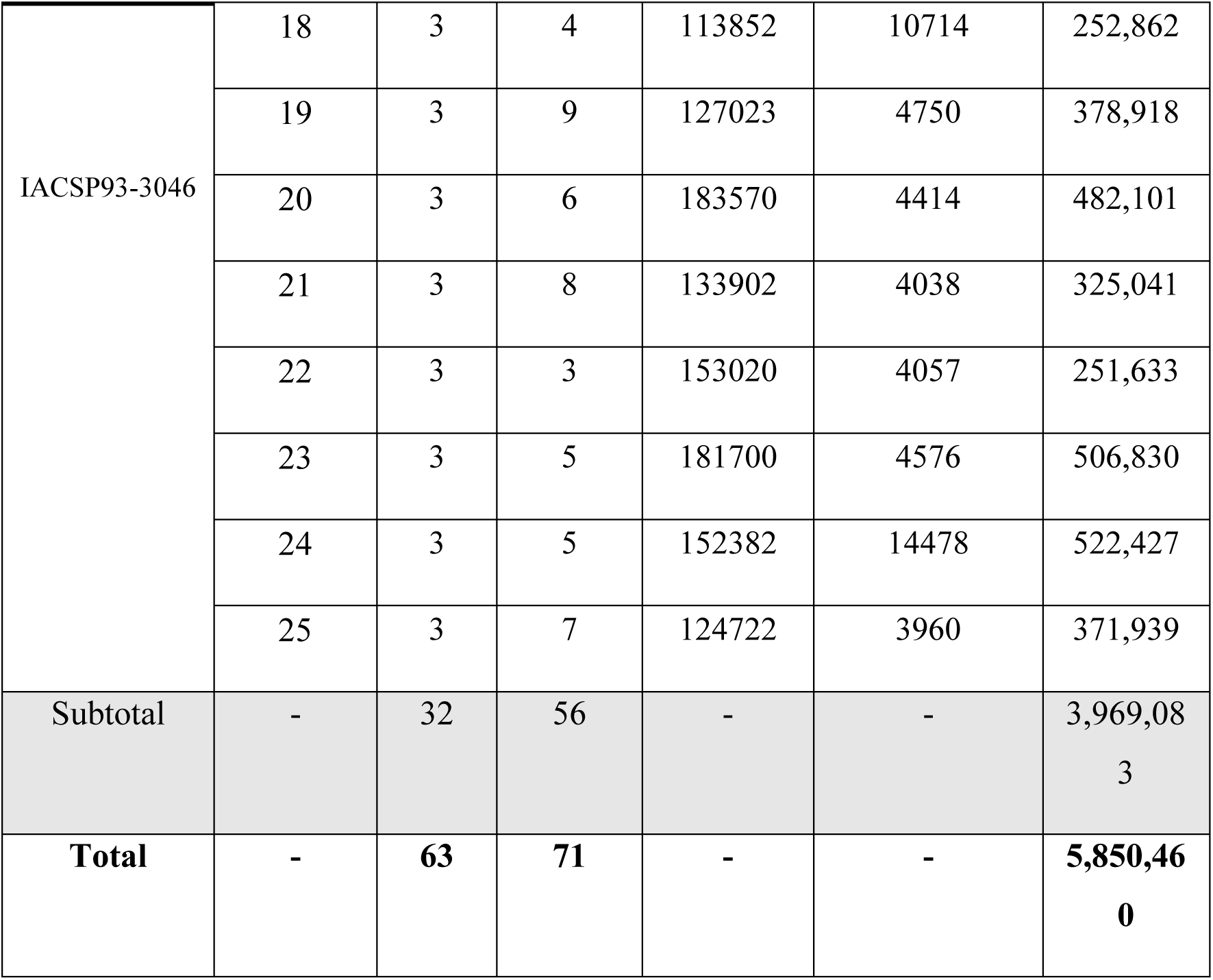
Summary of PacBio® Sequel sequencing and sequence assembly plus annotations.

**Supplementary Table 2:**
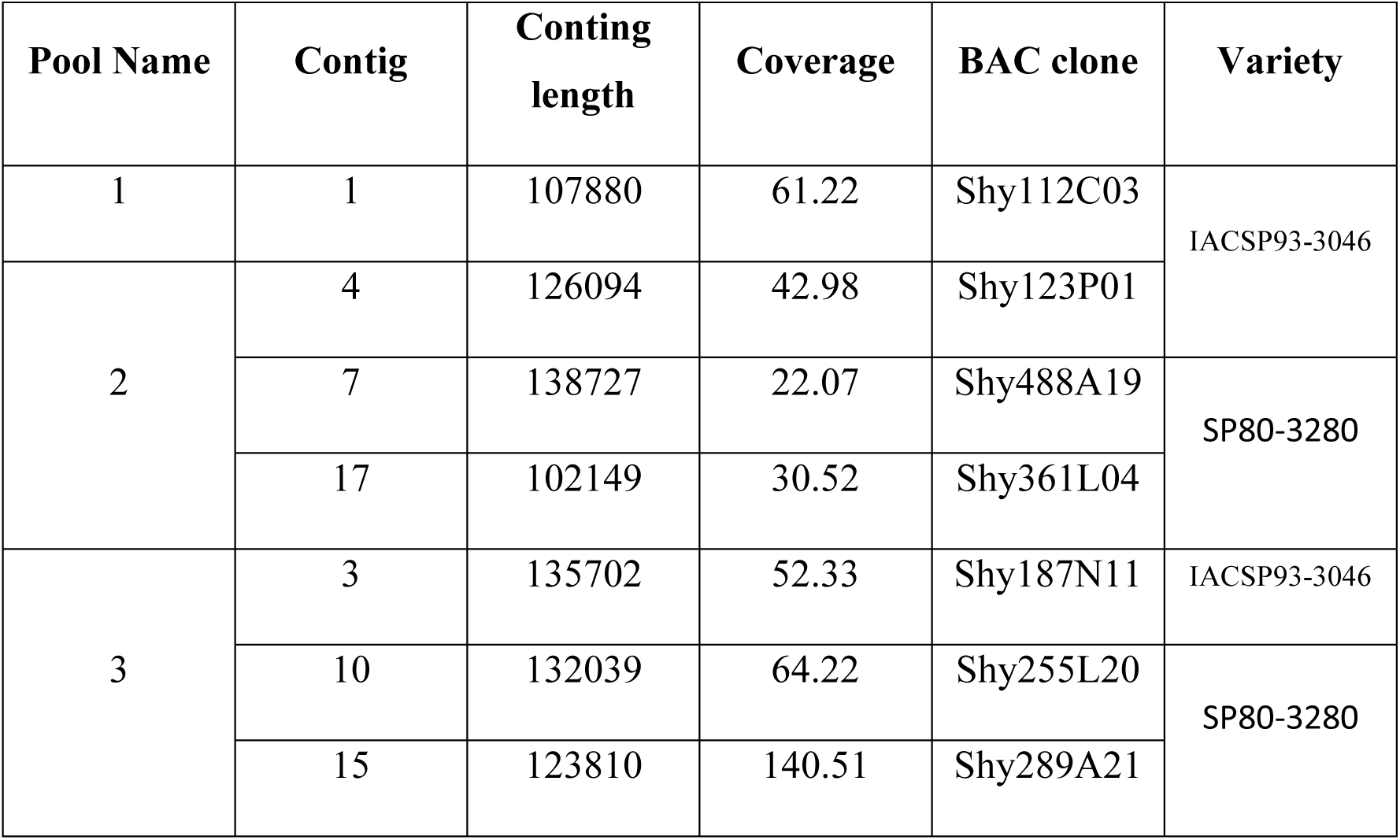
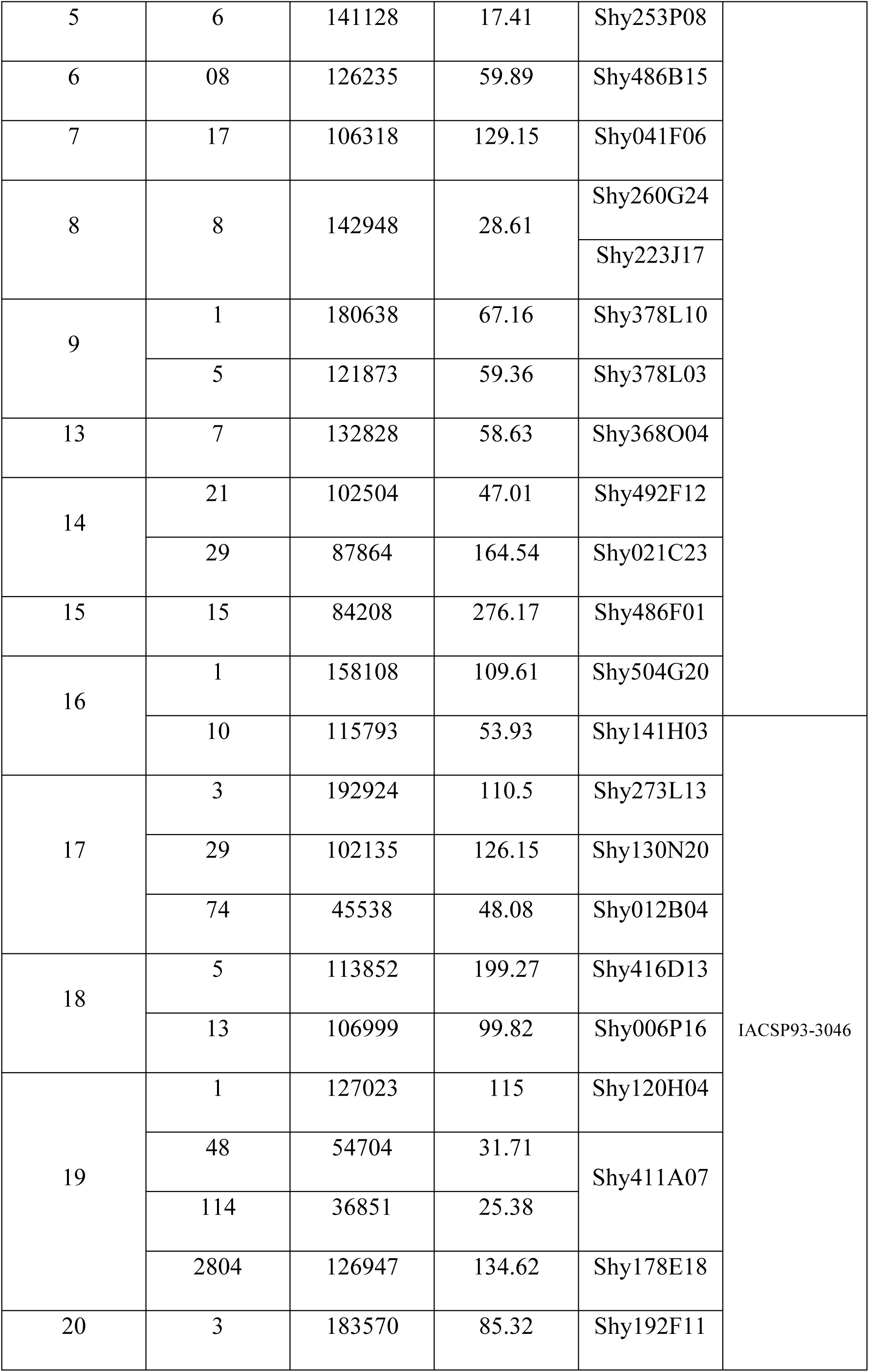
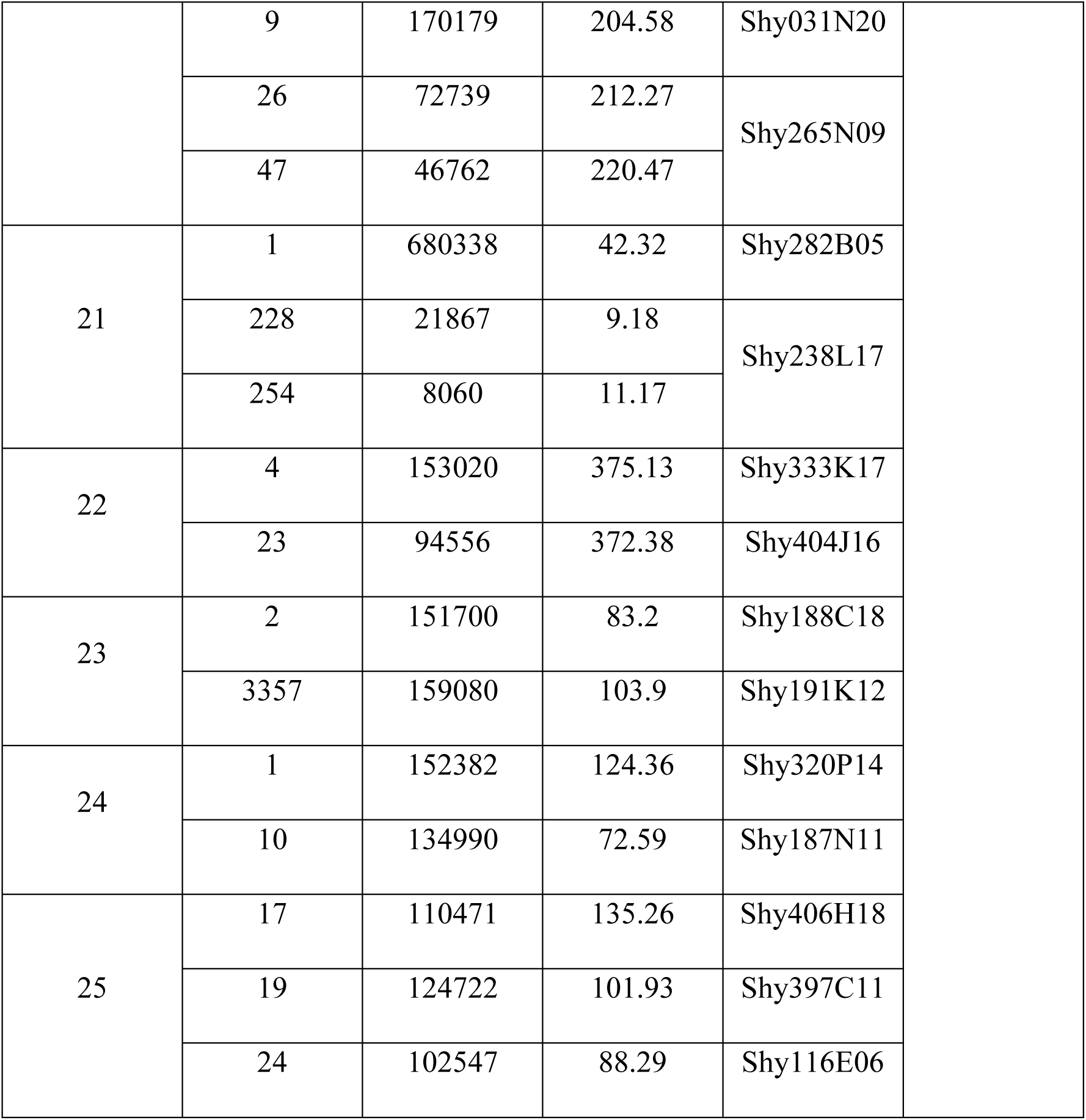
Results of PacBio® Sequel sequencing and sequence assembly plus annotation.

**Supplementary Table 03:** Summary of orthologous genes in sorghum and their proteins. Each gene was assigned a number following the order in which it appeared in the sorghum QTL. There are some absences in the column listing the names of the genes in *S. spontaneum*; however, this does not indicate that the genes are absent in the genomic region but rather that they were not detected by automated annotation (Zhang et al., 2018). In fact, only gene 14 was absent from chromosome Sspon.2 in *S. spontaneum.* The other genes (3, 5, 24, 27, 30, 31, 35, 36, 43, 51 and 55) were visualized and annotated manually (Supplementary Figures 1, 2, 3 and 4).

| Gene number | *S. bicolor* gene | *S. spontaneum* gene | Protein (Phytozome 13) |
| --- | --- | --- | --- |
| **01** | Sobic.002G223401 | Sspon.02G0013480 | Similar to a protein of the prolyl oligopeptidase family (**POP**) |
| **02** | Sobic.002G223500 | Sspon.02G0013470 | Predicted **AST**-like aspartate aminotransferase protein |
| **03** | Sobic.002G223550 | - | Predicted, uncharacterized protein. |
| **04** | Sobic.002G223600 | Sspon.02G0013460 | Predicted phosphatidylinositol-4-phosphate 5-kinase-related protein (**PIP5Ks**). |
| **05** | Sobic.002G223650 | - | Hypothetical, uncharacterized protein. |
| **06** | Sobic.002G223700 | Sspon.02G0013450 | Protein inferred by homology. Rhomboid-like protein RBL10. |
| **07** | Sobic.002G223800 | Sspon.02G0013440 | Hypothetical uncharacterized protein. |
| **08** | Sobic.002G223900 | Sspon.02G0013430 | Hypothetical protein. Similar to F-box-like proteins (FBPs). |
| **09** | Sobic.002G224000 | Sspon.02G0014420 | Similar to alpha-carbonic anhydrase domain-containing protein (CA). |
| **10** | Sobic.002G224100 | Sspon.02G0013410 | Similar to **Phosphate Transporter 3, PHT3**. |
| **11** | Sobic.002G224200 | Sspon.02G0013390 | Similar to **Heat shock factor B 4 (HSFB4) protein**. |
| **12** | Sobic.002G224300 | Sspon.02G0013400 | Predicted protein similar to NAM protein, an NAC transcription factor. |
| **13** | Sobic.002G224400 | Sspon.02G0037420 | **Receptor-like serine/threonine-protein kinase, lectin type G** |
| **14** | Sobic.002G224450 | - | Uncharacterized predicted protein. |
| **15** | Sobic.002G224500 | Sspon.02G0049360 | Similar to **BHLH domain-containing protein** (Basic helix-loop-helix). HEC |
| **16** | Sobic.002G224700 | Sspon.02G0013380 | **Inference by homology, NF - Kappa B activating protein** |
| **17** | Sobic.002G224800 | Sspon.02G0013370 | Protein belonging to the GDXG family, a family of lipolytic enzymes. Similar to alpha/beta hydrolase (with folded domain. Alpha/beta hydrolase fold domain - containing (ABH). |
| **18** | Sobic.002G224900 | Sspon.02G0037410 | Protein belonging to the GDXG family, a family of lipolytic enzymes. Similar to alpha/beta hydrolase (with folded domain. Alpha/beta hydrolase fold domain - containing (ABH). |
| **19** | Sobic.002G225000 | Sspon.02G0013350 | Protein inferred by homology. Similar to C-terminal remorin protein. Remorin C-Terminal like (REM). |
| **20** | Sobic.002G225100 | Sspon.02G0013320 | Similar to Absciscic Acid - insensitive 5 - like protein 4, **basic - leucine zipper domain (ABF1)**. |
| **21** | Sobic.002G225200 | Sspon.02G0013280 | Similar to putative uncharacterized protein B1342C04.33 |
| **22** | Sobic.002G225300 | Sspon.02G0013260 | Similar to **heat shock transcription factor B1 (HSB1) protein.** |
| **23** | Sobic.002G225400 | Sspon.02G0013240 | Similar to **abscisic acid 8'-hydroxylase (AA8' OH) protein.** |
| **24** | Sobic.002G225500 | - | Similar to a protein containing a **zinc finger domain** |
| **25** | Sobic.002G225600/<br>650 | Sspon.02G0013220 | Similar to alpha-amylase (**AMY**). |
| **26** | Sobic.002G225700 | Sspon.02G0013210 | Similar to **ethylene responsive factor 109 - (ERF109)** |
| **27** | Sobic.002G225800 | - | E3 ubiquitin ligase involved in syntaxin degradation |
| **28** | Sobic.002G225900 | Sspon.02G0013200 | Similar to fucosyltransferase (FUT). GO:0008417 |
| **29** | Sobic.002G226000 | Sspon.02G0013190 | Malectin-like domain-containing protein (**MLD**) |
| **30** | Sobic.002G226100 | - | Protein disulfide isomerase (**SCO2**) |
| **31** | Sobic.002G226200 | - | Uncharacterized protein |
| **32** | Sobic.002G226300 | Sspon.02G0013180 | Calmodulin - like protein (**CML**). |
| **33** | Sobic.002G226400 | Sspon.02G0013170 | Calmodulin - like protein (**CML**). |
| **34** | Sobic.002G226500 | Sspon.02G0013160 | Putative uncharacterized protein similar to basic leucine zipper |
|  |  |  | (BZIP) domain - containing protein. |
| **35** | Sobic.002G226600 | - | Uncharacterized protein |
| **36** | Sobic.002G226700 | - | similar to BRI1 **kinase inhibitor 1 (BKI1)** |
| **37** | Sobic.002G226800 | Sspon.02G0013130 | **PP2C**- protein phosphatase 2C // subfamily not named |
| **38** | Sobic.002G226900 | Sspon.02G0013140 | Similar to Enoyl-CoA reductase (**ECR**) |
| **39** | Sobic.002G227000 | Sspon.02G37380/37400 | Similar to Enoyl-CoA reductase (**ECR**) |
| **40** | Sobic.002G227100 | Sspon.02G0013150 | Cyclin-dependent kinase inhibitor (**CDK-inhibitor**) |
| **41** | Sobic.002G227200 | Sspon.02G0037390 | Similar to DNAJ-proteins or heat shock protein 40 (**HSP40**) |
| **42** | Sobic.002G227300 | Sspon.02G0037370 | Uncharacterized protein - predicted |
| **43** | Sobic.002G227400 | - | Similar to pentatricopeptide repeat (**PPR**) |
| **44** | Sobic.002G227500 | Sspon.02G0013040 | Similar to putative **SEC23** protein transport protein SEC23 |
| **45** | Sobic.002G227700 | Sspon.02G0013050 | Protein belonging to the GDXG family, a family of lipolytic enzymes. Similar to alpha/beta hydrolase (with folded domain. Alpha/beta hydrolase fold domain - containing (**ABH**). |
| **46** | Sobic.002G227800 | Sspon.02G0013060 | Protein belonging to the GDXG family, a family of lipolytic enzymes. Similar to alpha/beta hydrolase (with folded domain. Alpha/beta hydrolase fold domain - containing (**ABH**). |
| **47** | Sobic.002G227900 | Sspon.02G0013070 | Similar to alpha/beta hydrolase fold domain - containing (ABH). Gibberellin receptor (**GID1L2**). |
| **48** | Sobic.002G228000 | Sspon.02G0013080 | Similar to alpha/beta hydrolase fold domain - containing (ABH). Carboxylesterase (**CXE**). |
| **49** | Sobic.002G228100 | Sspon.02G0013090 | Similar to alpha/beta hydrolase fold domain - containing (ABH). Carboxylesterase (**CXE**). |
| **50** | Sobic.002G228200 | Sspon.02G0013090 | Similar to alpha/beta hydrolase fold domain - containing (ABH). Carboxylesterase (**CXE**). |
| **51** | Sobic.002G228300 | - | Uncharacterized and predicted protein. |
| **52** | Sobic.002G228400 | Sspon.02G0013100 | Similar to alpha/beta hydrolase fold domain - containing (ABH). Carboxylesterase (**CXE**). |
| **53** | Sobic.002G228500 | Sspon.02G0013110 | Similar to alpha/beta hydrolase fold domain - containing (ABH). Carboxylesterase (**CXE**). |
| **54** | Sobic.002G228600 | Sspon.02G0013120 | Similar to alpha/beta hydrolase fold domain - containing (ABH). Carboxylesterase (**CXE**). |
| **55** | Sobic.002G228700 | - | Similar to phospholipase A / patatin-related (**pPLA**) |
| **56** | Sobic.002G228800 | Sspon.02G0037310 | Uncharacterized protein -<br>predicted |
| **57** | Sobic.002G228900 | Sspon.02G0013030 | RNA Binding Protein - **RBP47** |

<u>**Supplementary Table 4.**</u> Differentially expressed genes (DEGs) identified between *Saccharum spontaneum* and *Saccharum officinarum*, IACSP93-3046 and SP80-3280. Log_2_(fold change) (Log_2_(FC)) values and false discovery rate (FDR)-corrected *p* values are provided for each gene.

