## Supplementary Table 4 for "Identifying Candidate Genes for Sugar Accumulation in Sugarcane Cultivars: From a Syntenic Genomic Region to a Gene Coexpression Network"

| Gene | log2(FC) | *p*-value |
|---|---|---|
| Sspon.01G0000070 | 20,426 | 2,27E-20 |
| Sspon.01G0000090 | -3,793 | 2,27E-20 |
| Sspon.01G0000190 | 21,122 | 2,45E-20 |
| Sspon.01G0000260 | -2,760 | 1,18E-18 |
| Sspon.01G0000280 | 5,223 | 2,61E-18 |
| Sspon.01G0000310 | 21,512 | 3,42E-18 |
| Sspon.01G0000350 | 3,238 | 5,51E-18 |
| Sspon.01G0000410 | 20,604 | 1,14E-17 |
| Sspon.01G0000490 | 3,626 | 1,55E-17 |
| Sspon.01G0000500 | 3,795 | 1,55E-17 |
| Sspon.01G0000610 | 20,589 | 1,73E-17 |
| Sspon.01G0000630 | 19,401 | 2,49E-17 |
| Sspon.01G0000720 | 19,455 | 2,49E-17 |
| Sspon.01G0000770 | 19,106 | 2,83E-17 |
| Sspon.01G0000820 | 18,811 | 4,12E-17 |
| Sspon.01G0000830 | 18,750 | 5,89E-17 |
| Sspon.01G0000880 | 18,773 | 3,11E-16 |
| Sspon.01G0000950 | -3,476 | 3,11E-16 |
| Sspon.01G0000960 | 3,503 | 3,11E-16 |
| Sspon.01G0000990 | -3,452 | 3,94E-16 |
| Sspon.01G0001000 | 18,232 | 5,47E-16 |
| Sspon.01G0001010 | 4,363 | 7,84E-16 |
| Sspon.01G0001090 | 5,304 | 8,12E-16 |
| Sspon.01G0001110 | 18,096 | 9,48E-16 |
| Sspon.01G0001120 | 19,141 | 9,48E-16 |
| Sspon.01G0001200 | 18,328 | 1,09E-15 |
| Sspon.01G0001240 | 3,363 | 1,29E-15 |
| Sspon.01G0001290 | 18,644 | 1,54E-15 |
| Sspon.01G0001400 | 3,142 | 1,54E-15 |
| Sspon.01G0001440 | 19,514 | 1,66E-15 |
| Sspon.01G0001460 | 18,772 | 1,66E-15 |
| Sspon.01G0001470 | 18,259 | 2,30E-15 |
| Sspon.01G0001540 | 18,433 | 2,64E-15 |
| Sspon.01G0001550 | 18,666 | 2,81E-15 |
| Sspon.01G0001590 | 20,041 | 3,52E-15 |
| Sspon.01G0001600 | 7,902 | 3,56E-15 |
| Sspon.01G0001650 | 18,425 | 3,77E-15 |
| Sspon.01G0001670 | 6,250 | 3,99E-15 |
| Sspon.01G0001700 | 2,814 | 4,75E-15 |
| Sspon.01G0001730 | 5,319 | 4,75E-15 |
| Sspon.01G0001780 | 2,738 | 4,87E-15 |
| Sspon.01G0002030 | 22,862 | 5,17E-15 |
| Sspon.01G0002040 | 2,633 | 6,29E-15 |

| | | |
|---|---|---|
| Sspon.01G0002180 | 18,756 | 6,78E-15 |
| Sspon.01G0002190 | -2,757 | 7,41E-15 |
| Sspon.01G0002210 | 18,587 | 7,57E-15 |
| Sspon.01G0002280 | 4,071 | 8,50E-15 |
| Sspon.01G0002310 | 19,924 | 9,08E-15 |
| Sspon.01G0002420 | -3,043 | 9,82E-15 |
| Sspon.01G0002430 | -2,634 | 9,82E-15 |
| Sspon.01G0002550 | -4,225 | 1,04E-14 |
| Sspon.01G0002570 | 18,140 | 1,09E-14 |
| Sspon.01G0002640 | 5,628 | 1,14E-14 |
| Sspon.01G0002660 | 17,775 | 1,20E-14 |
| Sspon.01G0002670 | 3,777 | 1,23E-14 |
| Sspon.01G0002690 | 21,288 | 1,32E-14 |
| Sspon.01G0002710 | 17,628 | 1,36E-14 |
| Sspon.01G0002730 | 2,604 | 1,43E-14 |
| Sspon.01G0002750 | -1,896 | 1,51E-14 |
| Sspon.01G0002770 | 17,537 | 1,55E-14 |
| Sspon.01G0002880 | 6,264 | 1,70E-14 |
| Sspon.01G0002890 | 18,699 | 1,70E-14 |
| Sspon.01G0003000 | 3,561 | 2,00E-14 |
| Sspon.01G0003070 | 17,571 | 2,17E-14 |
| Sspon.01G0003080 | -1,565 | 2,34E-14 |
| Sspon.01G0003100 | -1,778 | 3,13E-14 |
| Sspon.01G0003110 | 5,409 | 3,86E-14 |
| Sspon.01G0003170 | 20,838 | 4,33E-14 |
| Sspon.01G0003190 | 18,757 | 4,65E-14 |
| Sspon.01G0003200 | 4,662 | 4,65E-14 |
| Sspon.01G0003260 | -2,540 | 5,80E-14 |
| Sspon.01G0003320 | 18,909 | 6,31E-14 |
| Sspon.01G0003350 | 18,262 | 6,47E-14 |
| Sspon.01G0003530 | 17,826 | 7,59E-14 |
| Sspon.01G0003580 | -2,951 | 7,79E-14 |
| Sspon.01G0003620 | -3,947 | 9,08E-14 |
| Sspon.01G0003640 | 2,929 | 9,08E-14 |
| Sspon.01G0003760 | 3,096 | 9,53E-14 |
| Sspon.01G0003780 | 17,790 | 9,53E-14 |
| Sspon.01G0003840 | 8,098 | 9,70E-14 |
| Sspon.01G0003890 | 17,493 | 1,63E-13 |
| Sspon.01G0003920 | 5,666 | 1,68E-13 |
| Sspon.01G0003940 | -4,262 | 1,88E-13 |
| Sspon.01G0003950 | 3,900 | 2,23E-13 |
| Sspon.01G0004020 | -2,266 | 2,51E-13 |
| Sspon.01G0004070 | 4,075 | 2,80E-13 |

| | | |
|---|---|---|
| Sspon.01G0004180 | 5,833 | 3,51E-13 |
| Sspon.01G0004220 | -1,701 | 3,69E-13 |
| Sspon.01G0004250 | 5,137 | 3,70E-13 |
| Sspon.01G0004270 | 17,861 | 4,05E-13 |
| Sspon.01G0004330 | 3,196 | 4,23E-13 |
| Sspon.01G0004340 | 5,325 | 4,29E-13 |
| Sspon.01G0004360 | -1,900 | 4,43E-13 |
| Sspon.01G0004370 | 1,974 | 4,57E-13 |
| Sspon.01G0004380 | 18,696 | 4,57E-13 |
| Sspon.01G0004500 | -2,121 | 4,57E-13 |
| Sspon.01G0004520 | 4,256 | 4,59E-13 |
| Sspon.01G0004540 | -1,921 | 4,86E-13 |
| Sspon.01G0004550 | 4,185 | 5,35E-13 |
| Sspon.01G0004580 | 20,070 | 5,35E-13 |
| Sspon.01G0004670 | 18,870 | 5,82E-13 |
| Sspon.01G0004800 | 2,455 | 6,55E-13 |
| Sspon.01G0004840 | -3,691 | 7,03E-13 |
| Sspon.01G0004850 | 2,599 | 7,19E-13 |
| Sspon.01G0004860 | -1,574 | 7,19E-13 |
| Sspon.01G0004910 | 19,926 | 7,35E-13 |
| Sspon.01G0004970 | 2,991 | 7,69E-13 |
| Sspon.01G0005040 | 17,651 | 7,79E-13 |
| Sspon.01G0005150 | 18,450 | 7,81E-13 |
| Sspon.01G0005200 | 17,017 | 7,89E-13 |
| Sspon.01G0005230 | 4,112 | 7,89E-13 |
| Sspon.01G0005250 | 3,953 | 8,00E-13 |
| Sspon.01G0005290 | -3,007 | 8,36E-13 |
| Sspon.01G0005300 | 18,803 | 9,49E-13 |
| Sspon.01G0005310 | -1,389 | 1,02E-12 |
| Sspon.01G0005340 | 4,191 | 1,02E-12 |
| Sspon.01G0005430 | 3,313 | 1,09E-12 |
| Sspon.01G0005450 | -1,859 | 1,16E-12 |
| Sspon.01G0005530 | -3,228 | 1,29E-12 |
| Sspon.01G0005640 | -3,717 | 1,31E-12 |
| Sspon.01G0005730 | 3,268 | 1,35E-12 |
| Sspon.01G0005740 | 3,527 | 1,46E-12 |
| Sspon.01G0005840 | -1,295 | 1,52E-12 |
| Sspon.01G0005860 | 18,175 | 1,53E-12 |
| Sspon.01G0005990 | 16,762 | 1,63E-12 |
| Sspon.01G0006010 | 2,647 | 1,63E-12 |
| Sspon.01G0006030 | 3,144 | 1,67E-12 |
| Sspon.01G0006130 | 2,942 | 1,70E-12 |
| Sspon.01G0006250 | -2,449 | 1,77E-12 |

| | | |
|---|---|---|
| Sspon.01G0006310 | -2,497 | 1,88E-12 |
| Sspon.01G0006370 | 4,537 | 1,88E-12 |
| Sspon.01G0006410 | 5,966 | 2,00E-12 |
| Sspon.01G0006420 | 5,325 | 2,06E-12 |
| Sspon.01G0006470 | 3,611 | 2,07E-12 |
| Sspon.01G0006540 | -5,409 | 2,07E-12 |
| Sspon.01G0006550 | -3,427 | 2,08E-12 |
| Sspon.01G0006560 | 17,684 | 2,09E-12 |
| Sspon.01G0006640 | 5,780 | 2,11E-12 |
| Sspon.01G0006750 | -2,920 | 2,13E-12 |
| Sspon.01G0007110 | 17,665 | 2,20E-12 |
| Sspon.01G0007170 | 6,055 | 2,20E-12 |
| Sspon.01G0007230 | 3,669 | 2,21E-12 |
| Sspon.01G0007260 | 17,763 | 2,78E-12 |
| Sspon.01G0007290 | 18,522 | 2,78E-12 |
| Sspon.01G0007330 | 17,204 | 3,17E-12 |
| Sspon.01G0007360 | -1,898 | 3,25E-12 |
| Sspon.01G0007440 | -2,305 | 3,32E-12 |
| Sspon.01G0007450 | 17,511 | 3,39E-12 |
| Sspon.01G0007580 | -1,256 | 3,82E-12 |
| Sspon.01G0007610 | 3,824 | 3,83E-12 |
| Sspon.01G0007680 | 6,821 | 4,29E-12 |
| Sspon.01G0007690 | -2,545 | 4,29E-12 |
| Sspon.01G0007720 | 4,199 | 4,47E-12 |
| Sspon.01G0007740 | 17,743 | 4,60E-12 |
| Sspon.01G0007840 | 3,161 | 4,60E-12 |
| Sspon.01G0007920 | 2,908 | 4,77E-12 |
| Sspon.01G0007960 | 7,102 | 4,83E-12 |
| Sspon.01G0007990 | 17,344 | 4,91E-12 |
| Sspon.01G0008030 | -2,092 | 5,62E-12 |
| Sspon.01G0008120 | 2,204 | 5,62E-12 |
| Sspon.01G0008160 | 4,599 | 5,85E-12 |
| Sspon.01G0008200 | -1,493 | 5,85E-12 |
| Sspon.01G0008220 | 3,145 | 6,30E-12 |
| Sspon.01G0008230 | 18,369 | 6,56E-12 |
| Sspon.01G0008290 | -2,286 | 6,59E-12 |
| Sspon.01G0008300 | 2,609 | 6,59E-12 |
| Sspon.01G0008440 | 2,161 | 6,59E-12 |
| Sspon.01G0008450 | 3,719 | 6,63E-12 |
| Sspon.01G0008490 | 2,065 | 6,64E-12 |
| Sspon.01G0008520 | 17,237 | 6,81E-12 |
| Sspon.01G0008580 | 17,953 | 6,81E-12 |
| Sspon.01G0008610 | 19,156 | 6,81E-12 |

| | | |
|---|---|---|
| Sspon.01G0008710 | -2,359 | 7,62E-12 |
| Sspon.01G0008730 | 1,677 | 7,62E-12 |
| Sspon.01G0008790 | -2,307 | 7,62E-12 |
| Sspon.01G0008870 | 3,339 | 7,77E-12 |
| Sspon.01G0008880 | -3,372 | 9,12E-12 |
| Sspon.01G0008900 | -4,241 | 9,32E-12 |
| Sspon.01G0008910 | 4,278 | 9,32E-12 |
| Sspon.01G0008920 | 17,175 | 9,56E-12 |
| Sspon.01G0008950 | 4,630 | 9,75E-12 |
| Sspon.01G0009030 | 16,604 | 9,77E-12 |
| Sspon.01G0009070 | -2,958 | 1,01E-11 |
| Sspon.01G0009080 | -1,730 | 1,08E-11 |
| Sspon.01G0009140 | 18,738 | 1,09E-11 |
| Sspon.01G0009160 | 18,857 | 1,09E-11 |
| Sspon.01G0009170 | -1,464 | 1,11E-11 |
| Sspon.01G0009220 | 3,598 | 1,12E-11 |
| Sspon.01G0009250 | -2,209 | 1,18E-11 |
| Sspon.01G0009300 | -3,419 | 1,19E-11 |
| Sspon.01G0009350 | 3,324 | 1,23E-11 |
| Sspon.01G0009370 | 4,261 | 1,24E-11 |
| Sspon.01G0009430 | 4,870 | 1,38E-11 |
| Sspon.01G0009470 | 3,150 | 1,41E-11 |
| Sspon.01G0009480 | 3,202 | 1,45E-11 |
| Sspon.01G0009490 | -2,477 | 1,48E-11 |
| Sspon.01G0009500 | -1,625 | 1,48E-11 |
| Sspon.01G0009520 | 16,706 | 1,54E-11 |
| Sspon.01G0009790 | 5,610 | 1,59E-11 |
| Sspon.01G0009800 | -4,077 | 1,69E-11 |
| Sspon.01G0009810 | 1,647 | 1,97E-11 |
| Sspon.01G0009920 | 17,054 | 2,16E-11 |
| Sspon.01G0010010 | 2,673 | 2,17E-11 |
| Sspon.01G0010020 | 17,737 | 2,26E-11 |
| Sspon.01G0010060 | 17,334 | 2,42E-11 |
| Sspon.01G0010220 | 1,400 | 2,51E-11 |
| Sspon.01G0010240 | 2,515 | 2,60E-11 |
| Sspon.01G0010250 | 1,923 | 2,60E-11 |
| Sspon.01G0010320 | 2,671 | 2,81E-11 |
| Sspon.01G0010380 | 16,320 | 2,87E-11 |
| Sspon.01G0010420 | -2,654 | 2,88E-11 |
| Sspon.01G0010440 | -4,553 | 2,93E-11 |
| Sspon.01G0010460 | -1,354 | 2,95E-11 |
| Sspon.01G0010480 | -3,788 | 3,02E-11 |
| Sspon.01G0010510 | 16,971 | 3,19E-11 |

| | | |
|---|---|---|
| Sspon.01G0010520 | 3,619 | 3,22E-11 |
| Sspon.01G0010530 | -2,616 | 3,26E-11 |
| Sspon.01G0010570 | -2,316 | 3,26E-11 |
| Sspon.01G0010600 | -1,506 | 3,33E-11 |
| Sspon.01G0010610 | 17,020 | 3,35E-11 |
| Sspon.01G0010620 | 1,359 | 3,44E-11 |
| Sspon.01G0010640 | -2,584 | 3,44E-11 |
| Sspon.01G0010700 | -2,501 | 3,48E-11 |
| Sspon.01G0010770 | 3,536 | 3,48E-11 |
| Sspon.01G0010870 | 4,521 | 3,48E-11 |
| Sspon.01G0010880 | 17,424 | 3,58E-11 |
| Sspon.01G0010920 | -5,447 | 3,65E-11 |
| Sspon.01G0010960 | 4,264 | 3,91E-11 |
| Sspon.01G0011020 | 3,001 | 3,92E-11 |
| Sspon.01G0011030 | 8,997 | 4,10E-11 |
| Sspon.01G0011110 | -1,129 | 4,10E-11 |
| Sspon.01G0011130 | 4,223 | 4,33E-11 |
| Sspon.01G0011200 | -2,086 | 4,38E-11 |
| Sspon.01G0011240 | -4,427 | 4,38E-11 |
| Sspon.01G0011250 | 3,994 | 4,43E-11 |
| Sspon.01G0011270 | -1,098 | 4,90E-11 |
| Sspon.01G0011290 | -2,085 | 4,94E-11 |
| Sspon.01G0011300 | -1,875 | 4,95E-11 |
| Sspon.01G0011320 | 18,553 | 4,95E-11 |
| Sspon.01G0011330 | 1,143 | 5,02E-11 |
| Sspon.01G0011360 | -2,579 | 5,14E-11 |
| Sspon.01G0011410 | -3,464 | 5,26E-11 |
| Sspon.01G0011420 | 16,433 | 5,30E-11 |
| Sspon.01G0011520 | 2,994 | 5,36E-11 |
| Sspon.01G0011530 | 3,473 | 5,36E-11 |
| Sspon.01G0011650 | 3,819 | 5,37E-11 |
| Sspon.01G0011670 | 17,682 | 5,58E-11 |
| Sspon.01G0011850 | -1,095 | 6,15E-11 |
| Sspon.01G0012010 | 3,884 | 6,21E-11 |
| Sspon.01G0012050 | 3,793 | 6,53E-11 |
| Sspon.01G0012100 | 17,468 | 6,59E-11 |
| Sspon.01G0012140 | 5,145 | 6,66E-11 |
| Sspon.01G0012200 | 4,277 | 7,03E-11 |
| Sspon.01G0012220 | -4,075 | 7,04E-11 |
| Sspon.01G0012230 | -3,380 | 7,18E-11 |
| Sspon.01G0012240 | -2,555 | 7,36E-11 |
| Sspon.01G0012250 | 1,590 | 7,73E-11 |
| Sspon.01G0012280 | 4,752 | 8,40E-11 |

| | | |
|---|---|---|
| Sspon.01G0012290 | -2,147 | 8,40E-11 |
| Sspon.01G0012320 | 2,267 | 8,40E-11 |
| Sspon.01G0012360 | -1,599 | 8,52E-11 |
| Sspon.01G0012370 | 3,315 | 8,66E-11 |
| Sspon.01G0012470 | -2,101 | 9,01E-11 |
| Sspon.01G0012710 | 5,890 | 9,06E-11 |
| Sspon.01G0012730 | 19,551 | 9,10E-11 |
| Sspon.01G0012810 | -1,054 | 1,01E-10 |
| Sspon.01G0012870 | -1,854 | 1,01E-10 |
| Sspon.01G0012900 | 1,246 | 1,01E-10 |
| Sspon.01G0012930 | 3,325 | 1,02E-10 |
| Sspon.01G0012990 | 2,659 | 1,05E-10 |
| Sspon.01G0013070 | -1,509 | 1,07E-10 |
| Sspon.01G0013080 | -4,742 | 1,08E-10 |
| Sspon.01G0013110 | -1,665 | 1,22E-10 |
| Sspon.01G0013280 | 4,286 | 1,25E-10 |
| Sspon.01G0013290 | 5,604 | 1,25E-10 |
| Sspon.01G0013330 | 2,463 | 1,33E-10 |
| Sspon.01G0013350 | -3,066 | 1,35E-10 |
| Sspon.01G0013370 | 4,837 | 1,39E-10 |
| Sspon.01G0013410 | -3,059 | 1,40E-10 |
| Sspon.01G0013440 | 6,177 | 1,42E-10 |
| Sspon.01G0013620 | 2,761 | 1,43E-10 |
| Sspon.01G0013690 | -1,888 | 1,43E-10 |
| Sspon.01G0013750 | -1,002 | 1,45E-10 |
| Sspon.01G0013820 | 3,555 | 1,46E-10 |
| Sspon.01G0013870 | -2,734 | 1,47E-10 |
| Sspon.01G0014050 | 17,198 | 1,49E-10 |
| Sspon.01G0014060 | -1,347 | 1,52E-10 |
| Sspon.01G0014090 | 17,705 | 1,53E-10 |
| Sspon.01G0014120 | 16,162 | 1,54E-10 |
| Sspon.01G0014370 | 16,731 | 1,54E-10 |
| Sspon.01G0014490 | -1,148 | 1,56E-10 |
| Sspon.01G0014560 | 1,198 | 1,64E-10 |
| Sspon.01G0014690 | 18,857 | 1,64E-10 |
| Sspon.01G0014840 | -2,938 | 1,66E-10 |
| Sspon.01G0014860 | -1,684 | 1,69E-10 |
| Sspon.01G0014900 | -1,285 | 1,75E-10 |
| Sspon.01G0014980 | -1,398 | 1,75E-10 |
| Sspon.01G0015000 | 1,822 | 1,77E-10 |
| Sspon.01G0015120 | 3,024 | 1,77E-10 |
| Sspon.01G0015190 | -1,198 | 1,78E-10 |
| Sspon.01G0015220 | 2,764 | 1,85E-10 |

| | | |
|---|---|---|
| Sspon.01G0015240 | -1,013 | 1,87E-10 |
| Sspon.01G0015260 | 1,103 | 1,89E-10 |
| Sspon.01G0015290 | -1,383 | 1,91E-10 |
| Sspon.01G0015300 | 16,159 | 2,04E-10 |
| Sspon.01G0015320 | 18,332 | 2,06E-10 |
| Sspon.01G0015380 | -5,152 | 2,06E-10 |
| Sspon.01G0015400 | 3,616 | 2,06E-10 |
| Sspon.01G0015410 | -1,811 | 2,09E-10 |
| Sspon.01G0015420 | 16,147 | 2,14E-10 |
| Sspon.01G0015440 | 5,808 | 2,17E-10 |
| Sspon.01G0015450 | 2,391 | 2,25E-10 |
| Sspon.01G0015480 | 2,269 | 2,28E-10 |
| Sspon.01G0015530 | 5,383 | 2,36E-10 |
| Sspon.01G0015580 | 2,173 | 2,41E-10 |
| Sspon.01G0015590 | 2,736 | 2,42E-10 |
| Sspon.01G0015680 | -2,757 | 2,45E-10 |
| Sspon.01G0015740 | -1,668 | 2,46E-10 |
| Sspon.01G0015880 | 2,951 | 2,49E-10 |
| Sspon.01G0015970 | -3,383 | 2,49E-10 |
| Sspon.01G0016030 | 4,107 | 2,66E-10 |
| Sspon.01G0016110 | -1,603 | 2,66E-10 |
| Sspon.01G0016120 | -2,707 | 2,66E-10 |
| Sspon.01G0016160 | 2,192 | 2,73E-10 |
| Sspon.01G0016170 | 3,387 | 2,78E-10 |
| Sspon.01G0016260 | 17,748 | 2,78E-10 |
| Sspon.01G0016470 | -3,289 | 2,87E-10 |
| Sspon.01G0016490 | 16,054 | 2,90E-10 |
| Sspon.01G0016500 | -3,917 | 2,91E-10 |
| Sspon.01G0016520 | -20,892 | 2,96E-10 |
| Sspon.01G0016550 | 2,807 | 3,03E-10 |
| Sspon.01G0016570 | 2,182 | 3,05E-10 |
| Sspon.01G0016630 | 2,121 | 3,11E-10 |
| Sspon.01G0016680 | 1,980 | 3,40E-10 |
| Sspon.01G0016700 | 2,461 | 3,59E-10 |
| Sspon.01G0016760 | 4,413 | 3,66E-10 |
| Sspon.01G0016790 | -4,417 | 3,78E-10 |
| Sspon.01G0016830 | 5,186 | 3,82E-10 |
| Sspon.01G0016850 | -2,000 | 4,12E-10 |
| Sspon.01G0016920 | -1,763 | 4,28E-10 |
| Sspon.01G0016960 | 16,812 | 4,35E-10 |
| Sspon.01G0017040 | 5,335 | 4,45E-10 |
| Sspon.01G0017090 | -1,880 | 4,59E-10 |
| Sspon.01G0017130 | -1,841 | 4,62E-10 |

| | | |
|---|---|---|
| Sspon.01G0017160 | -20,708 | 4,81E-10 |
| Sspon.01G0017170 | -3,449 | 4,81E-10 |
| Sspon.01G0017180 | -3,435 | 4,88E-10 |
| Sspon.01G0017190 | -2,773 | 4,89E-10 |
| Sspon.01G0017200 | -2,468 | 4,91E-10 |
| Sspon.01G0017210 | -5,330 | 4,99E-10 |
| Sspon.01G0017240 | -3,216 | 5,01E-10 |
| Sspon.01G0017270 | 4,564 | 5,13E-10 |
| Sspon.01G0017360 | 1,349 | 5,24E-10 |
| Sspon.01G0017370 | 5,073 | 5,25E-10 |
| Sspon.01G0017400 | 3,283 | 5,75E-10 |
| Sspon.01G0017470 | -1,442 | 5,75E-10 |
| Sspon.01G0017510 | 1,120 | 6,11E-10 |
| Sspon.01G0017570 | -2,996 | 6,14E-10 |
| Sspon.01G0017630 | 1,403 | 6,20E-10 |
| Sspon.01G0017680 | 4,065 | 6,31E-10 |
| Sspon.01G0017750 | -1,100 | 6,38E-10 |
| Sspon.01G0017930 | 3,733 | 6,43E-10 |
| Sspon.01G0018020 | 2,247 | 6,46E-10 |
| Sspon.01G0018080 | 5,825 | 6,48E-10 |
| Sspon.01G0018090 | -3,704 | 6,50E-10 |
| Sspon.01G0018100 | 3,161 | 6,57E-10 |
| Sspon.01G0018120 | 3,978 | 6,68E-10 |
| Sspon.01G0018170 | 2,709 | 6,68E-10 |
| Sspon.01G0018300 | -2,196 | 6,70E-10 |
| Sspon.01G0018310 | -2,738 | 7,06E-10 |
| Sspon.01G0018350 | -2,091 | 7,14E-10 |
| Sspon.01G0018360 | 1,939 | 7,24E-10 |
| Sspon.01G0018410 | -2,495 | 7,24E-10 |
| Sspon.01G0018480 | 2,686 | 7,27E-10 |
| Sspon.01G0018490 | 3,534 | 7,59E-10 |
| Sspon.01G0018500 | 16,665 | 7,90E-10 |
| Sspon.01G0018580 | 4,653 | 7,95E-10 |
| Sspon.01G0018650 | 2,795 | 8,16E-10 |
| Sspon.01G0018680 | 2,759 | 8,16E-10 |
| Sspon.01G0018720 | -1,654 | 8,48E-10 |
| Sspon.01G0018780 | -6,733 | 8,63E-10 |
| Sspon.01G0018800 | -1,148 | 8,63E-10 |
| Sspon.01G0018990 | 3,273 | 8,63E-10 |
| Sspon.01G0019010 | -1,240 | 9,34E-10 |
| Sspon.01G0019040 | -2,812 | 9,34E-10 |
| Sspon.01G0019050 | 2,211 | 9,46E-10 |
| Sspon.01G0019160 | 2,390 | 9,48E-10 |

| | | |
|---|---|---|
| Sspon.01G0019260 | -1,506 | 9,53E-10 |
| Sspon.01G0019320 | -1,160 | 9,64E-10 |
| Sspon.01G0019330 | 5,263 | 9,76E-10 |
| Sspon.01G0019350 | -2,065 | 9,92E-10 |
| Sspon.01G0019360 | 16,548 | 9,92E-10 |
| Sspon.01G0019440 | 1,408 | 1,01E-09 |
| Sspon.01G0019470 | 2,070 | 1,01E-09 |
| Sspon.01G0019520 | -1,876 | 1,02E-09 |
| Sspon.01G0019540 | 1,889 | 1,02E-09 |
| Sspon.01G0019580 | 1,974 | 1,03E-09 |
| Sspon.01G0019700 | 3,912 | 1,07E-09 |
| Sspon.01G0019720 | -1,999 | 1,07E-09 |
| Sspon.01G0019750 | 2,109 | 1,07E-09 |
| Sspon.01G0019760 | 3,082 | 1,07E-09 |
| Sspon.01G0019780 | 2,813 | 1,08E-09 |
| Sspon.01G0019960 | -1,356 | 1,11E-09 |
| Sspon.01G0019970 | -1,528 | 1,11E-09 |
| Sspon.01G0020170 | 2,123 | 1,15E-09 |
| Sspon.01G0020200 | 1,373 | 1,15E-09 |
| Sspon.01G0020210 | -1,843 | 1,15E-09 |
| Sspon.01G0020240 | 2,322 | 1,16E-09 |
| Sspon.01G0020570 | -1,036 | 1,16E-09 |
| Sspon.01G0020630 | 2,123 | 1,19E-09 |
| Sspon.01G0020700 | 5,502 | 1,19E-09 |
| Sspon.01G0020790 | -1,197 | 1,21E-09 |
| Sspon.01G0020810 | -2,934 | 1,21E-09 |
| Sspon.01G0020910 | -4,906 | 1,22E-09 |
| Sspon.01G0020970 | 2,628 | 1,22E-09 |
| Sspon.01G0020980 | -2,449 | 1,27E-09 |
| Sspon.01G0021000 | -21,561 | 1,28E-09 |
| Sspon.01G0021020 | -1,249 | 1,34E-09 |
| Sspon.01G0021140 | 2,478 | 1,37E-09 |
| Sspon.01G0021200 | -1,451 | 1,37E-09 |
| Sspon.01G0021220 | -2,416 | 1,39E-09 |
| Sspon.01G0021230 | 17,050 | 1,39E-09 |
| Sspon.01G0021250 | 3,100 | 1,40E-09 |
| Sspon.01G0021260 | -1,962 | 1,40E-09 |
| Sspon.01G0021310 | 3,600 | 1,40E-09 |
| Sspon.01G0021410 | -2,005 | 1,41E-09 |
| Sspon.01G0021460 | 16,808 | 1,41E-09 |
| Sspon.01G0021530 | 2,843 | 1,44E-09 |
| Sspon.01G0021540 | 2,007 | 1,45E-09 |
| Sspon.01G0021650 | -1,514 | 1,48E-09 |

| | | |
|---|---|---|
| Sspon.01G0021670 | 2,317 | 1,49E-09 |
| Sspon.01G0021700 | 4,494 | 1,50E-09 |
| Sspon.01G0021830 | -1,901 | 1,52E-09 |
| Sspon.01G0021860 | -1,351 | 1,52E-09 |
| Sspon.01G0021920 | -3,297 | 1,52E-09 |
| Sspon.01G0022030 | 1,806 | 1,60E-09 |
| Sspon.01G0022060 | -1,730 | 1,68E-09 |
| Sspon.01G0022110 | -1,705 | 1,70E-09 |
| Sspon.01G0022220 | 15,539 | 1,72E-09 |
| Sspon.01G0022250 | -1,003 | 1,75E-09 |
| Sspon.01G0022300 | 3,696 | 1,75E-09 |
| Sspon.01G0022380 | -1,862 | 1,75E-09 |
| Sspon.01G0022520 | -2,350 | 1,78E-09 |
| Sspon.01G0022600 | 16,354 | 1,84E-09 |
| Sspon.01G0022910 | -4,880 | 1,86E-09 |
| Sspon.01G0022940 | 1,102 | 1,87E-09 |
| Sspon.01G0022980 | -1,641 | 1,88E-09 |
| Sspon.01G0023070 | -3,228 | 1,88E-09 |
| Sspon.01G0023110 | -2,253 | 1,89E-09 |
| Sspon.01G0023180 | -2,267 | 1,89E-09 |
| Sspon.01G0023350 | -3,734 | 1,89E-09 |
| Sspon.01G0023360 | 3,171 | 1,89E-09 |
| Sspon.01G0023440 | -2,534 | 1,94E-09 |
| Sspon.01G0023570 | 16,557 | 1,94E-09 |
| Sspon.01G0023580 | 17,070 | 1,94E-09 |
| Sspon.01G0023610 | 1,879 | 1,94E-09 |
| Sspon.01G0023620 | 2,826 | 1,97E-09 |
| Sspon.01G0023690 | 2,595 | 1,99E-09 |
| Sspon.01G0023720 | -2,808 | 1,99E-09 |
| Sspon.01G0023800 | -1,141 | 2,00E-09 |
| Sspon.01G0023830 | 18,286 | 2,02E-09 |
| Sspon.01G0023950 | 6,222 | 2,05E-09 |
| Sspon.01G0023970 | -1,223 | 2,05E-09 |
| Sspon.01G0023980 | 3,954 | 2,06E-09 |
| Sspon.01G0023990 | 2,979 | 2,07E-09 |
| Sspon.01G0024020 | 1,647 | 2,13E-09 |
| Sspon.01G0024060 | 1,488 | 2,13E-09 |
| Sspon.01G0024110 | -1,741 | 2,15E-09 |
| Sspon.01G0024140 | -1,444 | 2,19E-09 |
| Sspon.01G0024170 | -3,078 | 2,25E-09 |
| Sspon.01G0024190 | -2,283 | 2,27E-09 |
| Sspon.01G0024240 | -4,123 | 2,31E-09 |
| Sspon.01G0024250 | 3,884 | 2,32E-09 |

| | | |
|---|---|---|
| Sspon.01G0024290 | -2,668 | 2,35E-09 |
| Sspon.01G0024330 | -2,518 | 2,38E-09 |
| Sspon.01G0024410 | -1,712 | 2,42E-09 |
| Sspon.01G0024440 | -1,442 | 2,46E-09 |
| Sspon.01G0024520 | 18,310 | 2,46E-09 |
| Sspon.01G0024530 | -1,468 | 2,52E-09 |
| Sspon.01G0024550 | -19,912 | 2,52E-09 |
| Sspon.01G0024590 | 1,632 | 2,68E-09 |
| Sspon.01G0024650 | 2,089 | 2,70E-09 |
| Sspon.01G0024770 | 2,381 | 2,75E-09 |
| Sspon.01G0024870 | 2,902 | 2,80E-09 |
| Sspon.01G0024970 | 1,259 | 2,86E-09 |
| Sspon.01G0024990 | -2,769 | 2,95E-09 |
| Sspon.01G0025030 | -1,261 | 3,02E-09 |
| Sspon.01G0025060 | -1,424 | 3,02E-09 |
| Sspon.01G0025090 | -1,230 | 3,03E-09 |
| Sspon.01G0025140 | 2,108 | 3,13E-09 |
| Sspon.01G0025210 | 2,268 | 3,13E-09 |
| Sspon.01G0025220 | 2,064 | 3,16E-09 |
| Sspon.01G0025230 | -1,695 | 3,17E-09 |
| Sspon.01G0025260 | -1,596 | 3,19E-09 |
| Sspon.01G0025310 | 2,853 | 3,22E-09 |
| Sspon.01G0025320 | -1,729 | 3,22E-09 |
| Sspon.01G0025350 | 3,924 | 3,23E-09 |
| Sspon.01G0025390 | -3,038 | 3,26E-09 |
| Sspon.01G0025460 | -1,479 | 3,29E-09 |
| Sspon.01G0025590 | -1,072 | 3,34E-09 |
| Sspon.01G0025830 | -1,408 | 3,39E-09 |
| Sspon.01G0025840 | 3,423 | 3,39E-09 |
| Sspon.01G0025930 | -1,077 | 3,54E-09 |
| Sspon.01G0026010 | -2,247 | 3,54E-09 |
| Sspon.01G0026110 | -1,794 | 3,57E-09 |
| Sspon.01G0026430 | -2,674 | 3,60E-09 |
| Sspon.01G0026460 | 1,875 | 3,60E-09 |
| Sspon.01G0026480 | 1,783 | 3,62E-09 |
| Sspon.01G0026590 | 2,005 | 3,65E-09 |
| Sspon.01G0026600 | -1,762 | 3,69E-09 |
| Sspon.01G0026630 | 3,798 | 3,84E-09 |
| Sspon.01G0026870 | 1,968 | 3,87E-09 |
| Sspon.01G0026950 | 4,446 | 3,89E-09 |
| Sspon.01G0027100 | -2,227 | 3,96E-09 |
| Sspon.01G0027140 | 2,735 | 3,96E-09 |
| Sspon.01G0027150 | 2,391 | 4,11E-09 |

| | | |
|---|---|---|
| Sspon.01G0027220 | 1,156 | 4,11E-09 |
| Sspon.01G0027230 | -1,611 | 4,24E-09 |
| Sspon.01G0027280 | -1,153 | 4,25E-09 |
| Sspon.01G0027300 | 2,283 | 4,25E-09 |
| Sspon.01G0027310 | 1,117 | 4,27E-09 |
| Sspon.01G0027410 | 1,889 | 4,28E-09 |
| Sspon.01G0027430 | 1,385 | 4,33E-09 |
| Sspon.01G0027440 | 4,177 | 4,61E-09 |
| Sspon.01G0027520 | 2,765 | 4,74E-09 |
| Sspon.01G0027550 | 1,950 | 4,91E-09 |
| Sspon.01G0027570 | 1,546 | 4,95E-09 |
| Sspon.01G0027580 | 3,751 | 4,99E-09 |
| Sspon.01G0027610 | 3,037 | 5,02E-09 |
| Sspon.01G0027630 | -1,839 | 5,03E-09 |
| Sspon.01G0027640 | 3,579 | 5,08E-09 |
| Sspon.01G0027670 | -1,042 | 5,14E-09 |
| Sspon.01G0027810 | -1,230 | 5,15E-09 |
| Sspon.01G0027840 | 3,198 | 5,20E-09 |
| Sspon.01G0027850 | 3,423 | 5,22E-09 |
| Sspon.01G0027890 | 3,839 | 5,26E-09 |
| Sspon.01G0027900 | -2,535 | 5,27E-09 |
| Sspon.01G0027940 | -1,236 | 5,44E-09 |
| Sspon.01G0027970 | 3,899 | 5,50E-09 |
| Sspon.01G0027980 | 2,739 | 5,55E-09 |
| Sspon.01G0028000 | -1,142 | 5,65E-09 |
| Sspon.01G0028080 | -1,269 | 5,69E-09 |
| Sspon.01G0028090 | -19,486 | 5,70E-09 |
| Sspon.01G0028100 | 1,411 | 5,70E-09 |
| Sspon.01G0028120 | 3,111 | 5,77E-09 |
| Sspon.01G0028230 | -4,401 | 5,87E-09 |
| Sspon.01G0028270 | 2,167 | 5,90E-09 |
| Sspon.01G0028310 | 3,087 | 5,90E-09 |
| Sspon.01G0028440 | -2,982 | 5,90E-09 |
| Sspon.01G0028450 | -3,556 | 5,97E-09 |
| Sspon.01G0028770 | -1,572 | 6,09E-09 |
| Sspon.01G0028820 | 3,089 | 6,39E-09 |
| Sspon.01G0028830 | -1,175 | 6,51E-09 |
| Sspon.01G0028840 | -2,790 | 6,55E-09 |
| Sspon.01G0028860 | -1,081 | 6,66E-09 |
| Sspon.01G0028870 | -1,050 | 6,75E-09 |
| Sspon.01G0028970 | 2,319 | 6,76E-09 |
| Sspon.01G0029020 | 1,941 | 7,03E-09 |
| Sspon.01G0029090 | -1,370 | 7,05E-09 |

| | | |
|---|---|---|
| Sspon.01G0029100 | -2,729 | 7,05E-09 |
| Sspon.01G0029150 | 2,586 | 7,07E-09 |
| Sspon.01G0029160 | -2,108 | 7,09E-09 |
| Sspon.01G0029230 | 1,438 | 7,22E-09 |
| Sspon.01G0029240 | 3,175 | 7,25E-09 |
| Sspon.01G0029290 | -1,523 | 7,26E-09 |
| Sspon.01G0029350 | -1,329 | 7,26E-09 |
| Sspon.01G0029370 | -2,072 | 7,26E-09 |
| Sspon.01G0029440 | -2,160 | 7,26E-09 |
| Sspon.01G0029500 | -2,017 | 7,44E-09 |
| Sspon.01G0029520 | 1,251 | 7,44E-09 |
| Sspon.01G0029540 | 1,407 | 7,44E-09 |
| Sspon.01G0029560 | -1,254 | 7,52E-09 |
| Sspon.01G0029610 | 2,154 | 7,55E-09 |
| Sspon.01G0029640 | 3,153 | 7,61E-09 |
| Sspon.01G0029770 | -1,707 | 7,61E-09 |
| Sspon.01G0029780 | 1,884 | 7,67E-09 |
| Sspon.01G0029790 | 2,378 | 8,05E-09 |
| Sspon.01G0029820 | 1,373 | 8,54E-09 |
| Sspon.01G0029840 | 6,292 | 8,66E-09 |
| Sspon.01G0029850 | 1,187 | 8,76E-09 |
| Sspon.01G0029890 | 1,014 | 8,89E-09 |
| Sspon.01G0029900 | 2,164 | 8,97E-09 |
| Sspon.01G0030090 | -3,221 | 9,00E-09 |
| Sspon.01G0030100 | 3,951 | 9,07E-09 |
| Sspon.01G0030110 | 1,586 | 9,30E-09 |
| Sspon.01G0030140 | -2,103 | 9,34E-09 |
| Sspon.01G0030260 | 5,631 | 9,34E-09 |
| Sspon.01G0030290 | 3,284 | 9,41E-09 |
| Sspon.01G0030370 | -2,066 | 9,50E-09 |
| Sspon.01G0030390 | 1,820 | 9,76E-09 |
| Sspon.01G0030510 | -2,370 | 9,86E-09 |
| Sspon.01G0030560 | 1,778 | 9,90E-09 |
| Sspon.01G0030640 | 3,767 | 1,01E-08 |
| Sspon.01G0030690 | -2,869 | 1,05E-08 |
| Sspon.01G0030700 | -1,358 | 1,05E-08 |
| Sspon.01G0030710 | -1,204 | 1,07E-08 |
| Sspon.01G0030720 | 1,665 | 1,09E-08 |
| Sspon.01G0030760 | -1,474 | 1,11E-08 |
| Sspon.01G0030820 | 1,238 | 1,12E-08 |
| Sspon.01G0030830 | -2,016 | 1,12E-08 |
| Sspon.01G0030920 | 1,296 | 1,14E-08 |
| Sspon.01G0030940 | 1,793 | 1,16E-08 |

| | | |
|---|---|---|
| Sspon.01G0031050 | -1,769 | 1,17E-08 |
| Sspon.01G0031080 | 1,165 | 1,18E-08 |
| Sspon.01G0031140 | 1,736 | 1,22E-08 |
| Sspon.01G0031150 | 1,935 | 1,22E-08 |
| Sspon.01G0031180 | -2,555 | 1,24E-08 |
| Sspon.01G0031270 | 2,087 | 1,26E-08 |
| Sspon.01G0031290 | 1,110 | 1,26E-08 |
| Sspon.01G0031420 | 10,367 | 1,27E-08 |
| Sspon.01G0031430 | -1,111 | 1,27E-08 |
| Sspon.01G0031500 | 2,771 | 1,27E-08 |
| Sspon.01G0031510 | -1,659 | 1,28E-08 |
| Sspon.01G0031630 | -1,498 | 1,29E-08 |
| Sspon.01G0031650 | -2,591 | 1,30E-08 |
| Sspon.01G0031710 | 2,509 | 1,31E-08 |
| Sspon.01G0031750 | -1,185 | 1,39E-08 |
| Sspon.01G0031760 | 1,654 | 1,40E-08 |
| Sspon.01G0031880 | -5,664 | 1,40E-08 |
| Sspon.01G0031890 | -2,046 | 1,40E-08 |
| Sspon.01G0031900 | -3,226 | 1,41E-08 |
| Sspon.01G0032000 | -1,172 | 1,41E-08 |
| Sspon.01G0032080 | -20,922 | 1,45E-08 |
| Sspon.01G0032100 | 1,357 | 1,45E-08 |
| Sspon.01G0032360 | -19,533 | 1,50E-08 |
| Sspon.01G0032410 | 17,278 | 1,52E-08 |
| Sspon.01G0032430 | -2,705 | 1,52E-08 |
| Sspon.01G0032510 | -3,092 | 1,56E-08 |
| Sspon.01G0032540 | -2,158 | 1,56E-08 |
| Sspon.01G0032570 | -2,600 | 1,58E-08 |
| Sspon.01G0032700 | 3,049 | 1,60E-08 |
| Sspon.01G0032720 | -3,948 | 1,64E-08 |
| Sspon.01G0032730 | -1,373 | 1,66E-08 |
| Sspon.01G0032750 | -1,419 | 1,66E-08 |
| Sspon.01G0032760 | 1,557 | 1,66E-08 |
| Sspon.01G0032780 | -2,766 | 1,67E-08 |
| Sspon.01G0032790 | -1,422 | 1,68E-08 |
| Sspon.01G0032830 | 1,576 | 1,70E-08 |
| Sspon.01G0032860 | 1,699 | 1,72E-08 |
| Sspon.01G0032870 | -1,795 | 1,75E-08 |
| Sspon.01G0032940 | 1,430 | 1,76E-08 |
| Sspon.01G0032990 | -1,426 | 1,76E-08 |
| Sspon.01G0033020 | -1,805 | 1,77E-08 |
| Sspon.01G0033060 | -4,918 | 1,79E-08 |
| Sspon.01G0033130 | -1,048 | 1,81E-08 |

| | | |
|---|---|---|
| Sspon.01G0033170 | -2,202 | 1,84E-08 |
| Sspon.01G0033180 | 4,517 | 1,88E-08 |
| Sspon.01G0033210 | 4,135 | 1,94E-08 |
| Sspon.01G0033220 | 1,438 | 1,94E-08 |
| Sspon.01G0033230 | -1,708 | 1,96E-08 |
| Sspon.01G0033260 | 2,689 | 1,96E-08 |
| Sspon.01G0033280 | 2,724 | 1,96E-08 |
| Sspon.01G0033360 | 3,441 | 1,96E-08 |
| Sspon.01G0033400 | 3,314 | 1,97E-08 |
| Sspon.01G0033500 | -1,469 | 2,03E-08 |
| Sspon.01G0033540 | 3,219 | 2,06E-08 |
| Sspon.01G0033590 | -1,697 | 2,09E-08 |
| Sspon.01G0033680 | 3,682 | 2,10E-08 |
| Sspon.01G0033700 | 1,860 | 2,10E-08 |
| Sspon.01G0033880 | -1,026 | 2,10E-08 |
| Sspon.01G0033900 | -1,187 | 2,10E-08 |
| Sspon.01G0033930 | 4,129 | 2,12E-08 |
| Sspon.01G0033990 | 2,329 | 2,13E-08 |
| Sspon.01G0034070 | -2,130 | 2,15E-08 |
| Sspon.01G0034150 | -2,594 | 2,15E-08 |
| Sspon.01G0034170 | 2,555 | 2,22E-08 |
| Sspon.01G0034200 | 18,634 | 2,25E-08 |
| Sspon.01G0034230 | -2,423 | 2,28E-08 |
| Sspon.01G0034270 | -2,809 | 2,28E-08 |
| Sspon.01G0034320 | -1,061 | 2,36E-08 |
| Sspon.01G0034330 | 3,156 | 2,36E-08 |
| Sspon.01G0034340 | 4,286 | 2,37E-08 |
| Sspon.01G0034350 | -4,232 | 2,38E-08 |
| Sspon.01G0034400 | 3,029 | 2,40E-08 |
| Sspon.01G0034420 | 1,395 | 2,40E-08 |
| Sspon.01G0034490 | 3,417 | 2,40E-08 |
| Sspon.01G0034540 | -1,659 | 2,44E-08 |
| Sspon.01G0034620 | -2,223 | 2,48E-08 |
| Sspon.01G0034630 | -1,676 | 2,52E-08 |
| Sspon.01G0034860 | -1,340 | 2,52E-08 |
| Sspon.01G0035020 | -1,200 | 2,61E-08 |
| Sspon.01G0035040 | 2,858 | 2,63E-08 |
| Sspon.01G0035100 | -19,124 | 2,63E-08 |
| Sspon.01G0035180 | 1,852 | 2,64E-08 |
| Sspon.01G0035300 | 3,989 | 2,64E-08 |
| Sspon.01G0035340 | -1,775 | 2,65E-08 |
| Sspon.01G0035450 | 2,379 | 2,66E-08 |
| Sspon.01G0035520 | -1,330 | 2,66E-08 |

| | | |
|---|---|---|
| Sspon.01G0035580 | 2,774 | 2,67E-08 |
| Sspon.01G0035610 | -2,207 | 2,67E-08 |
| Sspon.01G0035640 | 1,036 | 2,70E-08 |
| Sspon.01G0035680 | 3,925 | 2,71E-08 |
| Sspon.01G0035780 | 3,848 | 2,72E-08 |
| Sspon.01G0035910 | -1,606 | 2,82E-08 |
| Sspon.01G0036030 | 3,909 | 2,82E-08 |
| Sspon.01G0036060 | 4,455 | 2,83E-08 |
| Sspon.01G0036090 | -1,326 | 2,92E-08 |
| Sspon.01G0036230 | -2,389 | 2,94E-08 |
| Sspon.01G0036320 | -1,149 | 2,95E-08 |
| Sspon.01G0036350 | -1,745 | 3,01E-08 |
| Sspon.01G0036380 | 2,437 | 3,07E-08 |
| Sspon.01G0036420 | -1,587 | 3,08E-08 |
| Sspon.01G0036430 | -2,218 | 3,12E-08 |
| Sspon.01G0036500 | 19,923 | 3,15E-08 |
| Sspon.01G0036580 | -1,093 | 3,16E-08 |
| Sspon.01G0036610 | -2,754 | 3,16E-08 |
| Sspon.01G0036760 | -1,240 | 3,17E-08 |
| Sspon.01G0036770 | 3,181 | 3,24E-08 |
| Sspon.01G0036780 | 2,261 | 3,25E-08 |
| Sspon.01G0036800 | -2,444 | 3,26E-08 |
| Sspon.01G0036820 | -2,651 | 3,26E-08 |
| Sspon.01G0036860 | -1,095 | 3,27E-08 |
| Sspon.01G0036950 | 2,945 | 3,27E-08 |
| Sspon.01G0036960 | 3,834 | 3,32E-08 |
| Sspon.01G0037070 | -1,349 | 3,38E-08 |
| Sspon.01G0037090 | 2,108 | 3,41E-08 |
| Sspon.01G0037220 | 1,143 | 3,44E-08 |
| Sspon.01G0037290 | -1,613 | 3,44E-08 |
| Sspon.01G0037470 | 17,123 | 3,44E-08 |
| Sspon.01G0037480 | -1,396 | 3,44E-08 |
| Sspon.01G0037550 | -3,566 | 3,52E-08 |
| Sspon.01G0037660 | -4,265 | 3,53E-08 |
| Sspon.01G0037690 | 1,502 | 3,60E-08 |
| Sspon.01G0037710 | -18,910 | 3,61E-08 |
| Sspon.01G0037730 | -1,201 | 3,61E-08 |
| Sspon.01G0037780 | -1,418 | 3,64E-08 |
| Sspon.01G0037800 | 5,009 | 3,67E-08 |
| Sspon.01G0037810 | 1,876 | 3,67E-08 |
| Sspon.01G0037830 | -3,714 | 3,69E-08 |
| Sspon.01G0037850 | 16,798 | 3,70E-08 |
| Sspon.01G0037930 | -3,037 | 3,70E-08 |

| | | |
|---|---|---|
| Sspon.01G0037950 | 1,937 | 3,70E-08 |
| Sspon.01G0037960 | 2,985 | 3,70E-08 |
| Sspon.01G0038030 | -2,977 | 3,72E-08 |
| Sspon.01G0038040 | -1,346 | 3,74E-08 |
| Sspon.01G0038050 | -1,180 | 3,75E-08 |
| Sspon.01G0038060 | -1,189 | 3,76E-08 |
| Sspon.01G0038070 | -2,279 | 3,77E-08 |
| Sspon.01G0038080 | 2,115 | 3,77E-08 |
| Sspon.01G0038100 | 2,062 | 3,80E-08 |
| Sspon.01G0038210 | 2,393 | 3,83E-08 |
| Sspon.01G0038270 | -2,069 | 3,84E-08 |
| Sspon.01G0038280 | -1,535 | 3,86E-08 |
| Sspon.01G0038320 | -1,795 | 3,89E-08 |
| Sspon.01G0038390 | 1,277 | 3,95E-08 |
| Sspon.01G0038430 | -1,492 | 3,95E-08 |
| Sspon.01G0038520 | 1,318 | 3,96E-08 |
| Sspon.01G0038600 | 17,714 | 3,98E-08 |
| Sspon.01G0038620 | -1,457 | 3,99E-08 |
| Sspon.01G0038650 | 1,080 | 4,01E-08 |
| Sspon.01G0038680 | -1,249 | 4,03E-08 |
| Sspon.01G0038710 | -1,333 | 4,05E-08 |
| Sspon.01G0038770 | -1,340 | 4,07E-08 |
| Sspon.01G0038820 | 17,049 | 4,07E-08 |
| Sspon.01G0038900 | 3,635 | 4,16E-08 |
| Sspon.01G0038920 | 2,259 | 4,16E-08 |
| Sspon.01G0038970 | 2,791 | 4,22E-08 |
| Sspon.01G0038990 | -1,362 | 4,22E-08 |
| Sspon.01G0039020 | -1,655 | 4,22E-08 |
| Sspon.01G0039060 | 1,553 | 4,23E-08 |
| Sspon.01G0039070 | -3,385 | 4,31E-08 |
| Sspon.01G0039080 | 1,521 | 4,39E-08 |
| Sspon.01G0039090 | 1,853 | 4,47E-08 |
| Sspon.01G0039110 | -1,022 | 4,54E-08 |
| Sspon.01G0039180 | -1,507 | 4,54E-08 |
| Sspon.01G0039230 | -1,304 | 4,57E-08 |
| Sspon.01G0039280 | -1,875 | 4,62E-08 |
| Sspon.01G0039380 | -1,108 | 4,65E-08 |
| Sspon.01G0039410 | -1,238 | 4,73E-08 |
| Sspon.01G0039430 | -18,813 | 4,73E-08 |
| Sspon.01G0039480 | 17,139 | 4,79E-08 |
| Sspon.01G0039510 | 2,049 | 4,85E-08 |
| Sspon.01G0039580 | 1,112 | 4,97E-08 |
| Sspon.01G0039590 | -2,391 | 5,08E-08 |

| | | |
|---|---|---|
| Sspon.01G0039640 | -2,967 | 5,11E-08 |
| Sspon.01G0039660 | 2,106 | 5,11E-08 |
| Sspon.01G0039670 | 1,470 | 5,11E-08 |
| Sspon.01G0039790 | 3,160 | 5,13E-08 |
| Sspon.01G0039840 | -1,003 | 5,17E-08 |
| Sspon.01G0039850 | 2,984 | 5,20E-08 |
| Sspon.01G0039870 | -4,243 | 5,22E-08 |
| Sspon.01G0039880 | 1,192 | 5,24E-08 |
| Sspon.01G0039900 | -3,565 | 5,37E-08 |
| Sspon.01G0039980 | 4,300 | 5,37E-08 |
| Sspon.01G0039990 | 2,437 | 5,47E-08 |
| Sspon.01G0040000 | 2,818 | 5,56E-08 |
| Sspon.01G0040040 | -1,147 | 5,57E-08 |
| Sspon.01G0040240 | 4,226 | 5,64E-08 |
| Sspon.01G0040330 | 1,007 | 5,70E-08 |
| Sspon.01G0040370 | 16,919 | 5,70E-08 |
| Sspon.01G0040400 | 17,149 | 5,71E-08 |
| Sspon.01G0040410 | -1,395 | 5,72E-08 |
| Sspon.01G0040490 | -1,584 | 5,72E-08 |
| Sspon.01G0040520 | -3,778 | 5,79E-08 |
| Sspon.01G0040530 | -2,621 | 5,79E-08 |
| Sspon.01G0040560 | 17,802 | 5,96E-08 |
| Sspon.01G0040670 | 1,715 | 5,99E-08 |
| Sspon.01G0040710 | -1,377 | 6,06E-08 |
| Sspon.01G0040790 | -2,762 | 6,07E-08 |
| Sspon.01G0040830 | -1,212 | 6,07E-08 |
| Sspon.01G0040850 | -1,578 | 6,17E-08 |
| Sspon.01G0041000 | -2,400 | 6,22E-08 |
| Sspon.01G0041090 | 1,789 | 6,24E-08 |
| Sspon.01G0041120 | 2,498 | 6,27E-08 |
| Sspon.01G0041150 | -2,030 | 6,28E-08 |
| Sspon.01G0041190 | 1,219 | 6,62E-08 |
| Sspon.01G0041280 | -2,179 | 6,63E-08 |
| Sspon.01G0041290 | 1,590 | 6,73E-08 |
| Sspon.01G0041490 | 2,270 | 6,85E-08 |
| Sspon.01G0041530 | -1,302 | 6,93E-08 |
| Sspon.01G0041550 | 1,631 | 6,94E-08 |
| Sspon.01G0041610 | -1,055 | 7,12E-08 |
| Sspon.01G0041670 | -2,470 | 7,15E-08 |
| Sspon.01G0041760 | 2,930 | 7,20E-08 |
| Sspon.01G0041860 | 16,899 | 7,22E-08 |
| Sspon.01G0041950 | -1,993 | 7,38E-08 |
| Sspon.01G0042090 | 2,016 | 7,42E-08 |

| | | |
|---|---|---|
| Sspon.01G0042230 | 1,798 | 7,52E-08 |
| Sspon.01G0042310 | 3,111 | 7,52E-08 |
| Sspon.01G0042430 | 3,286 | 7,64E-08 |
| Sspon.01G0042440 | -1,737 | 7,69E-08 |
| Sspon.01G0042480 | 16,279 | 7,69E-08 |
| Sspon.01G0042510 | 16,500 | 7,70E-08 |
| Sspon.01G0042540 | 4,887 | 7,78E-08 |
| Sspon.01G0042620 | 1,098 | 7,82E-08 |
| Sspon.01G0042750 | -1,405 | 7,86E-08 |
| Sspon.01G0042790 | -4,339 | 7,91E-08 |
| Sspon.01G0042830 | 2,706 | 7,92E-08 |
| Sspon.01G0042850 | 1,199 | 7,94E-08 |
| Sspon.01G0042860 | 1,209 | 7,94E-08 |
| Sspon.01G0042880 | -1,212 | 8,10E-08 |
| Sspon.01G0042890 | 5,179 | 8,10E-08 |
| Sspon.01G0042900 | -3,647 | 8,10E-08 |
| Sspon.01G0042920 | -6,814 | 8,19E-08 |
| Sspon.01G0042970 | 2,608 | 8,21E-08 |
| Sspon.01G0043000 | 1,864 | 8,31E-08 |
| Sspon.01G0043040 | -1,132 | 8,31E-08 |
| Sspon.01G0043120 | 17,213 | 8,33E-08 |
| Sspon.01G0043250 | -1,442 | 8,36E-08 |
| Sspon.01G0043330 | -1,031 | 8,44E-08 |
| Sspon.01G0043450 | 3,320 | 8,44E-08 |
| Sspon.01G0043460 | -1,142 | 8,49E-08 |
| Sspon.01G0043540 | 3,352 | 8,60E-08 |
| Sspon.01G0043590 | 3,872 | 8,67E-08 |
| Sspon.01G0043720 | 3,242 | 8,69E-08 |
| Sspon.01G0043750 | 1,879 | 8,69E-08 |
| Sspon.01G0043790 | 3,049 | 8,70E-08 |
| Sspon.01G0043820 | 4,663 | 8,71E-08 |
| Sspon.01G0043840 | -4,705 | 8,71E-08 |
| Sspon.01G0043890 | 3,756 | 8,79E-08 |
| Sspon.01G0043930 | -1,483 | 8,83E-08 |
| Sspon.01G0044010 | -1,597 | 8,85E-08 |
| Sspon.01G0044110 | -2,778 | 8,86E-08 |
| Sspon.01G0044260 | -18,560 | 8,89E-08 |
| Sspon.01G0044320 | 16,527 | 8,91E-08 |
| Sspon.01G0044630 | 2,279 | 9,04E-08 |
| Sspon.01G0044780 | 1,813 | 9,06E-08 |
| Sspon.01G0044940 | 2,284 | 9,10E-08 |
| Sspon.01G0045090 | 1,328 | 9,11E-08 |
| Sspon.01G0045150 | 1,549 | 9,12E-08 |

| | | |
|---|---|---|
| Sspon.01G0045160 | -1,608 | 9,14E-08 |
| Sspon.01G0045230 | -2,847 | 9,41E-08 |
| Sspon.01G0045250 | -1,065 | 9,53E-08 |
| Sspon.01G0045270 | -1,891 | 9,58E-08 |
| Sspon.01G0045340 | 6,220 | 9,58E-08 |
| Sspon.01G0045380 | -1,141 | 9,66E-08 |
| Sspon.01G0045520 | -2,553 | 9,66E-08 |
| Sspon.01G0045580 | 2,714 | 9,68E-08 |
| Sspon.01G0045630 | -2,164 | 9,84E-08 |
| Sspon.01G0045740 | 3,433 | 9,84E-08 |
| Sspon.01G0045850 | 2,968 | 1,00E-07 |
| Sspon.01G0045880 | 1,867 | 1,01E-07 |
| Sspon.01G0045910 | -1,271 | 1,02E-07 |
| Sspon.01G0045920 | 16,602 | 1,02E-07 |
| Sspon.01G0045980 | -1,374 | 1,02E-07 |
| Sspon.01G0046000 | -2,172 | 1,03E-07 |
| Sspon.01G0046050 | -1,348 | 1,04E-07 |
| Sspon.01G0046090 | -1,710 | 1,05E-07 |
| Sspon.01G0046110 | 15,841 | 1,05E-07 |
| Sspon.01G0046130 | 1,271 | 1,07E-07 |
| Sspon.01G0046150 | 18,141 | 1,08E-07 |
| Sspon.01G0046200 | 2,320 | 1,08E-07 |
| Sspon.01G0046220 | -2,132 | 1,09E-07 |
| Sspon.01G0046260 | -1,845 | 1,12E-07 |
| Sspon.01G0046430 | 1,409 | 1,12E-07 |
| Sspon.01G0046490 | 16,423 | 1,13E-07 |
| Sspon.01G0046500 | -1,314 | 1,14E-07 |
| Sspon.01G0046650 | 3,200 | 1,14E-07 |
| Sspon.01G0046960 | -2,061 | 1,14E-07 |
| Sspon.01G0047000 | 1,727 | 1,15E-07 |
| Sspon.01G0047030 | -3,454 | 1,16E-07 |
| Sspon.01G0047050 | 2,255 | 1,17E-07 |
| Sspon.01G0047090 | 1,272 | 1,17E-07 |
| Sspon.01G0047110 | 1,748 | 1,17E-07 |
| Sspon.01G0047140 | -19,378 | 1,18E-07 |
| Sspon.01G0047230 | -18,515 | 1,19E-07 |
| Sspon.01G0047330 | 2,014 | 1,19E-07 |
| Sspon.01G0047340 | -1,925 | 1,20E-07 |
| Sspon.01G0047350 | -1,510 | 1,21E-07 |
| Sspon.01G0047380 | 1,445 | 1,22E-07 |
| Sspon.01G0047430 | 2,690 | 1,23E-07 |
| Sspon.01G0047530 | 16,540 | 1,24E-07 |
| Sspon.01G0047560 | -1,806 | 1,25E-07 |

| | | |
|---|---|---|
| Sspon.01G0047900 | 1,145 | 1,25E-07 |
| Sspon.01G0047910 | 16,604 | 1,26E-07 |
| Sspon.01G0047940 | -1,423 | 1,26E-07 |
| Sspon.01G0048080 | -1,047 | 1,26E-07 |
| Sspon.01G0048090 | 6,063 | 1,27E-07 |
| Sspon.01G0048100 | 17,430 | 1,27E-07 |
| Sspon.01G0048170 | 1,819 | 1,28E-07 |
| Sspon.01G0048250 | 3,227 | 1,28E-07 |
| Sspon.01G0048300 | -2,170 | 1,29E-07 |
| Sspon.01G0048500 | -1,083 | 1,29E-07 |
| Sspon.01G0048520 | -1,010 | 1,29E-07 |
| Sspon.01G0048530 | -1,367 | 1,29E-07 |
| Sspon.01G0048600 | 1,836 | 1,30E-07 |
| Sspon.01G0048620 | -1,084 | 1,31E-07 |
| Sspon.01G0048790 | -1,596 | 1,32E-07 |
| Sspon.01G0048830 | -1,547 | 1,33E-07 |
| Sspon.01G0048890 | -1,075 | 1,33E-07 |
| Sspon.01G0048910 | 1,672 | 1,33E-07 |
| Sspon.01G0048980 | -1,218 | 1,34E-07 |
| Sspon.01G0049040 | 1,815 | 1,35E-07 |
| Sspon.01G0049050 | -1,000 | 1,35E-07 |
| Sspon.01G0049120 | 3,901 | 1,39E-07 |
| Sspon.01G0049330 | -1,215 | 1,40E-07 |
| Sspon.01G0049340 | -1,357 | 1,41E-07 |
| Sspon.01G0049400 | -18,568 | 1,41E-07 |
| Sspon.01G0049490 | -1,059 | 1,42E-07 |
| Sspon.01G0049590 | 1,918 | 1,44E-07 |
| Sspon.01G0049620 | 1,752 | 1,44E-07 |
| Sspon.01G0049670 | 4,176 | 1,45E-07 |
| Sspon.01G0049690 | 1,436 | 1,46E-07 |
| Sspon.01G0049780 | 16,216 | 1,47E-07 |
| Sspon.01G0049830 | 1,625 | 1,47E-07 |
| Sspon.01G0049870 | -1,033 | 1,47E-07 |
| Sspon.01G0049960 | -1,267 | 1,47E-07 |
| Sspon.01G0049990 | -1,369 | 1,48E-07 |
| Sspon.01G0050010 | -3,090 | 1,48E-07 |
| Sspon.01G0050040 | 2,687 | 1,48E-07 |
| Sspon.01G0050090 | 2,310 | 1,49E-07 |
| Sspon.01G0050220 | -1,364 | 1,49E-07 |
| Sspon.01G0050300 | 7,683 | 1,49E-07 |
| Sspon.01G0050310 | -1,654 | 1,52E-07 |
| Sspon.01G0050350 | 1,567 | 1,53E-07 |
| Sspon.01G0050390 | -1,188 | 1,53E-07 |

| | | |
|---|---|---|
| Sspon.01G0050410 | 1,132 | 1,56E-07 |
| Sspon.01G0050460 | 6,801 | 1,57E-07 |
| Sspon.01G0050470 | 1,632 | 1,60E-07 |
| Sspon.01G0050510 | 1,071 | 1,60E-07 |
| Sspon.01G0050560 | 5,064 | 1,62E-07 |
| Sspon.01G0050630 | -1,755 | 1,63E-07 |
| Sspon.01G0050660 | 2,441 | 1,64E-07 |
| Sspon.01G0050680 | 1,823 | 1,68E-07 |
| Sspon.01G0050750 | -2,017 | 1,71E-07 |
| Sspon.01G0050830 | 5,229 | 1,71E-07 |
| Sspon.01G0050880 | 1,247 | 1,72E-07 |
| Sspon.01G0050890 | -1,291 | 1,76E-07 |
| Sspon.01G0050960 | 18,344 | 1,76E-07 |
| Sspon.01G0050980 | -2,127 | 1,77E-07 |
| Sspon.01G0051050 | 16,484 | 1,80E-07 |
| Sspon.01G0051090 | 1,410 | 1,80E-07 |
| Sspon.01G0051130 | 16,872 | 1,81E-07 |
| Sspon.01G0051140 | 2,053 | 1,81E-07 |
| Sspon.01G0051180 | 2,721 | 1,81E-07 |
| Sspon.01G0051300 | 5,791 | 1,81E-07 |
| Sspon.01G0051340 | 4,750 | 1,81E-07 |
| Sspon.01G0051370 | -1,461 | 1,88E-07 |
| Sspon.01G0051420 | -2,147 | 1,91E-07 |
| Sspon.01G0051430 | 1,707 | 1,91E-07 |
| Sspon.01G0051460 | 1,895 | 1,94E-07 |
| Sspon.01G0051560 | -1,309 | 1,94E-07 |
| Sspon.01G0051570 | -1,572 | 1,95E-07 |
| Sspon.01G0051580 | -1,377 | 1,96E-07 |
| Sspon.01G0051690 | 17,771 | 1,96E-07 |
| Sspon.01G0051700 | -2,088 | 1,97E-07 |
| Sspon.01G0051720 | -1,054 | 1,97E-07 |
| Sspon.01G0051760 | 1,859 | 2,01E-07 |
| Sspon.01G0051810 | 17,825 | 2,03E-07 |
| Sspon.01G0051940 | 16,655 | 2,03E-07 |
| Sspon.01G0051950 | -1,118 | 2,03E-07 |
| Sspon.01G0051970 | 1,848 | 2,04E-07 |
| Sspon.01G0052010 | -1,494 | 2,06E-07 |
| Sspon.01G0052020 | 2,441 | 2,10E-07 |
| Sspon.01G0052030 | 3,171 | 2,11E-07 |
| Sspon.01G0052040 | 3,261 | 2,11E-07 |
| Sspon.01G0052120 | 1,154 | 2,11E-07 |
| Sspon.01G0052150 | -1,878 | 2,12E-07 |
| Sspon.01G0052190 | 2,679 | 2,14E-07 |

| | | |
|---|---|---|
| Sspon.01G0052220 | 3,362 | 2,16E-07 |
| Sspon.01G0052290 | 16,432 | 2,18E-07 |
| Sspon.01G0052320 | 3,728 | 2,21E-07 |
| Sspon.01G0052350 | 2,258 | 2,23E-07 |
| Sspon.01G0052450 | -2,925 | 2,23E-07 |
| Sspon.01G0052510 | 3,086 | 2,25E-07 |
| Sspon.01G0052540 | 1,450 | 2,25E-07 |
| Sspon.01G0052600 | -18,506 | 2,27E-07 |
| Sspon.01G0052610 | 2,708 | 2,27E-07 |
| Sspon.01G0052620 | -2,752 | 2,27E-07 |
| Sspon.01G0052690 | 1,088 | 2,27E-07 |
| Sspon.01G0052720 | 3,287 | 2,27E-07 |
| Sspon.01G0052730 | -2,261 | 2,29E-07 |
| Sspon.01G0052780 | 16,170 | 2,30E-07 |
| Sspon.01G0052940 | -1,163 | 2,31E-07 |
| Sspon.01G0052980 | -1,066 | 2,43E-07 |
| Sspon.01G0053040 | -1,153 | 2,44E-07 |
| Sspon.01G0053050 | -3,242 | 2,46E-07 |
| Sspon.01G0053120 | -1,633 | 2,49E-07 |
| Sspon.01G0053140 | 1,526 | 2,53E-07 |
| Sspon.01G0053150 | 2,298 | 2,54E-07 |
| Sspon.01G0053170 | -1,494 | 2,54E-07 |
| Sspon.01G0053200 | 3,584 | 2,56E-07 |
| Sspon.01G0053210 | -1,137 | 2,59E-07 |
| Sspon.01G0053410 | -1,199 | 2,61E-07 |
| Sspon.01G0053560 | 2,429 | 2,61E-07 |
| Sspon.01G0053610 | -1,486 | 2,63E-07 |
| Sspon.01G0053620 | -1,003 | 2,64E-07 |
| Sspon.01G0053630 | 4,237 | 2,66E-07 |
| Sspon.01G0053730 | 4,243 | 2,66E-07 |
| Sspon.01G0053820 | -1,310 | 2,66E-07 |
| Sspon.01G0053840 | 2,117 | 2,66E-07 |
| Sspon.01G0053860 | 1,096 | 2,67E-07 |
| Sspon.01G0053870 | -3,765 | 2,67E-07 |
| Sspon.01G0053950 | 3,668 | 2,69E-07 |
| Sspon.01G0054020 | 3,600 | 2,69E-07 |
| Sspon.01G0054110 | 16,353 | 2,69E-07 |
| Sspon.01G0054120 | 1,152 | 2,69E-07 |
| Sspon.01G0054140 | 1,716 | 2,69E-07 |
| Sspon.01G0054170 | -19,289 | 2,70E-07 |
| Sspon.01G0054190 | -1,341 | 2,73E-07 |
| Sspon.01G0054230 | 1,681 | 2,75E-07 |
| Sspon.01G0054280 | -2,447 | 2,75E-07 |

| | | |
|---|---|---|
| Sspon.01G0054320 | -1,622 | 2,78E-07 |
| Sspon.01G0054380 | 2,318 | 2,80E-07 |
| Sspon.01G0054440 | 16,169 | 2,82E-07 |
| Sspon.01G0054650 | -1,343 | 2,90E-07 |
| Sspon.01G0054700 | 1,708 | 2,91E-07 |
| Sspon.01G0054800 | 1,498 | 2,92E-07 |
| Sspon.01G0055050 | -1,013 | 2,92E-07 |
| Sspon.01G0055170 | 3,303 | 2,93E-07 |
| Sspon.01G0055210 | -2,207 | 2,94E-07 |
| Sspon.01G0055230 | 4,323 | 2,95E-07 |
| Sspon.01G0055320 | 3,190 | 2,98E-07 |
| Sspon.01G0055370 | -1,594 | 2,99E-07 |
| Sspon.01G0055390 | -2,058 | 3,00E-07 |
| Sspon.01G0055540 | -2,459 | 3,00E-07 |
| Sspon.01G0055650 | 3,049 | 3,01E-07 |
| Sspon.01G0055660 | 2,993 | 3,01E-07 |
| Sspon.01G0055820 | -1,339 | 3,01E-07 |
| Sspon.01G0055870 | 1,811 | 3,03E-07 |
| Sspon.01G0055930 | 2,255 | 3,03E-07 |
| Sspon.01G0055990 | 3,693 | 3,04E-07 |
| Sspon.01G0056010 | 1,999 | 3,04E-07 |
| Sspon.01G0056050 | -2,596 | 3,07E-07 |
| Sspon.01G0056310 | 1,036 | 3,10E-07 |
| Sspon.01G0056370 | 3,636 | 3,10E-07 |
| Sspon.01G0056460 | 2,767 | 3,11E-07 |
| Sspon.01G0056550 | 2,236 | 3,11E-07 |
| Sspon.01G0056600 | -1,748 | 3,12E-07 |
| Sspon.01G0056610 | 16,004 | 3,13E-07 |
| Sspon.01G0056630 | 3,104 | 3,14E-07 |
| Sspon.01G0056640 | -1,982 | 3,14E-07 |
| Sspon.01G0056740 | 2,471 | 3,15E-07 |
| Sspon.01G0056850 | -1,663 | 3,18E-07 |
| Sspon.01G0057040 | -1,817 | 3,18E-07 |
| Sspon.01G0057090 | 1,917 | 3,18E-07 |
| Sspon.01G0057330 | 2,436 | 3,19E-07 |
| Sspon.01G0057350 | -2,180 | 3,26E-07 |
| Sspon.01G0057420 | 4,124 | 3,28E-07 |
| Sspon.01G0057450 | -2,214 | 3,29E-07 |
| Sspon.01G0057510 | -2,051 | 3,31E-07 |
| Sspon.01G0057580 | -1,398 | 3,32E-07 |
| Sspon.01G0057660 | -2,547 | 3,34E-07 |
| Sspon.01G0057740 | 2,785 | 3,35E-07 |
| Sspon.01G0057760 | 2,642 | 3,42E-07 |

| | | |
|---|---|---|
| Sspon.01G0057770 | 1,038 | 3,42E-07 |
| Sspon.01G0057850 | -2,823 | 3,44E-07 |
| Sspon.01G0057870 | 3,427 | 3,44E-07 |
| Sspon.01G0057880 | 2,447 | 3,48E-07 |
| Sspon.01G0057950 | -1,921 | 3,48E-07 |
| Sspon.01G0057970 | 1,074 | 3,49E-07 |
| Sspon.01G0057990 | 1,394 | 3,49E-07 |
| Sspon.01G0058190 | -2,117 | 3,49E-07 |
| Sspon.01G0058290 | 18,221 | 3,49E-07 |
| Sspon.01G0058410 | -1,487 | 3,52E-07 |
| Sspon.01G0058430 | -1,238 | 3,53E-07 |
| Sspon.01G0058440 | 1,412 | 3,53E-07 |
| Sspon.01G0058810 | -3,678 | 3,62E-07 |
| Sspon.01G0058850 | 1,602 | 3,63E-07 |
| Sspon.01G0058860 | 1,027 | 3,64E-07 |
| Sspon.01G0058970 | -1,730 | 3,66E-07 |
| Sspon.01G0058980 | -1,035 | 3,66E-07 |
| Sspon.01G0059000 | 15,736 | 3,70E-07 |
| Sspon.01G0059080 | 2,075 | 3,73E-07 |
| Sspon.01G0059140 | 2,786 | 3,73E-07 |
| Sspon.01G0059200 | -3,686 | 3,73E-07 |
| Sspon.01G0059290 | 15,775 | 3,73E-07 |
| Sspon.01G0059300 | 15,878 | 3,78E-07 |
| Sspon.01G0059330 | 4,729 | 3,79E-07 |
| Sspon.01G0059400 | -1,492 | 3,81E-07 |
| Sspon.01G0059480 | -1,886 | 3,84E-07 |
| Sspon.01G0059590 | -1,324 | 3,95E-07 |
| Sspon.01G0059620 | -1,555 | 3,95E-07 |
| Sspon.01G0059630 | 5,729 | 3,96E-07 |
| Sspon.01G0059640 | -1,531 | 3,98E-07 |
| Sspon.01G0059680 | -1,524 | 3,98E-07 |
| Sspon.01G0059690 | 1,447 | 3,98E-07 |
| Sspon.01G0059720 | -1,233 | 3,98E-07 |
| Sspon.01G0059740 | 6,016 | 4,06E-07 |
| Sspon.01G0059790 | 4,838 | 4,07E-07 |
| Sspon.01G0059960 | 2,307 | 4,08E-07 |
| Sspon.01G0059970 | 16,747 | 4,10E-07 |
| Sspon.01G0060000 | 2,212 | 4,13E-07 |
| Sspon.01G0060030 | -1,611 | 4,15E-07 |
| Sspon.01G0060040 | 1,514 | 4,16E-07 |
| Sspon.01G0060110 | -2,172 | 4,18E-07 |
| Sspon.01G0060270 | 18,316 | 4,19E-07 |
| Sspon.01G0060440 | -1,489 | 4,19E-07 |

| | | |
|---|---|---|
| Sspon.01G0060540 | 4,466 | 4,19E-07 |
| Sspon.01G0060620 | -1,458 | 4,27E-07 |
| Sspon.01G0060760 | 3,403 | 4,32E-07 |
| Sspon.01G0060830 | 3,025 | 4,33E-07 |
| Sspon.01G0060930 | 18,777 | 4,33E-07 |
| Sspon.01G0061080 | 16,348 | 4,34E-07 |
| Sspon.01G0061120 | -2,293 | 4,36E-07 |
| Sspon.01G0061130 | 1,369 | 4,37E-07 |
| Sspon.01G0061190 | 1,425 | 4,40E-07 |
| Sspon.01G0061230 | 15,891 | 4,40E-07 |
| Sspon.01G0061350 | 21,442 | 4,44E-07 |
| Sspon.01G0061430 | 15,836 | 4,45E-07 |
| Sspon.01G0061450 | 1,656 | 4,49E-07 |
| Sspon.01G0061480 | 3,200 | 4,52E-07 |
| Sspon.01G0061610 | -13,600 | 4,57E-07 |
| Sspon.01G0061650 | -1,058 | 4,57E-07 |
| Sspon.01G0061660 | 5,119 | 4,58E-07 |
| Sspon.01G0061720 | 2,454 | 4,58E-07 |
| Sspon.01G0061730 | 1,127 | 4,60E-07 |
| Sspon.01G0061740 | -18,229 | 4,62E-07 |
| Sspon.01G0061750 | -1,117 | 4,63E-07 |
| Sspon.01G0062010 | 4,081 | 4,63E-07 |
| Sspon.01G0062090 | 3,485 | 4,64E-07 |
| Sspon.01G0062110 | -2,861 | 4,73E-07 |
| Sspon.01G0062280 | -1,343 | 4,81E-07 |
| Sspon.01G0062320 | -1,306 | 4,85E-07 |
| Sspon.01G0062330 | -2,094 | 4,88E-07 |
| Sspon.01G0062340 | 2,124 | 4,89E-07 |
| Sspon.01G0062470 | 1,795 | 4,90E-07 |
| Sspon.01G0062520 | -1,689 | 4,97E-07 |
| Sspon.01G0062640 | 2,011 | 4,99E-07 |
| Sspon.01G0062840 | 3,660 | 5,07E-07 |
| Sspon.01G0062880 | 2,192 | 5,07E-07 |
| Sspon.01G0062940 | -1,327 | 5,09E-07 |
| Sspon.01G0063120 | 2,085 | 5,13E-07 |
| Sspon.01G0063130 | 2,819 | 5,14E-07 |
| Sspon.01G0063170 | -2,307 | 5,28E-07 |
| Sspon.01G0063190 | 1,813 | 5,28E-07 |
| Sspon.01G0063290 | 16,814 | 5,35E-07 |
| Sspon.01G0063410 | -1,004 | 5,38E-07 |
| Sspon.01G0063420 | -2,358 | 5,39E-07 |
| Sspon.01G0063430 | 2,278 | 5,42E-07 |
| Sspon.01G0063440 | 2,156 | 5,48E-07 |

| | | |
|---|---|---|
| Sspon.01G0063500 | 1,919 | 5,49E-07 |
| Sspon.02G0000010 | 17,135 | 5,49E-07 |
| Sspon.02G0000040 | -2,023 | 5,49E-07 |
| Sspon.02G0000050 | 3,222 | 5,53E-07 |
| Sspon.02G0000080 | 2,688 | 5,54E-07 |
| Sspon.02G0000180 | 2,670 | 5,56E-07 |
| Sspon.02G0000470 | -1,670 | 5,57E-07 |
| Sspon.02G0000550 | 1,994 | 5,65E-07 |
| Sspon.02G0000570 | 15,851 | 5,68E-07 |
| Sspon.02G0000600 | -3,748 | 5,69E-07 |
| Sspon.02G0000610 | 1,564 | 5,75E-07 |
| Sspon.02G0000740 | 1,124 | 5,79E-07 |
| Sspon.02G0000760 | 16,283 | 5,79E-07 |
| Sspon.02G0000840 | 4,263 | 5,84E-07 |
| Sspon.02G0000860 | -2,462 | 5,85E-07 |
| Sspon.02G0000870 | -4,280 | 5,85E-07 |
| Sspon.02G0000900 | 1,367 | 5,86E-07 |
| Sspon.02G0000950 | -1,755 | 5,86E-07 |
| Sspon.02G0000960 | -2,813 | 5,88E-07 |
| Sspon.02G0000970 | 2,933 | 5,96E-07 |
| Sspon.02G0001010 | -3,009 | 5,97E-07 |
| Sspon.02G0001050 | 1,955 | 5,99E-07 |
| Sspon.02G0001060 | -1,220 | 6,00E-07 |
| Sspon.02G0001160 | -1,889 | 6,00E-07 |
| Sspon.02G0001190 | -1,719 | 6,00E-07 |
| Sspon.02G0001250 | 16,023 | 6,04E-07 |
| Sspon.02G0001280 | -1,574 | 6,05E-07 |
| Sspon.02G0001290 | -2,847 | 6,07E-07 |
| Sspon.02G0001360 | -1,735 | 6,15E-07 |
| Sspon.02G0001440 | -3,159 | 6,18E-07 |
| Sspon.02G0001450 | -1,059 | 6,21E-07 |
| Sspon.02G0001540 | 2,003 | 6,26E-07 |
| Sspon.02G0001590 | 1,608 | 6,39E-07 |
| Sspon.02G0001720 | -2,796 | 6,54E-07 |
| Sspon.02G0001750 | -1,317 | 6,54E-07 |
| Sspon.02G0001760 | 1,340 | 6,56E-07 |
| Sspon.02G0001930 | -2,514 | 6,61E-07 |
| Sspon.02G0001940 | 2,526 | 6,64E-07 |
| Sspon.02G0001960 | 3,305 | 6,68E-07 |
| Sspon.02G0001980 | -2,144 | 6,68E-07 |
| Sspon.02G0002010 | -2,788 | 6,69E-07 |
| Sspon.02G0002020 | 15,674 | 6,70E-07 |
| Sspon.02G0002130 | 18,238 | 6,75E-07 |

| | | |
|---|---|---|
| Sspon.02G0002220 | 1,459 | 6,76E-07 |
| Sspon.02G0002240 | 2,056 | 6,76E-07 |
| Sspon.02G0002300 | 3,123 | 6,78E-07 |
| Sspon.02G0002360 | -1,838 | 6,83E-07 |
| Sspon.02G0002510 | -18,189 | 6,84E-07 |
| Sspon.02G0002520 | 1,881 | 6,88E-07 |
| Sspon.02G0002560 | -1,088 | 6,91E-07 |
| Sspon.02G0002630 | 2,064 | 6,94E-07 |
| Sspon.02G0002750 | 2,281 | 7,00E-07 |
| Sspon.02G0002830 | 4,195 | 7,00E-07 |
| Sspon.02G0002840 | -1,542 | 7,02E-07 |
| Sspon.02G0002890 | 2,175 | 7,14E-07 |
| Sspon.02G0002990 | -1,791 | 7,14E-07 |
| Sspon.02G0003080 | -1,619 | 7,18E-07 |
| Sspon.02G0003130 | 1,717 | 7,32E-07 |
| Sspon.02G0003140 | 12,390 | 7,35E-07 |
| Sspon.02G0003160 | -1,736 | 7,40E-07 |
| Sspon.02G0003230 | 1,675 | 7,41E-07 |
| Sspon.02G0003280 | -1,602 | 7,41E-07 |
| Sspon.02G0003340 | 1,281 | 7,49E-07 |
| Sspon.02G0003350 | 3,517 | 7,50E-07 |
| Sspon.02G0003370 | 1,415 | 7,50E-07 |
| Sspon.02G0003390 | 2,898 | 7,50E-07 |
| Sspon.02G0003500 | 1,990 | 7,58E-07 |
| Sspon.02G0003590 | -1,079 | 7,58E-07 |
| Sspon.02G0003600 | 1,472 | 7,62E-07 |
| Sspon.02G0003650 | -2,028 | 7,65E-07 |
| Sspon.02G0003710 | 3,151 | 7,65E-07 |
| Sspon.02G0003740 | -1,249 | 7,73E-07 |
| Sspon.02G0003810 | 1,116 | 7,84E-07 |
| Sspon.02G0003820 | 1,289 | 7,86E-07 |
| Sspon.02G0003830 | 4,962 | 7,87E-07 |
| Sspon.02G0003880 | -1,026 | 7,89E-07 |
| Sspon.02G0003910 | -2,377 | 7,90E-07 |
| Sspon.02G0003920 | 2,282 | 8,10E-07 |
| Sspon.02G0003970 | 1,764 | 8,19E-07 |
| Sspon.02G0004090 | -1,496 | 8,21E-07 |
| Sspon.02G0004160 | -1,470 | 8,23E-07 |
| Sspon.02G0004200 | -1,871 | 8,24E-07 |
| Sspon.02G0004290 | -1,026 | 8,24E-07 |
| Sspon.02G0004300 | -1,093 | 8,27E-07 |
| Sspon.02G0004320 | 2,135 | 8,28E-07 |
| Sspon.02G0004350 | 3,256 | 8,30E-07 |

| | | |
|---|---|---|
| Sspon.02G0004390 | 1,745 | 8,30E-07 |
| Sspon.02G0004560 | -1,211 | 8,30E-07 |
| Sspon.02G0004610 | 1,397 | 8,31E-07 |
| Sspon.02G0004660 | -1,952 | 8,32E-07 |
| Sspon.02G0004670 | 1,183 | 8,32E-07 |
| Sspon.02G0004680 | 2,033 | 8,39E-07 |
| Sspon.02G0004700 | -1,270 | 8,46E-07 |
| Sspon.02G0004810 | -1,224 | 8,50E-07 |
| Sspon.02G0004830 | -1,071 | 8,50E-07 |
| Sspon.02G0004870 | -1,061 | 8,53E-07 |
| Sspon.02G0004900 | -2,406 | 8,54E-07 |
| Sspon.02G0004920 | 2,757 | 8,54E-07 |
| Sspon.02G0004930 | 2,061 | 8,55E-07 |
| Sspon.02G0005110 | 1,963 | 8,60E-07 |
| Sspon.02G0005230 | 5,232 | 8,60E-07 |
| Sspon.02G0005240 | 2,643 | 8,61E-07 |
| Sspon.02G0005320 | -20,074 | 8,84E-07 |
| Sspon.02G0005350 | -1,958 | 8,84E-07 |
| Sspon.02G0005490 | 1,751 | 8,85E-07 |
| Sspon.02G0005570 | 1,673 | 8,95E-07 |
| Sspon.02G0005610 | 16,641 | 8,96E-07 |
| Sspon.02G0005650 | -1,070 | 9,09E-07 |
| Sspon.02G0005770 | 1,314 | 9,10E-07 |
| Sspon.02G0005840 | -1,846 | 9,11E-07 |
| Sspon.02G0005900 | 3,495 | 9,12E-07 |
| Sspon.02G0006060 | 1,891 | 9,23E-07 |
| Sspon.02G0006100 | -1,849 | 9,27E-07 |
| Sspon.02G0006160 | 1,536 | 9,27E-07 |
| Sspon.02G0006200 | -4,141 | 9,31E-07 |
| Sspon.02G0006440 | -2,219 | 9,32E-07 |
| Sspon.02G0006530 | 2,127 | 9,33E-07 |
| Sspon.02G0006580 | -1,420 | 9,45E-07 |
| Sspon.02G0006660 | -1,297 | 9,47E-07 |
| Sspon.02G0006690 | 1,861 | 9,52E-07 |
| Sspon.02G0006790 | -2,021 | 9,54E-07 |
| Sspon.02G0006820 | -3,240 | 9,56E-07 |
| Sspon.02G0006860 | 15,588 | 9,57E-07 |
| Sspon.02G0006970 | -1,581 | 9,57E-07 |
| Sspon.02G0006990 | -2,619 | 9,62E-07 |
| Sspon.02G0007000 | 1,165 | 9,62E-07 |
| Sspon.02G0007040 | 7,596 | 9,64E-07 |
| Sspon.02G0007160 | -1,418 | 9,66E-07 |
| Sspon.02G0007270 | 2,061 | 9,67E-07 |

| | | |
|---|---|---|
| Sspon.02G0007360 | -2,428 | 9,78E-07 |
| Sspon.02G0007370 | 3,433 | 9,78E-07 |
| Sspon.02G0007380 | 2,103 | 9,84E-07 |
| Sspon.02G0007440 | 4,036 | 9,89E-07 |
| Sspon.02G0007510 | 1,710 | 9,96E-07 |
| Sspon.02G0007590 | 2,684 | 9,98E-07 |
| Sspon.02G0007680 | 2,874 | 9,99E-07 |
| Sspon.02G0007780 | 1,794 | 1,00E-06 |
| Sspon.02G0008000 | 2,024 | 1,00E-06 |
| Sspon.02G0008010 | -1,325 | 1,01E-06 |
| Sspon.02G0008030 | 1,413 | 1,01E-06 |
| Sspon.02G0008100 | 2,363 | 1,02E-06 |
| Sspon.02G0008120 | 4,009 | 1,02E-06 |
| Sspon.02G0008230 | 1,570 | 1,03E-06 |
| Sspon.02G0008240 | 2,065 | 1,03E-06 |
| Sspon.02G0008260 | -1,289 | 1,04E-06 |
| Sspon.02G0008370 | 2,674 | 1,04E-06 |
| Sspon.02G0008390 | -1,215 | 1,06E-06 |
| Sspon.02G0008480 | 1,668 | 1,06E-06 |
| Sspon.02G0008490 | 1,819 | 1,06E-06 |
| Sspon.02G0008520 | -1,848 | 1,07E-06 |
| Sspon.02G0008530 | -5,150 | 1,08E-06 |
| Sspon.02G0008590 | 2,110 | 1,09E-06 |
| Sspon.02G0008710 | -1,022 | 1,10E-06 |
| Sspon.02G0008820 | -1,281 | 1,10E-06 |
| Sspon.02G0008910 | 3,450 | 1,10E-06 |
| Sspon.02G0008930 | -1,278 | 1,11E-06 |
| Sspon.02G0008970 | 1,698 | 1,11E-06 |
| Sspon.02G0009050 | 1,434 | 1,12E-06 |
| Sspon.02G0009070 | -18,056 | 1,13E-06 |
| Sspon.02G0009090 | -3,290 | 1,13E-06 |
| Sspon.02G0009100 | 1,640 | 1,15E-06 |
| Sspon.02G0009220 | 1,021 | 1,15E-06 |
| Sspon.02G0009250 | 1,801 | 1,15E-06 |
| Sspon.02G0009380 | 2,463 | 1,17E-06 |
| Sspon.02G0009520 | 2,495 | 1,17E-06 |
| Sspon.02G0009590 | 17,313 | 1,17E-06 |
| Sspon.02G0009610 | 3,063 | 1,17E-06 |
| Sspon.02G0009650 | -1,884 | 1,17E-06 |
| Sspon.02G0009680 | 1,529 | 1,18E-06 |
| Sspon.02G0009690 | -1,509 | 1,18E-06 |
| Sspon.02G0009740 | -17,670 | 1,18E-06 |
| Sspon.02G0009750 | -1,933 | 1,18E-06 |

| | | |
|---|---|---|
| Sspon.02G0009810 | -1,167 | 1,19E-06 |
| Sspon.02G0009940 | 2,017 | 1,19E-06 |
| Sspon.02G0010060 | -2,745 | 1,20E-06 |
| Sspon.02G0010150 | -2,242 | 1,20E-06 |
| Sspon.02G0010180 | 1,484 | 1,21E-06 |
| Sspon.02G0010230 | -2,113 | 1,21E-06 |
| Sspon.02G0010240 | -1,383 | 1,22E-06 |
| Sspon.02G0010270 | -1,667 | 1,22E-06 |
| Sspon.02G0010320 | -1,615 | 1,22E-06 |
| Sspon.02G0010430 | 2,233 | 1,22E-06 |
| Sspon.02G0010480 | 15,853 | 1,22E-06 |
| Sspon.02G0010500 | -2,333 | 1,23E-06 |
| Sspon.02G0010590 | 3,124 | 1,23E-06 |
| Sspon.02G0010630 | 3,014 | 1,23E-06 |
| Sspon.02G0010710 | -2,414 | 1,23E-06 |
| Sspon.02G0010850 | -1,185 | 1,23E-06 |
| Sspon.02G0011050 | 3,061 | 1,27E-06 |
| Sspon.02G0011090 | 2,303 | 1,28E-06 |
| Sspon.02G0011110 | 1,987 | 1,28E-06 |
| Sspon.02G0011130 | -2,625 | 1,28E-06 |
| Sspon.02G0011160 | 16,328 | 1,29E-06 |
| Sspon.02G0011250 | -2,379 | 1,31E-06 |
| Sspon.02G0011310 | 2,133 | 1,33E-06 |
| Sspon.02G0011360 | 1,580 | 1,34E-06 |
| Sspon.02G0011390 | 1,108 | 1,34E-06 |
| Sspon.02G0011420 | 1,053 | 1,34E-06 |
| Sspon.02G0011470 | 1,347 | 1,34E-06 |
| Sspon.02G0011480 | 2,030 | 1,35E-06 |
| Sspon.02G0011490 | -1,371 | 1,35E-06 |
| Sspon.02G0011500 | 1,908 | 1,35E-06 |
| Sspon.02G0011510 | 1,795 | 1,36E-06 |
| Sspon.02G0011610 | -1,071 | 1,36E-06 |
| Sspon.02G0011620 | -2,913 | 1,36E-06 |
| Sspon.02G0011650 | -1,558 | 1,37E-06 |
| Sspon.02G0011660 | 1,036 | 1,37E-06 |
| Sspon.02G0011700 | 5,816 | 1,39E-06 |
| Sspon.02G0011710 | 1,387 | 1,40E-06 |
| Sspon.02G0011730 | 1,553 | 1,40E-06 |
| Sspon.02G0011750 | -1,624 | 1,41E-06 |
| Sspon.02G0011790 | 2,091 | 1,41E-06 |
| Sspon.02G0011800 | -2,462 | 1,41E-06 |
| Sspon.02G0011810 | -1,408 | 1,42E-06 |
| Sspon.02G0011870 | 3,975 | 1,42E-06 |

| | | |
|---|---|---|
| Sspon.02G0011890 | -2,135 | 1,42E-06 |
| Sspon.02G0011940 | 1,612 | 1,42E-06 |
| Sspon.02G0012000 | 1,376 | 1,42E-06 |
| Sspon.02G0012040 | -1,126 | 1,42E-06 |
| Sspon.02G0012190 | 15,788 | 1,42E-06 |
| Sspon.02G0012260 | -2,155 | 1,43E-06 |
| Sspon.02G0012270 | -1,775 | 1,43E-06 |
| Sspon.02G0012300 | -1,485 | 1,43E-06 |
| Sspon.02G0012320 | -1,627 | 1,44E-06 |
| Sspon.02G0012360 | -1,740 | 1,44E-06 |
| Sspon.02G0012390 | 1,064 | 1,47E-06 |
| Sspon.02G0012420 | -2,579 | 1,47E-06 |
| Sspon.02G0012490 | 1,424 | 1,48E-06 |
| Sspon.02G0012550 | -3,550 | 1,48E-06 |
| Sspon.02G0012630 | 1,891 | 1,48E-06 |
| Sspon.02G0012660 | 1,191 | 1,49E-06 |
| Sspon.02G0012730 | 3,369 | 1,51E-06 |
| Sspon.02G0012750 | -17,556 | 1,51E-06 |
| Sspon.02G0012770 | -1,611 | 1,52E-06 |
| Sspon.02G0012890 | 1,810 | 1,53E-06 |
| Sspon.02G0013210 | 1,596 | 1,53E-06 |
| Sspon.02G0013240 | -1,232 | 1,54E-06 |
| Sspon.02G0013260 | 1,904 | 1,54E-06 |
| Sspon.02G0013290 | 5,027 | 1,55E-06 |
| Sspon.02G0013350 | -1,412 | 1,55E-06 |
| Sspon.02G0013480 | -1,788 | 1,56E-06 |
| Sspon.02G0013520 | 2,510 | 1,57E-06 |
| Sspon.02G0013580 | 1,779 | 1,58E-06 |
| Sspon.02G0013700 | -1,699 | 1,58E-06 |
| Sspon.02G0013710 | -1,323 | 1,59E-06 |
| Sspon.02G0013750 | -3,579 | 1,60E-06 |
| Sspon.02G0013840 | -1,657 | 1,61E-06 |
| Sspon.02G0013870 | -18,439 | 1,62E-06 |
| Sspon.02G0013880 | 3,194 | 1,65E-06 |
| Sspon.02G0013900 | 6,965 | 1,66E-06 |
| Sspon.02G0014000 | 1,040 | 1,68E-06 |
| Sspon.02G0014030 | 2,756 | 1,68E-06 |
| Sspon.02G0014070 | -1,363 | 1,68E-06 |
| Sspon.02G0014120 | -1,234 | 1,68E-06 |
| Sspon.02G0014130 | 1,557 | 1,70E-06 |
| Sspon.02G0014210 | 3,922 | 1,73E-06 |
| Sspon.02G0014220 | -1,375 | 1,75E-06 |
| Sspon.02G0014330 | 1,646 | 1,77E-06 |

| | | |
|---|---|---|
| Sspon.02G0014360 | -2,386 | 1,77E-06 |
| Sspon.02G0014370 | 1,964 | 1,77E-06 |
| Sspon.02G0014420 | -2,772 | 1,78E-06 |
| Sspon.02G0014510 | 3,744 | 1,78E-06 |
| Sspon.02G0014520 | 2,297 | 1,78E-06 |
| Sspon.02G0014530 | 1,151 | 1,79E-06 |
| Sspon.02G0014600 | -1,650 | 1,79E-06 |
| Sspon.02G0014660 | 1,518 | 1,80E-06 |
| Sspon.02G0014680 | -2,700 | 1,82E-06 |
| Sspon.02G0014700 | 1,221 | 1,83E-06 |
| Sspon.02G0014770 | 3,157 | 1,84E-06 |
| Sspon.02G0014800 | 1,331 | 1,85E-06 |
| Sspon.02G0014840 | -1,379 | 1,85E-06 |
| Sspon.02G0014850 | -2,985 | 1,85E-06 |
| Sspon.02G0014930 | -1,634 | 1,86E-06 |
| Sspon.02G0014950 | -4,330 | 1,86E-06 |
| Sspon.02G0015000 | 1,121 | 1,87E-06 |
| Sspon.02G0015010 | 2,193 | 1,87E-06 |
| Sspon.02G0015060 | 1,941 | 1,88E-06 |
| Sspon.02G0015070 | 16,549 | 1,88E-06 |
| Sspon.02G0015090 | 2,376 | 1,89E-06 |
| Sspon.02G0015140 | 1,426 | 1,90E-06 |
| Sspon.02G0015160 | 1,386 | 1,91E-06 |
| Sspon.02G0015210 | -1,706 | 1,93E-06 |
| Sspon.02G0015320 | -1,666 | 1,93E-06 |
| Sspon.02G0015340 | 3,823 | 1,93E-06 |
| Sspon.02G0015350 | -1,216 | 1,93E-06 |
| Sspon.02G0015360 | -1,570 | 1,96E-06 |
| Sspon.02G0015410 | -4,577 | 1,97E-06 |
| Sspon.02G0015420 | 1,396 | 2,00E-06 |
| Sspon.02G0015460 | -2,924 | 2,00E-06 |
| Sspon.02G0015500 | -2,447 | 2,00E-06 |
| Sspon.02G0015510 | -1,339 | 2,03E-06 |
| Sspon.02G0015550 | -1,743 | 2,03E-06 |
| Sspon.02G0015570 | -1,192 | 2,07E-06 |
| Sspon.02G0015610 | 1,857 | 2,07E-06 |
| Sspon.02G0015690 | 1,468 | 2,08E-06 |
| Sspon.02G0015760 | 4,131 | 2,09E-06 |
| Sspon.02G0015870 | -2,269 | 2,09E-06 |
| Sspon.02G0015890 | 4,632 | 2,11E-06 |
| Sspon.02G0015910 | -6,230 | 2,11E-06 |
| Sspon.02G0015970 | 1,022 | 2,12E-06 |
| Sspon.02G0016030 | 2,374 | 2,12E-06 |

| | | |
|---|---|---|
| Sspon.02G0016040 | -1,235 | 2,13E-06 |
| Sspon.02G0016150 | -1,321 | 2,14E-06 |
| Sspon.02G0016300 | 3,475 | 2,15E-06 |
| Sspon.02G0016360 | 1,166 | 2,15E-06 |
| Sspon.02G0016370 | 6,590 | 2,15E-06 |
| Sspon.02G0016380 | 1,729 | 2,15E-06 |
| Sspon.02G0016390 | 1,514 | 2,15E-06 |
| Sspon.02G0016400 | -1,073 | 2,16E-06 |
| Sspon.02G0016450 | 15,425 | 2,18E-06 |
| Sspon.02G0016570 | -4,584 | 2,19E-06 |
| Sspon.02G0016670 | 9,708 | 2,21E-06 |
| Sspon.02G0016730 | 10,025 | 2,21E-06 |
| Sspon.02G0016770 | 1,359 | 2,22E-06 |
| Sspon.02G0016810 | 1,190 | 2,22E-06 |
| Sspon.02G0016850 | -2,021 | 2,25E-06 |
| Sspon.02G0016940 | 2,615 | 2,26E-06 |
| Sspon.02G0016950 | -2,316 | 2,26E-06 |
| Sspon.02G0017000 | 1,696 | 2,26E-06 |
| Sspon.02G0017080 | 1,392 | 2,26E-06 |
| Sspon.02G0017090 | 2,553 | 2,28E-06 |
| Sspon.02G0017220 | 3,108 | 2,28E-06 |
| Sspon.02G0017280 | 2,781 | 2,29E-06 |
| Sspon.02G0017370 | 2,516 | 2,29E-06 |
| Sspon.02G0017460 | 2,099 | 2,30E-06 |
| Sspon.02G0017490 | 1,182 | 2,30E-06 |
| Sspon.02G0017520 | -1,481 | 2,31E-06 |
| Sspon.02G0017580 | -1,431 | 2,31E-06 |
| Sspon.02G0017670 | 1,322 | 2,31E-06 |
| Sspon.02G0017700 | 4,796 | 2,34E-06 |
| Sspon.02G0017740 | 6,716 | 2,34E-06 |
| Sspon.02G0017750 | 1,840 | 2,36E-06 |
| Sspon.02G0017760 | 1,716 | 2,37E-06 |
| Sspon.02G0017780 | -1,031 | 2,37E-06 |
| Sspon.02G0017890 | 15,390 | 2,39E-06 |
| Sspon.02G0017940 | 1,761 | 2,41E-06 |
| Sspon.02G0018000 | -1,445 | 2,41E-06 |
| Sspon.02G0018030 | 7,893 | 2,41E-06 |
| Sspon.02G0018160 | -1,918 | 2,42E-06 |
| Sspon.02G0018210 | -1,620 | 2,42E-06 |
| Sspon.02G0018220 | -1,002 | 2,44E-06 |
| Sspon.02G0018280 | 1,163 | 2,50E-06 |
| Sspon.02G0018300 | 2,561 | 2,50E-06 |
| Sspon.02G0018310 | 6,417 | 2,52E-06 |

| | | |
|---|---|---|
| Sspon.02G0018430 | -1,258 | 2,52E-06 |
| Sspon.02G0018450 | -1,259 | 2,52E-06 |
| Sspon.02G0018530 | 9,902 | 2,53E-06 |
| Sspon.02G0018540 | 4,191 | 2,53E-06 |
| Sspon.02G0018550 | 3,387 | 2,54E-06 |
| Sspon.02G0018560 | -2,296 | 2,55E-06 |
| Sspon.02G0018610 | -2,154 | 2,56E-06 |
| Sspon.02G0018770 | -1,519 | 2,56E-06 |
| Sspon.02G0018980 | 4,246 | 2,56E-06 |
| Sspon.02G0019060 | -2,930 | 2,56E-06 |
| Sspon.02G0019120 | 1,830 | 2,59E-06 |
| Sspon.02G0019150 | 1,450 | 2,61E-06 |
| Sspon.02G0019180 | 3,237 | 2,61E-06 |
| Sspon.02G0019230 | -7,924 | 2,61E-06 |
| Sspon.02G0019280 | -1,195 | 2,62E-06 |
| Sspon.02G0019410 | 3,145 | 2,63E-06 |
| Sspon.02G0019440 | 2,781 | 2,65E-06 |
| Sspon.02G0019480 | 2,850 | 2,68E-06 |
| Sspon.02G0019500 | -4,815 | 2,69E-06 |
| Sspon.02G0019530 | -1,836 | 2,74E-06 |
| Sspon.02G0019660 | -1,642 | 2,77E-06 |
| Sspon.02G0019740 | 1,402 | 2,78E-06 |
| Sspon.02G0019810 | -1,826 | 2,80E-06 |
| Sspon.02G0019890 | -1,043 | 2,81E-06 |
| Sspon.02G0019970 | 1,369 | 2,81E-06 |
| Sspon.02G0020040 | 2,143 | 2,83E-06 |
| Sspon.02G0020300 | -1,983 | 2,83E-06 |
| Sspon.02G0020370 | -1,502 | 2,84E-06 |
| Sspon.02G0020500 | -1,046 | 2,86E-06 |
| Sspon.02G0020590 | 1,123 | 2,87E-06 |
| Sspon.02G0020600 | 3,486 | 2,87E-06 |
| Sspon.02G0020630 | -1,057 | 2,88E-06 |
| Sspon.02G0020700 | 3,920 | 2,89E-06 |
| Sspon.02G0020760 | 3,204 | 2,92E-06 |
| Sspon.02G0020920 | -1,732 | 2,93E-06 |
| Sspon.02G0020940 | -1,307 | 2,93E-06 |
| Sspon.02G0020970 | 1,471 | 2,94E-06 |
| Sspon.02G0021000 | 2,662 | 2,94E-06 |
| Sspon.02G0021050 | 6,001 | 2,95E-06 |
| Sspon.02G0021070 | -1,330 | 2,97E-06 |
| Sspon.02G0021230 | -1,118 | 2,98E-06 |
| Sspon.02G0021270 | 2,106 | 2,98E-06 |
| Sspon.02G0021300 | 1,144 | 2,99E-06 |

| | | |
|---|---|---|
| Sspon.02G0021310 | 8,310 | 3,00E-06 |
| Sspon.02G0021420 | -4,789 | 3,00E-06 |
| Sspon.02G0021510 | -2,496 | 3,00E-06 |
| Sspon.02G0021530 | -1,940 | 3,00E-06 |
| Sspon.02G0021540 | -1,886 | 3,03E-06 |
| Sspon.02G0021560 | 12,330 | 3,04E-06 |
| Sspon.02G0021650 | 1,560 | 3,06E-06 |
| Sspon.02G0021820 | 3,056 | 3,06E-06 |
| Sspon.02G0022000 | 1,072 | 3,07E-06 |
| Sspon.02G0022060 | 1,823 | 3,07E-06 |
| Sspon.02G0022180 | 15,508 | 3,08E-06 |
| Sspon.02G0022250 | 1,892 | 3,08E-06 |
| Sspon.02G0022330 | 4,181 | 3,10E-06 |
| Sspon.02G0022400 | -1,427 | 3,14E-06 |
| Sspon.02G0022420 | 16,725 | 3,14E-06 |
| Sspon.02G0022440 | 2,458 | 3,14E-06 |
| Sspon.02G0022450 | 16,360 | 3,15E-06 |
| Sspon.02G0022530 | 2,459 | 3,19E-06 |
| Sspon.02G0022560 | 1,998 | 3,19E-06 |
| Sspon.02G0022570 | -17,344 | 3,19E-06 |
| Sspon.02G0022600 | 4,270 | 3,19E-06 |
| Sspon.02G0022680 | 1,051 | 3,19E-06 |
| Sspon.02G0022750 | 1,051 | 3,19E-06 |
| Sspon.02G0022850 | 1,354 | 3,22E-06 |
| Sspon.02G0022900 | -1,855 | 3,24E-06 |
| Sspon.02G0022930 | 1,111 | 3,24E-06 |
| Sspon.02G0023090 | -2,118 | 3,24E-06 |
| Sspon.02G0023120 | 7,783 | 3,26E-06 |
| Sspon.02G0023140 | -1,421 | 3,31E-06 |
| Sspon.02G0023270 | -1,383 | 3,31E-06 |
| Sspon.02G0023300 | 1,943 | 3,31E-06 |
| Sspon.02G0023350 | -1,811 | 3,32E-06 |
| Sspon.02G0023370 | 3,078 | 3,33E-06 |
| Sspon.02G0023450 | 5,384 | 3,33E-06 |
| Sspon.02G0023530 | 1,225 | 3,35E-06 |
| Sspon.02G0023630 | 15,341 | 3,38E-06 |
| Sspon.02G0023750 | 5,328 | 3,38E-06 |
| Sspon.02G0023870 | -17,242 | 3,40E-06 |
| Sspon.02G0023920 | -1,622 | 3,40E-06 |
| Sspon.02G0023970 | 7,452 | 3,41E-06 |
| Sspon.02G0023980 | 1,832 | 3,41E-06 |
| Sspon.02G0023990 | 2,185 | 3,44E-06 |
| Sspon.02G0024030 | 2,279 | 3,44E-06 |

| | | |
|---|---|---|
| Sspon.02G0024040 | -1,347 | 3,44E-06 |
| Sspon.02G0024060 | 1,260 | 3,46E-06 |
| Sspon.02G0024090 | 7,616 | 3,46E-06 |
| Sspon.02G0024140 | -2,385 | 3,46E-06 |
| Sspon.02G0024150 | -3,070 | 3,50E-06 |
| Sspon.02G0024180 | -3,587 | 3,51E-06 |
| Sspon.02G0024300 | 1,785 | 3,53E-06 |
| Sspon.02G0024390 | -2,181 | 3,54E-06 |
| Sspon.02G0024450 | 1,674 | 3,54E-06 |
| Sspon.02G0024540 | -1,033 | 3,55E-06 |
| Sspon.02G0024660 | 17,026 | 3,55E-06 |
| Sspon.02G0024740 | 2,490 | 3,62E-06 |
| Sspon.02G0024820 | 2,550 | 3,64E-06 |
| Sspon.02G0024830 | 2,574 | 3,64E-06 |
| Sspon.02G0024850 | 3,679 | 3,64E-06 |
| Sspon.02G0024860 | -1,307 | 3,66E-06 |
| Sspon.02G0024870 | 1,143 | 3,67E-06 |
| Sspon.02G0024880 | -1,146 | 3,67E-06 |
| Sspon.02G0024890 | -1,705 | 3,69E-06 |
| Sspon.02G0024970 | 2,952 | 3,72E-06 |
| Sspon.02G0025010 | -1,324 | 3,72E-06 |
| Sspon.02G0025070 | -1,060 | 3,74E-06 |
| Sspon.02G0025100 | 1,873 | 3,74E-06 |
| Sspon.02G0025120 | -1,923 | 3,75E-06 |
| Sspon.02G0025210 | 1,278 | 3,77E-06 |
| Sspon.02G0025250 | 1,531 | 3,78E-06 |
| Sspon.02G0025290 | 2,291 | 3,79E-06 |
| Sspon.02G0025300 | 1,553 | 3,80E-06 |
| Sspon.02G0025310 | 1,153 | 3,80E-06 |
| Sspon.02G0025330 | -17,327 | 3,81E-06 |
| Sspon.02G0025360 | 2,714 | 3,82E-06 |
| Sspon.02G0025380 | 2,439 | 3,82E-06 |
| Sspon.02G0025400 | -1,719 | 3,83E-06 |
| Sspon.02G0025540 | -2,277 | 3,89E-06 |
| Sspon.02G0025660 | 1,682 | 3,91E-06 |
| Sspon.02G0025730 | -1,048 | 3,91E-06 |
| Sspon.02G0025760 | -1,478 | 3,92E-06 |
| Sspon.02G0025790 | -1,588 | 3,93E-06 |
| Sspon.02G0025800 | 11,122 | 3,93E-06 |
| Sspon.02G0025840 | 1,291 | 3,93E-06 |
| Sspon.02G0025910 | -1,509 | 3,93E-06 |
| Sspon.02G0026010 | -1,479 | 3,93E-06 |
| Sspon.02G0026020 | -1,132 | 3,96E-06 |

| | | |
|---|---|---|
| Sspon.02G0026080 | -1,815 | 3,97E-06 |
| Sspon.02G0026140 | 1,324 | 3,99E-06 |
| Sspon.02G0026180 | -1,270 | 4,00E-06 |
| Sspon.02G0026380 | 4,845 | 4,01E-06 |
| Sspon.02G0026440 | 3,182 | 4,07E-06 |
| Sspon.02G0026480 | -2,557 | 4,09E-06 |
| Sspon.02G0026530 | -1,325 | 4,12E-06 |
| Sspon.02G0026640 | 10,567 | 4,12E-06 |
| Sspon.02G0026650 | -2,367 | 4,13E-06 |
| Sspon.02G0026790 | 1,735 | 4,14E-06 |
| Sspon.02G0026970 | 1,309 | 4,14E-06 |
| Sspon.02G0026980 | 1,563 | 4,15E-06 |
| Sspon.02G0027030 | 18,397 | 4,17E-06 |
| Sspon.02G0027040 | 3,867 | 4,18E-06 |
| Sspon.02G0027090 | 2,403 | 4,20E-06 |
| Sspon.02G0027100 | -1,487 | 4,21E-06 |
| Sspon.02G0027150 | 17,478 | 4,23E-06 |
| Sspon.02G0027280 | -2,782 | 4,23E-06 |
| Sspon.02G0027290 | -1,355 | 4,27E-06 |
| Sspon.02G0027330 | -1,053 | 4,28E-06 |
| Sspon.02G0027400 | 1,581 | 4,29E-06 |
| Sspon.02G0027480 | 4,861 | 4,31E-06 |
| Sspon.02G0027560 | 1,668 | 4,34E-06 |
| Sspon.02G0027620 | 17,025 | 4,35E-06 |
| Sspon.02G0027690 | 2,228 | 4,35E-06 |
| Sspon.02G0027790 | 1,513 | 4,35E-06 |
| Sspon.02G0027810 | -1,537 | 4,37E-06 |
| Sspon.02G0027820 | 5,609 | 4,37E-06 |
| Sspon.02G0027840 | 2,443 | 4,42E-06 |
| Sspon.02G0027860 | -1,642 | 4,45E-06 |
| Sspon.02G0027900 | 2,459 | 4,45E-06 |
| Sspon.02G0027920 | 6,442 | 4,48E-06 |
| Sspon.02G0028520 | 1,992 | 4,51E-06 |
| Sspon.02G0028570 | 1,593 | 4,52E-06 |
| Sspon.02G0028620 | 5,836 | 4,54E-06 |
| Sspon.02G0028650 | 1,636 | 4,55E-06 |
| Sspon.02G0028660 | -1,609 | 4,55E-06 |
| Sspon.02G0028740 | -17,147 | 4,57E-06 |
| Sspon.02G0028800 | -1,102 | 4,57E-06 |
| Sspon.02G0028900 | 16,406 | 4,64E-06 |
| Sspon.02G0028990 | 5,130 | 4,67E-06 |
| Sspon.02G0029060 | 1,710 | 4,67E-06 |
| Sspon.02G0029130 | 1,154 | 4,68E-06 |

| | | |
|---|---|---|
| Sspon.02G0029180 | 15,598 | 4,70E-06 |
| Sspon.02G0029240 | -1,044 | 4,72E-06 |
| Sspon.02G0029330 | 1,135 | 4,72E-06 |
| Sspon.02G0029340 | -1,607 | 4,72E-06 |
| Sspon.02G0029400 | 11,425 | 4,72E-06 |
| Sspon.02G0029460 | 3,294 | 4,74E-06 |
| Sspon.02G0029480 | -1,080 | 4,76E-06 |
| Sspon.02G0029490 | 1,547 | 4,78E-06 |
| Sspon.02G0029510 | 16,179 | 4,78E-06 |
| Sspon.02G0029580 | 4,096 | 4,83E-06 |
| Sspon.02G0029670 | 1,279 | 4,83E-06 |
| Sspon.02G0030030 | 16,043 | 4,87E-06 |
| Sspon.02G0030060 | -1,577 | 4,92E-06 |
| Sspon.02G0030160 | -1,428 | 4,93E-06 |
| Sspon.02G0030170 | 1,664 | 4,93E-06 |
| Sspon.02G0030180 | 4,792 | 4,98E-06 |
| Sspon.02G0030190 | -1,786 | 4,99E-06 |
| Sspon.02G0030200 | 1,469 | 5,00E-06 |
| Sspon.02G0030210 | 3,232 | 5,01E-06 |
| Sspon.02G0030240 | 2,985 | 5,01E-06 |
| Sspon.02G0030250 | 16,084 | 5,05E-06 |
| Sspon.02G0030330 | 2,026 | 5,07E-06 |
| Sspon.02G0030350 | 2,507 | 5,11E-06 |
| Sspon.02G0030360 | 16,262 | 5,11E-06 |
| Sspon.02G0030420 | 6,952 | 5,15E-06 |
| Sspon.02G0030490 | 9,026 | 5,15E-06 |
| Sspon.02G0030550 | -1,323 | 5,16E-06 |
| Sspon.02G0030600 | -3,784 | 5,16E-06 |
| Sspon.02G0030640 | -1,222 | 5,20E-06 |
| Sspon.02G0030670 | 1,453 | 5,24E-06 |
| Sspon.02G0030740 | 1,036 | 5,25E-06 |
| Sspon.02G0030760 | -2,038 | 5,27E-06 |
| Sspon.02G0030830 | -17,678 | 5,29E-06 |
| Sspon.02G0030910 | 15,861 | 5,29E-06 |
| Sspon.02G0030920 | 1,891 | 5,30E-06 |
| Sspon.02G0030950 | -2,176 | 5,31E-06 |
| Sspon.02G0030960 | -1,109 | 5,31E-06 |
| Sspon.02G0030990 | 1,719 | 5,33E-06 |
| Sspon.02G0031090 | 1,429 | 5,33E-06 |
| Sspon.02G0031110 | 1,363 | 5,33E-06 |
| Sspon.02G0031230 | 2,756 | 5,35E-06 |
| Sspon.02G0031270 | -1,607 | 5,35E-06 |
| Sspon.02G0031360 | 3,415 | 5,37E-06 |

| | | |
|---|---|---|
| Sspon.02G0031370 | 1,501 | 5,40E-06 |
| Sspon.02G0031400 | -3,513 | 5,42E-06 |
| Sspon.02G0031500 | -6,095 | 5,42E-06 |
| Sspon.02G0031510 | 1,757 | 5,45E-06 |
| Sspon.02G0031520 | 2,586 | 5,48E-06 |
| Sspon.02G0031540 | -1,741 | 5,49E-06 |
| Sspon.02G0031780 | -1,694 | 5,54E-06 |
| Sspon.02G0031860 | 1,305 | 5,56E-06 |
| Sspon.02G0031870 | 2,331 | 5,60E-06 |
| Sspon.02G0031880 | -1,573 | 5,65E-06 |
| Sspon.02G0031910 | -17,078 | 5,69E-06 |
| Sspon.02G0031950 | 1,519 | 5,77E-06 |
| Sspon.02G0031960 | -17,561 | 5,83E-06 |
| Sspon.02G0032100 | -3,525 | 5,87E-06 |
| Sspon.02G0032150 | 10,728 | 5,88E-06 |
| Sspon.02G0032280 | 2,538 | 5,92E-06 |
| Sspon.02G0032370 | 1,200 | 5,96E-06 |
| Sspon.02G0032380 | 1,581 | 5,96E-06 |
| Sspon.02G0032430 | 1,560 | 5,97E-06 |
| Sspon.02G0032450 | 1,259 | 5,97E-06 |
| Sspon.02G0032490 | 1,704 | 5,97E-06 |
| Sspon.02G0032550 | -1,103 | 5,97E-06 |
| Sspon.02G0032570 | -1,667 | 5,97E-06 |
| Sspon.02G0032600 | 1,609 | 6,00E-06 |
| Sspon.02G0032620 | 1,762 | 6,02E-06 |
| Sspon.02G0032660 | 16,090 | 6,02E-06 |
| Sspon.02G0032780 | 1,363 | 6,03E-06 |
| Sspon.02G0032930 | 2,161 | 6,04E-06 |
| Sspon.02G0032950 | 1,230 | 6,06E-06 |
| Sspon.02G0032990 | 4,778 | 6,06E-06 |
| Sspon.02G0033000 | -1,191 | 6,08E-06 |
| Sspon.02G0033010 | -3,467 | 6,09E-06 |
| Sspon.02G0033020 | -1,449 | 6,11E-06 |
| Sspon.02G0033070 | 11,178 | 6,15E-06 |
| Sspon.02G0033110 | -3,920 | 6,17E-06 |
| Sspon.02G0033140 | -1,210 | 6,17E-06 |
| Sspon.02G0033190 | -17,046 | 6,23E-06 |
| Sspon.02G0033320 | 3,175 | 6,25E-06 |
| Sspon.02G0033390 | 3,036 | 6,28E-06 |
| Sspon.02G0033640 | 1,933 | 6,28E-06 |
| Sspon.02G0033980 | 1,577 | 6,28E-06 |
| Sspon.02G0033990 | 2,197 | 6,29E-06 |
| Sspon.02G0034010 | -3,480 | 6,31E-06 |

| | | |
|---|---|---|
| Sspon.02G0034050 | 2,055 | 6,31E-06 |
| Sspon.02G0034160 | 2,903 | 6,40E-06 |
| Sspon.02G0034170 | -1,915 | 6,42E-06 |
| Sspon.02G0034190 | -1,965 | 6,44E-06 |
| Sspon.02G0034510 | 16,151 | 6,48E-06 |
| Sspon.02G0034630 | -1,666 | 6,51E-06 |
| Sspon.02G0034640 | -1,970 | 6,52E-06 |
| Sspon.02G0034710 | -1,158 | 6,52E-06 |
| Sspon.02G0034760 | 18,038 | 6,54E-06 |
| Sspon.02G0034810 | 2,994 | 6,56E-06 |
| Sspon.02G0034860 | 5,482 | 6,56E-06 |
| Sspon.02G0034880 | 1,848 | 6,60E-06 |
| Sspon.02G0034900 | 7,209 | 6,61E-06 |
| Sspon.02G0034950 | 1,717 | 6,62E-06 |
| Sspon.02G0035040 | 1,853 | 6,69E-06 |
| Sspon.02G0035200 | 18,237 | 6,69E-06 |
| Sspon.02G0035230 | -2,911 | 6,69E-06 |
| Sspon.02G0035240 | 2,161 | 6,69E-06 |
| Sspon.02G0035280 | -2,495 | 6,75E-06 |
| Sspon.02G0035350 | 5,631 | 6,78E-06 |
| Sspon.02G0035390 | -1,643 | 6,88E-06 |
| Sspon.02G0035500 | 10,435 | 6,92E-06 |
| Sspon.02G0035520 | 5,205 | 7,08E-06 |
| Sspon.02G0035620 | 5,484 | 7,08E-06 |
| Sspon.02G0035640 | 5,536 | 7,08E-06 |
| Sspon.02G0035740 | -3,300 | 7,11E-06 |
| Sspon.02G0035800 | -1,305 | 7,11E-06 |
| Sspon.02G0035840 | 15,645 | 7,15E-06 |
| Sspon.02G0035910 | -2,703 | 7,17E-06 |
| Sspon.02G0035920 | 1,358 | 7,18E-06 |
| Sspon.02G0035930 | 1,120 | 7,18E-06 |
| Sspon.02G0036030 | 2,547 | 7,23E-06 |
| Sspon.02G0036190 | 1,362 | 7,23E-06 |
| Sspon.02G0036250 | 11,591 | 7,24E-06 |
| Sspon.02G0036340 | -1,137 | 7,26E-06 |
| Sspon.02G0036490 | 2,518 | 7,28E-06 |
| Sspon.02G0036570 | 3,006 | 7,31E-06 |
| Sspon.02G0036610 | -1,648 | 7,31E-06 |
| Sspon.02G0036640 | 2,822 | 7,31E-06 |
| Sspon.02G0036710 | 1,250 | 7,35E-06 |
| Sspon.02G0036770 | 2,444 | 7,36E-06 |
| Sspon.02G0036830 | 15,819 | 7,42E-06 |
| Sspon.02G0036840 | -1,880 | 7,46E-06 |

| | | |
|---|---|---|
| Sspon.02G0036910 | 2,295 | 7,48E-06 |
| Sspon.02G0036970 | 2,960 | 7,51E-06 |
| Sspon.02G0036980 | 2,169 | 7,53E-06 |
| Sspon.02G0037020 | 16,507 | 7,53E-06 |
| Sspon.02G0037060 | 20,959 | 7,54E-06 |
| Sspon.02G0037070 | -1,816 | 7,55E-06 |
| Sspon.02G0037080 | 16,508 | 7,55E-06 |
| Sspon.02G0037110 | 10,306 | 7,60E-06 |
| Sspon.02G0037250 | 2,169 | 7,60E-06 |
| Sspon.02G0037270 | -1,719 | 7,63E-06 |
| Sspon.02G0037290 | 1,324 | 7,68E-06 |
| Sspon.02G0037370 | 2,576 | 7,73E-06 |
| Sspon.02G0037520 | -1,309 | 7,73E-06 |
| Sspon.02G0037540 | -1,492 | 7,75E-06 |
| Sspon.02G0037580 | 4,265 | 7,78E-06 |
| Sspon.02G0037610 | -1,628 | 7,81E-06 |
| Sspon.02G0037890 | 1,816 | 7,81E-06 |
| Sspon.02G0037910 | 1,388 | 7,90E-06 |
| Sspon.02G0037960 | 2,996 | 7,90E-06 |
| Sspon.02G0038070 | 2,618 | 7,93E-06 |
| Sspon.02G0038080 | 2,184 | 7,97E-06 |
| Sspon.02G0038290 | -1,154 | 7,97E-06 |
| Sspon.02G0038300 | -2,034 | 7,97E-06 |
| Sspon.02G0038390 | -1,497 | 7,98E-06 |
| Sspon.02G0038410 | -2,084 | 8,01E-06 |
| Sspon.02G0038420 | -2,083 | 8,01E-06 |
| Sspon.02G0038450 | -17,232 | 8,01E-06 |
| Sspon.02G0038470 | 7,733 | 8,09E-06 |
| Sspon.02G0038500 | 2,751 | 8,16E-06 |
| Sspon.02G0038610 | -1,277 | 8,16E-06 |
| Sspon.02G0038650 | -2,780 | 8,19E-06 |
| Sspon.02G0038700 | -1,153 | 8,19E-06 |
| Sspon.02G0038730 | -16,978 | 8,21E-06 |
| Sspon.02G0038770 | 1,424 | 8,21E-06 |
| Sspon.02G0038800 | 9,252 | 8,24E-06 |
| Sspon.02G0038810 | 15,883 | 8,32E-06 |
| Sspon.02G0038820 | 2,005 | 8,32E-06 |
| Sspon.02G0038830 | 2,143 | 8,34E-06 |
| Sspon.02G0038880 | -1,938 | 8,37E-06 |
| Sspon.02G0038890 | -1,586 | 8,37E-06 |
| Sspon.02G0038980 | 1,447 | 8,44E-06 |
| Sspon.02G0039040 | -1,379 | 8,45E-06 |
| Sspon.02G0039060 | -1,773 | 8,47E-06 |

| | | |
|---|---|---|
| Sspon.02G0039280 | -1,285 | 8,48E-06 |
| Sspon.02G0039370 | -3,390 | 8,48E-06 |
| Sspon.02G0039400 | -1,387 | 8,58E-06 |
| Sspon.02G0039490 | 4,893 | 8,67E-06 |
| Sspon.02G0039640 | -3,618 | 8,70E-06 |
| Sspon.02G0039670 | 15,820 | 8,71E-06 |
| Sspon.02G0039730 | 2,041 | 8,74E-06 |
| Sspon.02G0039800 | 3,651 | 8,76E-06 |
| Sspon.02G0039820 | 1,689 | 8,76E-06 |
| Sspon.02G0039960 | -2,053 | 8,79E-06 |
| Sspon.02G0040090 | -1,536 | 8,80E-06 |
| Sspon.02G0040200 | -1,437 | 8,80E-06 |
| Sspon.02G0040240 | 3,555 | 8,80E-06 |
| Sspon.02G0040310 | 3,777 | 8,84E-06 |
| Sspon.02G0040370 | -1,209 | 8,87E-06 |
| Sspon.02G0040410 | 15,472 | 8,91E-06 |
| Sspon.02G0040480 | -2,728 | 8,95E-06 |
| Sspon.02G0040540 | 1,528 | 8,96E-06 |
| Sspon.02G0040550 | -1,688 | 8,96E-06 |
| Sspon.02G0040650 | -1,061 | 8,99E-06 |
| Sspon.02G0040820 | 1,019 | 9,14E-06 |
| Sspon.02G0040830 | -1,255 | 9,17E-06 |
| Sspon.02G0040860 | -16,908 | 9,17E-06 |
| Sspon.02G0040910 | -17,748 | 9,18E-06 |
| Sspon.02G0041020 | -1,613 | 9,18E-06 |
| Sspon.02G0041050 | 1,067 | 9,20E-06 |
| Sspon.02G0041070 | 1,206 | 9,21E-06 |
| Sspon.02G0041160 | -1,488 | 9,38E-06 |
| Sspon.02G0041200 | 1,367 | 9,56E-06 |
| Sspon.02G0041210 | -1,030 | 9,57E-06 |
| Sspon.02G0041240 | 1,177 | 9,58E-06 |
| Sspon.02G0041260 | -1,211 | 9,58E-06 |
| Sspon.02G0041280 | 1,708 | 9,63E-06 |
| Sspon.02G0041420 | 10,395 | 9,69E-06 |
| Sspon.02G0041580 | 15,688 | 9,71E-06 |
| Sspon.02G0041590 | 1,528 | 9,77E-06 |
| Sspon.02G0041620 | -1,791 | 9,77E-06 |
| Sspon.02G0041800 | -1,947 | 9,78E-06 |
| Sspon.02G0041810 | 1,753 | 9,85E-06 |
| Sspon.02G0041830 | -2,685 | 9,87E-06 |
| Sspon.02G0041920 | -1,074 | 9,94E-06 |
| Sspon.02G0042010 | 1,584 | 1,00E-05 |
| Sspon.02G0042100 | 1,561 | 1,00E-05 |

| | | |
|---|---|---|
| Sspon.02G0042190 | 15,536 | 1,00E-05 |
| Sspon.02G0042310 | 3,264 | 1,00E-05 |
| Sspon.02G0042330 | -1,487 | 1,01E-05 |
| Sspon.02G0042380 | -1,216 | 1,01E-05 |
| Sspon.02G0042460 | 1,134 | 1,01E-05 |
| Sspon.02G0042600 | 10,953 | 1,01E-05 |
| Sspon.02G0042650 | 15,827 | 1,01E-05 |
| Sspon.02G0042670 | 1,805 | 1,03E-05 |
| Sspon.02G0042730 | 2,784 | 1,03E-05 |
| Sspon.02G0042810 | 1,011 | 1,04E-05 |
| Sspon.02G0042850 | -1,550 | 1,05E-05 |
| Sspon.02G0042920 | 9,923 | 1,05E-05 |
| Sspon.02G0042970 | 1,691 | 1,05E-05 |
| Sspon.02G0042990 | 7,246 | 1,05E-05 |
| Sspon.02G0043020 | -3,681 | 1,05E-05 |
| Sspon.02G0043030 | 4,152 | 1,05E-05 |
| Sspon.02G0043040 | 8,682 | 1,06E-05 |
| Sspon.02G0043210 | 1,326 | 1,06E-05 |
| Sspon.02G0043230 | 1,986 | 1,07E-05 |
| Sspon.02G0043300 | 2,518 | 1,07E-05 |
| Sspon.02G0043650 | 2,472 | 1,07E-05 |
| Sspon.02G0043680 | -1,071 | 1,08E-05 |
| Sspon.02G0043710 | 1,242 | 1,09E-05 |
| Sspon.02G0043810 | -1,612 | 1,09E-05 |
| Sspon.02G0043840 | 15,948 | 1,09E-05 |
| Sspon.02G0043860 | 1,829 | 1,10E-05 |
| Sspon.02G0043900 | 1,429 | 1,11E-05 |
| Sspon.02G0044210 | -1,316 | 1,12E-05 |
| Sspon.02G0044250 | 2,058 | 1,13E-05 |
| Sspon.02G0044310 | 2,042 | 1,15E-05 |
| Sspon.02G0044350 | 4,133 | 1,16E-05 |
| Sspon.02G0044470 | -1,424 | 1,16E-05 |
| Sspon.02G0044510 | 3,864 | 1,16E-05 |
| Sspon.02G0044550 | -1,748 | 1,17E-05 |
| Sspon.02G0044580 | 8,798 | 1,17E-05 |
| Sspon.02G0044660 | -16,806 | 1,17E-05 |
| Sspon.02G0044750 | -1,082 | 1,18E-05 |
| Sspon.02G0044760 | 1,823 | 1,18E-05 |
| Sspon.02G0044810 | -1,476 | 1,19E-05 |
| Sspon.02G0044820 | 1,525 | 1,19E-05 |
| Sspon.02G0044960 | 1,210 | 1,20E-05 |
| Sspon.02G0045010 | 1,734 | 1,22E-05 |
| Sspon.02G0045020 | 1,625 | 1,22E-05 |

| | | |
|---|---|---|
| Sspon.02G0045240 | -1,333 | 1,23E-05 |
| Sspon.02G0045340 | 2,090 | 1,24E-05 |
| Sspon.02G0045400 | -2,544 | 1,24E-05 |
| Sspon.02G0045410 | 1,583 | 1,24E-05 |
| Sspon.02G0045420 | -1,637 | 1,24E-05 |
| Sspon.02G0045430 | 1,754 | 1,24E-05 |
| Sspon.02G0045500 | -2,110 | 1,25E-05 |
| Sspon.02G0045530 | 2,023 | 1,25E-05 |
| Sspon.02G0045540 | 1,867 | 1,25E-05 |
| Sspon.02G0045580 | 1,187 | 1,26E-05 |
| Sspon.02G0045610 | 2,526 | 1,26E-05 |
| Sspon.02G0045810 | -2,452 | 1,27E-05 |
| Sspon.02G0045870 | 1,457 | 1,27E-05 |
| Sspon.02G0045880 | 15,768 | 1,27E-05 |
| Sspon.02G0045920 | -1,725 | 1,28E-05 |
| Sspon.02G0045930 | 15,429 | 1,28E-05 |
| Sspon.02G0045940 | -3,518 | 1,30E-05 |
| Sspon.02G0046060 | 10,921 | 1,30E-05 |
| Sspon.02G0046160 | 1,272 | 1,31E-05 |
| Sspon.02G0046170 | -5,440 | 1,31E-05 |
| Sspon.02G0046180 | 1,716 | 1,31E-05 |
| Sspon.02G0046250 | 9,997 | 1,32E-05 |
| Sspon.02G0046310 | -1,410 | 1,32E-05 |
| Sspon.02G0046420 | -1,222 | 1,32E-05 |
| Sspon.02G0046450 | -1,100 | 1,32E-05 |
| Sspon.02G0046490 | 1,121 | 1,32E-05 |
| Sspon.02G0046540 | -1,877 | 1,33E-05 |
| Sspon.02G0046580 | 4,879 | 1,34E-05 |
| Sspon.02G0046640 | 1,820 | 1,34E-05 |
| Sspon.02G0046720 | 1,476 | 1,35E-05 |
| Sspon.02G0046780 | 1,196 | 1,36E-05 |
| Sspon.02G0046970 | 1,072 | 1,37E-05 |
| Sspon.02G0047000 | 1,301 | 1,37E-05 |
| Sspon.02G0047040 | -3,734 | 1,37E-05 |
| Sspon.02G0047120 | 3,224 | 1,38E-05 |
| Sspon.02G0047180 | -2,274 | 1,38E-05 |
| Sspon.02G0047200 | -1,236 | 1,38E-05 |
| Sspon.02G0047310 | -1,042 | 1,39E-05 |
| Sspon.02G0047370 | -1,231 | 1,39E-05 |
| Sspon.02G0047470 | 1,339 | 1,40E-05 |
| Sspon.02G0047560 | 1,693 | 1,40E-05 |
| Sspon.02G0047570 | -16,961 | 1,40E-05 |
| Sspon.02G0047610 | -2,415 | 1,40E-05 |

| | | |
|---|---|---|
| Sspon.02G0047680 | -2,521 | 1,40E-05 |
| Sspon.02G0047690 | 2,283 | 1,40E-05 |
| Sspon.02G0047820 | -1,321 | 1,40E-05 |
| Sspon.02G0048010 | -2,189 | 1,40E-05 |
| Sspon.02G0048040 | 1,016 | 1,41E-05 |
| Sspon.02G0048120 | -16,755 | 1,41E-05 |
| Sspon.02G0048130 | -1,213 | 1,41E-05 |
| Sspon.02G0048140 | -2,282 | 1,42E-05 |
| Sspon.02G0048300 | -3,217 | 1,42E-05 |
| Sspon.02G0048320 | 1,393 | 1,42E-05 |
| Sspon.02G0048370 | 5,850 | 1,43E-05 |
| Sspon.02G0048460 | -1,864 | 1,43E-05 |
| Sspon.02G0048500 | -3,651 | 1,44E-05 |
| Sspon.02G0048530 | 1,083 | 1,44E-05 |
| Sspon.02G0048560 | -4,765 | 1,44E-05 |
| Sspon.02G0048590 | -2,840 | 1,45E-05 |
| Sspon.02G0048700 | 1,454 | 1,45E-05 |
| Sspon.02G0048930 | -1,481 | 1,45E-05 |
| Sspon.02G0049020 | 1,399 | 1,45E-05 |
| Sspon.02G0049100 | -1,170 | 1,45E-05 |
| Sspon.02G0049170 | -1,127 | 1,45E-05 |
| Sspon.02G0049180 | 2,049 | 1,46E-05 |
| Sspon.02G0049210 | 3,385 | 1,46E-05 |
| Sspon.02G0049250 | 2,155 | 1,46E-05 |
| Sspon.02G0049270 | -1,390 | 1,46E-05 |
| Sspon.02G0049310 | -16,977 | 1,47E-05 |
| Sspon.02G0049510 | -16,755 | 1,48E-05 |
| Sspon.02G0049540 | -7,391 | 1,48E-05 |
| Sspon.02G0049760 | 4,999 | 1,48E-05 |
| Sspon.02G0049850 | -2,770 | 1,51E-05 |
| Sspon.02G0049870 | -1,664 | 1,52E-05 |
| Sspon.02G0049910 | 2,592 | 1,52E-05 |
| Sspon.02G0049930 | -1,963 | 1,53E-05 |
| Sspon.02G0050040 | 5,550 | 1,56E-05 |
| Sspon.02G0050050 | -1,329 | 1,56E-05 |
| Sspon.02G0050240 | 1,373 | 1,56E-05 |
| Sspon.02G0050260 | 1,234 | 1,56E-05 |
| Sspon.02G0050280 | -1,343 | 1,57E-05 |
| Sspon.02G0050420 | -1,853 | 1,57E-05 |
| Sspon.02G0050480 | -2,763 | 1,57E-05 |
| Sspon.02G0050570 | 2,539 | 1,57E-05 |
| Sspon.02G0050610 | -2,310 | 1,59E-05 |
| Sspon.02G0050760 | -1,002 | 1,59E-05 |

| | | |
|---|---|---|
| Sspon.02G0050800 | 1,084 | 1,59E-05 |
| Sspon.02G0050880 | -1,027 | 1,59E-05 |
| Sspon.02G0050970 | -1,630 | 1,59E-05 |
| Sspon.02G0051010 | -2,806 | 1,60E-05 |
| Sspon.02G0051040 | 4,954 | 1,61E-05 |
| Sspon.02G0051130 | 1,074 | 1,62E-05 |
| Sspon.02G0051180 | -2,517 | 1,63E-05 |
| Sspon.02G0051400 | 1,345 | 1,63E-05 |
| Sspon.02G0051460 | 4,399 | 1,64E-05 |
| Sspon.02G0051630 | 1,992 | 1,64E-05 |
| Sspon.02G0051640 | 2,232 | 1,64E-05 |
| Sspon.02G0051660 | 2,646 | 1,64E-05 |
| Sspon.02G0051740 | 2,840 | 1,64E-05 |
| Sspon.02G0051750 | 1,965 | 1,65E-05 |
| Sspon.02G0051780 | -2,936 | 1,65E-05 |
| Sspon.02G0051810 | 10,520 | 1,65E-05 |
| Sspon.02G0051930 | -1,129 | 1,66E-05 |
| Sspon.02G0051940 | 17,069 | 1,66E-05 |
| Sspon.02G0051980 | 2,520 | 1,66E-05 |
| Sspon.02G0052030 | 5,680 | 1,67E-05 |
| Sspon.02G0052060 | 1,220 | 1,67E-05 |
| Sspon.02G0052170 | 1,704 | 1,68E-05 |
| Sspon.02G0052210 | 1,929 | 1,68E-05 |
| Sspon.02G0052230 | 1,471 | 1,69E-05 |
| Sspon.02G0052260 | 9,731 | 1,69E-05 |
| Sspon.02G0052380 | 2,479 | 1,69E-05 |
| Sspon.02G0052410 | -1,113 | 1,72E-05 |
| Sspon.02G0052440 | -16,636 | 1,73E-05 |
| Sspon.02G0052470 | 3,555 | 1,73E-05 |
| Sspon.02G0052480 | 1,185 | 1,73E-05 |
| Sspon.02G0052520 | -16,700 | 1,75E-05 |
| Sspon.02G0052600 | 2,725 | 1,77E-05 |
| Sspon.02G0052620 | -7,813 | 1,79E-05 |
| Sspon.02G0052680 | 1,850 | 1,79E-05 |
| Sspon.02G0052730 | -2,945 | 1,79E-05 |
| Sspon.02G0052750 | 2,384 | 1,79E-05 |
| Sspon.02G0052770 | 1,172 | 1,80E-05 |
| Sspon.02G0052830 | 4,612 | 1,80E-05 |
| Sspon.02G0052870 | -1,069 | 1,80E-05 |
| Sspon.02G0052960 | 3,555 | 1,81E-05 |
| Sspon.02G0053080 | 2,450 | 1,81E-05 |
| Sspon.02G0053130 | 3,069 | 1,82E-05 |
| Sspon.02G0053300 | -16,634 | 1,82E-05 |

| | | |
|---|---|---|
| Sspon.02G0053310 | -1,387 | 1,84E-05 |
| Sspon.02G0053510 | 1,551 | 1,84E-05 |
| Sspon.02G0053550 | -1,039 | 1,85E-05 |
| Sspon.02G0053570 | 2,998 | 1,87E-05 |
| Sspon.02G0053630 | 1,834 | 1,87E-05 |
| Sspon.02G0053720 | 1,338 | 1,88E-05 |
| Sspon.02G0053730 | 2,309 | 1,89E-05 |
| Sspon.02G0053740 | -1,414 | 1,96E-05 |
| Sspon.02G0053820 | -1,789 | 1,98E-05 |
| Sspon.02G0054040 | 1,062 | 1,98E-05 |
| Sspon.02G0054120 | 1,455 | 1,99E-05 |
| Sspon.02G0054280 | -1,259 | 1,99E-05 |
| Sspon.02G0054320 | -1,412 | 2,00E-05 |
| Sspon.02G0054360 | -2,846 | 2,00E-05 |
| Sspon.02G0054370 | 1,219 | 2,02E-05 |
| Sspon.02G0054410 | 1,145 | 2,02E-05 |
| Sspon.02G0054460 | -1,222 | 2,02E-05 |
| Sspon.02G0054650 | -1,087 | 2,04E-05 |
| Sspon.02G0054660 | 2,140 | 2,05E-05 |
| Sspon.02G0054680 | 2,881 | 2,05E-05 |
| Sspon.02G0054730 | 1,214 | 2,06E-05 |
| Sspon.02G0054830 | 2,388 | 2,06E-05 |
| Sspon.02G0054860 | 1,324 | 2,07E-05 |
| Sspon.02G0054900 | 1,126 | 2,07E-05 |
| Sspon.02G0054950 | 1,257 | 2,07E-05 |
| Sspon.02G0055000 | -1,588 | 2,08E-05 |
| Sspon.02G0055050 | -1,373 | 2,08E-05 |
| Sspon.02G0055060 | -3,009 | 2,08E-05 |
| Sspon.02G0055160 | 4,234 | 2,09E-05 |
| Sspon.02G0055300 | 15,507 | 2,09E-05 |
| Sspon.02G0055330 | -1,389 | 2,09E-05 |
| Sspon.02G0055370 | -1,570 | 2,10E-05 |
| Sspon.02G0055460 | 16,404 | 2,11E-05 |
| Sspon.02G0055550 | 1,184 | 2,12E-05 |
| Sspon.02G0055620 | 1,061 | 2,13E-05 |
| Sspon.02G0055720 | 2,574 | 2,13E-05 |
| Sspon.02G0055780 | -1,638 | 2,14E-05 |
| Sspon.02G0055830 | -1,426 | 2,14E-05 |
| Sspon.02G0055890 | 1,054 | 2,14E-05 |
| Sspon.02G0056000 | 3,122 | 2,15E-05 |
| Sspon.02G0056050 | -1,381 | 2,15E-05 |
| Sspon.02G0056090 | -1,133 | 2,15E-05 |
| Sspon.02G0056240 | -1,668 | 2,16E-05 |

| | | |
|---|---|---|
| Sspon.02G0056330 | 1,271 | 2,16E-05 |
| Sspon.02G0056350 | 4,768 | 2,17E-05 |
| Sspon.02G0056380 | 4,712 | 2,18E-05 |
| Sspon.02G0056420 | -1,695 | 2,18E-05 |
| Sspon.02G0056440 | 3,101 | 2,18E-05 |
| Sspon.02G0056490 | 1,552 | 2,20E-05 |
| Sspon.02G0056660 | 15,177 | 2,20E-05 |
| Sspon.02G0056770 | 5,707 | 2,22E-05 |
| Sspon.02G0056830 | 1,874 | 2,25E-05 |
| Sspon.02G0056860 | -1,041 | 2,26E-05 |
| Sspon.02G0056890 | -1,047 | 2,26E-05 |
| Sspon.02G0056970 | 3,031 | 2,26E-05 |
| Sspon.02G0056980 | -1,670 | 2,27E-05 |
| Sspon.02G0057150 | 1,063 | 2,27E-05 |
| Sspon.02G0057200 | 1,388 | 2,27E-05 |
| Sspon.02G0057240 | -1,645 | 2,28E-05 |
| Sspon.02G0057410 | 1,470 | 2,29E-05 |
| Sspon.02G0057450 | -2,212 | 2,32E-05 |
| Sspon.02G0057520 | 2,736 | 2,33E-05 |
| Sspon.02G0057530 | 1,380 | 2,35E-05 |
| Sspon.02G0057570 | -1,010 | 2,38E-05 |
| Sspon.02G0057650 | 2,611 | 2,38E-05 |
| Sspon.02G0057750 | -1,128 | 2,39E-05 |
| Sspon.02G0057830 | -7,841 | 2,39E-05 |
| Sspon.02G0057900 | 1,949 | 2,41E-05 |
| Sspon.02G0057980 | -1,523 | 2,42E-05 |
| Sspon.02G0058010 | -1,164 | 2,42E-05 |
| Sspon.02G0058020 | -1,112 | 2,43E-05 |
| Sspon.02G0058030 | 1,352 | 2,43E-05 |
| Sspon.02G0058040 | -2,004 | 2,44E-05 |
| Sspon.02G0058070 | 1,687 | 2,44E-05 |
| Sspon.02G0058090 | 1,834 | 2,44E-05 |
| Sspon.02G0058220 | 1,653 | 2,45E-05 |
| Sspon.02G0058240 | 1,164 | 2,45E-05 |
| Sspon.02G0058350 | 1,625 | 2,46E-05 |
| Sspon.02G0058410 | 1,099 | 2,47E-05 |
| Sspon.02G0058460 | 1,957 | 2,48E-05 |
| Sspon.02G0058490 | 6,229 | 2,49E-05 |
| Sspon.02G0058500 | 1,467 | 2,49E-05 |
| Sspon.02G0058530 | 1,696 | 2,49E-05 |
| Sspon.02G0058540 | 2,354 | 2,49E-05 |
| Sspon.02G0058810 | 10,188 | 2,49E-05 |
| Sspon.02G0058860 | -2,363 | 2,49E-05 |

| | | |
|---|---|---|
| Sspon.02G0058930 | -1,830 | 2,54E-05 |
| Sspon.02G0058980 | 3,120 | 2,54E-05 |
| Sspon.02G0059070 | 2,170 | 2,54E-05 |
| Sspon.02G0059120 | -2,072 | 2,55E-05 |
| Sspon.02G0059150 | 2,082 | 2,55E-05 |
| Sspon.02G0059160 | 1,781 | 2,56E-05 |
| Sspon.02G0059250 | 4,046 | 2,56E-05 |
| Sspon.02G0059320 | -2,216 | 2,56E-05 |
| Sspon.02G0059330 | 1,195 | 2,56E-05 |
| Sspon.02G0059380 | 2,345 | 2,58E-05 |
| Sspon.02G0059550 | -1,002 | 2,58E-05 |
| Sspon.02G0059660 | 1,878 | 2,58E-05 |
| Sspon.02G0059690 | 2,499 | 2,60E-05 |
| Sspon.02G0059710 | 1,289 | 2,61E-05 |
| Sspon.02G0059720 | 21,200 | 2,63E-05 |
| Sspon.02G0059840 | 2,501 | 2,64E-05 |
| Sspon.02G0059870 | 7,348 | 2,64E-05 |
| Sspon.02G0059880 | -16,626 | 2,65E-05 |
| Sspon.02G0059910 | 10,240 | 2,65E-05 |
| Sspon.02G0059940 | -4,816 | 2,66E-05 |
| Sspon.02G0059960 | -3,567 | 2,66E-05 |
| Sspon.02G0059970 | -3,217 | 2,66E-05 |
| Sspon.02G0060010 | 1,150 | 2,67E-05 |
| Sspon.02G0060050 | 1,927 | 2,68E-05 |
| Sspon.02G0060060 | 5,828 | 2,68E-05 |
| Sspon.02G0060080 | -1,582 | 2,69E-05 |
| Sspon.02G0060110 | 1,111 | 2,70E-05 |
| Sspon.02G0060130 | 2,347 | 2,71E-05 |
| Sspon.03G0000010 | -1,261 | 2,71E-05 |
| Sspon.03G0000050 | 4,715 | 2,72E-05 |
| Sspon.03G0000060 | 2,001 | 2,73E-05 |
| Sspon.03G0000100 | -2,352 | 2,73E-05 |
| Sspon.03G0000140 | 2,356 | 2,74E-05 |
| Sspon.03G0000150 | 2,965 | 2,75E-05 |
| Sspon.03G0000370 | 5,428 | 2,75E-05 |
| Sspon.03G0000780 | -1,237 | 2,77E-05 |
| Sspon.03G0000810 | -2,183 | 2,77E-05 |
| Sspon.03G0000820 | 2,078 | 2,79E-05 |
| Sspon.03G0000860 | 3,909 | 2,80E-05 |
| Sspon.03G0000870 | 10,513 | 2,80E-05 |
| Sspon.03G0000880 | -1,336 | 2,82E-05 |
| Sspon.03G0000960 | 1,323 | 2,82E-05 |
| Sspon.03G0000970 | 1,907 | 2,82E-05 |

| | | |
|---|---|---|
| Sspon.03G0001050 | -4,933 | 2,83E-05 |
| Sspon.03G0001160 | 2,061 | 2,87E-05 |
| Sspon.03G0001210 | 1,282 | 2,88E-05 |
| Sspon.03G0001250 | 5,423 | 2,90E-05 |
| Sspon.03G0001280 | 2,177 | 2,90E-05 |
| Sspon.03G0001340 | 17,097 | 2,94E-05 |
| Sspon.03G0001350 | 1,851 | 2,95E-05 |
| Sspon.03G0001400 | -1,312 | 2,96E-05 |
| Sspon.03G0001440 | -1,011 | 2,97E-05 |
| Sspon.03G0001510 | 2,984 | 2,97E-05 |
| Sspon.03G0001670 | -2,501 | 2,98E-05 |
| Sspon.03G0001830 | 1,103 | 2,99E-05 |
| Sspon.03G0001860 | -1,154 | 2,99E-05 |
| Sspon.03G0001920 | -1,708 | 2,99E-05 |
| Sspon.03G0001970 | -2,569 | 2,99E-05 |
| Sspon.03G0002010 | -2,530 | 3,00E-05 |
| Sspon.03G0002040 | -16,658 | 3,02E-05 |
| Sspon.03G0002090 | 6,112 | 3,02E-05 |
| Sspon.03G0002160 | -8,135 | 3,04E-05 |
| Sspon.03G0002200 | 2,564 | 3,04E-05 |
| Sspon.03G0002220 | 1,992 | 3,05E-05 |
| Sspon.03G0002230 | -7,961 | 3,05E-05 |
| Sspon.03G0002250 | 1,729 | 3,05E-05 |
| Sspon.03G0002260 | 1,708 | 3,05E-05 |
| Sspon.03G0002310 | -1,578 | 3,05E-05 |
| Sspon.03G0002320 | 8,920 | 3,06E-05 |
| Sspon.03G0002470 | 5,500 | 3,07E-05 |
| Sspon.03G0002480 | 1,217 | 3,08E-05 |
| Sspon.03G0002530 | 1,928 | 3,08E-05 |
| Sspon.03G0002560 | 2,040 | 3,08E-05 |
| Sspon.03G0002600 | 1,395 | 3,08E-05 |
| Sspon.03G0002610 | 9,926 | 3,10E-05 |
| Sspon.03G0002630 | 10,093 | 3,10E-05 |
| Sspon.03G0002660 | 1,185 | 3,11E-05 |
| Sspon.03G0002680 | 1,015 | 3,11E-05 |
| Sspon.03G0002710 | 1,961 | 3,11E-05 |
| Sspon.03G0002730 | -1,371 | 3,12E-05 |
| Sspon.03G0002740 | -1,994 | 3,13E-05 |
| Sspon.03G0002750 | -1,245 | 3,15E-05 |
| Sspon.03G0002760 | 5,063 | 3,19E-05 |
| Sspon.03G0002930 | 1,073 | 3,21E-05 |
| Sspon.03G0002940 | 1,667 | 3,21E-05 |
| Sspon.03G0002980 | 1,864 | 3,23E-05 |

| | | |
|---|---|---|
| Sspon.03G0003050 | -1,326 | 3,24E-05 |
| Sspon.03G0003100 | 6,316 | 3,24E-05 |
| Sspon.03G0003170 | -1,058 | 3,25E-05 |
| Sspon.03G0003180 | 8,978 | 3,26E-05 |
| Sspon.03G0003230 | 1,409 | 3,26E-05 |
| Sspon.03G0003360 | 7,201 | 3,26E-05 |
| Sspon.03G0003390 | 1,013 | 3,29E-05 |
| Sspon.03G0003400 | 3,270 | 3,30E-05 |
| Sspon.03G0003440 | 1,165 | 3,31E-05 |
| Sspon.03G0003450 | 2,355 | 3,31E-05 |
| Sspon.03G0003490 | -16,466 | 3,36E-05 |
| Sspon.03G0003530 | 4,216 | 3,38E-05 |
| Sspon.03G0003550 | -1,395 | 3,39E-05 |
| Sspon.03G0003590 | -5,532 | 3,40E-05 |
| Sspon.03G0003600 | 1,417 | 3,44E-05 |
| Sspon.03G0003620 | -4,077 | 3,44E-05 |
| Sspon.03G0003740 | 2,514 | 3,44E-05 |
| Sspon.03G0003750 | 1,170 | 3,45E-05 |
| Sspon.03G0003820 | 1,684 | 3,47E-05 |
| Sspon.03G0003900 | 2,634 | 3,47E-05 |
| Sspon.03G0003980 | 2,615 | 3,48E-05 |
| Sspon.03G0004060 | -1,105 | 3,48E-05 |
| Sspon.03G0004090 | 2,241 | 3,48E-05 |
| Sspon.03G0004150 | 2,568 | 3,51E-05 |
| Sspon.03G0004190 | -1,782 | 3,53E-05 |
| Sspon.03G0004200 | -2,216 | 3,53E-05 |
| Sspon.03G0004210 | 6,463 | 3,55E-05 |
| Sspon.03G0004250 | -1,177 | 3,57E-05 |
| Sspon.03G0004310 | -1,390 | 3,59E-05 |
| Sspon.03G0004410 | -1,115 | 3,60E-05 |
| Sspon.03G0004450 | -2,473 | 3,60E-05 |
| Sspon.03G0004460 | 1,838 | 3,62E-05 |
| Sspon.03G0004480 | 1,697 | 3,65E-05 |
| Sspon.03G0004640 | 4,323 | 3,66E-05 |
| Sspon.03G0004710 | 1,389 | 3,66E-05 |
| Sspon.03G0004730 | 1,585 | 3,67E-05 |
| Sspon.03G0004740 | -1,666 | 3,68E-05 |
| Sspon.03G0004870 | 2,296 | 3,69E-05 |
| Sspon.03G0005080 | -1,606 | 3,69E-05 |
| Sspon.03G0005090 | 1,275 | 3,69E-05 |
| Sspon.03G0005140 | 1,832 | 3,70E-05 |
| Sspon.03G0005190 | 2,060 | 3,70E-05 |
| Sspon.03G0005270 | -4,243 | 3,71E-05 |

| | | |
|---|---|---|
| Sspon.03G0005300 | 4,872 | 3,72E-05 |
| Sspon.03G0005310 | 2,172 | 3,77E-05 |
| Sspon.03G0005350 | -1,972 | 3,77E-05 |
| Sspon.03G0005360 | -2,189 | 3,78E-05 |
| Sspon.03G0005390 | 2,797 | 3,78E-05 |
| Sspon.03G0005400 | 1,685 | 3,80E-05 |
| Sspon.03G0005430 | 2,287 | 3,86E-05 |
| Sspon.03G0005450 | 2,078 | 3,87E-05 |
| Sspon.03G0005460 | 1,057 | 3,89E-05 |
| Sspon.03G0005700 | 2,964 | 3,89E-05 |
| Sspon.03G0005720 | 3,105 | 3,90E-05 |
| Sspon.03G0005810 | -1,862 | 3,90E-05 |
| Sspon.03G0005840 | 16,271 | 3,91E-05 |
| Sspon.03G0005960 | 1,422 | 3,92E-05 |
| Sspon.03G0006150 | -2,585 | 3,92E-05 |
| Sspon.03G0006160 | 1,388 | 3,94E-05 |
| Sspon.03G0006190 | -1,155 | 3,95E-05 |
| Sspon.03G0006250 | 1,675 | 3,95E-05 |
| Sspon.03G0006260 | 2,519 | 3,95E-05 |
| Sspon.03G0006400 | 3,559 | 3,99E-05 |
| Sspon.03G0006440 | 1,483 | 3,99E-05 |
| Sspon.03G0006490 | 1,128 | 4,00E-05 |
| Sspon.03G0006530 | 1,977 | 4,01E-05 |
| Sspon.03G0006570 | 1,545 | 4,01E-05 |
| Sspon.03G0006650 | -1,066 | 4,01E-05 |
| Sspon.03G0006680 | -2,170 | 4,02E-05 |
| Sspon.03G0006700 | 1,913 | 4,04E-05 |
| Sspon.03G0006840 | 2,065 | 4,05E-05 |
| Sspon.03G0006850 | -1,197 | 4,07E-05 |
| Sspon.03G0006860 | 1,829 | 4,08E-05 |
| Sspon.03G0006900 | 10,279 | 4,09E-05 |
| Sspon.03G0006910 | 1,672 | 4,09E-05 |
| Sspon.03G0007050 | 1,100 | 4,09E-05 |
| Sspon.03G0007090 | 1,339 | 4,09E-05 |
| Sspon.03G0007100 | -2,053 | 4,10E-05 |
| Sspon.03G0007130 | -2,154 | 4,11E-05 |
| Sspon.03G0007310 | -1,380 | 4,13E-05 |
| Sspon.03G0007410 | 1,174 | 4,14E-05 |
| Sspon.03G0007620 | 3,862 | 4,14E-05 |
| Sspon.03G0007640 | 5,002 | 4,15E-05 |
| Sspon.03G0007670 | 3,279 | 4,20E-05 |
| Sspon.03G0007710 | -3,722 | 4,21E-05 |
| Sspon.03G0007730 | 3,520 | 4,21E-05 |

| | | |
|---|---|---|
| Sspon.03G0007820 | 10,411 | 4,21E-05 |
| Sspon.03G0008120 | -1,281 | 4,21E-05 |
| Sspon.03G0008210 | 2,490 | 4,23E-05 |
| Sspon.03G0008220 | -1,685 | 4,24E-05 |
| Sspon.03G0008340 | 9,035 | 4,24E-05 |
| Sspon.03G0008350 | 1,557 | 4,25E-05 |
| Sspon.03G0008360 | 4,194 | 4,28E-05 |
| Sspon.03G0008370 | 1,875 | 4,29E-05 |
| Sspon.03G0008390 | 2,420 | 4,29E-05 |
| Sspon.03G0008480 | 2,497 | 4,29E-05 |
| Sspon.03G0008490 | 1,617 | 4,30E-05 |
| Sspon.03G0008620 | -1,782 | 4,31E-05 |
| Sspon.03G0008630 | 3,103 | 4,36E-05 |
| Sspon.03G0008780 | -1,754 | 4,36E-05 |
| Sspon.03G0008840 | 2,148 | 4,37E-05 |
| Sspon.03G0008900 | 3,156 | 4,37E-05 |
| Sspon.03G0008970 | 5,888 | 4,37E-05 |
| Sspon.03G0009070 | 2,583 | 4,37E-05 |
| Sspon.03G0009120 | 2,807 | 4,39E-05 |
| Sspon.03G0009270 | 2,559 | 4,39E-05 |
| Sspon.03G0009400 | 1,420 | 4,39E-05 |
| Sspon.03G0009510 | 1,100 | 4,40E-05 |
| Sspon.03G0009530 | 2,077 | 4,43E-05 |
| Sspon.03G0009830 | -3,321 | 4,44E-05 |
| Sspon.03G0009840 | -1,950 | 4,44E-05 |
| Sspon.03G0009880 | 15,983 | 4,45E-05 |
| Sspon.03G0009900 | 1,164 | 4,45E-05 |
| Sspon.03G0009910 | 1,460 | 4,48E-05 |
| Sspon.03G0009920 | 2,573 | 4,48E-05 |
| Sspon.03G0009960 | 2,833 | 4,50E-05 |
| Sspon.03G0009970 | 15,920 | 4,51E-05 |
| Sspon.03G0009980 | -1,801 | 4,52E-05 |
| Sspon.03G0010040 | 2,763 | 4,53E-05 |
| Sspon.03G0010050 | 1,843 | 4,54E-05 |
| Sspon.03G0010180 | 3,074 | 4,55E-05 |
| Sspon.03G0010200 | 2,435 | 4,59E-05 |
| Sspon.03G0010220 | -1,980 | 4,60E-05 |
| Sspon.03G0010250 | 1,908 | 4,62E-05 |
| Sspon.03G0010260 | 1,019 | 4,62E-05 |
| Sspon.03G0010420 | 2,126 | 4,63E-05 |
| Sspon.03G0010450 | 1,286 | 4,63E-05 |
| Sspon.03G0010480 | -1,128 | 4,64E-05 |
| Sspon.03G0010520 | 6,733 | 4,64E-05 |

| | | |
|---|---|---|
| Sspon.03G0010550 | -2,900 | 4,64E-05 |
| Sspon.03G0010600 | 9,995 | 4,66E-05 |
| Sspon.03G0010650 | -1,128 | 4,68E-05 |
| Sspon.03G0010660 | 1,798 | 4,73E-05 |
| Sspon.03G0011060 | 2,756 | 4,73E-05 |
| Sspon.03G0011120 | -1,443 | 4,74E-05 |
| Sspon.03G0011300 | 1,039 | 4,74E-05 |
| Sspon.03G0011310 | 2,028 | 4,75E-05 |
| Sspon.03G0011490 | 2,191 | 4,75E-05 |
| Sspon.03G0011500 | 1,515 | 4,76E-05 |
| Sspon.03G0011590 | 17,251 | 4,76E-05 |
| Sspon.03G0011620 | 1,426 | 4,76E-05 |
| Sspon.03G0011790 | 3,528 | 4,76E-05 |
| Sspon.03G0011810 | 1,836 | 4,77E-05 |
| Sspon.03G0011820 | 1,414 | 4,79E-05 |
| Sspon.03G0011930 | -1,745 | 4,79E-05 |
| Sspon.03G0012020 | -1,044 | 4,81E-05 |
| Sspon.03G0012080 | -1,524 | 4,83E-05 |
| Sspon.03G0012120 | -2,407 | 4,84E-05 |
| Sspon.03G0012140 | 15,865 | 4,85E-05 |
| Sspon.03G0012160 | -1,624 | 4,88E-05 |
| Sspon.03G0012190 | -1,006 | 4,90E-05 |
| Sspon.03G0012210 | 1,387 | 4,91E-05 |
| Sspon.03G0012440 | 9,944 | 4,91E-05 |
| Sspon.03G0012450 | -2,040 | 4,91E-05 |
| Sspon.03G0012470 | 1,050 | 4,92E-05 |
| Sspon.03G0012580 | 1,855 | 4,92E-05 |
| Sspon.03G0012590 | 8,883 | 4,93E-05 |
| Sspon.03G0012630 | 1,054 | 4,94E-05 |
| Sspon.03G0012640 | -1,314 | 4,94E-05 |
| Sspon.03G0012690 | -1,769 | 5,04E-05 |
| Sspon.03G0012820 | -1,379 | 5,05E-05 |
| Sspon.03G0012900 | 2,152 | 5,05E-05 |
| Sspon.03G0012970 | 1,622 | 5,06E-05 |
| Sspon.03G0012990 | 1,554 | 5,07E-05 |
| Sspon.03G0013060 | -3,049 | 5,08E-05 |
| Sspon.03G0013240 | 2,853 | 5,12E-05 |
| Sspon.03G0013260 | 3,024 | 5,12E-05 |
| Sspon.03G0013360 | 9,773 | 5,16E-05 |
| Sspon.03G0013450 | 1,101 | 5,17E-05 |
| Sspon.03G0013480 | -5,071 | 5,18E-05 |
| Sspon.03G0013500 | 1,004 | 5,18E-05 |
| Sspon.03G0013560 | 6,571 | 5,19E-05 |

| | | |
|---|---|---|
| Sspon.03G0013570 | -3,074 | 5,20E-05 |
| Sspon.03G0013600 | -1,288 | 5,22E-05 |
| Sspon.03G0013640 | -10,766 | 5,23E-05 |
| Sspon.03G0013730 | 1,943 | 5,23E-05 |
| Sspon.03G0013760 | -16,330 | 5,24E-05 |
| Sspon.03G0013770 | 2,367 | 5,25E-05 |
| Sspon.03G0013780 | 8,718 | 5,25E-05 |
| Sspon.03G0013820 | -1,059 | 5,26E-05 |
| Sspon.03G0013830 | -2,034 | 5,32E-05 |
| Sspon.03G0013850 | -1,182 | 5,32E-05 |
| Sspon.03G0013920 | 17,080 | 5,35E-05 |
| Sspon.03G0013950 | -3,185 | 5,35E-05 |
| Sspon.03G0013980 | 16,602 | 5,35E-05 |
| Sspon.03G0014050 | 1,885 | 5,35E-05 |
| Sspon.03G0014080 | 2,120 | 5,38E-05 |
| Sspon.03G0014100 | -1,573 | 5,40E-05 |
| Sspon.03G0014150 | 1,422 | 5,41E-05 |
| Sspon.03G0014230 | -1,690 | 5,42E-05 |
| Sspon.03G0014240 | 2,524 | 5,45E-05 |
| Sspon.03G0014250 | -2,127 | 5,46E-05 |
| Sspon.03G0014320 | 1,610 | 5,52E-05 |
| Sspon.03G0014340 | -1,710 | 5,54E-05 |
| Sspon.03G0014400 | -1,352 | 5,56E-05 |
| Sspon.03G0014470 | 3,890 | 5,56E-05 |
| Sspon.03G0014500 | 1,124 | 5,57E-05 |
| Sspon.03G0014520 | 2,725 | 5,57E-05 |
| Sspon.03G0014560 | -1,117 | 5,59E-05 |
| Sspon.03G0014590 | -3,651 | 5,59E-05 |
| Sspon.03G0014600 | 1,533 | 5,62E-05 |
| Sspon.03G0014610 | -2,105 | 5,63E-05 |
| Sspon.03G0014620 | 4,311 | 5,66E-05 |
| Sspon.03G0014630 | 1,227 | 5,66E-05 |
| Sspon.03G0014650 | -1,051 | 5,66E-05 |
| Sspon.03G0014660 | 2,112 | 5,66E-05 |
| Sspon.03G0014810 | 1,885 | 5,66E-05 |
| Sspon.03G0014820 | -1,440 | 5,67E-05 |
| Sspon.03G0014880 | 1,013 | 5,67E-05 |
| Sspon.03G0015040 | 3,933 | 5,67E-05 |
| Sspon.03G0015070 | -1,776 | 5,69E-05 |
| Sspon.03G0015110 | -1,033 | 5,71E-05 |
| Sspon.03G0015140 | 6,595 | 5,73E-05 |
| Sspon.03G0015200 | 1,154 | 5,73E-05 |
| Sspon.03G0015260 | -4,814 | 5,74E-05 |

| | | |
|---|---|---|
| Sspon.03G0015310 | 1,151 | 5,77E-05 |
| Sspon.03G0015380 | 1,765 | 5,77E-05 |
| Sspon.03G0015420 | -1,195 | 5,77E-05 |
| Sspon.03G0015510 | -1,220 | 5,77E-05 |
| Sspon.03G0015540 | -6,137 | 5,78E-05 |
| Sspon.03G0015550 | 1,532 | 5,82E-05 |
| Sspon.03G0015670 | 1,559 | 5,84E-05 |
| Sspon.03G0015710 | -2,516 | 5,85E-05 |
| Sspon.03G0015730 | 3,402 | 5,85E-05 |
| Sspon.03G0015810 | -3,128 | 5,85E-05 |
| Sspon.03G0015850 | 1,469 | 5,87E-05 |
| Sspon.03G0016000 | -1,400 | 5,87E-05 |
| Sspon.03G0016030 | 1,178 | 5,88E-05 |
| Sspon.03G0016060 | 1,091 | 5,89E-05 |
| Sspon.03G0016170 | -1,291 | 5,89E-05 |
| Sspon.03G0016200 | 1,017 | 5,91E-05 |
| Sspon.03G0016270 | 1,446 | 5,94E-05 |
| Sspon.03G0016310 | 2,858 | 5,95E-05 |
| Sspon.03G0016330 | 5,067 | 6,00E-05 |
| Sspon.03G0016340 | 15,479 | 6,03E-05 |
| Sspon.03G0016410 | 1,738 | 6,04E-05 |
| Sspon.03G0016420 | 1,937 | 6,07E-05 |
| Sspon.03G0016490 | 2,652 | 6,08E-05 |
| Sspon.03G0016500 | 4,172 | 6,09E-05 |
| Sspon.03G0016510 | 2,345 | 6,10E-05 |
| Sspon.03G0016520 | 1,738 | 6,11E-05 |
| Sspon.03G0016610 | 2,964 | 6,21E-05 |
| Sspon.03G0016710 | 9,047 | 6,23E-05 |
| Sspon.03G0016730 | 1,946 | 6,23E-05 |
| Sspon.03G0016780 | 1,754 | 6,27E-05 |
| Sspon.03G0016790 | -16,185 | 6,29E-05 |
| Sspon.03G0016880 | 6,012 | 6,30E-05 |
| Sspon.03G0016980 | 2,153 | 6,34E-05 |
| Sspon.03G0017070 | 3,039 | 6,35E-05 |
| Sspon.03G0017090 | -1,515 | 6,36E-05 |
| Sspon.03G0017140 | -3,774 | 6,37E-05 |
| Sspon.03G0017170 | 3,740 | 6,40E-05 |
| Sspon.03G0017380 | -2,426 | 6,42E-05 |
| Sspon.03G0017650 | 2,407 | 6,45E-05 |
| Sspon.03G0017730 | 2,050 | 6,46E-05 |
| Sspon.03G0017830 | 1,920 | 6,50E-05 |
| Sspon.03G0017840 | 2,523 | 6,53E-05 |
| Sspon.03G0017940 | 7,958 | 6,53E-05 |

| | | |
|---|---|---|
| Sspon.03G0017950 | 1,103 | 6,53E-05 |
| Sspon.03G0018030 | 2,109 | 6,58E-05 |
| Sspon.03G0018060 | 1,485 | 6,62E-05 |
| Sspon.03G0018080 | 1,142 | 6,62E-05 |
| Sspon.03G0018100 | 1,309 | 6,63E-05 |
| Sspon.03G0018230 | 9,551 | 6,70E-05 |
| Sspon.03G0018240 | 1,466 | 6,73E-05 |
| Sspon.03G0018290 | 2,654 | 6,73E-05 |
| Sspon.03G0018310 | 4,133 | 6,74E-05 |
| Sspon.03G0018320 | 9,786 | 6,77E-05 |
| Sspon.03G0018360 | 9,617 | 6,78E-05 |
| Sspon.03G0018410 | -1,521 | 6,85E-05 |
| Sspon.03G0018420 | -10,328 | 6,86E-05 |
| Sspon.03G0018430 | -3,722 | 6,87E-05 |
| Sspon.03G0018450 | -16,146 | 6,87E-05 |
| Sspon.03G0018530 | 6,961 | 6,89E-05 |
| Sspon.03G0018620 | 2,695 | 6,89E-05 |
| Sspon.03G0018640 | 2,322 | 6,90E-05 |
| Sspon.03G0018740 | 1,205 | 6,97E-05 |
| Sspon.03G0018780 | -1,207 | 6,99E-05 |
| Sspon.03G0018820 | -1,494 | 7,04E-05 |
| Sspon.03G0018870 | -2,163 | 7,07E-05 |
| Sspon.03G0018900 | 1,651 | 7,08E-05 |
| Sspon.03G0018940 | 2,387 | 7,09E-05 |
| Sspon.03G0018950 | 15,855 | 7,09E-05 |
| Sspon.03G0018990 | 2,100 | 7,12E-05 |
| Sspon.03G0019000 | -4,792 | 7,12E-05 |
| Sspon.03G0019060 | 2,955 | 7,13E-05 |
| Sspon.03G0019100 | -1,026 | 7,15E-05 |
| Sspon.03G0019210 | 3,590 | 7,18E-05 |
| Sspon.03G0019230 | -1,050 | 7,18E-05 |
| Sspon.03G0019340 | 9,542 | 7,20E-05 |
| Sspon.03G0019350 | 2,466 | 7,21E-05 |
| Sspon.03G0019430 | 9,625 | 7,21E-05 |
| Sspon.03G0019480 | -1,121 | 7,21E-05 |
| Sspon.03G0019490 | -1,499 | 7,29E-05 |
| Sspon.03G0019510 | 1,315 | 7,30E-05 |
| Sspon.03G0019520 | 4,329 | 7,31E-05 |
| Sspon.03G0019530 | -2,311 | 7,32E-05 |
| Sspon.03G0019550 | 1,592 | 7,34E-05 |
| Sspon.03G0019560 | 1,569 | 7,38E-05 |
| Sspon.03G0019570 | 4,586 | 7,38E-05 |
| Sspon.03G0019630 | 2,844 | 7,40E-05 |

| | | |
|---|---|---|
| Sspon.03G0019660 | 1,438 | 7,40E-05 |
| Sspon.03G0019710 | -1,606 | 7,42E-05 |
| Sspon.03G0019730 | -1,536 | 7,44E-05 |
| Sspon.03G0019740 | -1,024 | 7,45E-05 |
| Sspon.03G0019750 | 2,493 | 7,45E-05 |
| Sspon.03G0019770 | -2,560 | 7,48E-05 |
| Sspon.03G0020080 | 7,839 | 7,58E-05 |
| Sspon.03G0020090 | 15,372 | 7,61E-05 |
| Sspon.03G0020110 | 1,591 | 7,64E-05 |
| Sspon.03G0020210 | -1,310 | 7,64E-05 |
| Sspon.03G0020250 | 2,880 | 7,65E-05 |
| Sspon.03G0020290 | 1,700 | 7,72E-05 |
| Sspon.03G0020330 | -2,257 | 7,75E-05 |
| Sspon.03G0020460 | 2,737 | 7,75E-05 |
| Sspon.03G0020530 | -1,138 | 7,85E-05 |
| Sspon.03G0020600 | 2,199 | 7,85E-05 |
| Sspon.03G0020640 | 9,032 | 7,86E-05 |
| Sspon.03G0020700 | 1,871 | 7,86E-05 |
| Sspon.03G0020720 | -6,624 | 7,94E-05 |
| Sspon.03G0020780 | -5,596 | 7,96E-05 |
| Sspon.03G0020800 | 5,768 | 8,02E-05 |
| Sspon.03G0020820 | 1,122 | 8,03E-05 |
| Sspon.03G0020910 | 2,092 | 8,04E-05 |
| Sspon.03G0020980 | 1,151 | 8,04E-05 |
| Sspon.03G0021010 | -1,497 | 8,05E-05 |
| Sspon.03G0021020 | -1,194 | 8,08E-05 |
| Sspon.03G0021140 | 1,617 | 8,09E-05 |
| Sspon.03G0021190 | 2,149 | 8,14E-05 |
| Sspon.03G0021220 | -1,230 | 8,17E-05 |
| Sspon.03G0021280 | -1,632 | 8,17E-05 |
| Sspon.03G0021300 | 2,591 | 8,18E-05 |
| Sspon.03G0021320 | 9,431 | 8,22E-05 |
| Sspon.03G0021330 | 2,020 | 8,28E-05 |
| Sspon.03G0021420 | 1,111 | 8,28E-05 |
| Sspon.03G0021430 | 2,175 | 8,30E-05 |
| Sspon.03G0021490 | -4,588 | 8,33E-05 |
| Sspon.03G0021690 | -1,495 | 8,33E-05 |
| Sspon.03G0021780 | 1,564 | 8,38E-05 |
| Sspon.03G0021810 | -3,882 | 8,39E-05 |
| Sspon.03G0021820 | -1,389 | 8,42E-05 |
| Sspon.03G0021910 | -5,648 | 8,45E-05 |
| Sspon.03G0021980 | 2,574 | 8,48E-05 |
| Sspon.03G0021990 | 1,142 | 8,49E-05 |

| | | |
|---|---|---|
| Sspon.03G0022030 | 16,543 | 8,50E-05 |
| Sspon.03G0022090 | -3,016 | 8,50E-05 |
| Sspon.03G0022140 | 7,225 | 8,56E-05 |
| Sspon.03G0022230 | 1,851 | 8,59E-05 |
| Sspon.03G0022290 | 1,060 | 8,60E-05 |
| Sspon.03G0022300 | -1,861 | 8,60E-05 |
| Sspon.03G0022310 | 3,329 | 8,64E-05 |
| Sspon.03G0022340 | -1,355 | 8,72E-05 |
| Sspon.03G0022350 | 7,305 | 8,72E-05 |
| Sspon.03G0022430 | 3,852 | 8,72E-05 |
| Sspon.03G0022610 | -9,984 | 8,72E-05 |
| Sspon.03G0022660 | 3,761 | 8,72E-05 |
| Sspon.03G0022730 | 1,471 | 8,73E-05 |
| Sspon.03G0022750 | 1,237 | 8,78E-05 |
| Sspon.03G0022760 | -1,510 | 8,80E-05 |
| Sspon.03G0022770 | 1,331 | 8,82E-05 |
| Sspon.03G0022840 | 1,774 | 8,83E-05 |
| Sspon.03G0022860 | 2,025 | 8,86E-05 |
| Sspon.03G0022930 | 6,208 | 8,89E-05 |
| Sspon.03G0022950 | -3,570 | 8,97E-05 |
| Sspon.03G0022970 | 3,247 | 9,00E-05 |
| Sspon.03G0023020 | 1,843 | 9,01E-05 |
| Sspon.03G0023040 | -1,127 | 9,01E-05 |
| Sspon.03G0023110 | -1,430 | 9,03E-05 |
| Sspon.03G0023210 | 7,870 | 9,07E-05 |
| Sspon.03G0023230 | 9,321 | 9,08E-05 |
| Sspon.03G0023340 | -1,028 | 9,08E-05 |
| Sspon.03G0023630 | 1,782 | 9,09E-05 |
| Sspon.03G0023660 | 5,436 | 9,11E-05 |
| Sspon.03G0023670 | -1,306 | 9,11E-05 |
| Sspon.03G0023760 | 1,100 | 9,11E-05 |
| Sspon.03G0023860 | 1,429 | 9,13E-05 |
| Sspon.03G0023870 | -1,412 | 9,15E-05 |
| Sspon.03G0023890 | 4,657 | 9,25E-05 |
| Sspon.03G0023950 | 1,125 | 9,26E-05 |
| Sspon.03G0023970 | 1,221 | 9,28E-05 |
| Sspon.03G0023980 | 8,538 | 9,28E-05 |
| Sspon.03G0023990 | 17,009 | 9,29E-05 |
| Sspon.03G0024070 | 2,899 | 9,32E-05 |
| Sspon.03G0024130 | 1,632 | 9,33E-05 |
| Sspon.03G0024210 | -1,662 | 9,35E-05 |
| Sspon.03G0024250 | -1,122 | 9,35E-05 |
| Sspon.03G0024280 | -1,812 | 9,36E-05 |

| | | |
|---|---|---|
| Sspon.03G0024320 | 1,513 | 9,39E-05 |
| Sspon.03G0024330 | 7,435 | 9,39E-05 |
| Sspon.03G0024400 | -1,790 | 9,42E-05 |
| Sspon.03G0024550 | 1,085 | 9,42E-05 |
| Sspon.03G0024640 | -1,170 | 9,43E-05 |
| Sspon.03G0024680 | -1,500 | 9,46E-05 |
| Sspon.03G0024730 | 8,426 | 9,48E-05 |
| Sspon.03G0024800 | 3,641 | 9,49E-05 |
| Sspon.03G0024860 | -1,466 | 9,50E-05 |
| Sspon.03G0024950 | 1,302 | 9,50E-05 |
| Sspon.03G0024970 | -1,329 | 9,58E-05 |
| Sspon.03G0025110 | 1,267 | 9,58E-05 |
| Sspon.03G0025140 | -9,147 | 9,59E-05 |
| Sspon.03G0025210 | -2,018 | 9,65E-05 |
| Sspon.03G0025350 | 1,772 | 9,70E-05 |
| Sspon.03G0025380 | 1,619 | 9,70E-05 |
| Sspon.03G0025610 | -10,475 | 9,70E-05 |
| Sspon.03G0025630 | 3,496 | 9,71E-05 |
| Sspon.03G0025650 | 1,377 | 9,71E-05 |
| Sspon.03G0025660 | -1,442 | 9,71E-05 |
| Sspon.03G0025680 | 1,486 | 9,76E-05 |
| Sspon.03G0025690 | -1,004 | 9,82E-05 |
| Sspon.03G0025710 | -4,576 | 9,84E-05 |
| Sspon.03G0025750 | 1,912 | 9,84E-05 |
| Sspon.03G0025790 | -2,971 | 9,87E-05 |
| Sspon.03G0026000 | 1,473 | 9,88E-05 |
| Sspon.03G0026050 | 15,382 | 9,93E-05 |
| Sspon.03G0026080 | 8,373 | 9,93E-05 |
| Sspon.03G0026100 | 1,145 | 9,97E-05 |
| Sspon.03G0026170 | -8,632 | 0,0001 |
| Sspon.03G0026180 | -1,337 | 0,0001 |
| Sspon.03G0026200 | 2,629 | 0,0001 |
| Sspon.03G0026220 | 1,455 | 0,0001 |
| Sspon.03G0026290 | -10,376 | 0,0001 |
| Sspon.03G0026410 | -1,227 | 0,0001 |
| Sspon.03G0026440 | 1,168 | 0,0001 |
| Sspon.03G0026530 | 1,828 | 0,0001 |
| Sspon.03G0026650 | 5,820 | 0,0001 |
| Sspon.03G0026670 | 2,623 | 0,0001 |
| Sspon.03G0026680 | 3,146 | 0,0001 |
| Sspon.03G0026820 | 3,356 | 0,0001 |
| Sspon.03G0026830 | -1,167 | 0,0001 |
| Sspon.03G0026870 | 1,831 | 0,0001 |

| | | |
|---|---|---|
| Sspon.03G0026900 | -1,572 | 0,0001 |
| Sspon.03G0026960 | 3,047 | 0,0001 |
| Sspon.03G0026990 | 2,353 | 0,0001 |
| Sspon.03G0027030 | -1,989 | 0,0001 |
| Sspon.03G0027050 | -2,237 | 0,0001 |
| Sspon.03G0027060 | 8,330 | 0,0001 |
| Sspon.03G0027250 | -1,617 | 0,0001 |
| Sspon.03G0027300 | 4,028 | 0,0001 |
| Sspon.03G0027360 | 4,519 | 0,0001 |
| Sspon.03G0027380 | 3,858 | 0,0001 |
| Sspon.03G0027390 | -15,999 | 0,0001 |
| Sspon.03G0027430 | 3,445 | 0,0001 |
| Sspon.03G0027480 | 1,300 | 0,0001 |
| Sspon.03G0027500 | 2,633 | 0,0001 |
| Sspon.03G0027510 | -1,589 | 0,0001 |
| Sspon.03G0027620 | 1,867 | 0,0001 |
| Sspon.03G0027640 | 2,227 | 0,0001 |
| Sspon.03G0027710 | -1,659 | 0,0001 |
| Sspon.03G0027780 | -15,961 | 0,0001 |
| Sspon.03G0027790 | 3,198 | 0,0001 |
| Sspon.03G0027820 | 7,474 | 0,0001 |
| Sspon.03G0027830 | -1,542 | 0,0001 |
| Sspon.03G0027960 | 2,084 | 0,0001 |
| Sspon.03G0028000 | -1,052 | 0,0001 |
| Sspon.03G0028120 | 1,205 | 0,0001 |
| Sspon.03G0028130 | 5,947 | 0,0001 |
| Sspon.03G0028140 | 2,040 | 0,0001 |
| Sspon.03G0028210 | 1,265 | 0,0001 |
| Sspon.03G0028270 | -1,217 | 0,0001 |
| Sspon.03G0028310 | 15,809 | 0,0001 |
| Sspon.03G0028450 | 1,686 | 0,0001 |
| Sspon.03G0028710 | 1,970 | 0,0001 |
| Sspon.03G0028730 | 2,291 | 0,0001 |
| Sspon.03G0028770 | 1,127 | 0,0001 |
| Sspon.03G0028780 | 1,105 | 0,0001 |
| Sspon.03G0028790 | -5,110 | 0,0001 |
| Sspon.03G0028830 | -1,855 | 0,0001 |
| Sspon.03G0028860 | -15,953 | 0,0001 |
| Sspon.03G0028910 | 2,384 | 0,0001 |
| Sspon.03G0028970 | -2,616 | 0,0001 |
| Sspon.03G0029070 | 4,781 | 0,0001 |
| Sspon.03G0029090 | 6,266 | 0,0001 |
| Sspon.03G0029140 | 1,261 | 0,0001 |

| | | |
|---|---|---|
| Sspon.03G0029160 | -1,016 | 0,0001 |
| Sspon.03G0029240 | -1,133 | 0,0001 |
| Sspon.03G0029270 | -18,173 | 0,0001 |
| Sspon.03G0029330 | 1,654 | 0,0001 |
| Sspon.03G0029380 | 2,561 | 0,0001 |
| Sspon.03G0029430 | -1,797 | 0,0001 |
| Sspon.03G0029440 | 5,479 | 0,0001 |
| Sspon.03G0029510 | 3,686 | 0,0001 |
| Sspon.03G0029690 | 1,374 | 0,0001 |
| Sspon.03G0029750 | -1,757 | 0,0001 |
| Sspon.03G0029790 | 5,855 | 0,0001 |
| Sspon.03G0029860 | -2,120 | 0,0001 |
| Sspon.03G0029930 | 1,204 | 0,0001 |
| Sspon.03G0029990 | 4,564 | 0,0001 |
| Sspon.03G0030050 | 3,427 | 0,0001 |
| Sspon.03G0030130 | 1,683 | 0,0001 |
| Sspon.03G0030210 | 6,164 | 0,0001 |
| Sspon.03G0030260 | -1,430 | 0,0001 |
| Sspon.03G0030320 | 2,772 | 0,0001 |
| Sspon.03G0030370 | 1,187 | 0,0001 |
| Sspon.03G0030380 | -1,235 | 0,0001 |
| Sspon.03G0030470 | 1,762 | 0,0001 |
| Sspon.03G0030500 | 18,619 | 0,0001 |
| Sspon.03G0030510 | 1,188 | 0,0001 |
| Sspon.03G0030520 | 2,184 | 0,0001 |
| Sspon.03G0030530 | -1,054 | 0,0001 |
| Sspon.03G0030640 | 2,399 | 0,0001 |
| Sspon.03G0030760 | 2,489 | 0,0001 |
| Sspon.03G0030800 | 1,715 | 0,0001 |
| Sspon.03G0030880 | -1,575 | 0,0001 |
| Sspon.03G0030930 | 9,172 | 0,0001 |
| Sspon.03G0031040 | -1,207 | 0,0001 |
| Sspon.03G0031100 | 6,389 | 0,0001 |
| Sspon.03G0031160 | -2,560 | 0,0001 |
| Sspon.03G0031210 | 1,433 | 0,0001 |
| Sspon.03G0031230 | 1,277 | 0,0001 |
| Sspon.03G0031240 | -1,977 | 0,0001 |
| Sspon.03G0031340 | 2,872 | 0,0001 |
| Sspon.03G0031370 | 2,297 | 0,0001 |
| Sspon.03G0031390 | 1,502 | 0,0001 |
| Sspon.03G0031440 | 6,966 | 0,0001 |
| Sspon.03G0031450 | -9,061 | 0,0001 |
| Sspon.03G0031480 | 3,477 | 0,0001 |

| | | |
|---|---|---|
| Sspon.03G0031550 | 1,145 | 0,0001 |
| Sspon.03G0031710 | 2,355 | 0,0001 |
| Sspon.03G0031740 | -2,132 | 0,0001 |
| Sspon.03G0031790 | 1,378 | 0,0001 |
| Sspon.03G0031810 | -9,363 | 0,0001 |
| Sspon.03G0031870 | 3,955 | 0,0001 |
| Sspon.03G0031900 | 5,189 | 0,0001 |
| Sspon.03G0032020 | -1,045 | 0,0001 |
| Sspon.03G0032110 | 1,540 | 0,0001 |
| Sspon.03G0032180 | -1,916 | 0,0001 |
| Sspon.03G0032300 | 1,066 | 0,0001 |
| Sspon.03G0032380 | 2,445 | 0,0001 |
| Sspon.03G0032390 | 1,115 | 0,0001 |
| Sspon.03G0032550 | 3,714 | 0,0001 |
| Sspon.03G0032590 | -15,876 | 0,0001 |
| Sspon.03G0032650 | 16,315 | 0,0001 |
| Sspon.03G0032660 | 1,545 | 0,0001 |
| Sspon.03G0032860 | 1,029 | 0,0001 |
| Sspon.03G0033040 | 4,275 | 0,0001 |
| Sspon.03G0033090 | 2,059 | 0,0001 |
| Sspon.03G0033110 | 2,015 | 0,0001 |
| Sspon.03G0033120 | 1,205 | 0,0001 |
| Sspon.03G0033140 | 2,318 | 0,0001 |
| Sspon.03G0033150 | 8,186 | 0,0001 |
| Sspon.03G0033230 | 1,734 | 0,0001 |
| Sspon.03G0033330 | 1,318 | 0,0001 |
| Sspon.03G0033380 | -3,632 | 0,0001 |
| Sspon.03G0033450 | -1,366 | 0,0001 |
| Sspon.03G0033800 | -1,456 | 0,0001 |
| Sspon.03G0033860 | -2,170 | 0,0001 |
| Sspon.03G0033870 | 1,418 | 0,0001 |
| Sspon.03G0033910 | 1,872 | 0,0001 |
| Sspon.03G0033920 | 17,135 | 0,0001 |
| Sspon.03G0034010 | 1,207 | 0,0001 |
| Sspon.03G0034070 | 2,067 | 0,0001 |
| Sspon.03G0034130 | 3,921 | 0,0001 |
| Sspon.03G0034220 | 1,630 | 0,0001 |
| Sspon.03G0034250 | 1,053 | 0,0001 |
| Sspon.03G0034340 | -6,856 | 0,0001 |
| Sspon.03G0034430 | 2,344 | 0,0001 |
| Sspon.03G0034520 | 2,090 | 0,0001 |
| Sspon.03G0034530 | 18,382 | 0,0001 |
| Sspon.03G0034540 | 2,157 | 0,0001 |

| | | |
|---|---|---|
| Sspon.03G0034600 | 1,963 | 0,0001 |
| Sspon.03G0034620 | 2,450 | 0,0001 |
| Sspon.03G0034690 | 3,423 | 0,0001 |
| Sspon.03G0034760 | 1,688 | 0,0001 |
| Sspon.03G0034780 | 1,329 | 0,0001 |
| Sspon.03G0034920 | 4,405 | 0,0001 |
| Sspon.03G0035100 | -1,537 | 0,0001 |
| Sspon.03G0035130 | 4,030 | 0,0001 |
| Sspon.03G0035150 | -1,193 | 0,0001 |
| Sspon.03G0035240 | 3,404 | 0,0001 |
| Sspon.03G0035350 | -1,629 | 0,0001 |
| Sspon.03G0035380 | 1,142 | 0,0001 |
| Sspon.03G0035410 | 17,107 | 0,0001 |
| Sspon.03G0035460 | 2,372 | 0,0001 |
| Sspon.03G0035470 | 1,385 | 0,0001 |
| Sspon.03G0035540 | 1,381 | 0,0001 |
| Sspon.03G0035590 | -1,091 | 0,0001 |
| Sspon.03G0035600 | 1,868 | 0,0001 |
| Sspon.03G0035630 | -1,094 | 0,0001 |
| Sspon.03G0035680 | -2,206 | 0,0001 |
| Sspon.03G0035770 | 3,595 | 0,0001 |
| Sspon.03G0035850 | 2,576 | 0,0001 |
| Sspon.03G0035930 | 1,872 | 0,0001 |
| Sspon.03G0035990 | 3,065 | 0,0001 |
| Sspon.03G0036310 | 1,657 | 0,0001 |
| Sspon.03G0036440 | 1,592 | 0,0001 |
| Sspon.03G0036680 | 2,294 | 0,0001 |
| Sspon.03G0036700 | -1,061 | 0,0001 |
| Sspon.03G0036710 | 1,157 | 0,0001 |
| Sspon.03G0036850 | 5,938 | 0,0001 |
| Sspon.03G0036900 | 1,696 | 0,0001 |
| Sspon.03G0037060 | -1,628 | 0,0001 |
| Sspon.03G0037120 | 1,137 | 0,0001 |
| Sspon.03G0037170 | -3,715 | 0,0002 |
| Sspon.03G0037200 | 3,142 | 0,0002 |
| Sspon.03G0037210 | 1,376 | 0,0002 |
| Sspon.03G0037320 | -4,612 | 0,0002 |
| Sspon.03G0037340 | 1,807 | 0,0002 |
| Sspon.03G0037560 | 4,810 | 0,0002 |
| Sspon.03G0037640 | -1,185 | 0,0002 |
| Sspon.03G0037710 | 1,682 | 0,0002 |
| Sspon.03G0037760 | 1,278 | 0,0002 |
| Sspon.03G0037790 | -3,022 | 0,0002 |

| | | |
|---|---|---|
| Sspon.03G0037860 | -2,382 | 0,0002 |
| Sspon.03G0037900 | 1,894 | 0,0002 |
| Sspon.03G0038290 | 3,100 | 0,0002 |
| Sspon.03G0038320 | -1,557 | 0,0002 |
| Sspon.03G0038370 | 2,852 | 0,0002 |
| Sspon.03G0038380 | 4,191 | 0,0002 |
| Sspon.03G0038420 | 17,846 | 0,0002 |
| Sspon.03G0038450 | -3,324 | 0,0002 |
| Sspon.03G0038580 | 1,550 | 0,0002 |
| Sspon.03G0038590 | 5,501 | 0,0002 |
| Sspon.03G0038620 | 17,028 | 0,0002 |
| Sspon.03G0038740 | 8,974 | 0,0002 |
| Sspon.03G0038800 | 8,011 | 0,0002 |
| Sspon.03G0038850 | 5,458 | 0,0002 |
| Sspon.03G0039020 | -3,026 | 0,0002 |
| Sspon.03G0039030 | 2,102 | 0,0002 |
| Sspon.03G0039060 | 1,585 | 0,0002 |
| Sspon.03G0039070 | 1,354 | 0,0002 |
| Sspon.03G0039110 | -6,190 | 0,0002 |
| Sspon.03G0039150 | 1,188 | 0,0002 |
| Sspon.03G0039170 | 1,284 | 0,0002 |
| Sspon.03G0039270 | -2,459 | 0,0002 |
| Sspon.03G0039300 | -1,121 | 0,0002 |
| Sspon.03G0039350 | 1,522 | 0,0002 |
| Sspon.03G0039360 | 1,909 | 0,0002 |
| Sspon.03G0039430 | -9,700 | 0,0002 |
| Sspon.03G0039440 | 17,423 | 0,0002 |
| Sspon.03G0039450 | 6,380 | 0,0002 |
| Sspon.03G0039500 | 1,502 | 0,0002 |
| Sspon.03G0039510 | 3,333 | 0,0002 |
| Sspon.03G0039680 | 2,417 | 0,0002 |
| Sspon.03G0039710 | 1,535 | 0,0002 |
| Sspon.03G0039740 | -1,124 | 0,0002 |
| Sspon.03G0039750 | 1,399 | 0,0002 |
| Sspon.03G0039800 | 15,708 | 0,0002 |
| Sspon.03G0039830 | -1,436 | 0,0002 |
| Sspon.03G0039850 | 2,398 | 0,0002 |
| Sspon.03G0039860 | 1,227 | 0,0002 |
| Sspon.03G0039890 | 3,409 | 0,0002 |
| Sspon.03G0039950 | 1,468 | 0,0002 |
| Sspon.03G0040040 | -1,340 | 0,0002 |
| Sspon.03G0040130 | -2,475 | 0,0002 |
| Sspon.03G0040140 | 1,238 | 0,0002 |

| | | |
|---|---|---|
| Sspon.03G0040160 | 3,301 | 0,0002 |
| Sspon.03G0040260 | 1,770 | 0,0002 |
| Sspon.03G0040450 | 2,232 | 0,0002 |
| Sspon.03G0040650 | 9,315 | 0,0002 |
| Sspon.03G0040670 | 5,675 | 0,0002 |
| Sspon.03G0040690 | 2,624 | 0,0002 |
| Sspon.03G0040750 | 1,901 | 0,0002 |
| Sspon.03G0040760 | 2,498 | 0,0002 |
| Sspon.03G0040830 | 3,063 | 0,0002 |
| Sspon.03G0040880 | 1,569 | 0,0002 |
| Sspon.03G0040900 | 2,295 | 0,0002 |
| Sspon.03G0040910 | 1,810 | 0,0002 |
| Sspon.03G0041110 | 1,883 | 0,0002 |
| Sspon.03G0041200 | -1,308 | 0,0002 |
| Sspon.03G0041210 | 1,651 | 0,0002 |
| Sspon.03G0041420 | 8,876 | 0,0002 |
| Sspon.03G0041440 | 4,247 | 0,0002 |
| Sspon.03G0041470 | 3,123 | 0,0002 |
| Sspon.03G0041580 | 1,819 | 0,0002 |
| Sspon.03G0041590 | 1,829 | 0,0002 |
| Sspon.03G0041660 | -1,385 | 0,0002 |
| Sspon.03G0041720 | -9,654 | 0,0002 |
| Sspon.03G0041750 | -1,374 | 0,0002 |
| Sspon.03G0041840 | 1,705 | 0,0002 |
| Sspon.03G0041850 | -1,600 | 0,0002 |
| Sspon.03G0041880 | 1,105 | 0,0002 |
| Sspon.03G0041890 | 1,758 | 0,0002 |
| Sspon.03G0041960 | -1,146 | 0,0002 |
| Sspon.03G0042140 | 1,942 | 0,0002 |
| Sspon.03G0042240 | -1,477 | 0,0002 |
| Sspon.03G0042280 | 1,778 | 0,0002 |
| Sspon.03G0042330 | -1,400 | 0,0002 |
| Sspon.03G0042360 | 1,516 | 0,0002 |
| Sspon.03G0042370 | 1,211 | 0,0002 |
| Sspon.03G0042380 | 1,685 | 0,0002 |
| Sspon.03G0042390 | -1,780 | 0,0002 |
| Sspon.03G0042440 | -1,084 | 0,0002 |
| Sspon.03G0042830 | 1,670 | 0,0002 |
| Sspon.03G0042960 | 3,277 | 0,0002 |
| Sspon.03G0042980 | 7,914 | 0,0002 |
| Sspon.03G0043060 | -15,761 | 0,0002 |
| Sspon.03G0043070 | 1,738 | 0,0002 |
| Sspon.03G0043080 | 16,239 | 0,0002 |

| | | |
|---|---|---|
| Sspon.03G0043230 | 9,271 | 0,0002 |
| Sspon.03G0043410 | -1,744 | 0,0002 |
| Sspon.03G0043440 | -1,825 | 0,0002 |
| Sspon.03G0043450 | 2,078 | 0,0002 |
| Sspon.03G0043480 | -2,604 | 0,0002 |
| Sspon.03G0043540 | -4,903 | 0,0002 |
| Sspon.03G0043590 | -1,305 | 0,0002 |
| Sspon.03G0043600 | 15,663 | 0,0002 |
| Sspon.03G0043700 | 4,120 | 0,0002 |
| Sspon.03G0043760 | 6,986 | 0,0002 |
| Sspon.03G0043850 | 2,614 | 0,0002 |
| Sspon.03G0043910 | -1,662 | 0,0002 |
| Sspon.03G0044030 | -1,665 | 0,0002 |
| Sspon.03G0044200 | 8,011 | 0,0002 |
| Sspon.03G0044220 | 2,498 | 0,0002 |
| Sspon.03G0044250 | 2,820 | 0,0002 |
| Sspon.03G0044290 | 9,117 | 0,0002 |
| Sspon.03G0044330 | -1,910 | 0,0002 |
| Sspon.03G0044370 | 1,368 | 0,0002 |
| Sspon.03G0044520 | 1,207 | 0,0002 |
| Sspon.03G0044540 | 1,936 | 0,0002 |
| Sspon.03G0044680 | 2,414 | 0,0002 |
| Sspon.03G0044690 | -1,096 | 0,0002 |
| Sspon.03G0044940 | 1,386 | 0,0002 |
| Sspon.03G0044980 | -1,094 | 0,0002 |
| Sspon.03G0045050 | 5,335 | 0,0002 |
| Sspon.03G0045200 | 1,391 | 0,0002 |
| Sspon.03G0045270 | -1,648 | 0,0002 |
| Sspon.03G0045290 | 1,392 | 0,0002 |
| Sspon.03G0045300 | 17,326 | 0,0002 |
| Sspon.03G0045360 | -1,081 | 0,0002 |
| Sspon.03G0045420 | 1,100 | 0,0002 |
| Sspon.03G0045610 | 4,262 | 0,0002 |
| Sspon.03G0045620 | -3,964 | 0,0002 |
| Sspon.03G0045660 | -1,490 | 0,0002 |
| Sspon.03G0045680 | 1,681 | 0,0002 |
| Sspon.03G0045740 | 1,380 | 0,0002 |
| Sspon.03G0045760 | 3,050 | 0,0002 |
| Sspon.03G0045830 | 1,840 | 0,0002 |
| Sspon.03G0045890 | 1,734 | 0,0002 |
| Sspon.03G0045930 | 2,893 | 0,0002 |
| Sspon.03G0046150 | -1,697 | 0,0002 |
| Sspon.03G0046580 | -4,552 | 0,0002 |

| | | |
|---|---|---|
| Sspon.03G0046620 | 7,895 | 0,0002 |
| Sspon.03G0046660 | 1,404 | 0,0002 |
| Sspon.03G0046690 | -1,759 | 0,0002 |
| Sspon.03G0046930 | -1,872 | 0,0002 |
| Sspon.03G0047080 | 2,494 | 0,0002 |
| Sspon.03G0047090 | 3,349 | 0,0002 |
| Sspon.03G0047270 | -8,670 | 0,0002 |
| Sspon.03G0047340 | -1,357 | 0,0002 |
| Sspon.03G0047360 | -1,947 | 0,0002 |
| Sspon.03G0047390 | -1,533 | 0,0002 |
| Sspon.03G0047430 | 6,912 | 0,0002 |
| Sspon.03G0047480 | 1,127 | 0,0002 |
| Sspon.03G0047500 | 16,422 | 0,0002 |
| Sspon.04G0000060 | -1,916 | 0,0002 |
| Sspon.04G0000070 | 1,536 | 0,0002 |
| Sspon.04G0000150 | 1,207 | 0,0002 |
| Sspon.04G0000180 | -1,027 | 0,0002 |
| Sspon.04G0000240 | 2,877 | 0,0002 |
| Sspon.04G0000270 | 1,650 | 0,0002 |
| Sspon.04G0000460 | -1,284 | 0,0002 |
| Sspon.04G0000570 | 1,306 | 0,0002 |
| Sspon.04G0000640 | 2,393 | 0,0002 |
| Sspon.04G0000720 | -1,798 | 0,0002 |
| Sspon.04G0000820 | 1,897 | 0,0002 |
| Sspon.04G0000950 | -1,385 | 0,0002 |
| Sspon.04G0000960 | 1,063 | 0,0002 |
| Sspon.04G0000970 | 15,532 | 0,0002 |
| Sspon.04G0000980 | -2,592 | 0,0002 |
| Sspon.04G0000990 | 2,344 | 0,0002 |
| Sspon.04G0001000 | -1,233 | 0,0002 |
| Sspon.04G0001010 | 3,186 | 0,0002 |
| Sspon.04G0001020 | 2,026 | 0,0002 |
| Sspon.04G0001110 | 4,428 | 0,0002 |
| Sspon.04G0001140 | 1,729 | 0,0002 |
| Sspon.04G0001190 | -2,681 | 0,0002 |
| Sspon.04G0001270 | 1,004 | 0,0002 |
| Sspon.04G0001340 | -2,009 | 0,0002 |
| Sspon.04G0001570 | 1,029 | 0,0002 |
| Sspon.04G0001670 | 1,757 | 0,0002 |
| Sspon.04G0001700 | 2,194 | 0,0002 |
| Sspon.04G0001770 | 2,887 | 0,0002 |
| Sspon.04G0001880 | -2,284 | 0,0002 |
| Sspon.04G0001910 | 3,802 | 0,0002 |

| | | |
|---|---|---|
| Sspon.04G0001990 | -1,638 | 0,0002 |
| Sspon.04G0002070 | 2,290 | 0,0002 |
| Sspon.04G0002100 | -3,123 | 0,0002 |
| Sspon.04G0002160 | 1,440 | 0,0002 |
| Sspon.04G0002230 | -1,148 | 0,0002 |
| Sspon.04G0002240 | 2,215 | 0,0002 |
| Sspon.04G0002330 | 10,179 | 0,0002 |
| Sspon.04G0002390 | -7,423 | 0,0002 |
| Sspon.04G0002500 | 1,324 | 0,0002 |
| Sspon.04G0002510 | 1,746 | 0,0002 |
| Sspon.04G0002530 | -1,476 | 0,0002 |
| Sspon.04G0002590 | 2,990 | 0,0002 |
| Sspon.04G0002600 | -4,371 | 0,0002 |
| Sspon.04G0002610 | 2,012 | 0,0002 |
| Sspon.04G0002690 | -1,012 | 0,0002 |
| Sspon.04G0002730 | -2,044 | 0,0002 |
| Sspon.04G0002940 | 15,690 | 0,0002 |
| Sspon.04G0002960 | -3,205 | 0,0002 |
| Sspon.04G0002980 | -1,455 | 0,0002 |
| Sspon.04G0003000 | -3,014 | 0,0002 |
| Sspon.04G0003030 | 1,141 | 0,0002 |
| Sspon.04G0003110 | 2,101 | 0,0002 |
| Sspon.04G0003230 | -8,786 | 0,0002 |
| Sspon.04G0003250 | 2,227 | 0,0002 |
| Sspon.04G0003270 | 2,954 | 0,0002 |
| Sspon.04G0003500 | 1,543 | 0,0003 |
| Sspon.04G0003610 | -1,244 | 0,0003 |
| Sspon.04G0003690 | 3,196 | 0,0003 |
| Sspon.04G0003700 | 3,722 | 0,0003 |
| Sspon.04G0003740 | 1,655 | 0,0003 |
| Sspon.04G0003760 | -3,781 | 0,0003 |
| Sspon.04G0003800 | 1,927 | 0,0003 |
| Sspon.04G0003820 | -1,060 | 0,0003 |
| Sspon.04G0003850 | -2,719 | 0,0003 |
| Sspon.04G0003860 | -1,088 | 0,0003 |
| Sspon.04G0003890 | 1,776 | 0,0003 |
| Sspon.04G0004080 | -3,058 | 0,0003 |
| Sspon.04G0004230 | -2,152 | 0,0003 |
| Sspon.04G0004290 | -5,937 | 0,0003 |
| Sspon.04G0004400 | -1,782 | 0,0003 |
| Sspon.04G0004450 | 1,358 | 0,0003 |
| Sspon.04G0004530 | 9,614 | 0,0003 |
| Sspon.04G0004590 | 1,627 | 0,0003 |

| | | |
|---|---|---|
| Sspon.04G0004600 | -1,239 | 0,0003 |
| Sspon.04G0004630 | 15,322 | 0,0003 |
| Sspon.04G0004700 | 2,348 | 0,0003 |
| Sspon.04G0004740 | 1,667 | 0,0003 |
| Sspon.04G0004760 | -1,406 | 0,0003 |
| Sspon.04G0004850 | 5,417 | 0,0003 |
| Sspon.04G0004910 | 1,106 | 0,0003 |
| Sspon.04G0004940 | 1,510 | 0,0003 |
| Sspon.04G0005000 | -6,089 | 0,0003 |
| Sspon.04G0005010 | 3,162 | 0,0003 |
| Sspon.04G0005020 | 1,209 | 0,0003 |
| Sspon.04G0005050 | 1,500 | 0,0003 |
| Sspon.04G0005060 | 1,110 | 0,0003 |
| Sspon.04G0005190 | 6,794 | 0,0003 |
| Sspon.04G0005250 | 1,286 | 0,0003 |
| Sspon.04G0005260 | 1,876 | 0,0003 |
| Sspon.04G0005270 | 1,503 | 0,0003 |
| Sspon.04G0005280 | 1,608 | 0,0003 |
| Sspon.04G0005320 | 1,853 | 0,0003 |
| Sspon.04G0005350 | 1,029 | 0,0003 |
| Sspon.04G0005380 | 1,219 | 0,0003 |
| Sspon.04G0005520 | 1,619 | 0,0003 |
| Sspon.04G0005530 | 1,270 | 0,0003 |
| Sspon.04G0005570 | -1,743 | 0,0003 |
| Sspon.04G0005590 | 1,050 | 0,0003 |
| Sspon.04G0005700 | -2,187 | 0,0003 |
| Sspon.04G0005740 | 1,808 | 0,0003 |
| Sspon.04G0005780 | 5,467 | 0,0003 |
| Sspon.04G0005860 | 2,027 | 0,0003 |
| Sspon.04G0005900 | -2,680 | 0,0003 |
| Sspon.04G0005990 | 1,641 | 0,0003 |
| Sspon.04G0006000 | 2,593 | 0,0003 |
| Sspon.04G0006120 | -1,258 | 0,0003 |
| Sspon.04G0006150 | 1,702 | 0,0003 |
| Sspon.04G0006220 | -1,052 | 0,0003 |
| Sspon.04G0006230 | 2,613 | 0,0003 |
| Sspon.04G0006240 | -9,183 | 0,0003 |
| Sspon.04G0006270 | -3,328 | 0,0003 |
| Sspon.04G0006320 | 1,366 | 0,0003 |
| Sspon.04G0006340 | 3,256 | 0,0003 |
| Sspon.04G0006520 | 2,818 | 0,0003 |
| Sspon.04G0006580 | 1,411 | 0,0003 |
| Sspon.04G0006620 | 1,404 | 0,0003 |

| | | |
|---|---|---|
| Sspon.04G0006710 | -1,765 | 0,0003 |
| Sspon.04G0006890 | 1,170 | 0,0003 |
| Sspon.04G0006900 | 1,121 | 0,0003 |
| Sspon.04G0006940 | -1,291 | 0,0003 |
| Sspon.04G0006980 | -2,817 | 0,0003 |
| Sspon.04G0007040 | 1,587 | 0,0003 |
| Sspon.04G0007060 | -1,803 | 0,0003 |
| Sspon.04G0007090 | 1,030 | 0,0003 |
| Sspon.04G0007180 | -1,409 | 0,0003 |
| Sspon.04G0007190 | 1,894 | 0,0003 |
| Sspon.04G0007240 | 2,263 | 0,0003 |
| Sspon.04G0007280 | 1,842 | 0,0003 |
| Sspon.04G0007320 | 6,833 | 0,0003 |
| Sspon.04G0007360 | -1,182 | 0,0003 |
| Sspon.04G0007410 | -1,478 | 0,0003 |
| Sspon.04G0007590 | 1,843 | 0,0003 |
| Sspon.04G0007630 | 4,166 | 0,0003 |
| Sspon.04G0007650 | 1,494 | 0,0003 |
| Sspon.04G0007710 | 1,027 | 0,0003 |
| Sspon.04G0007830 | -6,086 | 0,0003 |
| Sspon.04G0007860 | 1,212 | 0,0003 |
| Sspon.04G0007940 | 2,808 | 0,0003 |
| Sspon.04G0007950 | 2,285 | 0,0003 |
| Sspon.04G0008020 | 1,213 | 0,0003 |
| Sspon.04G0008050 | -17,552 | 0,0003 |
| Sspon.04G0008080 | 15,849 | 0,0003 |
| Sspon.04G0008150 | 1,163 | 0,0003 |
| Sspon.04G0008170 | 1,037 | 0,0003 |
| Sspon.04G0008190 | -1,080 | 0,0003 |
| Sspon.04G0008200 | 2,085 | 0,0003 |
| Sspon.04G0008210 | 1,704 | 0,0003 |
| Sspon.04G0008240 | 16,746 | 0,0003 |
| Sspon.04G0008260 | 2,487 | 0,0003 |
| Sspon.04G0008290 | -1,730 | 0,0003 |
| Sspon.04G0008300 | 8,499 | 0,0003 |
| Sspon.04G0008450 | -1,010 | 0,0003 |
| Sspon.04G0008500 | -3,811 | 0,0003 |
| Sspon.04G0008670 | -1,092 | 0,0003 |
| Sspon.04G0008850 | 8,553 | 0,0003 |
| Sspon.04G0008880 | 4,077 | 0,0003 |
| Sspon.04G0008940 | -10,343 | 0,0003 |
| Sspon.04G0008980 | -1,223 | 0,0003 |
| Sspon.04G0009110 | 4,676 | 0,0003 |

| | | |
|---|---|---|
| Sspon.04G0009160 | -4,330 | 0,0003 |
| Sspon.04G0009220 | 2,624 | 0,0003 |
| Sspon.04G0009240 | 8,959 | 0,0003 |
| Sspon.04G0009270 | 3,848 | 0,0003 |
| Sspon.04G0009460 | 1,312 | 0,0003 |
| Sspon.04G0009510 | -1,059 | 0,0003 |
| Sspon.04G0009590 | -1,096 | 0,0003 |
| Sspon.04G0009620 | -1,792 | 0,0003 |
| Sspon.04G0009640 | 1,205 | 0,0003 |
| Sspon.04G0009690 | -1,705 | 0,0003 |
| Sspon.04G0009710 | -1,452 | 0,0003 |
| Sspon.04G0009720 | 4,092 | 0,0003 |
| Sspon.04G0009770 | -9,695 | 0,0003 |
| Sspon.04G0009870 | 16,957 | 0,0003 |
| Sspon.04G0009880 | 1,708 | 0,0003 |
| Sspon.04G0009920 | 1,234 | 0,0003 |
| Sspon.04G0009950 | 1,219 | 0,0003 |
| Sspon.04G0010000 | -1,800 | 0,0003 |
| Sspon.04G0010020 | -1,278 | 0,0003 |
| Sspon.04G0010030 | -1,259 | 0,0003 |
| Sspon.04G0010040 | 15,382 | 0,0003 |
| Sspon.04G0010060 | -4,147 | 0,0003 |
| Sspon.04G0010210 | 1,996 | 0,0003 |
| Sspon.04G0010290 | 1,244 | 0,0003 |
| Sspon.04G0010340 | 1,929 | 0,0003 |
| Sspon.04G0010370 | 1,202 | 0,0003 |
| Sspon.04G0010380 | 3,940 | 0,0003 |
| Sspon.04G0010430 | 8,678 | 0,0003 |
| Sspon.04G0010500 | -1,930 | 0,0003 |
| Sspon.04G0010510 | 1,483 | 0,0003 |
| Sspon.04G0010530 | 1,561 | 0,0003 |
| Sspon.04G0010590 | 1,874 | 0,0003 |
| Sspon.04G0010600 | -2,659 | 0,0003 |
| Sspon.04G0010700 | 2,717 | 0,0003 |
| Sspon.04G0010730 | -9,347 | 0,0003 |
| Sspon.04G0010830 | 2,817 | 0,0003 |
| Sspon.04G0010890 | 7,538 | 0,0003 |
| Sspon.04G0010960 | -1,229 | 0,0003 |
| Sspon.04G0010980 | -2,316 | 0,0003 |
| Sspon.04G0011020 | 1,181 | 0,0003 |
| Sspon.04G0011030 | 16,931 | 0,0003 |
| Sspon.04G0011050 | 1,621 | 0,0003 |
| Sspon.04G0011130 | -2,036 | 0,0003 |

| | | |
|---|---|---|
| Sspon.04G0011180 | -1,304 | 0,0003 |
| Sspon.04G0011240 | -2,415 | 0,0003 |
| Sspon.04G0011330 | -6,743 | 0,0003 |
| Sspon.04G0011380 | 1,622 | 0,0003 |
| Sspon.04G0011460 | 1,803 | 0,0003 |
| Sspon.04G0011490 | -1,122 | 0,0003 |
| Sspon.04G0011590 | -3,958 | 0,0003 |
| Sspon.04G0011660 | 2,791 | 0,0003 |
| Sspon.04G0011710 | 2,161 | 0,0003 |
| Sspon.04G0011730 | 1,360 | 0,0003 |
| Sspon.04G0011910 | 5,426 | 0,0003 |
| Sspon.04G0011940 | 2,396 | 0,0003 |
| Sspon.04G0011990 | 4,449 | 0,0003 |
| Sspon.04G0012000 | 4,351 | 0,0003 |
| Sspon.04G0012020 | 1,698 | 0,0003 |
| Sspon.04G0012050 | -1,592 | 0,0003 |
| Sspon.04G0012090 | -3,627 | 0,0004 |
| Sspon.04G0012130 | 1,112 | 0,0004 |
| Sspon.04G0012140 | 1,241 | 0,0004 |
| Sspon.04G0012150 | -1,131 | 0,0004 |
| Sspon.04G0012200 | -1,019 | 0,0004 |
| Sspon.04G0012320 | -1,645 | 0,0004 |
| Sspon.04G0012370 | 8,731 | 0,0004 |
| Sspon.04G0012420 | 2,137 | 0,0004 |
| Sspon.04G0012440 | -1,390 | 0,0004 |
| Sspon.04G0012620 | 1,035 | 0,0004 |
| Sspon.04G0012630 | -2,677 | 0,0004 |
| Sspon.04G0012680 | -2,935 | 0,0004 |
| Sspon.04G0012760 | 5,620 | 0,0004 |
| Sspon.04G0012850 | -1,866 | 0,0004 |
| Sspon.04G0012860 | 1,361 | 0,0004 |
| Sspon.04G0012880 | -1,456 | 0,0004 |
| Sspon.04G0012920 | -1,233 | 0,0004 |
| Sspon.04G0012990 | -2,169 | 0,0004 |
| Sspon.04G0013030 | 5,276 | 0,0004 |
| Sspon.04G0013040 | 15,801 | 0,0004 |
| Sspon.04G0013080 | -5,428 | 0,0004 |
| Sspon.04G0013100 | -3,105 | 0,0004 |
| Sspon.04G0013150 | -1,250 | 0,0004 |
| Sspon.04G0013160 | 8,495 | 0,0004 |
| Sspon.04G0013170 | 7,155 | 0,0004 |
| Sspon.04G0013200 | 1,757 | 0,0004 |
| Sspon.04G0013280 | 8,350 | 0,0004 |

| | | |
|---|---|---|
| Sspon.04G0013320 | 4,227 | 0,0004 |
| Sspon.04G0013350 | 1,128 | 0,0004 |
| Sspon.04G0013410 | -1,028 | 0,0004 |
| Sspon.04G0013460 | 5,377 | 0,0004 |
| Sspon.04G0013550 | 1,207 | 0,0004 |
| Sspon.04G0013630 | 1,383 | 0,0004 |
| Sspon.04G0013710 | 16,642 | 0,0004 |
| Sspon.04G0013770 | 5,931 | 0,0004 |
| Sspon.04G0013800 | 8,634 | 0,0004 |
| Sspon.04G0013810 | 2,270 | 0,0004 |
| Sspon.04G0013860 | -6,098 | 0,0004 |
| Sspon.04G0013970 | 2,103 | 0,0004 |
| Sspon.04G0014030 | -3,095 | 0,0004 |
| Sspon.04G0014080 | 2,547 | 0,0004 |
| Sspon.04G0014150 | 2,457 | 0,0004 |
| Sspon.04G0014320 | 5,757 | 0,0004 |
| Sspon.04G0014370 | 1,005 | 0,0004 |
| Sspon.04G0014400 | 1,491 | 0,0004 |
| Sspon.04G0014430 | -2,243 | 0,0004 |
| Sspon.04G0014450 | -2,723 | 0,0004 |
| Sspon.04G0014600 | 5,655 | 0,0004 |
| Sspon.04G0014610 | 8,436 | 0,0004 |
| Sspon.04G0014650 | 1,839 | 0,0004 |
| Sspon.04G0014660 | 3,646 | 0,0004 |
| Sspon.04G0014700 | 5,359 | 0,0004 |
| Sspon.04G0014820 | 1,556 | 0,0004 |
| Sspon.04G0014890 | 1,045 | 0,0004 |
| Sspon.04G0014940 | -6,408 | 0,0004 |
| Sspon.04G0014990 | 1,550 | 0,0004 |
| Sspon.04G0015100 | -8,852 | 0,0004 |
| Sspon.04G0015130 | 18,252 | 0,0004 |
| Sspon.04G0015140 | -3,326 | 0,0004 |
| Sspon.04G0015250 | -1,650 | 0,0004 |
| Sspon.04G0015330 | 3,628 | 0,0004 |
| Sspon.04G0015380 | 1,024 | 0,0004 |
| Sspon.04G0015500 | 3,318 | 0,0004 |
| Sspon.04G0015560 | 3,763 | 0,0004 |
| Sspon.04G0015570 | 1,421 | 0,0004 |
| Sspon.04G0015690 | -8,554 | 0,0004 |
| Sspon.04G0015810 | 2,972 | 0,0004 |
| Sspon.04G0015970 | -1,665 | 0,0004 |
| Sspon.04G0016010 | 1,296 | 0,0004 |
| Sspon.04G0016160 | 3,544 | 0,0004 |

| | | |
|---|---|---|
| Sspon.04G0016180 | 1,297 | 0,0004 |
| Sspon.04G0016200 | -3,269 | 0,0004 |
| Sspon.04G0016280 | 1,135 | 0,0004 |
| Sspon.04G0016290 | 1,312 | 0,0004 |
| Sspon.04G0016330 | 1,520 | 0,0004 |
| Sspon.04G0016380 | 2,072 | 0,0004 |
| Sspon.04G0016410 | -1,041 | 0,0004 |
| Sspon.04G0016430 | 6,505 | 0,0004 |
| Sspon.04G0016540 | 1,318 | 0,0004 |
| Sspon.04G0016600 | -1,020 | 0,0004 |
| Sspon.04G0016610 | -1,018 | 0,0004 |
| Sspon.04G0016640 | -7,387 | 0,0004 |
| Sspon.04G0016750 | -4,075 | 0,0004 |
| Sspon.04G0016790 | 1,079 | 0,0004 |
| Sspon.04G0017010 | 1,679 | 0,0004 |
| Sspon.04G0017050 | 2,723 | 0,0004 |
| Sspon.04G0017080 | 2,245 | 0,0004 |
| Sspon.04G0017150 | -2,456 | 0,0004 |
| Sspon.04G0017170 | -1,443 | 0,0004 |
| Sspon.04G0017370 | 3,508 | 0,0004 |
| Sspon.04G0017470 | 1,659 | 0,0004 |
| Sspon.04G0017570 | -1,162 | 0,0005 |
| Sspon.04G0017580 | 1,929 | 0,0005 |
| Sspon.04G0017620 | -1,506 | 0,0005 |
| Sspon.04G0017640 | 15,541 | 0,0005 |
| Sspon.04G0017660 | 2,944 | 0,0005 |
| Sspon.04G0017780 | -1,530 | 0,0005 |
| Sspon.04G0017830 | 2,209 | 0,0005 |
| Sspon.04G0017870 | 1,431 | 0,0005 |
| Sspon.04G0017900 | -8,760 | 0,0005 |
| Sspon.04G0017910 | -1,469 | 0,0005 |
| Sspon.04G0017950 | 1,091 | 0,0005 |
| Sspon.04G0017970 | -1,229 | 0,0005 |
| Sspon.04G0018050 | 2,511 | 0,0005 |
| Sspon.04G0018200 | -1,652 | 0,0005 |
| Sspon.04G0018220 | -1,668 | 0,0005 |
| Sspon.04G0018250 | 1,526 | 0,0005 |
| Sspon.04G0018350 | -1,537 | 0,0005 |
| Sspon.04G0018390 | 8,273 | 0,0005 |
| Sspon.04G0018450 | -1,648 | 0,0005 |
| Sspon.04G0018500 | -1,475 | 0,0005 |
| Sspon.04G0018520 | 1,745 | 0,0005 |
| Sspon.04G0018580 | -2,579 | 0,0005 |

| | | |
|---|---|---|
| Sspon.04G0018610 | -7,296 | 0,0005 |
| Sspon.04G0018720 | 2,921 | 0,0005 |
| Sspon.04G0018820 | -1,588 | 0,0005 |
| Sspon.04G0018860 | -1,085 | 0,0005 |
| Sspon.04G0018880 | -9,404 | 0,0005 |
| Sspon.04G0018890 | 1,056 | 0,0005 |
| Sspon.04G0018900 | 2,578 | 0,0005 |
| Sspon.04G0018980 | 1,706 | 0,0005 |
| Sspon.04G0019030 | 1,501 | 0,0005 |
| Sspon.04G0019060 | 1,388 | 0,0005 |
| Sspon.04G0019220 | 3,327 | 0,0005 |
| Sspon.04G0019470 | 2,329 | 0,0005 |
| Sspon.04G0019480 | 1,509 | 0,0005 |
| Sspon.04G0019490 | 1,073 | 0,0005 |
| Sspon.04G0019500 | -2,881 | 0,0005 |
| Sspon.04G0019510 | 1,135 | 0,0005 |
| Sspon.04G0019610 | -3,430 | 0,0005 |
| Sspon.04G0019620 | -1,555 | 0,0005 |
| Sspon.04G0019660 | 8,754 | 0,0005 |
| Sspon.04G0019780 | 1,440 | 0,0005 |
| Sspon.04G0019840 | 1,921 | 0,0005 |
| Sspon.04G0019880 | -2,624 | 0,0005 |
| Sspon.04G0020070 | 2,814 | 0,0005 |
| Sspon.04G0020140 | -2,979 | 0,0005 |
| Sspon.04G0020150 | 1,474 | 0,0005 |
| Sspon.04G0020160 | -1,008 | 0,0005 |
| Sspon.04G0020280 | -1,209 | 0,0005 |
| Sspon.04G0020290 | 2,668 | 0,0005 |
| Sspon.04G0020370 | -9,360 | 0,0005 |
| Sspon.04G0020420 | 1,487 | 0,0005 |
| Sspon.04G0020460 | -2,564 | 0,0005 |
| Sspon.04G0020500 | 1,016 | 0,0005 |
| Sspon.04G0020540 | -1,884 | 0,0005 |
| Sspon.04G0020570 | -3,041 | 0,0005 |
| Sspon.04G0020600 | 1,324 | 0,0005 |
| Sspon.04G0020730 | 2,171 | 0,0005 |
| Sspon.04G0020750 | -2,579 | 0,0005 |
| Sspon.04G0020780 | 1,711 | 0,0005 |
| Sspon.04G0020820 | 8,414 | 0,0005 |
| Sspon.04G0020850 | -8,384 | 0,0005 |
| Sspon.04G0020870 | 2,889 | 0,0005 |
| Sspon.04G0020880 | 1,122 | 0,0005 |
| Sspon.04G0021100 | 1,227 | 0,0005 |

| | | |
|---|---|---|
| Sspon.04G0021320 | 2,037 | 0,0005 |
| Sspon.04G0021330 | 2,074 | 0,0005 |
| Sspon.04G0021340 | 1,000 | 0,0005 |
| Sspon.04G0021390 | 1,667 | 0,0005 |
| Sspon.04G0021420 | 8,121 | 0,0005 |
| Sspon.04G0021500 | 1,814 | 0,0005 |
| Sspon.04G0021560 | 1,254 | 0,0005 |
| Sspon.04G0021570 | 1,793 | 0,0005 |
| Sspon.04G0021700 | 1,348 | 0,0005 |
| Sspon.04G0021730 | 1,503 | 0,0005 |
| Sspon.04G0021860 | 6,349 | 0,0005 |
| Sspon.04G0021930 | -1,991 | 0,0005 |
| Sspon.04G0021950 | 1,410 | 0,0005 |
| Sspon.04G0022030 | -1,419 | 0,0005 |
| Sspon.04G0022070 | -1,385 | 0,0005 |
| Sspon.04G0022110 | 1,483 | 0,0005 |
| Sspon.04G0022120 | 1,691 | 0,0005 |
| Sspon.04G0022190 | -2,144 | 0,0005 |
| Sspon.04G0022220 | 2,131 | 0,0005 |
| Sspon.04G0022290 | -3,223 | 0,0005 |
| Sspon.04G0022300 | -1,501 | 0,0005 |
| Sspon.04G0022310 | 1,233 | 0,0006 |
| Sspon.04G0022330 | 1,078 | 0,0006 |
| Sspon.04G0022350 | -3,679 | 0,0006 |
| Sspon.04G0022400 | 1,755 | 0,0006 |
| Sspon.04G0022440 | -1,123 | 0,0006 |
| Sspon.04G0022450 | 1,882 | 0,0006 |
| Sspon.04G0022470 | 3,856 | 0,0006 |
| Sspon.04G0022540 | 2,116 | 0,0006 |
| Sspon.04G0022640 | 1,094 | 0,0006 |
| Sspon.04G0022660 | 3,984 | 0,0006 |
| Sspon.04G0022670 | -1,612 | 0,0006 |
| Sspon.04G0022800 | -1,596 | 0,0006 |
| Sspon.04G0022820 | 7,456 | 0,0006 |
| Sspon.04G0022840 | -3,048 | 0,0006 |
| Sspon.04G0022970 | 1,580 | 0,0006 |
| Sspon.04G0022990 | 3,816 | 0,0006 |
| Sspon.04G0023000 | -1,224 | 0,0006 |
| Sspon.04G0023030 | 1,424 | 0,0006 |
| Sspon.04G0023090 | -1,268 | 0,0006 |
| Sspon.04G0023160 | 2,153 | 0,0006 |
| Sspon.04G0023170 | -1,139 | 0,0006 |
| Sspon.04G0023180 | -1,543 | 0,0006 |

| | | |
|---|---|---|
| Sspon.04G0023210 | 1,013 | 0,0006 |
| Sspon.04G0023240 | 1,109 | 0,0006 |
| Sspon.04G0023260 | -4,767 | 0,0006 |
| Sspon.04G0023300 | 1,995 | 0,0006 |
| Sspon.04G0023370 | 1,213 | 0,0006 |
| Sspon.04G0023400 | -1,199 | 0,0006 |
| Sspon.04G0023470 | -1,095 | 0,0006 |
| Sspon.04G0023530 | 1,728 | 0,0006 |
| Sspon.04G0023570 | 1,032 | 0,0006 |
| Sspon.04G0023580 | 3,771 | 0,0006 |
| Sspon.04G0023650 | -1,672 | 0,0006 |
| Sspon.04G0023750 | 1,288 | 0,0006 |
| Sspon.04G0023800 | 1,236 | 0,0006 |
| Sspon.04G0023820 | 4,563 | 0,0006 |
| Sspon.04G0023850 | 1,482 | 0,0006 |
| Sspon.04G0024060 | 1,521 | 0,0006 |
| Sspon.04G0024080 | -1,863 | 0,0006 |
| Sspon.04G0024140 | 1,204 | 0,0006 |
| Sspon.04G0024150 | -9,228 | 0,0006 |
| Sspon.04G0024180 | -1,504 | 0,0006 |
| Sspon.04G0024200 | 1,231 | 0,0006 |
| Sspon.04G0024240 | 2,289 | 0,0006 |
| Sspon.04G0024250 | 15,597 | 0,0006 |
| Sspon.04G0024340 | 1,592 | 0,0006 |
| Sspon.04G0024410 | 5,488 | 0,0006 |
| Sspon.04G0024430 | -1,281 | 0,0006 |
| Sspon.04G0024510 | -1,225 | 0,0006 |
| Sspon.04G0024540 | 6,068 | 0,0006 |
| Sspon.04G0024550 | 1,174 | 0,0006 |
| Sspon.04G0024880 | -4,775 | 0,0006 |
| Sspon.04G0025050 | -8,699 | 0,0006 |
| Sspon.04G0025070 | 1,812 | 0,0006 |
| Sspon.04G0025120 | 1,702 | 0,0006 |
| Sspon.04G0025170 | 5,172 | 0,0006 |
| Sspon.04G0025420 | 1,478 | 0,0006 |
| Sspon.04G0025430 | -1,075 | 0,0006 |
| Sspon.04G0025470 | 4,570 | 0,0006 |
| Sspon.04G0025510 | -7,020 | 0,0006 |
| Sspon.04G0025540 | -3,522 | 0,0006 |
| Sspon.04G0025730 | 1,500 | 0,0006 |
| Sspon.04G0025740 | 2,469 | 0,0006 |
| Sspon.04G0025750 | 1,071 | 0,0006 |
| Sspon.04G0025800 | 1,036 | 0,0006 |

| | | |
|---|---|---|
| Sspon.04G0025900 | 1,793 | 0,0006 |
| Sspon.04G0026080 | 1,957 | 0,0006 |
| Sspon.04G0026090 | -2,643 | 0,0006 |
| Sspon.04G0026100 | -1,003 | 0,0006 |
| Sspon.04G0026150 | 1,400 | 0,0006 |
| Sspon.04G0026200 | 3,375 | 0,0006 |
| Sspon.04G0026310 | -8,159 | 0,0006 |
| Sspon.04G0026540 | -1,092 | 0,0006 |
| Sspon.04G0026600 | 6,619 | 0,0006 |
| Sspon.04G0026650 | 9,491 | 0,0006 |
| Sspon.04G0026790 | -2,815 | 0,0006 |
| Sspon.04G0026840 | -1,074 | 0,0006 |
| Sspon.04G0026880 | -1,622 | 0,0006 |
| Sspon.04G0026950 | 2,053 | 0,0006 |
| Sspon.04G0027120 | -1,569 | 0,0006 |
| Sspon.04G0027140 | 1,008 | 0,0006 |
| Sspon.04G0027150 | 1,617 | 0,0006 |
| Sspon.04G0027190 | 1,621 | 0,0006 |
| Sspon.04G0027250 | 3,442 | 0,0006 |
| Sspon.04G0027360 | -1,031 | 0,0006 |
| Sspon.04G0027400 | -6,428 | 0,0006 |
| Sspon.04G0027480 | 3,905 | 0,0006 |
| Sspon.04G0027510 | 1,268 | 0,0006 |
| Sspon.04G0027550 | -1,237 | 0,0006 |
| Sspon.04G0027720 | 2,817 | 0,0006 |
| Sspon.04G0027820 | -1,196 | 0,0006 |
| Sspon.04G0027850 | 17,052 | 0,0006 |
| Sspon.04G0027860 | 4,738 | 0,0006 |
| Sspon.04G0027870 | 3,850 | 0,0006 |
| Sspon.04G0027880 | 2,117 | 0,0006 |
| Sspon.04G0027940 | 16,755 | 0,0006 |
| Sspon.04G0028100 | -2,322 | 0,0006 |
| Sspon.04G0028320 | -1,595 | 0,0006 |
| Sspon.04G0028390 | 6,154 | 0,0006 |
| Sspon.04G0028420 | 1,736 | 0,0006 |
| Sspon.04G0028480 | 1,558 | 0,0006 |
| Sspon.04G0028520 | -1,147 | 0,0006 |
| Sspon.04G0028650 | -1,477 | 0,0007 |
| Sspon.04G0028710 | -1,639 | 0,0007 |
| Sspon.04G0028780 | 1,478 | 0,0007 |
| Sspon.04G0028900 | 2,552 | 0,0007 |
| Sspon.04G0028940 | 3,011 | 0,0007 |
| Sspon.04G0028970 | -2,650 | 0,0007 |

| | | |
|---|---|---|
| Sspon.04G0028980 | 1,382 | 0,0007 |
| Sspon.04G0029000 | -1,424 | 0,0007 |
| Sspon.04G0029050 | 1,524 | 0,0007 |
| Sspon.04G0029080 | -1,272 | 0,0007 |
| Sspon.04G0029090 | 1,459 | 0,0007 |
| Sspon.04G0029200 | -1,320 | 0,0007 |
| Sspon.04G0029380 | -1,871 | 0,0007 |
| Sspon.04G0029390 | 2,285 | 0,0007 |
| Sspon.04G0029410 | -1,341 | 0,0007 |
| Sspon.04G0029430 | -1,137 | 0,0007 |
| Sspon.04G0029440 | -2,588 | 0,0007 |
| Sspon.04G0029530 | 1,054 | 0,0007 |
| Sspon.04G0029550 | -5,346 | 0,0007 |
| Sspon.04G0029580 | 1,697 | 0,0007 |
| Sspon.04G0029590 | 1,583 | 0,0007 |
| Sspon.04G0029610 | 1,566 | 0,0007 |
| Sspon.04G0029660 | -1,198 | 0,0007 |
| Sspon.04G0029680 | 3,569 | 0,0007 |
| Sspon.04G0029690 | 1,658 | 0,0007 |
| Sspon.04G0029820 | 1,481 | 0,0007 |
| Sspon.04G0029850 | -2,609 | 0,0007 |
| Sspon.04G0029860 | 3,422 | 0,0007 |
| Sspon.04G0030000 | 1,466 | 0,0007 |
| Sspon.04G0030010 | 1,208 | 0,0007 |
| Sspon.04G0030020 | 1,240 | 0,0007 |
| Sspon.04G0030080 | 4,935 | 0,0007 |
| Sspon.04G0030090 | 1,247 | 0,0007 |
| Sspon.04G0030100 | 1,466 | 0,0007 |
| Sspon.04G0030190 | 1,657 | 0,0007 |
| Sspon.04G0030250 | 7,112 | 0,0007 |
| Sspon.04G0030300 | 3,128 | 0,0007 |
| Sspon.04G0030340 | -1,194 | 0,0007 |
| Sspon.04G0030360 | -3,213 | 0,0007 |
| Sspon.04G0030380 | 1,388 | 0,0007 |
| Sspon.04G0030400 | 3,746 | 0,0007 |
| Sspon.04G0030420 | -2,438 | 0,0007 |
| Sspon.04G0030660 | -3,468 | 0,0007 |
| Sspon.04G0030680 | -1,246 | 0,0007 |
| Sspon.04G0030690 | 8,083 | 0,0007 |
| Sspon.04G0030770 | 1,401 | 0,0007 |
| Sspon.04G0030800 | 2,797 | 0,0007 |
| Sspon.04G0030920 | -1,488 | 0,0007 |
| Sspon.04G0031010 | 6,247 | 0,0007 |

| | | |
|---|---|---|
| Sspon.04G0031030 | 2,033 | 0,0007 |
| Sspon.04G0031050 | 3,872 | 0,0007 |
| Sspon.04G0031060 | -3,966 | 0,0007 |
| Sspon.04G0031150 | -2,599 | 0,0007 |
| Sspon.04G0031160 | 2,179 | 0,0007 |
| Sspon.04G0031170 | 3,037 | 0,0007 |
| Sspon.04G0031240 | 1,119 | 0,0007 |
| Sspon.04G0031350 | 8,043 | 0,0007 |
| Sspon.04G0031380 | -1,664 | 0,0007 |
| Sspon.04G0031410 | 1,287 | 0,0007 |
| Sspon.04G0031440 | -1,308 | 0,0007 |
| Sspon.04G0031580 | -1,227 | 0,0007 |
| Sspon.04G0031590 | -3,745 | 0,0007 |
| Sspon.04G0031600 | 8,272 | 0,0007 |
| Sspon.04G0031610 | 2,133 | 0,0007 |
| Sspon.04G0031640 | -9,055 | 0,0007 |
| Sspon.04G0031680 | 8,580 | 0,0007 |
| Sspon.04G0032010 | -1,143 | 0,0007 |
| Sspon.04G0032050 | -1,004 | 0,0007 |
| Sspon.04G0032070 | 1,018 | 0,0007 |
| Sspon.04G0032120 | 3,825 | 0,0007 |
| Sspon.04G0032240 | 1,135 | 0,0007 |
| Sspon.04G0032270 | -1,203 | 0,0007 |
| Sspon.04G0032290 | 1,084 | 0,0007 |
| Sspon.04G0032380 | -1,340 | 0,0008 |
| Sspon.04G0032410 | 3,424 | 0,0008 |
| Sspon.04G0032530 | -1,499 | 0,0008 |
| Sspon.04G0032540 | 2,489 | 0,0008 |
| Sspon.04G0032590 | 1,861 | 0,0008 |
| Sspon.04G0032760 | 3,587 | 0,0008 |
| Sspon.04G0032770 | 2,031 | 0,0008 |
| Sspon.04G0032860 | 2,115 | 0,0008 |
| Sspon.04G0033030 | 19,056 | 0,0008 |
| Sspon.04G0033040 | 6,046 | 0,0008 |
| Sspon.04G0033080 | 1,813 | 0,0008 |
| Sspon.04G0033100 | 7,167 | 0,0008 |
| Sspon.04G0033130 | -7,818 | 0,0008 |
| Sspon.04G0033220 | -1,372 | 0,0008 |
| Sspon.04G0033400 | -1,166 | 0,0008 |
| Sspon.04G0033410 | 1,645 | 0,0008 |
| Sspon.04G0033420 | 1,402 | 0,0008 |
| Sspon.04G0033570 | 7,008 | 0,0008 |
| Sspon.04G0033670 | 1,383 | 0,0008 |

| | | |
|---|---|---|
| Sspon.04G0033680 | 1,003 | 0,0008 |
| Sspon.04G0033730 | 3,394 | 0,0008 |
| Sspon.04G0033770 | -2,720 | 0,0008 |
| Sspon.04G0033780 | -1,980 | 0,0008 |
| Sspon.04G0033840 | -4,510 | 0,0008 |
| Sspon.04G0033940 | 1,135 | 0,0008 |
| Sspon.04G0033960 | -1,737 | 0,0008 |
| Sspon.04G0033970 | -1,600 | 0,0008 |
| Sspon.04G0034070 | 1,119 | 0,0008 |
| Sspon.04G0034140 | -1,325 | 0,0008 |
| Sspon.04G0034180 | 4,733 | 0,0008 |
| Sspon.04G0034270 | -4,005 | 0,0008 |
| Sspon.04G0034510 | -4,882 | 0,0008 |
| Sspon.04G0034530 | -3,351 | 0,0008 |
| Sspon.04G0034540 | -2,024 | 0,0008 |
| Sspon.04G0034590 | 1,119 | 0,0008 |
| Sspon.04G0034600 | -1,005 | 0,0008 |
| Sspon.04G0034650 | -2,403 | 0,0008 |
| Sspon.04G0034960 | 5,618 | 0,0008 |
| Sspon.04G0034980 | 1,768 | 0,0008 |
| Sspon.04G0035000 | 3,501 | 0,0008 |
| Sspon.04G0035090 | 4,127 | 0,0008 |
| Sspon.04G0035120 | 1,386 | 0,0008 |
| Sspon.04G0035180 | -1,301 | 0,0008 |
| Sspon.04G0035220 | 1,930 | 0,0008 |
| Sspon.04G0035230 | 1,545 | 0,0008 |
| Sspon.04G0035250 | -1,229 | 0,0008 |
| Sspon.04G0035420 | 1,555 | 0,0008 |
| Sspon.04G0035470 | 2,518 | 0,0008 |
| Sspon.04G0035510 | -8,701 | 0,0008 |
| Sspon.04G0035540 | -1,612 | 0,0008 |
| Sspon.04G0035680 | -3,819 | 0,0008 |
| Sspon.04G0035690 | 1,282 | 0,0008 |
| Sspon.04G0035750 | 3,506 | 0,0008 |
| Sspon.04G0035830 | 2,266 | 0,0008 |
| Sspon.04G0035860 | -1,339 | 0,0008 |
| Sspon.04G0035880 | -15,825 | 0,0008 |
| Sspon.04G0035900 | 1,107 | 0,0008 |
| Sspon.04G0036030 | 1,196 | 0,0008 |
| Sspon.04G0036090 | 3,565 | 0,0008 |
| Sspon.04G0036100 | 3,341 | 0,0008 |
| Sspon.04G0036180 | 4,336 | 0,0008 |
| Sspon.04G0036200 | 1,373 | 0,0008 |

| | | |
|---|---|---|
| Sspon.04G0036220 | 1,033 | 0,0008 |
| Sspon.04G0036320 | 7,932 | 0,0008 |
| Sspon.04G0036340 | -1,443 | 0,0008 |
| Sspon.04G0036350 | 2,642 | 0,0008 |
| Sspon.04G0036390 | -8,883 | 0,0009 |
| Sspon.04G0036400 | -3,246 | 0,0009 |
| Sspon.04G0036530 | 1,448 | 0,0009 |
| Sspon.04G0036630 | 1,320 | 0,0009 |
| Sspon.04G0036740 | -1,260 | 0,0009 |
| Sspon.04G0036800 | 1,510 | 0,0009 |
| Sspon.04G0036970 | 1,267 | 0,0009 |
| Sspon.04G0037030 | -1,004 | 0,0009 |
| Sspon.04G0037120 | -7,874 | 0,0009 |
| Sspon.04G0037300 | 1,507 | 0,0009 |
| Sspon.04G0037370 | 6,077 | 0,0009 |
| Sspon.04G0037390 | 3,866 | 0,0009 |
| Sspon.04G0037440 | 1,300 | 0,0009 |
| Sspon.04G0037450 | 6,845 | 0,0009 |
| Sspon.04G0037500 | 2,838 | 0,0009 |
| Sspon.04G0037620 | 3,761 | 0,0009 |
| Sspon.04G0037740 | 2,311 | 0,0009 |
| Sspon.04G0037790 | -2,058 | 0,0009 |
| Sspon.04G0037800 | 1,106 | 0,0009 |
| Sspon.04G0037910 | 2,463 | 0,0009 |
| Sspon.04G0037940 | -2,397 | 0,0009 |
| Sspon.04G0037970 | 1,088 | 0,0009 |
| Sspon.04G0037980 | 2,256 | 0,0009 |
| Sspon.04G0038080 | 1,047 | 0,0009 |
| Sspon.05G0000010 | 16,858 | 0,0009 |
| Sspon.05G0000090 | 2,695 | 0,0009 |
| Sspon.05G0000120 | -2,711 | 0,0009 |
| Sspon.05G0000180 | 1,017 | 0,0009 |
| Sspon.05G0000240 | 1,067 | 0,0009 |
| Sspon.05G0000270 | -1,698 | 0,0009 |
| Sspon.05G0000340 | -3,790 | 0,0009 |
| Sspon.05G0000410 | 2,881 | 0,0009 |
| Sspon.05G0000520 | -7,397 | 0,0009 |
| Sspon.05G0000590 | 3,950 | 0,0009 |
| Sspon.05G0000660 | 5,891 | 0,0009 |
| Sspon.05G0000670 | -2,525 | 0,0009 |
| Sspon.05G0000690 | 1,002 | 0,0009 |
| Sspon.05G0000760 | 1,892 | 0,0009 |
| Sspon.05G0000780 | -1,323 | 0,0009 |

| | | |
|---|---|---|
| Sspon.05G0000820 | 1,717 | 0,0009 |
| Sspon.05G0000890 | 1,206 | 0,0009 |
| Sspon.05G0000970 | 1,849 | 0,0009 |
| Sspon.05G0001030 | -1,812 | 0,0009 |
| Sspon.05G0001070 | 1,307 | 0,0009 |
| Sspon.05G0001210 | 1,523 | 0,0009 |
| Sspon.05G0001360 | -1,952 | 0,0009 |
| Sspon.05G0001460 | -1,696 | 0,0009 |
| Sspon.05G0001500 | 2,780 | 0,0009 |
| Sspon.05G0001610 | -1,474 | 0,0009 |
| Sspon.05G0001710 | 18,842 | 0,0009 |
| Sspon.05G0001830 | 16,612 | 0,0009 |
| Sspon.05G0001840 | 1,222 | 0,0009 |
| Sspon.05G0001850 | 7,060 | 0,0009 |
| Sspon.05G0001980 | 1,556 | 0,0009 |
| Sspon.05G0002070 | -1,005 | 0,0009 |
| Sspon.05G0002180 | -2,699 | 0,0009 |
| Sspon.05G0002250 | -5,696 | 0,0009 |
| Sspon.05G0002310 | 15,178 | 0,0009 |
| Sspon.05G0002360 | 1,961 | 0,0009 |
| Sspon.05G0002370 | 1,147 | 0,0009 |
| Sspon.05G0002460 | 4,860 | 0,0009 |
| Sspon.05G0002520 | 2,303 | 0,0009 |
| Sspon.05G0002590 | -2,215 | 0,0009 |
| Sspon.05G0002610 | -8,570 | 0,0009 |
| Sspon.05G0002720 | 6,789 | 0,0009 |
| Sspon.05G0002740 | 3,470 | 0,0009 |
| Sspon.05G0002780 | 5,541 | 0,0010 |
| Sspon.05G0002840 | -1,325 | 0,0010 |
| Sspon.05G0002870 | -2,557 | 0,0010 |
| Sspon.05G0002900 | 3,810 | 0,0010 |
| Sspon.05G0002910 | 6,035 | 0,0010 |
| Sspon.05G0002930 | 2,184 | 0,0010 |
| Sspon.05G0002990 | 2,330 | 0,0010 |
| Sspon.05G0003010 | -1,930 | 0,0010 |
| Sspon.05G0003070 | 1,436 | 0,0010 |
| Sspon.05G0003080 | 1,731 | 0,0010 |
| Sspon.05G0003230 | 2,396 | 0,0010 |
| Sspon.05G0003270 | 1,442 | 0,0010 |
| Sspon.05G0003290 | 3,673 | 0,0010 |
| Sspon.05G0003320 | 2,443 | 0,0010 |
| Sspon.05G0003340 | -1,105 | 0,0010 |
| Sspon.05G0003350 | -1,628 | 0,0010 |

| | | |
|---|---|---|
| Sspon.05G0003430 | 1,843 | 0,0010 |
| Sspon.05G0003500 | 1,209 | 0,0010 |
| Sspon.05G0003540 | 1,341 | 0,0010 |
| Sspon.05G0003550 | 19,891 | 0,0010 |
| Sspon.05G0003560 | 2,092 | 0,0010 |
| Sspon.05G0003570 | 3,252 | 0,0010 |
| Sspon.05G0003610 | -1,767 | 0,0010 |
| Sspon.05G0003630 | 1,889 | 0,0010 |
| Sspon.05G0003680 | -2,544 | 0,0010 |
| Sspon.05G0003800 | 1,656 | 0,0010 |
| Sspon.05G0003810 | 3,714 | 0,0010 |
| Sspon.05G0003820 | -1,266 | 0,0010 |
| Sspon.05G0004040 | -1,469 | 0,0010 |
| Sspon.05G0004090 | 15,024 | 0,0010 |
| Sspon.05G0004270 | 1,041 | 0,0010 |
| Sspon.05G0004470 | 4,971 | 0,0010 |
| Sspon.05G0004520 | 1,380 | 0,0010 |
| Sspon.05G0004670 | 2,626 | 0,0010 |
| Sspon.05G0004720 | 1,350 | 0,0010 |
| Sspon.05G0004760 | 1,743 | 0,0010 |
| Sspon.05G0004820 | 1,301 | 0,0010 |
| Sspon.05G0004850 | -15,874 | 0,0010 |
| Sspon.05G0004860 | 7,626 | 0,0010 |
| Sspon.05G0004880 | 1,588 | 0,0010 |
| Sspon.05G0005020 | 3,298 | 0,0011 |
| Sspon.05G0005060 | 1,023 | 0,0011 |
| Sspon.05G0005070 | 1,205 | 0,0011 |
| Sspon.05G0005180 | 15,813 | 0,0011 |
| Sspon.05G0005220 | 1,937 | 0,0011 |
| Sspon.05G0005230 | 1,442 | 0,0011 |
| Sspon.05G0005260 | 3,249 | 0,0011 |
| Sspon.05G0005350 | -1,015 | 0,0011 |
| Sspon.05G0005390 | 1,188 | 0,0011 |
| Sspon.05G0005440 | 1,167 | 0,0011 |
| Sspon.05G0005450 | 1,224 | 0,0011 |
| Sspon.05G0005500 | -7,859 | 0,0011 |
| Sspon.05G0005530 | -2,702 | 0,0011 |
| Sspon.05G0005550 | 1,261 | 0,0011 |
| Sspon.05G0005610 | -1,064 | 0,0011 |
| Sspon.05G0005620 | -1,014 | 0,0011 |
| Sspon.05G0005640 | 3,966 | 0,0011 |
| Sspon.05G0005720 | 1,343 | 0,0011 |
| Sspon.05G0005770 | 1,553 | 0,0011 |

| | | |
|---|---|---|
| Sspon.05G0006050 | -4,469 | 0,0011 |
| Sspon.05G0006060 | 1,495 | 0,0011 |
| Sspon.05G0006160 | 15,390 | 0,0011 |
| Sspon.05G0006180 | 1,060 | 0,0011 |
| Sspon.05G0006220 | 6,613 | 0,0011 |
| Sspon.05G0006550 | 6,847 | 0,0011 |
| Sspon.05G0006560 | 7,679 | 0,0011 |
| Sspon.05G0006600 | 1,196 | 0,0011 |
| Sspon.05G0006620 | 15,649 | 0,0011 |
| Sspon.05G0006640 | -1,096 | 0,0011 |
| Sspon.05G0006660 | 7,569 | 0,0011 |
| Sspon.05G0006830 | 2,837 | 0,0011 |
| Sspon.05G0006860 | 1,308 | 0,0011 |
| Sspon.05G0006880 | 5,838 | 0,0011 |
| Sspon.05G0006900 | 6,256 | 0,0011 |
| Sspon.05G0006920 | 2,097 | 0,0011 |
| Sspon.05G0006960 | 5,115 | 0,0011 |
| Sspon.05G0006970 | -1,413 | 0,0011 |
| Sspon.05G0006990 | -2,968 | 0,0011 |
| Sspon.05G0007000 | -1,463 | 0,0011 |
| Sspon.05G0007010 | -6,085 | 0,0011 |
| Sspon.05G0007050 | 1,274 | 0,0011 |
| Sspon.05G0007060 | -6,222 | 0,0011 |
| Sspon.05G0007100 | -2,670 | 0,0011 |
| Sspon.05G0007110 | -1,661 | 0,0011 |
| Sspon.05G0007120 | 7,687 | 0,0011 |
| Sspon.05G0007150 | -1,189 | 0,0011 |
| Sspon.05G0007260 | 4,255 | 0,0011 |
| Sspon.05G0007310 | 15,512 | 0,0011 |
| Sspon.05G0007370 | 14,974 | 0,0011 |
| Sspon.05G0007430 | 3,970 | 0,0011 |
| Sspon.05G0007500 | 1,680 | 0,0011 |
| Sspon.05G0007520 | -1,625 | 0,0011 |
| Sspon.05G0007570 | 1,531 | 0,0012 |
| Sspon.05G0007580 | 1,041 | 0,0012 |
| Sspon.05G0007600 | 1,075 | 0,0012 |
| Sspon.05G0007730 | -1,455 | 0,0012 |
| Sspon.05G0007830 | 2,774 | 0,0012 |
| Sspon.05G0008060 | 1,343 | 0,0012 |
| Sspon.05G0008140 | 1,272 | 0,0012 |
| Sspon.05G0008180 | 15,321 | 0,0012 |
| Sspon.05G0008190 | 1,845 | 0,0012 |
| Sspon.05G0008250 | -1,542 | 0,0012 |

| | | |
|---|---|---|
| Sspon.05G0008290 | 1,100 | 0,0012 |
| Sspon.05G0008300 | -4,248 | 0,0012 |
| Sspon.05G0008320 | 2,400 | 0,0012 |
| Sspon.05G0008340 | 7,889 | 0,0012 |
| Sspon.05G0008350 | 3,375 | 0,0012 |
| Sspon.05G0008440 | 1,279 | 0,0012 |
| Sspon.05G0008500 | 1,428 | 0,0012 |
| Sspon.05G0008510 | 6,780 | 0,0012 |
| Sspon.05G0008540 | 5,149 | 0,0012 |
| Sspon.05G0008580 | -1,329 | 0,0012 |
| Sspon.05G0008610 | 1,652 | 0,0012 |
| Sspon.05G0008650 | -4,432 | 0,0012 |
| Sspon.05G0008710 | -2,269 | 0,0012 |
| Sspon.05G0008980 | 7,753 | 0,0012 |
| Sspon.05G0009020 | -6,191 | 0,0012 |
| Sspon.05G0009140 | 1,487 | 0,0012 |
| Sspon.05G0009330 | 2,574 | 0,0012 |
| Sspon.05G0009340 | -2,654 | 0,0012 |
| Sspon.05G0009400 | -7,079 | 0,0012 |
| Sspon.05G0009420 | 1,653 | 0,0012 |
| Sspon.05G0009440 | 1,526 | 0,0012 |
| Sspon.05G0009540 | 1,630 | 0,0012 |
| Sspon.05G0009880 | -2,678 | 0,0012 |
| Sspon.05G0010010 | 4,085 | 0,0012 |
| Sspon.05G0010120 | -4,385 | 0,0012 |
| Sspon.05G0010170 | -2,357 | 0,0012 |
| Sspon.05G0010230 | 1,077 | 0,0012 |
| Sspon.05G0010240 | -6,980 | 0,0012 |
| Sspon.05G0010270 | 1,800 | 0,0012 |
| Sspon.05G0010330 | -7,651 | 0,0012 |
| Sspon.05G0010410 | 1,662 | 0,0012 |
| Sspon.05G0010440 | 1,183 | 0,0012 |
| Sspon.05G0010460 | 1,349 | 0,0012 |
| Sspon.05G0010550 | 1,444 | 0,0012 |
| Sspon.05G0010720 | 1,064 | 0,0012 |
| Sspon.05G0010850 | 1,454 | 0,0012 |
| Sspon.05G0010870 | 1,564 | 0,0012 |
| Sspon.05G0010970 | -7,043 | 0,0012 |
| Sspon.05G0011020 | 1,154 | 0,0012 |
| Sspon.05G0011040 | 1,389 | 0,0012 |
| Sspon.05G0011050 | 16,458 | 0,0012 |
| Sspon.05G0011060 | 15,719 | 0,0013 |
| Sspon.05G0011080 | 3,645 | 0,0013 |

| | | |
|---|---|---|
| Sspon.05G0011100 | 1,239 | 0,0013 |
| Sspon.05G0011130 | 1,213 | 0,0013 |
| Sspon.05G0011330 | 1,837 | 0,0013 |
| Sspon.05G0011360 | 1,106 | 0,0013 |
| Sspon.05G0011370 | 1,601 | 0,0013 |
| Sspon.05G0011380 | 1,799 | 0,0013 |
| Sspon.05G0011400 | 4,337 | 0,0013 |
| Sspon.05G0011500 | 1,787 | 0,0013 |
| Sspon.05G0011530 | 2,782 | 0,0013 |
| Sspon.05G0011550 | 8,124 | 0,0013 |
| Sspon.05G0011560 | 1,974 | 0,0013 |
| Sspon.05G0011620 | 1,469 | 0,0013 |
| Sspon.05G0011650 | 4,847 | 0,0013 |
| Sspon.05G0011700 | 2,948 | 0,0013 |
| Sspon.05G0011710 | 2,555 | 0,0013 |
| Sspon.05G0011720 | -6,249 | 0,0013 |
| Sspon.05G0011880 | -3,288 | 0,0013 |
| Sspon.05G0011890 | -2,563 | 0,0013 |
| Sspon.05G0012010 | 1,335 | 0,0013 |
| Sspon.05G0012030 | -1,085 | 0,0013 |
| Sspon.05G0012050 | 2,591 | 0,0013 |
| Sspon.05G0012070 | 1,232 | 0,0013 |
| Sspon.05G0012120 | -3,144 | 0,0013 |
| Sspon.05G0012240 | 6,526 | 0,0013 |
| Sspon.05G0012310 | -1,738 | 0,0013 |
| Sspon.05G0012330 | -1,609 | 0,0013 |
| Sspon.05G0012350 | -2,135 | 0,0013 |
| Sspon.05G0012410 | 1,500 | 0,0013 |
| Sspon.05G0012500 | -1,663 | 0,0013 |
| Sspon.05G0012580 | 1,013 | 0,0013 |
| Sspon.05G0012690 | -8,012 | 0,0013 |
| Sspon.05G0012730 | 1,023 | 0,0013 |
| Sspon.05G0012760 | 3,279 | 0,0013 |
| Sspon.05G0012830 | 2,274 | 0,0013 |
| Sspon.05G0012840 | -1,266 | 0,0014 |
| Sspon.05G0012910 | 1,106 | 0,0014 |
| Sspon.05G0012940 | -6,087 | 0,0014 |
| Sspon.05G0013000 | 2,305 | 0,0014 |
| Sspon.05G0013010 | 2,061 | 0,0014 |
| Sspon.05G0013060 | 1,665 | 0,0014 |
| Sspon.05G0013080 | -1,405 | 0,0014 |
| Sspon.05G0013100 | 1,476 | 0,0014 |
| Sspon.05G0013120 | -15,475 | 0,0014 |

| | | |
|---|---|---|
| Sspon.05G0013130 | 2,099 | 0,0014 |
| Sspon.05G0013140 | 2,817 | 0,0014 |
| Sspon.05G0013150 | -1,015 | 0,0014 |
| Sspon.05G0013340 | 3,209 | 0,0014 |
| Sspon.05G0013460 | -1,281 | 0,0014 |
| Sspon.05G0013470 | 18,647 | 0,0014 |
| Sspon.05G0013590 | -8,564 | 0,0014 |
| Sspon.05G0013620 | -7,757 | 0,0014 |
| Sspon.05G0013670 | 3,828 | 0,0014 |
| Sspon.05G0013760 | 1,527 | 0,0014 |
| Sspon.05G0013860 | 1,402 | 0,0014 |
| Sspon.05G0013880 | 3,142 | 0,0014 |
| Sspon.05G0014030 | -1,300 | 0,0014 |
| Sspon.05G0014040 | -3,419 | 0,0014 |
| Sspon.05G0014070 | -2,374 | 0,0014 |
| Sspon.05G0014260 | -1,746 | 0,0014 |
| Sspon.05G0014270 | 4,215 | 0,0014 |
| Sspon.05G0014280 | 1,324 | 0,0014 |
| Sspon.05G0014300 | 4,414 | 0,0014 |
| Sspon.05G0014430 | -3,172 | 0,0014 |
| Sspon.05G0014500 | 1,056 | 0,0015 |
| Sspon.05G0014560 | 5,006 | 0,0015 |
| Sspon.05G0014570 | 4,299 | 0,0015 |
| Sspon.05G0014650 | -5,634 | 0,0015 |
| Sspon.05G0014720 | 1,814 | 0,0015 |
| Sspon.05G0014730 | -5,415 | 0,0015 |
| Sspon.05G0014760 | -1,797 | 0,0015 |
| Sspon.05G0014810 | -1,146 | 0,0015 |
| Sspon.05G0014900 | 7,386 | 0,0015 |
| Sspon.05G0014940 | 2,019 | 0,0015 |
| Sspon.05G0014980 | 17,227 | 0,0015 |
| Sspon.05G0015040 | 1,682 | 0,0015 |
| Sspon.05G0015150 | 6,942 | 0,0015 |
| Sspon.05G0015270 | 1,553 | 0,0015 |
| Sspon.05G0015380 | 1,301 | 0,0015 |
| Sspon.05G0015390 | -1,261 | 0,0015 |
| Sspon.05G0015410 | -1,083 | 0,0015 |
| Sspon.05G0015450 | 2,118 | 0,0015 |
| Sspon.05G0015460 | 1,929 | 0,0015 |
| Sspon.05G0015470 | 18,502 | 0,0015 |
| Sspon.05G0015490 | 2,020 | 0,0015 |
| Sspon.05G0015500 | -1,932 | 0,0015 |
| Sspon.05G0015520 | 1,379 | 0,0016 |

| | | |
|---|---|---|
| Sspon.05G0015540 | 6,739 | 0,0016 |
| Sspon.05G0015550 | 4,548 | 0,0016 |
| Sspon.05G0015680 | 2,864 | 0,0016 |
| Sspon.05G0015690 | -1,676 | 0,0016 |
| Sspon.05G0015720 | -5,983 | 0,0016 |
| Sspon.05G0015740 | -1,859 | 0,0016 |
| Sspon.05G0015760 | 1,013 | 0,0016 |
| Sspon.05G0015800 | 1,275 | 0,0016 |
| Sspon.05G0015830 | 1,062 | 0,0016 |
| Sspon.05G0015860 | 1,300 | 0,0016 |
| Sspon.05G0015900 | 2,223 | 0,0016 |
| Sspon.05G0015940 | -1,591 | 0,0016 |
| Sspon.05G0015950 | -1,525 | 0,0016 |
| Sspon.05G0015970 | 1,551 | 0,0016 |
| Sspon.05G0016070 | 2,038 | 0,0016 |
| Sspon.05G0016090 | 1,005 | 0,0016 |
| Sspon.05G0016180 | 2,510 | 0,0016 |
| Sspon.05G0016270 | 3,308 | 0,0016 |
| Sspon.05G0016280 | 1,555 | 0,0016 |
| Sspon.05G0016290 | 1,578 | 0,0016 |
| Sspon.05G0016310 | 2,348 | 0,0016 |
| Sspon.05G0016350 | -2,986 | 0,0016 |
| Sspon.05G0016370 | 17,004 | 0,0016 |
| Sspon.05G0016450 | 2,942 | 0,0016 |
| Sspon.05G0016490 | 1,666 | 0,0016 |
| Sspon.05G0016550 | 1,491 | 0,0016 |
| Sspon.05G0016560 | 1,895 | 0,0016 |
| Sspon.05G0016650 | 1,250 | 0,0016 |
| Sspon.05G0016680 | -1,379 | 0,0016 |
| Sspon.05G0016710 | 1,610 | 0,0016 |
| Sspon.05G0016720 | 1,128 | 0,0016 |
| Sspon.05G0016750 | -3,965 | 0,0016 |
| Sspon.05G0016800 | 2,870 | 0,0016 |
| Sspon.05G0016820 | 1,014 | 0,0016 |
| Sspon.05G0016840 | 3,084 | 0,0016 |
| Sspon.05G0016860 | -1,383 | 0,0016 |
| Sspon.05G0016880 | 7,842 | 0,0016 |
| Sspon.05G0016890 | 18,845 | 0,0016 |
| Sspon.05G0017000 | -1,459 | 0,0016 |
| Sspon.05G0017010 | 1,169 | 0,0016 |
| Sspon.05G0017020 | 1,656 | 0,0016 |
| Sspon.05G0017060 | -1,206 | 0,0016 |
| Sspon.05G0017070 | 2,265 | 0,0017 |

| | | |
|---|---|---|
| Sspon.05G0017160 | -1,025 | 0,0017 |
| Sspon.05G0017220 | -2,545 | 0,0017 |
| Sspon.05G0017310 | 8,094 | 0,0017 |
| Sspon.05G0017320 | -1,066 | 0,0017 |
| Sspon.05G0017400 | 1,570 | 0,0017 |
| Sspon.05G0017480 | 5,599 | 0,0017 |
| Sspon.05G0017510 | 4,773 | 0,0017 |
| Sspon.05G0017590 | 1,400 | 0,0017 |
| Sspon.05G0017700 | 1,960 | 0,0017 |
| Sspon.05G0017770 | 15,334 | 0,0017 |
| Sspon.05G0017780 | 1,860 | 0,0017 |
| Sspon.05G0017810 | 1,521 | 0,0017 |
| Sspon.05G0017880 | 1,279 | 0,0017 |
| Sspon.05G0017890 | -1,878 | 0,0017 |
| Sspon.05G0017930 | -7,312 | 0,0017 |
| Sspon.05G0018000 | 8,178 | 0,0017 |
| Sspon.05G0018020 | -1,524 | 0,0017 |
| Sspon.05G0018040 | 17,085 | 0,0017 |
| Sspon.05G0018080 | 8,234 | 0,0017 |
| Sspon.05G0018090 | 7,354 | 0,0017 |
| Sspon.05G0018140 | -3,816 | 0,0017 |
| Sspon.05G0018160 | 2,349 | 0,0017 |
| Sspon.05G0018200 | 3,030 | 0,0017 |
| Sspon.05G0018220 | 1,494 | 0,0017 |
| Sspon.05G0018300 | -3,027 | 0,0017 |
| Sspon.05G0018310 | -1,275 | 0,0017 |
| Sspon.05G0018320 | 6,625 | 0,0017 |
| Sspon.05G0018430 | 1,411 | 0,0017 |
| Sspon.05G0018460 | 5,622 | 0,0017 |
| Sspon.05G0018500 | 1,929 | 0,0018 |
| Sspon.05G0018550 | -1,073 | 0,0018 |
| Sspon.05G0018590 | 1,340 | 0,0018 |
| Sspon.05G0018610 | -7,837 | 0,0018 |
| Sspon.05G0018680 | -1,237 | 0,0018 |
| Sspon.05G0018770 | 1,307 | 0,0018 |
| Sspon.05G0018790 | 1,101 | 0,0018 |
| Sspon.05G0018880 | 2,704 | 0,0018 |
| Sspon.05G0018890 | 1,167 | 0,0018 |
| Sspon.05G0018930 | -8,517 | 0,0018 |
| Sspon.05G0018970 | -2,137 | 0,0018 |
| Sspon.05G0019010 | -1,087 | 0,0018 |
| Sspon.05G0019040 | 6,422 | 0,0018 |
| Sspon.05G0019080 | 1,503 | 0,0018 |

| | | |
|---|---|---|
| Sspon.05G0019100 | 1,401 | 0,0018 |
| Sspon.05G0019260 | 16,994 | 0,0018 |
| Sspon.05G0019280 | 1,575 | 0,0018 |
| Sspon.05G0019300 | -1,107 | 0,0018 |
| Sspon.05G0019310 | -3,061 | 0,0018 |
| Sspon.05G0019360 | 3,404 | 0,0018 |
| Sspon.05G0019390 | 2,081 | 0,0018 |
| Sspon.05G0019420 | 1,402 | 0,0018 |
| Sspon.05G0019430 | -5,790 | 0,0018 |
| Sspon.05G0019450 | 1,045 | 0,0018 |
| Sspon.05G0019550 | 1,044 | 0,0018 |
| Sspon.05G0019600 | -1,063 | 0,0018 |
| Sspon.05G0019610 | 7,537 | 0,0018 |
| Sspon.05G0019660 | -1,232 | 0,0018 |
| Sspon.05G0019690 | 7,506 | 0,0018 |
| Sspon.05G0019730 | 1,143 | 0,0018 |
| Sspon.05G0019810 | -1,326 | 0,0018 |
| Sspon.05G0019840 | -2,823 | 0,0018 |
| Sspon.05G0019880 | 16,023 | 0,0018 |
| Sspon.05G0019890 | -6,331 | 0,0018 |
| Sspon.05G0019920 | 1,905 | 0,0018 |
| Sspon.05G0019930 | 2,804 | 0,0018 |
| Sspon.05G0019950 | 4,525 | 0,0018 |
| Sspon.05G0020100 | -6,177 | 0,0018 |
| Sspon.05G0020110 | 1,513 | 0,0018 |
| Sspon.05G0020120 | -1,003 | 0,0018 |
| Sspon.05G0020170 | 1,427 | 0,0019 |
| Sspon.05G0020190 | 6,480 | 0,0019 |
| Sspon.05G0020220 | 4,558 | 0,0019 |
| Sspon.05G0020270 | -3,036 | 0,0019 |
| Sspon.05G0020400 | 2,039 | 0,0019 |
| Sspon.05G0020520 | -2,281 | 0,0019 |
| Sspon.05G0020620 | -1,171 | 0,0019 |
| Sspon.05G0020650 | -1,917 | 0,0019 |
| Sspon.05G0020690 | 1,248 | 0,0019 |
| Sspon.05G0020710 | 1,571 | 0,0019 |
| Sspon.05G0020720 | 1,232 | 0,0019 |
| Sspon.05G0020760 | 1,569 | 0,0019 |
| Sspon.05G0020800 | 3,332 | 0,0019 |
| Sspon.05G0020840 | -1,228 | 0,0019 |
| Sspon.05G0020850 | -1,454 | 0,0019 |
| Sspon.05G0020860 | 1,104 | 0,0019 |
| Sspon.05G0020910 | 2,964 | 0,0019 |

| | | |
|---|---|---|
| Sspon.05G0020940 | 1,396 | 0,0019 |
| Sspon.05G0020950 | 1,337 | 0,0020 |
| Sspon.05G0021100 | 3,380 | 0,0020 |
| Sspon.05G0021110 | -1,249 | 0,0020 |
| Sspon.05G0021160 | 2,330 | 0,0020 |
| Sspon.05G0021170 | 2,326 | 0,0020 |
| Sspon.05G0021220 | -3,199 | 0,0020 |
| Sspon.05G0021330 | 1,058 | 0,0020 |
| Sspon.05G0021350 | 2,320 | 0,0020 |
| Sspon.05G0021360 | 1,572 | 0,0020 |
| Sspon.05G0021470 | -1,759 | 0,0020 |
| Sspon.05G0021480 | 5,058 | 0,0020 |
| Sspon.05G0021530 | 1,865 | 0,0020 |
| Sspon.05G0021540 | 5,837 | 0,0020 |
| Sspon.05G0021550 | -1,113 | 0,0020 |
| Sspon.05G0021600 | 1,062 | 0,0020 |
| Sspon.05G0021610 | -3,956 | 0,0020 |
| Sspon.05G0021750 | -1,187 | 0,0020 |
| Sspon.05G0021850 | -2,172 | 0,0020 |
| Sspon.05G0021900 | 1,758 | 0,0020 |
| Sspon.05G0021920 | 1,938 | 0,0020 |
| Sspon.05G0022090 | -2,404 | 0,0020 |
| Sspon.05G0022150 | 1,798 | 0,0021 |
| Sspon.05G0022200 | -2,258 | 0,0021 |
| Sspon.05G0022210 | 4,616 | 0,0021 |
| Sspon.05G0022470 | 3,256 | 0,0021 |
| Sspon.05G0022480 | 2,122 | 0,0021 |
| Sspon.05G0022490 | 1,985 | 0,0021 |
| Sspon.05G0022510 | -1,110 | 0,0021 |
| Sspon.05G0022640 | -3,161 | 0,0021 |
| Sspon.05G0022660 | 1,353 | 0,0021 |
| Sspon.05G0022700 | -1,496 | 0,0021 |
| Sspon.05G0022770 | 1,185 | 0,0021 |
| Sspon.05G0022850 | 1,365 | 0,0021 |
| Sspon.05G0022930 | 19,145 | 0,0021 |
| Sspon.05G0023070 | 1,183 | 0,0021 |
| Sspon.05G0023090 | -1,013 | 0,0021 |
| Sspon.05G0023100 | 1,278 | 0,0021 |
| Sspon.05G0023140 | 6,081 | 0,0021 |
| Sspon.05G0023170 | -2,741 | 0,0021 |
| Sspon.05G0023260 | 17,901 | 0,0021 |
| Sspon.05G0023300 | 1,747 | 0,0021 |
| Sspon.05G0023360 | 1,458 | 0,0021 |

| | | |
|---|---|---|
| Sspon.05G0023380 | -1,434 | 0,0021 |
| Sspon.05G0023430 | -1,291 | 0,0021 |
| Sspon.05G0023530 | -1,227 | 0,0021 |
| Sspon.05G0023540 | 1,649 | 0,0021 |
| Sspon.05G0023570 | 1,707 | 0,0022 |
| Sspon.05G0023650 | -2,120 | 0,0022 |
| Sspon.05G0023760 | -5,795 | 0,0022 |
| Sspon.05G0023790 | -3,996 | 0,0022 |
| Sspon.05G0023880 | 1,059 | 0,0022 |
| Sspon.05G0023910 | 16,572 | 0,0022 |
| Sspon.05G0023920 | 1,561 | 0,0022 |
| Sspon.05G0023950 | 8,757 | 0,0022 |
| Sspon.05G0024110 | -1,116 | 0,0022 |
| Sspon.05G0024210 | 1,138 | 0,0022 |
| Sspon.05G0024250 | 1,253 | 0,0022 |
| Sspon.05G0024360 | 2,541 | 0,0022 |
| Sspon.05G0024380 | 1,650 | 0,0022 |
| Sspon.05G0024420 | -1,637 | 0,0022 |
| Sspon.05G0024460 | 6,687 | 0,0022 |
| Sspon.05G0024500 | -3,998 | 0,0022 |
| Sspon.05G0024560 | -1,409 | 0,0022 |
| Sspon.05G0024570 | 1,036 | 0,0022 |
| Sspon.05G0024580 | 1,280 | 0,0022 |
| Sspon.05G0024610 | 1,047 | 0,0022 |
| Sspon.05G0024670 | -1,182 | 0,0022 |
| Sspon.05G0024680 | 1,017 | 0,0022 |
| Sspon.05G0024700 | -7,922 | 0,0022 |
| Sspon.05G0025030 | 1,166 | 0,0022 |
| Sspon.05G0025040 | 5,369 | 0,0022 |
| Sspon.05G0025080 | -2,975 | 0,0022 |
| Sspon.05G0025210 | -1,090 | 0,0023 |
| Sspon.05G0025360 | -1,019 | 0,0023 |
| Sspon.05G0025460 | 2,351 | 0,0023 |
| Sspon.05G0025480 | 1,588 | 0,0023 |
| Sspon.05G0025500 | -1,043 | 0,0023 |
| Sspon.05G0025510 | 7,603 | 0,0023 |
| Sspon.05G0025580 | 1,799 | 0,0023 |
| Sspon.05G0025590 | 4,108 | 0,0023 |
| Sspon.05G0025670 | 14,954 | 0,0023 |
| Sspon.05G0025720 | -1,112 | 0,0023 |
| Sspon.05G0025750 | 2,618 | 0,0023 |
| Sspon.05G0025880 | 5,959 | 0,0023 |
| Sspon.05G0025890 | 1,586 | 0,0023 |

| | | |
|---|---|---|
| Sspon.05G0025920 | 3,239 | 0,0023 |
| Sspon.05G0025950 | 1,826 | 0,0023 |
| Sspon.05G0026000 | 1,083 | 0,0023 |
| Sspon.05G0026060 | 1,822 | 0,0023 |
| Sspon.05G0026100 | 3,817 | 0,0023 |
| Sspon.05G0026150 | -3,217 | 0,0023 |
| Sspon.05G0026250 | 1,333 | 0,0023 |
| Sspon.05G0026260 | 1,210 | 0,0023 |
| Sspon.05G0026410 | -1,175 | 0,0023 |
| Sspon.05G0026430 | 5,223 | 0,0023 |
| Sspon.05G0026660 | 6,442 | 0,0023 |
| Sspon.05G0026710 | -1,197 | 0,0023 |
| Sspon.05G0026790 | 1,319 | 0,0023 |
| Sspon.05G0026800 | 1,370 | 0,0023 |
| Sspon.05G0026910 | 1,441 | 0,0023 |
| Sspon.05G0026990 | -2,930 | 0,0023 |
| Sspon.05G0027050 | 2,578 | 0,0024 |
| Sspon.05G0027080 | -2,835 | 0,0024 |
| Sspon.05G0027310 | -1,126 | 0,0024 |
| Sspon.05G0027420 | 1,422 | 0,0024 |
| Sspon.05G0027440 | -2,439 | 0,0024 |
| Sspon.05G0027510 | 1,335 | 0,0024 |
| Sspon.05G0027620 | 1,063 | 0,0024 |
| Sspon.05G0027750 | 5,277 | 0,0024 |
| Sspon.05G0027770 | 1,184 | 0,0024 |
| Sspon.05G0027790 | 3,545 | 0,0024 |
| Sspon.05G0027940 | 1,306 | 0,0024 |
| Sspon.05G0027950 | 2,349 | 0,0024 |
| Sspon.05G0028010 | 14,463 | 0,0024 |
| Sspon.05G0028060 | 1,960 | 0,0024 |
| Sspon.05G0028090 | 1,936 | 0,0024 |
| Sspon.05G0028210 | 1,201 | 0,0024 |
| Sspon.05G0028230 | 1,206 | 0,0024 |
| Sspon.05G0028240 | -8,355 | 0,0024 |
| Sspon.05G0028320 | -7,007 | 0,0024 |
| Sspon.05G0028420 | 3,850 | 0,0024 |
| Sspon.05G0028520 | 1,724 | 0,0024 |
| Sspon.05G0028560 | 1,545 | 0,0024 |
| Sspon.05G0028570 | 1,705 | 0,0024 |
| Sspon.05G0028580 | 1,162 | 0,0024 |
| Sspon.05G0028630 | 18,768 | 0,0024 |
| Sspon.05G0028710 | -3,904 | 0,0024 |
| Sspon.05G0028730 | -1,729 | 0,0024 |

| | | |
|---|---|---|
| Sspon.05G0028740 | 6,343 | 0,0024 |
| Sspon.05G0028830 | 7,640 | 0,0024 |
| Sspon.05G0028870 | -1,011 | 0,0024 |
| Sspon.05G0028920 | -2,538 | 0,0025 |
| Sspon.05G0028950 | 1,488 | 0,0025 |
| Sspon.05G0028960 | 1,004 | 0,0025 |
| Sspon.05G0028980 | 5,567 | 0,0025 |
| Sspon.05G0029030 | 1,870 | 0,0025 |
| Sspon.05G0029040 | 1,835 | 0,0025 |
| Sspon.05G0029050 | 15,192 | 0,0025 |
| Sspon.05G0029070 | -1,756 | 0,0025 |
| Sspon.05G0029080 | -1,312 | 0,0025 |
| Sspon.05G0029130 | -6,258 | 0,0025 |
| Sspon.05G0029180 | -2,872 | 0,0025 |
| Sspon.05G0029200 | 1,814 | 0,0025 |
| Sspon.05G0029230 | 6,073 | 0,0025 |
| Sspon.05G0029250 | -1,239 | 0,0025 |
| Sspon.05G0029300 | 1,543 | 0,0025 |
| Sspon.05G0029310 | 3,520 | 0,0025 |
| Sspon.05G0029340 | 1,302 | 0,0025 |
| Sspon.05G0029360 | -1,179 | 0,0025 |
| Sspon.05G0029480 | 1,463 | 0,0025 |
| Sspon.05G0029510 | 1,212 | 0,0025 |
| Sspon.05G0029520 | 3,674 | 0,0025 |
| Sspon.05G0029560 | 7,909 | 0,0025 |
| Sspon.05G0029640 | 1,757 | 0,0025 |
| Sspon.05G0029650 | 8,289 | 0,0025 |
| Sspon.05G0029700 | -1,216 | 0,0025 |
| Sspon.05G0029760 | 4,586 | 0,0025 |
| Sspon.05G0029800 | 1,920 | 0,0025 |
| Sspon.05G0029810 | -1,049 | 0,0026 |
| Sspon.05G0029880 | -1,678 | 0,0026 |
| Sspon.05G0030020 | -5,698 | 0,0026 |
| Sspon.05G0030030 | -1,142 | 0,0026 |
| Sspon.05G0030090 | 1,705 | 0,0026 |
| Sspon.05G0030200 | -6,998 | 0,0026 |
| Sspon.05G0030250 | 15,367 | 0,0026 |
| Sspon.05G0030300 | 2,524 | 0,0026 |
| Sspon.05G0030340 | 3,442 | 0,0026 |
| Sspon.05G0030370 | 5,285 | 0,0026 |
| Sspon.05G0030390 | -1,539 | 0,0026 |
| Sspon.05G0030460 | 1,000 | 0,0026 |
| Sspon.05G0030480 | 2,280 | 0,0026 |

| | | |
|---|---|---|
| Sspon.05G0030500 | -1,304 | 0,0026 |
| Sspon.05G0030560 | 3,394 | 0,0026 |
| Sspon.05G0030570 | 1,186 | 0,0026 |
| Sspon.05G0030780 | 1,847 | 0,0026 |
| Sspon.05G0030810 | 1,087 | 0,0026 |
| Sspon.05G0030820 | 3,511 | 0,0026 |
| Sspon.05G0030870 | 15,624 | 0,0026 |
| Sspon.05G0030880 | 4,108 | 0,0026 |
| Sspon.05G0030950 | 1,575 | 0,0026 |
| Sspon.05G0030960 | -6,866 | 0,0026 |
| Sspon.05G0030970 | -1,085 | 0,0026 |
| Sspon.05G0031000 | 1,149 | 0,0026 |
| Sspon.05G0031050 | 1,227 | 0,0027 |
| Sspon.05G0031060 | 1,431 | 0,0027 |
| Sspon.05G0031140 | 6,699 | 0,0027 |
| Sspon.05G0031220 | 1,979 | 0,0027 |
| Sspon.05G0031240 | 1,793 | 0,0027 |
| Sspon.05G0031270 | 1,170 | 0,0027 |
| Sspon.05G0031280 | 14,892 | 0,0027 |
| Sspon.05G0031320 | 3,788 | 0,0027 |
| Sspon.05G0031330 | 1,815 | 0,0027 |
| Sspon.05G0031350 | 1,036 | 0,0027 |
| Sspon.05G0031370 | -3,000 | 0,0027 |
| Sspon.05G0031520 | -6,224 | 0,0027 |
| Sspon.05G0031530 | 2,668 | 0,0027 |
| Sspon.05G0031640 | 5,392 | 0,0027 |
| Sspon.05G0031670 | 1,089 | 0,0027 |
| Sspon.05G0031780 | 7,226 | 0,0027 |
| Sspon.05G0031910 | 1,223 | 0,0027 |
| Sspon.05G0031940 | 1,742 | 0,0027 |
| Sspon.05G0031960 | 1,499 | 0,0027 |
| Sspon.05G0032000 | 1,606 | 0,0027 |
| Sspon.05G0032060 | -2,098 | 0,0028 |
| Sspon.05G0032080 | -1,292 | 0,0028 |
| Sspon.05G0032110 | -1,006 | 0,0028 |
| Sspon.05G0032190 | -1,905 | 0,0028 |
| Sspon.05G0032230 | 1,600 | 0,0028 |
| Sspon.05G0032340 | 3,347 | 0,0028 |
| Sspon.05G0032390 | 6,956 | 0,0028 |
| Sspon.05G0032430 | 2,423 | 0,0028 |
| Sspon.05G0032510 | -2,105 | 0,0028 |
| Sspon.05G0032520 | 3,456 | 0,0028 |
| Sspon.05G0032590 | 1,019 | 0,0028 |

| | | |
|---|---|---|
| Sspon.05G0032600 | 1,788 | 0,0028 |
| Sspon.05G0032630 | 3,920 | 0,0028 |
| Sspon.05G0032660 | 1,437 | 0,0028 |
| Sspon.05G0032730 | -4,858 | 0,0028 |
| Sspon.05G0032820 | 4,275 | 0,0028 |
| Sspon.05G0032850 | 4,360 | 0,0028 |
| Sspon.05G0032860 | 1,518 | 0,0028 |
| Sspon.05G0033050 | 3,715 | 0,0028 |
| Sspon.05G0033120 | -4,335 | 0,0028 |
| Sspon.05G0033190 | 3,562 | 0,0028 |
| Sspon.05G0033240 | 1,956 | 0,0028 |
| Sspon.05G0033300 | 1,698 | 0,0028 |
| Sspon.05G0033310 | -2,044 | 0,0028 |
| Sspon.05G0033340 | 4,781 | 0,0028 |
| Sspon.05G0033350 | 7,548 | 0,0028 |
| Sspon.05G0033360 | 1,916 | 0,0028 |
| Sspon.05G0033480 | 2,188 | 0,0028 |
| Sspon.05G0033590 | 1,897 | 0,0029 |
| Sspon.05G0033680 | -1,267 | 0,0029 |
| Sspon.05G0033690 | -1,109 | 0,0029 |
| Sspon.05G0033720 | -2,428 | 0,0029 |
| Sspon.05G0033750 | -1,238 | 0,0029 |
| Sspon.05G0033770 | -2,196 | 0,0029 |
| Sspon.05G0033910 | 15,811 | 0,0029 |
| Sspon.05G0033980 | 2,260 | 0,0029 |
| Sspon.05G0034090 | 1,296 | 0,0029 |
| Sspon.05G0034100 | 1,441 | 0,0029 |
| Sspon.05G0034110 | -2,335 | 0,0029 |
| Sspon.05G0034130 | -6,277 | 0,0029 |
| Sspon.05G0034160 | 15,017 | 0,0029 |
| Sspon.05G0034180 | 2,169 | 0,0029 |
| Sspon.05G0034310 | 1,483 | 0,0029 |
| Sspon.05G0034370 | 2,868 | 0,0029 |
| Sspon.05G0034510 | 2,359 | 0,0029 |
| Sspon.05G0034520 | 1,482 | 0,0029 |
| Sspon.05G0034560 | -1,172 | 0,0029 |
| Sspon.05G0034580 | 6,079 | 0,0029 |
| Sspon.05G0034590 | 17,914 | 0,0029 |
| Sspon.05G0034610 | 1,009 | 0,0029 |
| Sspon.05G0034640 | -1,487 | 0,0030 |
| Sspon.05G0034650 | -2,515 | 0,0030 |
| Sspon.05G0034670 | 1,465 | 0,0030 |
| Sspon.05G0034680 | -1,441 | 0,0030 |

| | | |
|---|---|---|
| Sspon.05G0034740 | -1,106 | 0,0030 |
| Sspon.05G0034790 | -2,088 | 0,0030 |
| Sspon.05G0034800 | -1,162 | 0,0030 |
| Sspon.05G0035010 | -1,086 | 0,0030 |
| Sspon.05G0035030 | 15,334 | 0,0030 |
| Sspon.05G0035060 | -1,044 | 0,0030 |
| Sspon.05G0035110 | 1,667 | 0,0030 |
| Sspon.05G0035160 | -2,226 | 0,0030 |
| Sspon.05G0035170 | 17,278 | 0,0030 |
| Sspon.05G0035180 | 1,078 | 0,0030 |
| Sspon.05G0035230 | 1,125 | 0,0030 |
| Sspon.05G0035290 | -2,560 | 0,0030 |
| Sspon.05G0035300 | 1,727 | 0,0030 |
| Sspon.05G0035330 | 7,043 | 0,0030 |
| Sspon.05G0035370 | -1,428 | 0,0030 |
| Sspon.05G0035380 | 1,134 | 0,0030 |
| Sspon.05G0035400 | 1,271 | 0,0030 |
| Sspon.05G0035460 | -2,717 | 0,0030 |
| Sspon.05G0035500 | -1,370 | 0,0030 |
| Sspon.05G0035530 | -1,022 | 0,0030 |
| Sspon.05G0035590 | 2,062 | 0,0031 |
| Sspon.05G0035610 | 2,829 | 0,0031 |
| Sspon.05G0035630 | 1,423 | 0,0031 |
| Sspon.05G0035710 | 1,569 | 0,0031 |
| Sspon.05G0035740 | 2,268 | 0,0031 |
| Sspon.05G0035780 | -1,642 | 0,0031 |
| Sspon.05G0035790 | 1,337 | 0,0031 |
| Sspon.05G0035800 | 1,522 | 0,0031 |
| Sspon.05G0035870 | -2,075 | 0,0031 |
| Sspon.05G0035890 | 5,171 | 0,0031 |
| Sspon.05G0036030 | 1,180 | 0,0031 |
| Sspon.05G0036040 | 3,307 | 0,0031 |
| Sspon.05G0036080 | 2,417 | 0,0031 |
| Sspon.05G0036090 | 1,236 | 0,0031 |
| Sspon.05G0036100 | 6,022 | 0,0031 |
| Sspon.05G0036170 | -2,429 | 0,0031 |
| Sspon.05G0036260 | 3,909 | 0,0031 |
| Sspon.05G0036280 | -8,079 | 0,0031 |
| Sspon.05G0036320 | -1,213 | 0,0031 |
| Sspon.05G0036330 | -4,340 | 0,0031 |
| Sspon.05G0036470 | 2,003 | 0,0031 |
| Sspon.05G0036550 | -2,297 | 0,0032 |
| Sspon.05G0036580 | 1,127 | 0,0032 |

| | | |
|---|---|---|
| Sspon.05G0036680 | 1,628 | 0,0032 |
| Sspon.05G0036740 | -7,675 | 0,0032 |
| Sspon.05G0036760 | -1,030 | 0,0032 |
| Sspon.05G0036770 | 1,043 | 0,0032 |
| Sspon.05G0036800 | -3,791 | 0,0032 |
| Sspon.05G0036810 | -2,085 | 0,0032 |
| Sspon.05G0036860 | 15,661 | 0,0032 |
| Sspon.05G0036880 | 6,124 | 0,0032 |
| Sspon.05G0036960 | 4,454 | 0,0032 |
| Sspon.05G0037020 | 4,707 | 0,0032 |
| Sspon.05G0037180 | 17,747 | 0,0032 |
| Sspon.05G0037200 | -1,816 | 0,0032 |
| Sspon.05G0037250 | 2,040 | 0,0032 |
| Sspon.05G0037260 | -1,151 | 0,0032 |
| Sspon.05G0037290 | 1,010 | 0,0032 |
| Sspon.05G0037300 | 1,027 | 0,0032 |
| Sspon.05G0037320 | 1,118 | 0,0032 |
| Sspon.05G0037430 | -1,898 | 0,0032 |
| Sspon.05G0037470 | 5,248 | 0,0033 |
| Sspon.05G0037540 | 1,935 | 0,0033 |
| Sspon.05G0037630 | -4,649 | 0,0033 |
| Sspon.05G0037640 | 2,471 | 0,0033 |
| Sspon.05G0037670 | 1,214 | 0,0033 |
| Sspon.05G0037680 | 2,531 | 0,0033 |
| Sspon.05G0037760 | 1,095 | 0,0033 |
| Sspon.05G0037870 | -1,002 | 0,0033 |
| Sspon.05G0038370 | -4,635 | 0,0033 |
| Sspon.05G0038420 | 6,149 | 0,0033 |
| Sspon.05G0038550 | 3,290 | 0,0033 |
| Sspon.05G0038660 | -1,164 | 0,0033 |
| Sspon.05G0038730 | 3,675 | 0,0033 |
| Sspon.05G0038780 | 1,691 | 0,0033 |
| Sspon.05G0038800 | -1,018 | 0,0033 |
| Sspon.05G0038830 | -2,306 | 0,0033 |
| Sspon.05G0038860 | 5,305 | 0,0033 |
| Sspon.05G0038960 | 1,003 | 0,0033 |
| Sspon.05G0039060 | -1,420 | 0,0033 |
| Sspon.05G0039140 | 1,615 | 0,0033 |
| Sspon.05G0039280 | 1,356 | 0,0033 |
| Sspon.05G0039500 | 14,517 | 0,0033 |
| Sspon.05G0039550 | 1,762 | 0,0034 |
| Sspon.05G0039570 | 1,254 | 0,0034 |
| Sspon.05G0039620 | -6,050 | 0,0034 |

| | | |
|---|---|---|
| Sspon.05G0039630 | -2,343 | 0,0034 |
| Sspon.05G0039720 | 1,104 | 0,0034 |
| Sspon.05G0039730 | -7,522 | 0,0034 |
| Sspon.05G0039870 | 1,473 | 0,0034 |
| Sspon.05G0039880 | 2,017 | 0,0034 |
| Sspon.05G0039900 | 1,106 | 0,0034 |
| Sspon.05G0039920 | -2,453 | 0,0034 |
| Sspon.05G0039980 | -5,023 | 0,0034 |
| Sspon.05G0040000 | -1,453 | 0,0034 |
| Sspon.06G0000160 | -1,079 | 0,0034 |
| Sspon.06G0000290 | -7,607 | 0,0034 |
| Sspon.06G0000320 | 1,860 | 0,0035 |
| Sspon.06G0000400 | 8,507 | 0,0035 |
| Sspon.06G0000410 | -1,196 | 0,0035 |
| Sspon.06G0000450 | 2,672 | 0,0035 |
| Sspon.06G0000540 | -1,442 | 0,0035 |
| Sspon.06G0000570 | 1,524 | 0,0035 |
| Sspon.06G0000580 | 3,522 | 0,0035 |
| Sspon.06G0000590 | 1,485 | 0,0035 |
| Sspon.06G0000620 | 5,454 | 0,0035 |
| Sspon.06G0000660 | -2,420 | 0,0035 |
| Sspon.06G0000670 | 1,463 | 0,0035 |
| Sspon.06G0000680 | -7,633 | 0,0036 |
| Sspon.06G0000730 | 2,042 | 0,0036 |
| Sspon.06G0000790 | -1,023 | 0,0036 |
| Sspon.06G0000900 | -1,128 | 0,0036 |
| Sspon.06G0000910 | 1,207 | 0,0036 |
| Sspon.06G0000920 | -4,890 | 0,0036 |
| Sspon.06G0000950 | 2,108 | 0,0036 |
| Sspon.06G0000970 | -3,063 | 0,0036 |
| Sspon.06G0000980 | 1,254 | 0,0036 |
| Sspon.06G0001120 | 2,617 | 0,0036 |
| Sspon.06G0001160 | 1,319 | 0,0036 |
| Sspon.06G0001220 | 3,932 | 0,0036 |
| Sspon.06G0001290 | 1,052 | 0,0037 |
| Sspon.06G0001340 | 1,111 | 0,0037 |
| Sspon.06G0001470 | -1,201 | 0,0037 |
| Sspon.06G0001530 | -4,699 | 0,0037 |
| Sspon.06G0001540 | 2,178 | 0,0037 |
| Sspon.06G0001610 | 1,964 | 0,0037 |
| Sspon.06G0001620 | 1,120 | 0,0037 |
| Sspon.06G0001740 | 1,040 | 0,0037 |
| Sspon.06G0001770 | 1,168 | 0,0037 |

| | | |
|---|---|---|
| Sspon.06G0001800 | -3,023 | 0,0037 |
| Sspon.06G0001810 | 2,404 | 0,0037 |
| Sspon.06G0001820 | 7,642 | 0,0037 |
| Sspon.06G0001860 | 4,674 | 0,0037 |
| Sspon.06G0001930 | 1,141 | 0,0037 |
| Sspon.06G0001950 | 1,494 | 0,0037 |
| Sspon.06G0002040 | -1,017 | 0,0038 |
| Sspon.06G0002140 | 1,361 | 0,0038 |
| Sspon.06G0002150 | 7,541 | 0,0038 |
| Sspon.06G0002170 | -1,171 | 0,0038 |
| Sspon.06G0002250 | 1,011 | 0,0038 |
| Sspon.06G0002260 | -4,604 | 0,0038 |
| Sspon.06G0002270 | 5,184 | 0,0038 |
| Sspon.06G0002340 | 5,337 | 0,0038 |
| Sspon.06G0002390 | -2,296 | 0,0038 |
| Sspon.06G0002600 | 4,434 | 0,0038 |
| Sspon.06G0002830 | -1,110 | 0,0038 |
| Sspon.06G0002850 | -1,300 | 0,0039 |
| Sspon.06G0002870 | 4,367 | 0,0039 |
| Sspon.06G0002890 | -1,007 | 0,0039 |
| Sspon.06G0002930 | 3,154 | 0,0039 |
| Sspon.06G0003060 | 6,644 | 0,0039 |
| Sspon.06G0003110 | 15,754 | 0,0039 |
| Sspon.06G0003170 | 1,082 | 0,0039 |
| Sspon.06G0003180 | -1,203 | 0,0039 |
| Sspon.06G0003300 | 3,707 | 0,0039 |
| Sspon.06G0003360 | 2,422 | 0,0039 |
| Sspon.06G0003390 | 2,400 | 0,0039 |
| Sspon.06G0003430 | -2,406 | 0,0039 |
| Sspon.06G0003530 | -7,001 | 0,0039 |
| Sspon.06G0003650 | -2,407 | 0,0040 |
| Sspon.06G0003660 | 6,160 | 0,0040 |
| Sspon.06G0003670 | -5,055 | 0,0040 |
| Sspon.06G0003680 | 2,280 | 0,0040 |
| Sspon.06G0003720 | 6,519 | 0,0040 |
| Sspon.06G0003770 | 1,038 | 0,0040 |
| Sspon.06G0003820 | -6,978 | 0,0040 |
| Sspon.06G0003830 | -4,422 | 0,0040 |
| Sspon.06G0003840 | 1,179 | 0,0041 |
| Sspon.06G0003960 | 2,361 | 0,0041 |
| Sspon.06G0004020 | -2,449 | 0,0041 |
| Sspon.06G0004040 | 3,500 | 0,0041 |
| Sspon.06G0004070 | 1,916 | 0,0041 |

| | | |
|---|---|---|
| Sspon.06G0004100 | -4,833 | 0,0041 |
| Sspon.06G0004120 | 1,413 | 0,0041 |
| Sspon.06G0004210 | -4,595 | 0,0041 |
| Sspon.06G0004230 | 2,152 | 0,0041 |
| Sspon.06G0004260 | -1,547 | 0,0041 |
| Sspon.06G0004300 | -1,787 | 0,0041 |
| Sspon.06G0004390 | -2,262 | 0,0041 |
| Sspon.06G0004430 | -1,082 | 0,0041 |
| Sspon.06G0004440 | 15,441 | 0,0041 |
| Sspon.06G0004480 | -2,161 | 0,0041 |
| Sspon.06G0004510 | -2,401 | 0,0041 |
| Sspon.06G0004590 | -5,213 | 0,0042 |
| Sspon.06G0004720 | -1,129 | 0,0042 |
| Sspon.06G0004790 | -1,484 | 0,0042 |
| Sspon.06G0004860 | 1,937 | 0,0042 |
| Sspon.06G0004880 | 1,015 | 0,0042 |
| Sspon.06G0004900 | 1,306 | 0,0042 |
| Sspon.06G0004980 | 1,567 | 0,0042 |
| Sspon.06G0005130 | -1,068 | 0,0042 |
| Sspon.06G0005320 | 1,066 | 0,0042 |
| Sspon.06G0005360 | -1,029 | 0,0042 |
| Sspon.06G0005440 | -1,183 | 0,0042 |
| Sspon.06G0005480 | -3,465 | 0,0042 |
| Sspon.06G0005500 | -2,003 | 0,0042 |
| Sspon.06G0005520 | 1,869 | 0,0043 |
| Sspon.06G0005570 | 1,607 | 0,0043 |
| Sspon.06G0005620 | 1,966 | 0,0043 |
| Sspon.06G0005680 | 1,930 | 0,0043 |
| Sspon.06G0005780 | -1,595 | 0,0043 |
| Sspon.06G0005880 | 1,830 | 0,0043 |
| Sspon.06G0006200 | 1,314 | 0,0043 |
| Sspon.06G0006210 | -7,119 | 0,0043 |
| Sspon.06G0006340 | 2,198 | 0,0043 |
| Sspon.06G0006430 | 2,089 | 0,0043 |
| Sspon.06G0006440 | -1,258 | 0,0043 |
| Sspon.06G0006450 | 3,157 | 0,0043 |
| Sspon.06G0006490 | 3,296 | 0,0044 |
| Sspon.06G0006510 | 2,782 | 0,0044 |
| Sspon.06G0006560 | 1,613 | 0,0044 |
| Sspon.06G0006660 | 2,246 | 0,0044 |
| Sspon.06G0006680 | 6,606 | 0,0044 |
| Sspon.06G0006700 | 6,954 | 0,0044 |
| Sspon.06G0006750 | 4,110 | 0,0044 |

| | | |
|---|---|---|
| Sspon.06G0006780 | -2,102 | 0,0044 |
| Sspon.06G0006880 | 1,441 | 0,0044 |
| Sspon.06G0006960 | 1,930 | 0,0044 |
| Sspon.06G0006990 | 1,888 | 0,0044 |
| Sspon.06G0007080 | 3,462 | 0,0044 |
| Sspon.06G0007160 | 5,008 | 0,0044 |
| Sspon.06G0007210 | 2,865 | 0,0044 |
| Sspon.06G0007250 | 3,388 | 0,0044 |
| Sspon.06G0007310 | 1,305 | 0,0045 |
| Sspon.06G0007430 | 5,814 | 0,0045 |
| Sspon.06G0007450 | -1,109 | 0,0045 |
| Sspon.06G0007470 | -1,197 | 0,0045 |
| Sspon.06G0007630 | 1,235 | 0,0045 |
| Sspon.06G0007710 | -1,508 | 0,0045 |
| Sspon.06G0007750 | 1,009 | 0,0045 |
| Sspon.06G0007770 | 1,808 | 0,0045 |
| Sspon.06G0007830 | 6,490 | 0,0045 |
| Sspon.06G0007840 | -5,343 | 0,0045 |
| Sspon.06G0007870 | -2,944 | 0,0046 |
| Sspon.06G0007880 | 2,069 | 0,0046 |
| Sspon.06G0007890 | 6,693 | 0,0046 |
| Sspon.06G0007920 | 6,569 | 0,0046 |
| Sspon.06G0007940 | -1,933 | 0,0046 |
| Sspon.06G0008070 | 3,209 | 0,0046 |
| Sspon.06G0008140 | -7,851 | 0,0046 |
| Sspon.06G0008150 | -1,093 | 0,0046 |
| Sspon.06G0008200 | -3,539 | 0,0046 |
| Sspon.06G0008250 | -7,418 | 0,0047 |
| Sspon.06G0008370 | -3,286 | 0,0047 |
| Sspon.06G0008410 | -2,151 | 0,0047 |
| Sspon.06G0008440 | 1,003 | 0,0047 |
| Sspon.06G0008560 | 3,069 | 0,0047 |
| Sspon.06G0008570 | 1,016 | 0,0047 |
| Sspon.06G0008670 | 7,508 | 0,0047 |
| Sspon.06G0008730 | 1,936 | 0,0047 |
| Sspon.06G0008750 | 1,295 | 0,0047 |
| Sspon.06G0008910 | 1,397 | 0,0047 |
| Sspon.06G0008940 | -1,504 | 0,0048 |
| Sspon.06G0009100 | 1,228 | 0,0048 |
| Sspon.06G0009130 | -7,658 | 0,0048 |
| Sspon.06G0009140 | -1,106 | 0,0048 |
| Sspon.06G0009150 | -2,930 | 0,0048 |
| Sspon.06G0009180 | 1,600 | 0,0048 |

| | | |
|---|---|---|
| Sspon.06G0009240 | 1,918 | 0,0048 |
| Sspon.06G0009340 | -7,409 | 0,0048 |
| Sspon.06G0009370 | -1,450 | 0,0048 |
| Sspon.06G0009420 | 1,144 | 0,0048 |
| Sspon.06G0009440 | 1,050 | 0,0049 |
| Sspon.06G0009490 | 1,912 | 0,0049 |
| Sspon.06G0009500 | 1,797 | 0,0049 |
| Sspon.06G0009510 | -7,579 | 0,0049 |
| Sspon.06G0009530 | 1,560 | 0,0049 |
| Sspon.06G0009540 | -4,141 | 0,0049 |
| Sspon.06G0009550 | 1,780 | 0,0049 |
| Sspon.06G0009600 | 18,332 | 0,0049 |
| Sspon.06G0009620 | 1,357 | 0,0049 |
| Sspon.06G0009630 | -6,241 | 0,0049 |
| Sspon.06G0009650 | -2,799 | 0,0049 |
| Sspon.06G0009780 | 1,312 | 0,0049 |
| Sspon.06G0009800 | -1,258 | 0,0049 |
| Sspon.06G0009850 | 4,869 | 0,0049 |
| Sspon.06G0009860 | 1,151 | 0,0049 |
| Sspon.06G0009910 | 6,154 | 0,0050 |
| Sspon.06G0009970 | 2,499 | 0,0050 |
| Sspon.06G0010070 | 1,916 | 0,0050 |
| Sspon.06G0010120 | 3,058 | 0,0050 |
| Sspon.06G0010130 | -1,075 | 0,0050 |
| Sspon.06G0010160 | 1,961 | 0,0050 |
| Sspon.06G0010240 | -1,197 | 0,0050 |
| Sspon.06G0010300 | -3,713 | 0,0050 |
| Sspon.06G0010320 | 4,257 | 0,0050 |
| Sspon.06G0010370 | -3,403 | 0,0051 |
| Sspon.06G0010410 | 2,062 | 0,0051 |
| Sspon.06G0010490 | 5,711 | 0,0051 |
| Sspon.06G0010510 | 2,893 | 0,0051 |
| Sspon.06G0010560 | 1,612 | 0,0051 |
| Sspon.06G0010570 | 1,109 | 0,0051 |
| Sspon.06G0010620 | -1,642 | 0,0051 |
| Sspon.06G0010920 | 1,180 | 0,0052 |
| Sspon.06G0010970 | 7,503 | 0,0052 |
| Sspon.06G0010980 | -7,339 | 0,0052 |
| Sspon.06G0010990 | 2,699 | 0,0052 |
| Sspon.06G0011000 | 1,793 | 0,0052 |
| Sspon.06G0011030 | -1,045 | 0,0052 |
| Sspon.06G0011220 | -1,236 | 0,0052 |
| Sspon.06G0011400 | -1,802 | 0,0052 |

| | | |
|---|---|---|
| Sspon.06G0011460 | 5,695 | 0,0052 |
| Sspon.06G0011510 | 2,345 | 0,0052 |
| Sspon.06G0011520 | -1,175 | 0,0052 |
| Sspon.06G0011630 | -1,585 | 0,0053 |
| Sspon.06G0011640 | 1,273 | 0,0053 |
| Sspon.06G0011700 | -1,138 | 0,0053 |
| Sspon.06G0011760 | 14,894 | 0,0053 |
| Sspon.06G0011810 | 1,133 | 0,0053 |
| Sspon.06G0011850 | -1,221 | 0,0053 |
| Sspon.06G0011860 | 2,019 | 0,0053 |
| Sspon.06G0011910 | 2,981 | 0,0054 |
| Sspon.06G0011990 | 1,064 | 0,0054 |
| Sspon.06G0012140 | 6,717 | 0,0054 |
| Sspon.06G0012180 | 15,573 | 0,0054 |
| Sspon.06G0012270 | 16,326 | 0,0054 |
| Sspon.06G0012290 | -5,445 | 0,0054 |
| Sspon.06G0012590 | -2,328 | 0,0054 |
| Sspon.06G0012660 | 1,020 | 0,0054 |
| Sspon.06G0012670 | 15,020 | 0,0054 |
| Sspon.06G0012680 | 1,828 | 0,0054 |
| Sspon.06G0012710 | -1,016 | 0,0054 |
| Sspon.06G0012810 | 1,290 | 0,0055 |
| Sspon.06G0012820 | -6,722 | 0,0055 |
| Sspon.06G0012870 | 18,506 | 0,0055 |
| Sspon.06G0012900 | 1,317 | 0,0055 |
| Sspon.06G0012930 | -2,214 | 0,0055 |
| Sspon.06G0012940 | 2,856 | 0,0055 |
| Sspon.06G0012950 | -1,351 | 0,0055 |
| Sspon.06G0012980 | -1,189 | 0,0055 |
| Sspon.06G0013030 | -3,290 | 0,0055 |
| Sspon.06G0013040 | 1,701 | 0,0055 |
| Sspon.06G0013100 | 7,169 | 0,0055 |
| Sspon.06G0013110 | 4,737 | 0,0056 |
| Sspon.06G0013120 | -2,174 | 0,0056 |
| Sspon.06G0013150 | -5,362 | 0,0056 |
| Sspon.06G0013190 | 1,040 | 0,0056 |
| Sspon.06G0013230 | -5,521 | 0,0056 |
| Sspon.06G0013280 | -1,240 | 0,0056 |
| Sspon.06G0013290 | 1,352 | 0,0056 |
| Sspon.06G0013350 | 4,988 | 0,0056 |
| Sspon.06G0013370 | -4,231 | 0,0056 |
| Sspon.06G0013380 | 8,387 | 0,0056 |
| Sspon.06G0013390 | -1,201 | 0,0056 |

| | | |
|---|---|---|
| Sspon.06G0013410 | 1,370 | 0,0056 |
| Sspon.06G0013530 | -4,759 | 0,0056 |
| Sspon.06G0013560 | 1,026 | 0,0056 |
| Sspon.06G0013620 | -1,033 | 0,0056 |
| Sspon.06G0013670 | 1,663 | 0,0057 |
| Sspon.06G0013680 | -2,150 | 0,0057 |
| Sspon.06G0013830 | -1,291 | 0,0057 |
| Sspon.06G0013840 | -2,530 | 0,0057 |
| Sspon.06G0013860 | 5,606 | 0,0057 |
| Sspon.06G0013940 | 1,245 | 0,0057 |
| Sspon.06G0013970 | 1,390 | 0,0057 |
| Sspon.06G0014030 | 3,502 | 0,0057 |
| Sspon.06G0014160 | -1,126 | 0,0057 |
| Sspon.06G0014190 | -1,030 | 0,0057 |
| Sspon.06G0014210 | 1,374 | 0,0057 |
| Sspon.06G0014270 | -7,394 | 0,0057 |
| Sspon.06G0014340 | -2,477 | 0,0057 |
| Sspon.06G0014360 | 1,676 | 0,0057 |
| Sspon.06G0014400 | 2,812 | 0,0057 |
| Sspon.06G0014420 | -2,154 | 0,0057 |
| Sspon.06G0014450 | 4,696 | 0,0058 |
| Sspon.06G0014460 | 15,176 | 0,0058 |
| Sspon.06G0014550 | -2,965 | 0,0058 |
| Sspon.06G0014570 | -1,063 | 0,0058 |
| Sspon.06G0014650 | 1,958 | 0,0058 |
| Sspon.06G0014700 | 1,101 | 0,0058 |
| Sspon.06G0014830 | -6,516 | 0,0058 |
| Sspon.06G0014870 | -2,816 | 0,0058 |
| Sspon.06G0014880 | 1,060 | 0,0058 |
| Sspon.06G0014930 | -1,154 | 0,0058 |
| Sspon.06G0014960 | 5,503 | 0,0058 |
| Sspon.06G0014970 | 1,082 | 0,0058 |
| Sspon.06G0015120 | 2,296 | 0,0059 |
| Sspon.06G0015160 | -6,426 | 0,0059 |
| Sspon.06G0015180 | -1,395 | 0,0059 |
| Sspon.06G0015220 | 1,079 | 0,0059 |
| Sspon.06G0015230 | -3,462 | 0,0059 |
| Sspon.06G0015240 | -1,926 | 0,0059 |
| Sspon.06G0015310 | 2,899 | 0,0059 |
| Sspon.06G0015330 | -4,115 | 0,0059 |
| Sspon.06G0015480 | 5,332 | 0,0059 |
| Sspon.06G0015520 | 2,833 | 0,0059 |
| Sspon.06G0015570 | -1,278 | 0,0059 |

| | | |
|---|---|---|
| Sspon.06G0015610 | 14,295 | 0,0059 |
| Sspon.06G0015630 | -7,133 | 0,0059 |
| Sspon.06G0015780 | -2,347 | 0,0060 |
| Sspon.06G0015800 | 3,177 | 0,0060 |
| Sspon.06G0015820 | 1,141 | 0,0060 |
| Sspon.06G0015860 | -2,714 | 0,0060 |
| Sspon.06G0015870 | 1,312 | 0,0060 |
| Sspon.06G0016040 | -1,091 | 0,0060 |
| Sspon.06G0016170 | 1,065 | 0,0060 |
| Sspon.06G0016190 | 4,780 | 0,0060 |
| Sspon.06G0016200 | 1,224 | 0,0061 |
| Sspon.06G0016240 | 1,403 | 0,0061 |
| Sspon.06G0016250 | -5,349 | 0,0061 |
| Sspon.06G0016330 | 4,118 | 0,0061 |
| Sspon.06G0016360 | -1,971 | 0,0061 |
| Sspon.06G0016410 | 1,857 | 0,0061 |
| Sspon.06G0016420 | 1,088 | 0,0061 |
| Sspon.06G0016430 | 6,298 | 0,0061 |
| Sspon.06G0016590 | 1,363 | 0,0061 |
| Sspon.06G0016620 | -2,193 | 0,0061 |
| Sspon.06G0016810 | 1,628 | 0,0061 |
| Sspon.06G0016850 | 1,319 | 0,0061 |
| Sspon.06G0016930 | 2,499 | 0,0061 |
| Sspon.06G0016980 | -2,131 | 0,0062 |
| Sspon.06G0017020 | -1,052 | 0,0062 |
| Sspon.06G0017090 | 6,472 | 0,0062 |
| Sspon.06G0017140 | -2,644 | 0,0062 |
| Sspon.06G0017230 | 1,794 | 0,0062 |
| Sspon.06G0017330 | 19,128 | 0,0062 |
| Sspon.06G0017340 | 3,169 | 0,0062 |
| Sspon.06G0017430 | -1,375 | 0,0062 |
| Sspon.06G0017480 | 1,649 | 0,0062 |
| Sspon.06G0017510 | -1,054 | 0,0062 |
| Sspon.06G0017550 | 1,112 | 0,0062 |
| Sspon.06G0017610 | -2,195 | 0,0062 |
| Sspon.06G0017640 | 1,018 | 0,0062 |
| Sspon.06G0017650 | -7,272 | 0,0062 |
| Sspon.06G0017670 | 15,822 | 0,0062 |
| Sspon.06G0017680 | 1,057 | 0,0062 |
| Sspon.06G0017720 | 1,139 | 0,0063 |
| Sspon.06G0017770 | 2,189 | 0,0063 |
| Sspon.06G0017860 | -7,157 | 0,0063 |
| Sspon.06G0017930 | 1,011 | 0,0063 |

| | | |
|---|---|---|
| Sspon.06G0018020 | -2,284 | 0,0063 |
| Sspon.06G0018070 | 1,075 | 0,0063 |
| Sspon.06G0018100 | 1,502 | 0,0063 |
| Sspon.06G0018120 | 1,883 | 0,0063 |
| Sspon.06G0018130 | 15,181 | 0,0063 |
| Sspon.06G0018150 | -1,078 | 0,0063 |
| Sspon.06G0018160 | 1,422 | 0,0063 |
| Sspon.06G0018180 | 1,591 | 0,0064 |
| Sspon.06G0018220 | 9,055 | 0,0064 |
| Sspon.06G0018250 | 1,256 | 0,0064 |
| Sspon.06G0018420 | 5,002 | 0,0064 |
| Sspon.06G0018430 | 2,411 | 0,0064 |
| Sspon.06G0018450 | 1,360 | 0,0064 |
| Sspon.06G0018480 | -6,802 | 0,0064 |
| Sspon.06G0018510 | 2,594 | 0,0064 |
| Sspon.06G0018560 | -1,133 | 0,0064 |
| Sspon.06G0018590 | -1,967 | 0,0064 |
| Sspon.06G0018630 | 2,128 | 0,0065 |
| Sspon.06G0018650 | 1,513 | 0,0065 |
| Sspon.06G0018660 | -3,118 | 0,0065 |
| Sspon.06G0018710 | -1,059 | 0,0065 |
| Sspon.06G0018760 | -1,031 | 0,0065 |
| Sspon.06G0018800 | -1,235 | 0,0066 |
| Sspon.06G0019020 | -5,859 | 0,0066 |
| Sspon.06G0019190 | 1,119 | 0,0066 |
| Sspon.06G0019260 | 1,441 | 0,0066 |
| Sspon.06G0019320 | 2,149 | 0,0066 |
| Sspon.06G0019340 | -3,781 | 0,0066 |
| Sspon.06G0019360 | -6,817 | 0,0066 |
| Sspon.06G0019390 | 1,393 | 0,0066 |
| Sspon.06G0019400 | -1,974 | 0,0066 |
| Sspon.06G0019410 | -1,290 | 0,0066 |
| Sspon.06G0019460 | 1,089 | 0,0066 |
| Sspon.06G0019510 | 4,831 | 0,0066 |
| Sspon.06G0019570 | -1,033 | 0,0066 |
| Sspon.06G0019680 | 1,166 | 0,0066 |
| Sspon.06G0019760 | -1,042 | 0,0067 |
| Sspon.06G0019770 | -5,015 | 0,0067 |
| Sspon.06G0019800 | 4,720 | 0,0067 |
| Sspon.06G0019830 | 1,799 | 0,0067 |
| Sspon.06G0019840 | 3,145 | 0,0067 |
| Sspon.06G0019850 | -2,750 | 0,0067 |
| Sspon.06G0019920 | -4,442 | 0,0067 |

| | | |
|---|---|---|
| Sspon.06G0019970 | 2,558 | 0,0067 |
| Sspon.06G0020070 | 3,477 | 0,0067 |
| Sspon.06G0020170 | -2,335 | 0,0068 |
| Sspon.06G0020190 | -1,990 | 0,0068 |
| Sspon.06G0020210 | -1,165 | 0,0068 |
| Sspon.06G0020370 | -1,056 | 0,0068 |
| Sspon.06G0020460 | 2,884 | 0,0068 |
| Sspon.06G0020470 | -1,115 | 0,0068 |
| Sspon.06G0020640 | 3,117 | 0,0068 |
| Sspon.06G0020660 | 1,046 | 0,0069 |
| Sspon.06G0020690 | 5,511 | 0,0069 |
| Sspon.06G0020780 | 1,205 | 0,0069 |
| Sspon.06G0020860 | 1,954 | 0,0069 |
| Sspon.06G0021150 | -3,309 | 0,0069 |
| Sspon.06G0021200 | 1,151 | 0,0069 |
| Sspon.06G0021400 | 1,386 | 0,0069 |
| Sspon.06G0021410 | 1,774 | 0,0069 |
| Sspon.06G0021510 | -4,262 | 0,0069 |
| Sspon.06G0021550 | -1,968 | 0,0070 |
| Sspon.06G0021560 | 1,170 | 0,0070 |
| Sspon.06G0021570 | 1,259 | 0,0070 |
| Sspon.06G0021580 | -5,834 | 0,0070 |
| Sspon.06G0021590 | 5,274 | 0,0070 |
| Sspon.06G0021620 | 2,036 | 0,0070 |
| Sspon.06G0021790 | -4,887 | 0,0070 |
| Sspon.06G0021800 | 4,895 | 0,0071 |
| Sspon.06G0021810 | 1,155 | 0,0071 |
| Sspon.06G0021840 | -4,383 | 0,0071 |
| Sspon.06G0021860 | -6,220 | 0,0071 |
| Sspon.06G0021910 | 4,229 | 0,0071 |
| Sspon.06G0021990 | 1,041 | 0,0071 |
| Sspon.06G0022050 | -7,136 | 0,0071 |
| Sspon.06G0022060 | 2,796 | 0,0072 |
| Sspon.06G0022130 | -5,717 | 0,0072 |
| Sspon.06G0022200 | 1,716 | 0,0072 |
| Sspon.06G0022230 | 1,365 | 0,0072 |
| Sspon.06G0022260 | -1,220 | 0,0072 |
| Sspon.06G0022340 | 2,560 | 0,0072 |
| Sspon.06G0022370 | 2,081 | 0,0072 |
| Sspon.06G0022540 | 15,337 | 0,0072 |
| Sspon.06G0022560 | -2,284 | 0,0072 |
| Sspon.06G0022590 | -3,179 | 0,0072 |
| Sspon.06G0022600 | 1,419 | 0,0072 |

| | | |
|---|---|---|
| Sspon.06G0022650 | 2,235 | 0,0073 |
| Sspon.06G0022680 | 6,756 | 0,0073 |
| Sspon.06G0022690 | 2,783 | 0,0073 |
| Sspon.06G0022720 | -1,152 | 0,0073 |
| Sspon.06G0022810 | 17,107 | 0,0073 |
| Sspon.06G0022870 | 1,252 | 0,0073 |
| Sspon.06G0022890 | 1,074 | 0,0073 |
| Sspon.06G0022990 | 3,023 | 0,0073 |
| Sspon.06G0023090 | 1,200 | 0,0073 |
| Sspon.06G0023180 | 1,685 | 0,0073 |
| Sspon.06G0023220 | -4,421 | 0,0073 |
| Sspon.06G0023230 | -1,074 | 0,0074 |
| Sspon.06G0023240 | -1,863 | 0,0074 |
| Sspon.06G0023410 | 1,226 | 0,0074 |
| Sspon.06G0023600 | 1,257 | 0,0074 |
| Sspon.06G0023780 | 1,258 | 0,0074 |
| Sspon.06G0023810 | -1,173 | 0,0074 |
| Sspon.06G0023830 | 1,491 | 0,0074 |
| Sspon.06G0023940 | 2,624 | 0,0074 |
| Sspon.06G0023960 | -5,083 | 0,0074 |
| Sspon.06G0023970 | -1,313 | 0,0075 |
| Sspon.06G0023990 | 1,233 | 0,0075 |
| Sspon.06G0024100 | -4,027 | 0,0075 |
| Sspon.06G0024190 | -1,037 | 0,0075 |
| Sspon.06G0024210 | -6,696 | 0,0075 |
| Sspon.06G0024220 | -1,853 | 0,0076 |
| Sspon.06G0024250 | -6,687 | 0,0076 |
| Sspon.06G0024320 | -1,936 | 0,0076 |
| Sspon.06G0024370 | -3,290 | 0,0076 |
| Sspon.06G0024380 | 1,426 | 0,0076 |
| Sspon.06G0024410 | 1,097 | 0,0076 |
| Sspon.06G0024520 | 3,090 | 0,0076 |
| Sspon.06G0024580 | 4,003 | 0,0076 |
| Sspon.06G0024590 | -2,183 | 0,0076 |
| Sspon.06G0024610 | -1,845 | 0,0077 |
| Sspon.06G0024620 | -3,936 | 0,0077 |
| Sspon.06G0024750 | -1,980 | 0,0077 |
| Sspon.06G0025030 | 5,500 | 0,0077 |
| Sspon.06G0025060 | 1,553 | 0,0077 |
| Sspon.06G0025070 | 1,313 | 0,0077 |
| Sspon.06G0025090 | 1,651 | 0,0077 |
| Sspon.06G0025110 | 6,100 | 0,0077 |
| Sspon.06G0025180 | 2,586 | 0,0077 |

| | | |
|---|---|---|
| Sspon.06G0025190 | 1,312 | 0,0078 |
| Sspon.06G0025230 | 3,160 | 0,0078 |
| Sspon.06G0025260 | 1,341 | 0,0078 |
| Sspon.06G0025270 | 1,182 | 0,0078 |
| Sspon.06G0025300 | -6,148 | 0,0078 |
| Sspon.06G0025330 | 3,654 | 0,0078 |
| Sspon.06G0025360 | -2,641 | 0,0078 |
| Sspon.06G0025370 | 1,108 | 0,0078 |
| Sspon.06G0025380 | -2,730 | 0,0078 |
| Sspon.06G0025470 | 1,210 | 0,0079 |
| Sspon.06G0025530 | -2,464 | 0,0079 |
| Sspon.06G0025770 | -1,239 | 0,0079 |
| Sspon.06G0025840 | -5,165 | 0,0079 |
| Sspon.06G0025850 | -3,030 | 0,0079 |
| Sspon.06G0025870 | 3,289 | 0,0080 |
| Sspon.06G0025920 | 1,466 | 0,0080 |
| Sspon.06G0025990 | 3,820 | 0,0080 |
| Sspon.06G0026050 | -3,988 | 0,0080 |
| Sspon.06G0026060 | -6,934 | 0,0080 |
| Sspon.06G0026090 | 2,691 | 0,0080 |
| Sspon.06G0026130 | 1,294 | 0,0080 |
| Sspon.06G0026140 | 5,313 | 0,0080 |
| Sspon.06G0026150 | 6,657 | 0,0080 |
| Sspon.06G0026160 | -2,351 | 0,0080 |
| Sspon.06G0026300 | 4,720 | 0,0081 |
| Sspon.06G0026390 | 2,907 | 0,0081 |
| Sspon.06G0026410 | -4,932 | 0,0081 |
| Sspon.06G0026430 | 3,899 | 0,0081 |
| Sspon.06G0026460 | 1,325 | 0,0081 |
| Sspon.06G0026690 | -1,027 | 0,0081 |
| Sspon.06G0026770 | -3,961 | 0,0081 |
| Sspon.06G0026920 | -3,087 | 0,0081 |
| Sspon.06G0026930 | 2,042 | 0,0082 |
| Sspon.06G0027110 | -3,584 | 0,0082 |
| Sspon.06G0027140 | 1,289 | 0,0082 |
| Sspon.06G0027150 | -3,170 | 0,0082 |
| Sspon.06G0027200 | -2,013 | 0,0082 |
| Sspon.06G0027230 | -1,042 | 0,0082 |
| Sspon.06G0027240 | -1,125 | 0,0082 |
| Sspon.06G0027280 | 2,954 | 0,0082 |
| Sspon.06G0027340 | 1,155 | 0,0083 |
| Sspon.06G0027380 | 2,438 | 0,0083 |
| Sspon.06G0027410 | 16,534 | 0,0083 |

| | | |
|---|---|---|
| Sspon.06G0027470 | 2,001 | 0,0083 |
| Sspon.06G0027490 | 1,967 | 0,0083 |
| Sspon.06G0027530 | 1,169 | 0,0083 |
| Sspon.06G0027580 | 16,177 | 0,0084 |
| Sspon.06G0027590 | -2,032 | 0,0084 |
| Sspon.06G0027610 | 5,511 | 0,0084 |
| Sspon.06G0027750 | 1,031 | 0,0084 |
| Sspon.06G0027760 | -1,140 | 0,0084 |
| Sspon.06G0027810 | 1,195 | 0,0084 |
| Sspon.06G0027820 | 4,503 | 0,0084 |
| Sspon.06G0027900 | -1,543 | 0,0084 |
| Sspon.06G0027920 | 1,481 | 0,0084 |
| Sspon.06G0028010 | -2,075 | 0,0084 |
| Sspon.06G0028040 | 6,698 | 0,0084 |
| Sspon.06G0028110 | -1,257 | 0,0084 |
| Sspon.06G0028210 | 1,308 | 0,0085 |
| Sspon.06G0028240 | 5,984 | 0,0085 |
| Sspon.06G0028270 | -1,004 | 0,0085 |
| Sspon.06G0028290 | 2,522 | 0,0085 |
| Sspon.06G0028340 | 2,294 | 0,0085 |
| Sspon.06G0028420 | 6,133 | 0,0085 |
| Sspon.06G0028430 | -5,703 | 0,0086 |
| Sspon.06G0028460 | -4,400 | 0,0086 |
| Sspon.06G0028480 | 1,253 | 0,0086 |
| Sspon.06G0028490 | 6,788 | 0,0086 |
| Sspon.06G0028540 | 2,093 | 0,0086 |
| Sspon.06G0028570 | -2,073 | 0,0086 |
| Sspon.06G0028580 | 2,656 | 0,0086 |
| Sspon.06G0028640 | 1,094 | 0,0086 |
| Sspon.06G0028680 | -6,623 | 0,0087 |
| Sspon.06G0028710 | -4,284 | 0,0087 |
| Sspon.06G0028850 | 1,069 | 0,0087 |
| Sspon.06G0029090 | -5,101 | 0,0087 |
| Sspon.06G0029170 | 3,951 | 0,0087 |
| Sspon.06G0029310 | 6,611 | 0,0087 |
| Sspon.06G0029350 | -1,902 | 0,0087 |
| Sspon.06G0029410 | -3,555 | 0,0087 |
| Sspon.06G0029430 | 1,133 | 0,0087 |
| Sspon.06G0029440 | -6,337 | 0,0087 |
| Sspon.06G0029460 | 2,480 | 0,0087 |
| Sspon.06G0029480 | -1,357 | 0,0088 |
| Sspon.06G0029490 | 2,489 | 0,0088 |
| Sspon.06G0029500 | -3,408 | 0,0088 |

| | | |
|---|---|---|
| Sspon.06G0029550 | -3,568 | 0,0088 |
| Sspon.06G0029560 | -5,123 | 0,0088 |
| Sspon.06G0029750 | 4,994 | 0,0088 |
| Sspon.06G0029820 | -6,084 | 0,0088 |
| Sspon.06G0029860 | -2,952 | 0,0088 |
| Sspon.06G0029990 | -3,284 | 0,0088 |
| Sspon.06G0030100 | 4,421 | 0,0089 |
| Sspon.06G0030140 | 1,532 | 0,0089 |
| Sspon.06G0030230 | -1,963 | 0,0089 |
| Sspon.06G0030480 | 1,214 | 0,0089 |
| Sspon.06G0030540 | -2,552 | 0,0089 |
| Sspon.06G0030560 | -5,236 | 0,0090 |
| Sspon.06G0030590 | 4,476 | 0,0090 |
| Sspon.06G0030660 | -1,132 | 0,0090 |
| Sspon.06G0030680 | 1,319 | 0,0090 |
| Sspon.06G0030800 | 1,076 | 0,0090 |
| Sspon.06G0030920 | -4,154 | 0,0090 |
| Sspon.06G0030970 | 1,411 | 0,0090 |
| Sspon.06G0030980 | 6,810 | 0,0090 |
| Sspon.06G0031050 | 2,527 | 0,0090 |
| Sspon.06G0031080 | 1,470 | 0,0090 |
| Sspon.06G0031200 | -2,001 | 0,0090 |
| Sspon.06G0031230 | 1,171 | 0,0090 |
| Sspon.06G0031240 | 5,287 | 0,0090 |
| Sspon.06G0031280 | 1,483 | 0,0091 |
| Sspon.06G0031310 | 2,642 | 0,0091 |
| Sspon.06G0031330 | 6,493 | 0,0091 |
| Sspon.06G0031380 | -2,401 | 0,0091 |
| Sspon.06G0031430 | -2,651 | 0,0091 |
| Sspon.06G0031440 | 1,264 | 0,0092 |
| Sspon.06G0031450 | 1,053 | 0,0092 |
| Sspon.06G0031460 | 15,075 | 0,0093 |
| Sspon.06G0031480 | 1,855 | 0,0093 |
| Sspon.06G0031510 | -2,699 | 0,0093 |
| Sspon.06G0031520 | 3,809 | 0,0093 |
| Sspon.06G0031560 | 1,458 | 0,0093 |
| Sspon.06G0031570 | 6,672 | 0,0093 |
| Sspon.06G0031590 | -1,823 | 0,0093 |
| Sspon.06G0031670 | 5,472 | 0,0093 |
| Sspon.06G0031680 | 6,752 | 0,0093 |
| Sspon.06G0031720 | 15,619 | 0,0093 |
| Sspon.06G0031790 | 4,908 | 0,0093 |
| Sspon.06G0031810 | -6,275 | 0,0094 |

| | | |
|---|---|---|
| Sspon.06G0031840 | 1,212 | 0,0094 |
| Sspon.06G0031880 | 4,742 | 0,0094 |
| Sspon.06G0031910 | -5,060 | 0,0094 |
| Sspon.06G0031950 | 2,249 | 0,0095 |
| Sspon.06G0032040 | -1,060 | 0,0095 |
| Sspon.06G0032080 | -1,901 | 0,0095 |
| Sspon.06G0032260 | 1,156 | 0,0095 |
| Sspon.06G0032300 | -4,693 | 0,0095 |
| Sspon.06G0032310 | 1,275 | 0,0095 |
| Sspon.06G0032390 | 1,369 | 0,0095 |
| Sspon.06G0032450 | -1,849 | 0,0095 |
| Sspon.06G0032490 | -1,112 | 0,0096 |
| Sspon.06G0032510 | 1,203 | 0,0096 |
| Sspon.06G0032530 | -1,004 | 0,0096 |
| Sspon.06G0032580 | 1,366 | 0,0097 |
| Sspon.06G0032630 | -1,348 | 0,0097 |
| Sspon.06G0032650 | 6,033 | 0,0097 |
| Sspon.06G0032680 | -1,669 | 0,0097 |
| Sspon.06G0032800 | -5,355 | 0,0097 |
| Sspon.06G0032840 | -3,043 | 0,0097 |
| Sspon.06G0032850 | -4,420 | 0,0098 |
| Sspon.06G0032910 | 1,072 | 0,0098 |
| Sspon.06G0032930 | 4,550 | 0,0098 |
| Sspon.06G0032940 | -2,134 | 0,0098 |
| Sspon.06G0032950 | 1,380 | 0,0098 |
| Sspon.06G0032960 | 1,261 | 0,0098 |
| Sspon.06G0032970 | 2,295 | 0,0099 |
| Sspon.06G0032980 | -1,373 | 0,0099 |
| Sspon.06G0033080 | 2,669 | 0,0099 |
| Sspon.06G0033090 | -1,006 | 0,0099 |
| Sspon.06G0033110 | 15,231 | 0,0099 |
| Sspon.06G0033340 | -1,045 | 0,0100 |
| Sspon.06G0033450 | -2,624 | 0,0100 |
| Sspon.06G0033500 | -1,800 | 0,0100 |
| Sspon.06G0033540 | -3,052 | 0,0100 |
| Sspon.06G0033560 | 1,114 | 0,0101 |
| Sspon.06G0033570 | 2,828 | 0,0101 |
| Sspon.06G0033760 | 5,322 | 0,0101 |
| Sspon.06G0033780 | 6,411 | 0,0101 |
| Sspon.06G0033880 | -1,435 | 0,0101 |
| Sspon.06G0033960 | 1,549 | 0,0101 |
| Sspon.06G0034100 | -1,205 | 0,0101 |
| Sspon.06G0034160 | 1,621 | 0,0102 |

| | | |
|---|---|---|
| Sspon.06G0034420 | 1,310 | 0,0102 |
| Sspon.06G0034430 | 1,295 | 0,0102 |
| Sspon.06G0034480 | 1,579 | 0,0102 |
| Sspon.06G0034520 | 2,150 | 0,0102 |
| Sspon.06G0034540 | -1,707 | 0,0103 |
| Sspon.06G0034580 | 2,914 | 0,0103 |
| Sspon.06G0034600 | 1,325 | 0,0103 |
| Sspon.06G0034610 | 15,334 | 0,0103 |
| Sspon.06G0034630 | 1,675 | 0,0103 |
| Sspon.06G0034660 | 1,744 | 0,0103 |
| Sspon.06G0034680 | 2,826 | 0,0103 |
| Sspon.06G0034690 | 1,243 | 0,0104 |
| Sspon.06G0034710 | -3,157 | 0,0104 |
| Sspon.06G0034730 | 5,076 | 0,0104 |
| Sspon.06G0034790 | 3,660 | 0,0104 |
| Sspon.06G0034860 | -1,938 | 0,0104 |
| Sspon.06G0034880 | -6,720 | 0,0104 |
| Sspon.06G0034930 | 1,153 | 0,0105 |
| Sspon.06G0035180 | -4,538 | 0,0105 |
| Sspon.06G0035190 | -1,974 | 0,0105 |
| Sspon.06G0035210 | 1,498 | 0,0105 |
| Sspon.06G0035230 | 1,034 | 0,0105 |
| Sspon.06G0035240 | -1,086 | 0,0105 |
| Sspon.06G0035250 | 1,845 | 0,0105 |
| Sspon.06G0035310 | 1,532 | 0,0105 |
| Sspon.06G0035320 | 1,798 | 0,0105 |
| Sspon.06G0035340 | -4,718 | 0,0106 |
| Sspon.06G0035410 | -1,085 | 0,0106 |
| Sspon.06G0035440 | 5,524 | 0,0106 |
| Sspon.06G0035550 | 1,356 | 0,0106 |
| Sspon.06G0035560 | 1,454 | 0,0106 |
| Sspon.06G0035580 | 6,129 | 0,0106 |
| Sspon.06G0035600 | 1,032 | 0,0106 |
| Sspon.06G0035620 | 4,380 | 0,0106 |
| Sspon.06G0035710 | 1,428 | 0,0106 |
| Sspon.06G0035790 | -2,102 | 0,0106 |
| Sspon.06G0035810 | 1,541 | 0,0106 |
| Sspon.06G0035820 | 1,056 | 0,0106 |
| Sspon.06G0035890 | 1,822 | 0,0106 |
| Sspon.06G0035910 | -4,685 | 0,0106 |
| Sspon.06G0036060 | 2,326 | 0,0107 |
| Sspon.06G0036110 | -1,932 | 0,0107 |
| Sspon.06G0036130 | 3,148 | 0,0107 |

| | | |
|---|---|---|
| Sspon.06G0036140 | 2,683 | 0,0107 |
| Sspon.06G0036170 | 2,710 | 0,0107 |
| Sspon.06G0036210 | 5,947 | 0,0107 |
| Sspon.07G0000070 | -2,030 | 0,0108 |
| Sspon.07G0000100 | -1,986 | 0,0108 |
| Sspon.07G0000110 | 1,356 | 0,0108 |
| Sspon.07G0000120 | 6,656 | 0,0108 |
| Sspon.07G0000190 | 1,563 | 0,0108 |
| Sspon.07G0000300 | 7,406 | 0,0108 |
| Sspon.07G0000510 | 5,645 | 0,0108 |
| Sspon.07G0000750 | -4,686 | 0,0108 |
| Sspon.07G0000760 | 1,127 | 0,0108 |
| Sspon.07G0000800 | 1,327 | 0,0109 |
| Sspon.07G0000850 | -2,077 | 0,0109 |
| Sspon.07G0000860 | 3,162 | 0,0109 |
| Sspon.07G0001130 | -4,645 | 0,0109 |
| Sspon.07G0001190 | -7,367 | 0,0109 |
| Sspon.07G0001230 | 3,982 | 0,0109 |
| Sspon.07G0001240 | 6,582 | 0,0109 |
| Sspon.07G0001250 | -5,748 | 0,0110 |
| Sspon.07G0001270 | 1,381 | 0,0110 |
| Sspon.07G0001280 | 1,572 | 0,0110 |
| Sspon.07G0001290 | 1,172 | 0,0110 |
| Sspon.07G0001300 | 1,273 | 0,0110 |
| Sspon.07G0001310 | 1,582 | 0,0110 |
| Sspon.07G0001380 | 2,063 | 0,0110 |
| Sspon.07G0001540 | 5,753 | 0,0110 |
| Sspon.07G0001560 | -2,644 | 0,0110 |
| Sspon.07G0001590 | 7,719 | 0,0111 |
| Sspon.07G0001610 | -3,174 | 0,0111 |
| Sspon.07G0001620 | 1,015 | 0,0111 |
| Sspon.07G0001680 | 1,484 | 0,0111 |
| Sspon.07G0001720 | -6,439 | 0,0111 |
| Sspon.07G0001760 | -1,882 | 0,0111 |
| Sspon.07G0001780 | 6,591 | 0,0111 |
| Sspon.07G0001800 | 5,309 | 0,0111 |
| Sspon.07G0001820 | 1,178 | 0,0112 |
| Sspon.07G0001910 | 1,476 | 0,0112 |
| Sspon.07G0001980 | 1,151 | 0,0112 |
| Sspon.07G0002030 | 4,122 | 0,0112 |
| Sspon.07G0002100 | 1,341 | 0,0112 |
| Sspon.07G0002160 | 1,716 | 0,0112 |
| Sspon.07G0002170 | 1,873 | 0,0112 |

| | | |
|---|---|---|
| Sspon.07G0002190 | -2,795 | 0,0112 |
| Sspon.07G0002260 | -5,210 | 0,0113 |
| Sspon.07G0002310 | -2,826 | 0,0113 |
| Sspon.07G0002460 | 1,023 | 0,0113 |
| Sspon.07G0002490 | 1,206 | 0,0113 |
| Sspon.07G0002500 | 2,934 | 0,0113 |
| Sspon.07G0002530 | 1,608 | 0,0114 |
| Sspon.07G0002570 | 2,947 | 0,0114 |
| Sspon.07G0002600 | 2,202 | 0,0114 |
| Sspon.07G0002830 | 2,928 | 0,0114 |
| Sspon.07G0002840 | -4,835 | 0,0115 |
| Sspon.07G0002860 | 5,673 | 0,0115 |
| Sspon.07G0002890 | 1,338 | 0,0115 |
| Sspon.07G0002910 | 4,119 | 0,0115 |
| Sspon.07G0002940 | 5,233 | 0,0115 |
| Sspon.07G0002970 | 1,413 | 0,0116 |
| Sspon.07G0003000 | -1,297 | 0,0116 |
| Sspon.07G0003050 | 4,319 | 0,0116 |
| Sspon.07G0003090 | 2,409 | 0,0116 |
| Sspon.07G0003100 | 1,874 | 0,0116 |
| Sspon.07G0003190 | -1,509 | 0,0117 |
| Sspon.07G0003250 | 1,301 | 0,0117 |
| Sspon.07G0003260 | 1,065 | 0,0117 |
| Sspon.07G0003350 | -5,068 | 0,0117 |
| Sspon.07G0003410 | -1,003 | 0,0117 |
| Sspon.07G0003450 | -4,619 | 0,0117 |
| Sspon.07G0003470 | -5,851 | 0,0117 |
| Sspon.07G0003480 | 15,391 | 0,0117 |
| Sspon.07G0003540 | -2,008 | 0,0117 |
| Sspon.07G0003560 | -1,919 | 0,0117 |
| Sspon.07G0003580 | 9,952 | 0,0117 |
| Sspon.07G0003590 | 1,424 | 0,0117 |
| Sspon.07G0003620 | 1,134 | 0,0117 |
| Sspon.07G0003760 | 7,503 | 0,0117 |
| Sspon.07G0003780 | -5,617 | 0,0117 |
| Sspon.07G0003850 | 1,890 | 0,0118 |
| Sspon.07G0003860 | -2,102 | 0,0118 |
| Sspon.07G0003900 | -2,951 | 0,0118 |
| Sspon.07G0003910 | -2,649 | 0,0118 |
| Sspon.07G0003980 | 6,891 | 0,0118 |
| Sspon.07G0004150 | -4,172 | 0,0118 |
| Sspon.07G0004200 | -5,532 | 0,0118 |
| Sspon.07G0004210 | -2,529 | 0,0119 |

| | | |
|---|---|---|
| Sspon.07G0004260 | 1,106 | 0,0119 |
| Sspon.07G0004270 | -6,533 | 0,0119 |
| Sspon.07G0004310 | 5,949 | 0,0119 |
| Sspon.07G0004320 | 16,097 | 0,0119 |
| Sspon.07G0004440 | 2,517 | 0,0119 |
| Sspon.07G0004540 | -2,613 | 0,0119 |
| Sspon.07G0004590 | -2,961 | 0,0119 |
| Sspon.07G0004630 | -3,565 | 0,0120 |
| Sspon.07G0004650 | -1,173 | 0,0120 |
| Sspon.07G0004700 | 1,483 | 0,0120 |
| Sspon.07G0004750 | 1,118 | 0,0120 |
| Sspon.07G0004830 | -1,569 | 0,0120 |
| Sspon.07G0004840 | -6,767 | 0,0120 |
| Sspon.07G0004860 | 5,665 | 0,0120 |
| Sspon.07G0004910 | 1,653 | 0,0120 |
| Sspon.07G0004930 | -1,077 | 0,0121 |
| Sspon.07G0005040 | -5,231 | 0,0121 |
| Sspon.07G0005160 | 2,225 | 0,0121 |
| Sspon.07G0005190 | 2,089 | 0,0122 |
| Sspon.07G0005230 | 6,985 | 0,0122 |
| Sspon.07G0005240 | 1,616 | 0,0123 |
| Sspon.07G0005280 | -1,894 | 0,0123 |
| Sspon.07G0005330 | 2,830 | 0,0123 |
| Sspon.07G0005350 | 14,621 | 0,0123 |
| Sspon.07G0005470 | 1,043 | 0,0123 |
| Sspon.07G0005490 | -1,738 | 0,0123 |
| Sspon.07G0005550 | 1,738 | 0,0123 |
| Sspon.07G0005560 | 1,937 | 0,0123 |
| Sspon.07G0005740 | -2,488 | 0,0124 |
| Sspon.07G0005800 | 1,732 | 0,0124 |
| Sspon.07G0005860 | -3,665 | 0,0124 |
| Sspon.07G0005960 | 6,481 | 0,0125 |
| Sspon.07G0006000 | 1,049 | 0,0125 |
| Sspon.07G0006030 | 1,112 | 0,0125 |
| Sspon.07G0006070 | -5,806 | 0,0125 |
| Sspon.07G0006090 | 5,581 | 0,0125 |
| Sspon.07G0006210 | -4,479 | 0,0126 |
| Sspon.07G0006290 | 6,585 | 0,0126 |
| Sspon.07G0006330 | -2,781 | 0,0127 |
| Sspon.07G0006370 | 4,307 | 0,0127 |
| Sspon.07G0006380 | 3,484 | 0,0127 |
| Sspon.07G0006400 | 7,192 | 0,0128 |
| Sspon.07G0006450 | 1,049 | 0,0128 |

| | | |
|---|---|---|
| Sspon.07G0006500 | -2,360 | 0,0128 |
| Sspon.07G0006520 | 1,460 | 0,0128 |
| Sspon.07G0006570 | 15,917 | 0,0128 |
| Sspon.07G0006630 | 1,979 | 0,0129 |
| Sspon.07G0006670 | 2,567 | 0,0129 |
| Sspon.07G0006680 | 7,014 | 0,0129 |
| Sspon.07G0006740 | 2,063 | 0,0129 |
| Sspon.07G0006770 | -3,291 | 0,0130 |
| Sspon.07G0006820 | 1,406 | 0,0130 |
| Sspon.07G0006870 | 2,644 | 0,0131 |
| Sspon.07G0006880 | -1,032 | 0,0131 |
| Sspon.07G0006890 | -3,615 | 0,0131 |
| Sspon.07G0006900 | 1,113 | 0,0131 |
| Sspon.07G0007000 | 1,163 | 0,0132 |
| Sspon.07G0007050 | -1,136 | 0,0132 |
| Sspon.07G0007070 | -5,953 | 0,0132 |
| Sspon.07G0007080 | 1,697 | 0,0132 |
| Sspon.07G0007100 | 1,342 | 0,0133 |
| Sspon.07G0007200 | 17,477 | 0,0133 |
| Sspon.07G0007360 | -1,813 | 0,0133 |
| Sspon.07G0007410 | 3,301 | 0,0133 |
| Sspon.07G0007420 | -2,954 | 0,0133 |
| Sspon.07G0007480 | -1,999 | 0,0134 |
| Sspon.07G0007500 | 2,910 | 0,0134 |
| Sspon.07G0007570 | 3,345 | 0,0134 |
| Sspon.07G0007580 | 1,845 | 0,0134 |
| Sspon.07G0007620 | -1,045 | 0,0135 |
| Sspon.07G0007730 | -3,083 | 0,0135 |
| Sspon.07G0007770 | 5,782 | 0,0135 |
| Sspon.07G0007900 | -1,035 | 0,0136 |
| Sspon.07G0007930 | -4,741 | 0,0136 |
| Sspon.07G0007980 | 3,280 | 0,0137 |
| Sspon.07G0008000 | 18,198 | 0,0137 |
| Sspon.07G0008140 | 1,507 | 0,0137 |
| Sspon.07G0008160 | -2,396 | 0,0137 |
| Sspon.07G0008210 | -1,999 | 0,0137 |
| Sspon.07G0008260 | 1,119 | 0,0137 |
| Sspon.07G0008290 | 3,670 | 0,0137 |
| Sspon.07G0008300 | -2,680 | 0,0137 |
| Sspon.07G0008390 | -6,661 | 0,0138 |
| Sspon.07G0008410 | 1,211 | 0,0138 |
| Sspon.07G0008420 | -5,222 | 0,0138 |
| Sspon.07G0008450 | 2,304 | 0,0138 |

| | | |
|---|---|---|
| Sspon.07G0008580 | -1,023 | 0,0138 |
| Sspon.07G0008630 | 1,646 | 0,0138 |
| Sspon.07G0008700 | 1,480 | 0,0139 |
| Sspon.07G0008760 | 14,460 | 0,0139 |
| Sspon.07G0008810 | 2,689 | 0,0139 |
| Sspon.07G0008870 | 2,523 | 0,0139 |
| Sspon.07G0008880 | -2,452 | 0,0139 |
| Sspon.07G0008920 | -2,373 | 0,0139 |
| Sspon.07G0008980 | 3,385 | 0,0139 |
| Sspon.07G0009020 | -1,733 | 0,0140 |
| Sspon.07G0009150 | -2,504 | 0,0140 |
| Sspon.07G0009220 | -4,470 | 0,0140 |
| Sspon.07G0009230 | -4,000 | 0,0141 |
| Sspon.07G0009240 | 15,908 | 0,0141 |
| Sspon.07G0009290 | -3,178 | 0,0141 |
| Sspon.07G0009310 | 2,411 | 0,0142 |
| Sspon.07G0009320 | -2,736 | 0,0142 |
| Sspon.07G0009340 | 1,549 | 0,0142 |
| Sspon.07G0009360 | 17,974 | 0,0142 |
| Sspon.07G0009470 | -4,770 | 0,0142 |
| Sspon.07G0009530 | 4,335 | 0,0142 |
| Sspon.07G0009560 | 5,553 | 0,0143 |
| Sspon.07G0009640 | -5,375 | 0,0143 |
| Sspon.07G0009730 | 2,243 | 0,0144 |
| Sspon.07G0009770 | -1,142 | 0,0144 |
| Sspon.07G0009780 | 1,018 | 0,0144 |
| Sspon.07G0009890 | 2,429 | 0,0145 |
| Sspon.07G0010030 | 1,207 | 0,0145 |
| Sspon.07G0010130 | 14,315 | 0,0145 |
| Sspon.07G0010140 | 1,168 | 0,0145 |
| Sspon.07G0010160 | -1,061 | 0,0145 |
| Sspon.07G0010180 | -4,826 | 0,0145 |
| Sspon.07G0010240 | 1,115 | 0,0146 |
| Sspon.07G0010250 | 5,275 | 0,0146 |
| Sspon.07G0010260 | -1,907 | 0,0146 |
| Sspon.07G0010320 | 3,837 | 0,0146 |
| Sspon.07G0010340 | 1,156 | 0,0146 |
| Sspon.07G0010400 | 1,507 | 0,0147 |
| Sspon.07G0010720 | -2,422 | 0,0147 |
| Sspon.07G0010780 | 4,140 | 0,0147 |
| Sspon.07G0010800 | 1,791 | 0,0147 |
| Sspon.07G0010830 | -1,107 | 0,0147 |
| Sspon.07G0010880 | 1,356 | 0,0148 |

| | | |
|---|---|---|
| Sspon.07G0010890 | -2,392 | 0,0148 |
| Sspon.07G0010940 | 1,307 | 0,0148 |
| Sspon.07G0010980 | 5,564 | 0,0148 |
| Sspon.07G0010990 | -3,605 | 0,0148 |
| Sspon.07G0011020 | -6,111 | 0,0149 |
| Sspon.07G0011050 | -3,140 | 0,0149 |
| Sspon.07G0011210 | 16,255 | 0,0149 |
| Sspon.07G0011300 | 15,272 | 0,0150 |
| Sspon.07G0011310 | 2,196 | 0,0150 |
| Sspon.07G0011330 | -2,069 | 0,0150 |
| Sspon.07G0011420 | 15,569 | 0,0150 |
| Sspon.07G0011520 | -6,021 | 0,0150 |
| Sspon.07G0011570 | 4,299 | 0,0150 |
| Sspon.07G0011580 | -5,399 | 0,0151 |
| Sspon.07G0011630 | 4,969 | 0,0151 |
| Sspon.07G0011710 | 1,007 | 0,0152 |
| Sspon.07G0011750 | 6,393 | 0,0152 |
| Sspon.07G0011830 | 1,289 | 0,0152 |
| Sspon.07G0011850 | -3,944 | 0,0152 |
| Sspon.07G0011900 | 1,014 | 0,0152 |
| Sspon.07G0011970 | 2,578 | 0,0152 |
| Sspon.07G0012030 | -2,253 | 0,0152 |
| Sspon.07G0012120 | 1,037 | 0,0153 |
| Sspon.07G0012150 | 2,880 | 0,0153 |
| Sspon.07G0012300 | -1,914 | 0,0153 |
| Sspon.07G0012330 | -3,319 | 0,0154 |
| Sspon.07G0012340 | -1,759 | 0,0154 |
| Sspon.07G0012370 | -4,658 | 0,0154 |
| Sspon.07G0012490 | -5,800 | 0,0154 |
| Sspon.07G0012500 | -5,002 | 0,0154 |
| Sspon.07G0012510 | 4,111 | 0,0154 |
| Sspon.07G0012690 | 1,006 | 0,0155 |
| Sspon.07G0012710 | 15,036 | 0,0155 |
| Sspon.07G0012730 | 1,826 | 0,0155 |
| Sspon.07G0012760 | 1,195 | 0,0155 |
| Sspon.07G0012820 | 15,071 | 0,0155 |
| Sspon.07G0012960 | 5,581 | 0,0155 |
| Sspon.07G0013130 | -1,769 | 0,0155 |
| Sspon.07G0013140 | 1,468 | 0,0156 |
| Sspon.07G0013160 | 4,835 | 0,0156 |
| Sspon.07G0013280 | -3,337 | 0,0156 |
| Sspon.07G0013330 | 14,972 | 0,0156 |
| Sspon.07G0013510 | -1,066 | 0,0156 |

| | | |
|---|---|---|
| Sspon.07G0013570 | -1,016 | 0,0157 |
| Sspon.07G0013680 | 2,894 | 0,0157 |
| Sspon.07G0013730 | 16,187 | 0,0157 |
| Sspon.07G0013790 | 1,429 | 0,0157 |
| Sspon.07G0013800 | -1,154 | 0,0157 |
| Sspon.07G0013840 | -1,675 | 0,0157 |
| Sspon.07G0013900 | 1,482 | 0,0158 |
| Sspon.07G0013910 | 4,154 | 0,0158 |
| Sspon.07G0014020 | -4,862 | 0,0158 |
| Sspon.07G0014030 | 3,601 | 0,0159 |
| Sspon.07G0014040 | -2,658 | 0,0159 |
| Sspon.07G0014120 | 15,927 | 0,0159 |
| Sspon.07G0014130 | 6,289 | 0,0160 |
| Sspon.07G0014170 | 1,837 | 0,0160 |
| Sspon.07G0014200 | -3,301 | 0,0161 |
| Sspon.07G0014270 | -1,278 | 0,0161 |
| Sspon.07G0014300 | 1,080 | 0,0161 |
| Sspon.07G0014310 | -3,138 | 0,0161 |
| Sspon.07G0014340 | -6,346 | 0,0162 |
| Sspon.07G0014380 | 1,087 | 0,0162 |
| Sspon.07G0014420 | -4,673 | 0,0162 |
| Sspon.07G0014450 | 1,229 | 0,0162 |
| Sspon.07G0014470 | 1,901 | 0,0163 |
| Sspon.07G0014480 | -6,102 | 0,0163 |
| Sspon.07G0014590 | -6,368 | 0,0164 |
| Sspon.07G0014660 | 1,057 | 0,0164 |
| Sspon.07G0014710 | 2,112 | 0,0164 |
| Sspon.07G0014740 | 3,563 | 0,0164 |
| Sspon.07G0014790 | 2,538 | 0,0164 |
| Sspon.07G0014800 | -1,242 | 0,0164 |
| Sspon.07G0014830 | 1,746 | 0,0165 |
| Sspon.07G0014840 | -4,051 | 0,0165 |
| Sspon.07G0014940 | 4,318 | 0,0165 |
| Sspon.07G0014960 | 1,145 | 0,0165 |
| Sspon.07G0014970 | -2,316 | 0,0165 |
| Sspon.07G0014980 | -6,542 | 0,0165 |
| Sspon.07G0015050 | 1,730 | 0,0166 |
| Sspon.07G0015090 | 5,087 | 0,0166 |
| Sspon.07G0015110 | 2,363 | 0,0167 |
| Sspon.07G0015120 | -4,894 | 0,0167 |
| Sspon.07G0015150 | 1,211 | 0,0168 |
| Sspon.07G0015230 | 3,629 | 0,0168 |
| Sspon.07G0015240 | 1,419 | 0,0168 |

| | | |
|---|---|---|
| Sspon.07G0015250 | 1,146 | 0,0168 |
| Sspon.07G0015310 | 3,351 | 0,0168 |
| Sspon.07G0015380 | -1,494 | 0,0168 |
| Sspon.07G0015400 | 1,311 | 0,0169 |
| Sspon.07G0015410 | -2,404 | 0,0169 |
| Sspon.07G0015430 | -1,804 | 0,0169 |
| Sspon.07G0015520 | 4,188 | 0,0169 |
| Sspon.07G0015610 | 5,923 | 0,0170 |
| Sspon.07G0015660 | -1,282 | 0,0170 |
| Sspon.07G0015670 | 2,227 | 0,0170 |
| Sspon.07G0015720 | -4,779 | 0,0170 |
| Sspon.07G0015750 | 1,441 | 0,0170 |
| Sspon.07G0015800 | 1,664 | 0,0171 |
| Sspon.07G0015880 | 3,165 | 0,0171 |
| Sspon.07G0015990 | 3,462 | 0,0171 |
| Sspon.07G0016030 | -2,371 | 0,0171 |
| Sspon.07G0016120 | 1,079 | 0,0171 |
| Sspon.07G0016130 | 1,066 | 0,0171 |
| Sspon.07G0016300 | -4,029 | 0,0172 |
| Sspon.07G0016470 | -1,218 | 0,0172 |
| Sspon.07G0016560 | 6,666 | 0,0172 |
| Sspon.07G0016620 | -4,317 | 0,0172 |
| Sspon.07G0016680 | -2,436 | 0,0172 |
| Sspon.07G0016700 | 1,433 | 0,0173 |
| Sspon.07G0016740 | -1,377 | 0,0173 |
| Sspon.07G0016780 | -2,712 | 0,0174 |
| Sspon.07G0016790 | -3,303 | 0,0174 |
| Sspon.07G0016800 | -4,651 | 0,0174 |
| Sspon.07G0016810 | -4,758 | 0,0174 |
| Sspon.07G0016870 | 1,348 | 0,0174 |
| Sspon.07G0016930 | -4,431 | 0,0175 |
| Sspon.07G0017020 | 3,472 | 0,0175 |
| Sspon.07G0017060 | -2,847 | 0,0175 |
| Sspon.07G0017100 | -5,804 | 0,0175 |
| Sspon.07G0017110 | -2,319 | 0,0175 |
| Sspon.07G0017140 | 1,170 | 0,0176 |
| Sspon.07G0017190 | 2,003 | 0,0176 |
| Sspon.07G0017200 | -1,124 | 0,0176 |
| Sspon.07G0017250 | 3,270 | 0,0176 |
| Sspon.07G0017270 | -1,001 | 0,0177 |
| Sspon.07G0017280 | -2,804 | 0,0177 |
| Sspon.07G0017300 | 1,365 | 0,0177 |
| Sspon.07G0017330 | 1,082 | 0,0178 |

| | | |
|---|---|---|
| Sspon.07G0017440 | 1,517 | 0,0178 |
| Sspon.07G0017530 | 1,423 | 0,0178 |
| Sspon.07G0017580 | -2,792 | 0,0179 |
| Sspon.07G0017600 | 1,360 | 0,0179 |
| Sspon.07G0017650 | 2,581 | 0,0179 |
| Sspon.07G0017780 | -1,513 | 0,0179 |
| Sspon.07G0017790 | 1,764 | 0,0179 |
| Sspon.07G0017800 | 15,861 | 0,0179 |
| Sspon.07G0017810 | 5,576 | 0,0180 |
| Sspon.07G0017830 | 6,555 | 0,0180 |
| Sspon.07G0017840 | -7,192 | 0,0180 |
| Sspon.07G0017850 | -2,371 | 0,0180 |
| Sspon.07G0017860 | -2,344 | 0,0181 |
| Sspon.07G0017870 | -1,400 | 0,0181 |
| Sspon.07G0017890 | -2,552 | 0,0181 |
| Sspon.07G0017900 | -2,648 | 0,0181 |
| Sspon.07G0017920 | -2,743 | 0,0182 |
| Sspon.07G0017960 | -1,022 | 0,0182 |
| Sspon.07G0017980 | -4,306 | 0,0182 |
| Sspon.07G0018010 | -6,084 | 0,0182 |
| Sspon.07G0018050 | 1,336 | 0,0183 |
| Sspon.07G0018080 | 1,035 | 0,0183 |
| Sspon.07G0018090 | -5,040 | 0,0183 |
| Sspon.07G0018100 | 1,333 | 0,0183 |
| Sspon.07G0018110 | 2,301 | 0,0183 |
| Sspon.07G0018120 | -6,389 | 0,0183 |
| Sspon.07G0018130 | 1,039 | 0,0184 |
| Sspon.07G0018150 | 7,314 | 0,0184 |
| Sspon.07G0018210 | -3,224 | 0,0184 |
| Sspon.07G0018340 | 5,988 | 0,0185 |
| Sspon.07G0018490 | -2,087 | 0,0185 |
| Sspon.07G0018500 | -1,889 | 0,0185 |
| Sspon.07G0018610 | 2,652 | 0,0185 |
| Sspon.07G0018850 | -1,996 | 0,0185 |
| Sspon.07G0018990 | 2,009 | 0,0186 |
| Sspon.07G0019010 | -4,311 | 0,0186 |
| Sspon.07G0019110 | -1,927 | 0,0186 |
| Sspon.07G0019150 | 5,783 | 0,0186 |
| Sspon.07G0019170 | 6,231 | 0,0187 |
| Sspon.07G0019290 | 1,282 | 0,0187 |
| Sspon.07G0019300 | 1,120 | 0,0188 |
| Sspon.07G0019320 | 3,532 | 0,0188 |
| Sspon.07G0019360 | 5,039 | 0,0188 |

| | | |
|---|---|---|
| Sspon.07G0019370 | -1,041 | 0,0188 |
| Sspon.07G0019390 | -2,399 | 0,0189 |
| Sspon.07G0019440 | 2,034 | 0,0189 |
| Sspon.07G0019470 | 2,357 | 0,0189 |
| Sspon.07G0019540 | 1,305 | 0,0189 |
| Sspon.07G0019550 | 1,376 | 0,0190 |
| Sspon.07G0019640 | 2,395 | 0,0190 |
| Sspon.07G0019660 | 1,201 | 0,0191 |
| Sspon.07G0019680 | -1,657 | 0,0191 |
| Sspon.07G0019740 | -2,292 | 0,0191 |
| Sspon.07G0019750 | -1,888 | 0,0191 |
| Sspon.07G0019760 | 2,329 | 0,0191 |
| Sspon.07G0020060 | -3,805 | 0,0192 |
| Sspon.07G0020070 | -2,321 | 0,0192 |
| Sspon.07G0020270 | -5,743 | 0,0193 |
| Sspon.07G0020330 | -1,007 | 0,0193 |
| Sspon.07G0020400 | 3,822 | 0,0193 |
| Sspon.07G0020410 | -1,774 | 0,0193 |
| Sspon.07G0020550 | -4,166 | 0,0193 |
| Sspon.07G0020590 | -2,850 | 0,0194 |
| Sspon.07G0020610 | -5,671 | 0,0194 |
| Sspon.07G0020620 | -2,395 | 0,0194 |
| Sspon.07G0020700 | 1,782 | 0,0194 |
| Sspon.07G0020870 | 14,908 | 0,0194 |
| Sspon.07G0020890 | 5,392 | 0,0194 |
| Sspon.07G0020910 | -6,133 | 0,0194 |
| Sspon.07G0021050 | 2,470 | 0,0194 |
| Sspon.07G0021070 | 16,975 | 0,0195 |
| Sspon.07G0021140 | 1,332 | 0,0195 |
| Sspon.07G0021160 | -3,642 | 0,0195 |
| Sspon.07G0021190 | -3,141 | 0,0196 |
| Sspon.07G0021200 | 1,976 | 0,0196 |
| Sspon.07G0021210 | -2,826 | 0,0196 |
| Sspon.07G0021240 | -2,403 | 0,0196 |
| Sspon.07G0021290 | -3,608 | 0,0196 |
| Sspon.07G0021310 | 5,156 | 0,0196 |
| Sspon.07G0021320 | -1,047 | 0,0197 |
| Sspon.07G0021380 | -1,891 | 0,0197 |
| Sspon.07G0021390 | 6,008 | 0,0197 |
| Sspon.07G0021400 | 1,451 | 0,0197 |
| Sspon.07G0021440 | -1,795 | 0,0197 |
| Sspon.07G0021450 | 1,259 | 0,0198 |
| Sspon.07G0021480 | -1,863 | 0,0198 |

| | | |
|---|---|---|
| Sspon.07G0021520 | 2,279 | 0,0198 |
| Sspon.07G0021690 | 6,298 | 0,0199 |
| Sspon.07G0021730 | -1,638 | 0,0199 |
| Sspon.07G0021750 | 3,289 | 0,0199 |
| Sspon.07G0021870 | -3,431 | 0,0200 |
| Sspon.07G0021890 | -5,319 | 0,0200 |
| Sspon.07G0021900 | -5,177 | 0,0200 |
| Sspon.07G0021990 | 1,280 | 0,0200 |
| Sspon.07G0022000 | 2,373 | 0,0200 |
| Sspon.07G0022010 | -3,948 | 0,0201 |
| Sspon.07G0022020 | 1,016 | 0,0201 |
| Sspon.07G0022230 | 2,812 | 0,0201 |
| Sspon.07G0022290 | -5,188 | 0,0201 |
| Sspon.07G0022330 | -3,996 | 0,0201 |
| Sspon.07G0022350 | 5,314 | 0,0202 |
| Sspon.07G0022400 | -1,664 | 0,0202 |
| Sspon.07G0022500 | -1,527 | 0,0202 |
| Sspon.07G0022600 | -1,820 | 0,0202 |
| Sspon.07G0022640 | -5,661 | 0,0202 |
| Sspon.07G0022690 | 3,554 | 0,0202 |
| Sspon.07G0022780 | -5,460 | 0,0202 |
| Sspon.07G0022800 | -1,975 | 0,0203 |
| Sspon.07G0022850 | -5,162 | 0,0203 |
| Sspon.07G0022880 | 4,790 | 0,0203 |
| Sspon.07G0022900 | 1,073 | 0,0203 |
| Sspon.07G0023090 | -2,278 | 0,0203 |
| Sspon.07G0023110 | 6,006 | 0,0203 |
| Sspon.07G0023130 | 4,260 | 0,0204 |
| Sspon.07G0023180 | 7,741 | 0,0204 |
| Sspon.07G0023190 | 1,088 | 0,0204 |
| Sspon.07G0023210 | 2,201 | 0,0204 |
| Sspon.07G0023220 | 1,088 | 0,0205 |
| Sspon.07G0023230 | 3,603 | 0,0205 |
| Sspon.07G0023240 | 6,793 | 0,0205 |
| Sspon.07G0023450 | -5,319 | 0,0205 |
| Sspon.07G0023480 | 1,201 | 0,0205 |
| Sspon.07G0023530 | 1,221 | 0,0206 |
| Sspon.07G0023540 | 2,118 | 0,0206 |
| Sspon.07G0023690 | 1,309 | 0,0207 |
| Sspon.07G0023760 | 1,246 | 0,0207 |
| Sspon.07G0023860 | 2,288 | 0,0208 |
| Sspon.07G0023930 | 5,806 | 0,0208 |
| Sspon.07G0024000 | 5,791 | 0,0208 |

| | | |
|---|---|---|
| Sspon.07G0024070 | 1,210 | 0,0208 |
| Sspon.07G0024140 | 4,798 | 0,0208 |
| Sspon.07G0024290 | -4,553 | 0,0208 |
| Sspon.07G0024350 | 5,588 | 0,0208 |
| Sspon.07G0024560 | -1,999 | 0,0208 |
| Sspon.07G0024650 | 1,151 | 0,0209 |
| Sspon.07G0024750 | -4,205 | 0,0209 |
| Sspon.07G0024840 | 2,902 | 0,0209 |
| Sspon.07G0024970 | -5,231 | 0,0209 |
| Sspon.07G0025100 | -3,490 | 0,0209 |
| Sspon.07G0025330 | -5,617 | 0,0210 |
| Sspon.07G0025380 | 1,384 | 0,0210 |
| Sspon.07G0025400 | -2,705 | 0,0210 |
| Sspon.07G0025480 | 2,177 | 0,0210 |
| Sspon.07G0025500 | -2,972 | 0,0211 |
| Sspon.07G0025640 | -1,008 | 0,0211 |
| Sspon.07G0025730 | 2,546 | 0,0213 |
| Sspon.07G0025740 | 1,112 | 0,0213 |
| Sspon.07G0025760 | -4,361 | 0,0213 |
| Sspon.07G0025770 | -1,667 | 0,0214 |
| Sspon.07G0025890 | 1,973 | 0,0214 |
| Sspon.07G0025910 | 4,626 | 0,0214 |
| Sspon.07G0025930 | 1,989 | 0,0215 |
| Sspon.07G0025940 | 4,053 | 0,0215 |
| Sspon.07G0025980 | 6,369 | 0,0215 |
| Sspon.07G0025990 | 1,109 | 0,0215 |
| Sspon.07G0026060 | 3,039 | 0,0215 |
| Sspon.07G0026080 | 3,223 | 0,0217 |
| Sspon.07G0026130 | 6,619 | 0,0218 |
| Sspon.07G0026270 | -2,707 | 0,0218 |
| Sspon.07G0026320 | -1,023 | 0,0218 |
| Sspon.07G0026580 | -2,754 | 0,0219 |
| Sspon.07G0026590 | 1,225 | 0,0219 |
| Sspon.07G0026620 | -1,082 | 0,0219 |
| Sspon.07G0026650 | 1,463 | 0,0220 |
| Sspon.07G0026690 | -1,790 | 0,0220 |
| Sspon.07G0026700 | -3,046 | 0,0220 |
| Sspon.07G0026730 | -6,282 | 0,0220 |
| Sspon.07G0026840 | 1,001 | 0,0221 |
| Sspon.07G0026860 | 7,627 | 0,0221 |
| Sspon.07G0026880 | 1,313 | 0,0222 |
| Sspon.07G0026970 | -2,486 | 0,0222 |
| Sspon.07G0027100 | 5,901 | 0,0222 |

| | | |
|---|---|---|
| Sspon.07G0027190 | -1,142 | 0,0222 |
| Sspon.07G0027260 | -2,167 | 0,0222 |
| Sspon.07G0027340 | 4,478 | 0,0223 |
| Sspon.07G0027370 | -1,817 | 0,0223 |
| Sspon.07G0027440 | 1,842 | 0,0223 |
| Sspon.07G0027450 | 3,025 | 0,0223 |
| Sspon.07G0027460 | 1,221 | 0,0224 |
| Sspon.07G0027490 | -6,315 | 0,0224 |
| Sspon.07G0027510 | -1,908 | 0,0224 |
| Sspon.07G0027620 | -4,952 | 0,0224 |
| Sspon.07G0027780 | -4,268 | 0,0224 |
| Sspon.07G0027800 | 3,109 | 0,0224 |
| Sspon.07G0027820 | 2,566 | 0,0224 |
| Sspon.07G0027890 | -1,935 | 0,0225 |
| Sspon.07G0028160 | -3,355 | 0,0225 |
| Sspon.07G0028180 | -4,443 | 0,0225 |
| Sspon.07G0028240 | -1,871 | 0,0225 |
| Sspon.07G0028320 | 1,504 | 0,0226 |
| Sspon.07G0028370 | 1,427 | 0,0226 |
| Sspon.07G0028420 | 2,072 | 0,0226 |
| Sspon.07G0028460 | 5,656 | 0,0226 |
| Sspon.07G0028470 | 4,476 | 0,0227 |
| Sspon.07G0028510 | -1,744 | 0,0227 |
| Sspon.07G0028530 | 1,818 | 0,0227 |
| Sspon.07G0028640 | -3,646 | 0,0227 |
| Sspon.07G0028680 | 3,837 | 0,0228 |
| Sspon.07G0028690 | 2,533 | 0,0228 |
| Sspon.07G0028730 | -1,760 | 0,0229 |
| Sspon.07G0028800 | 4,669 | 0,0229 |
| Sspon.07G0028830 | -1,038 | 0,0229 |
| Sspon.07G0028880 | 15,011 | 0,0230 |
| Sspon.07G0028910 | 1,034 | 0,0230 |
| Sspon.07G0028930 | 1,165 | 0,0231 |
| Sspon.07G0028970 | 2,408 | 0,0231 |
| Sspon.07G0028980 | 1,355 | 0,0231 |
| Sspon.07G0029060 | 3,541 | 0,0231 |
| Sspon.07G0029180 | -1,017 | 0,0232 |
| Sspon.07G0029220 | 1,882 | 0,0233 |
| Sspon.07G0029230 | -1,676 | 0,0234 |
| Sspon.07G0029240 | 1,201 | 0,0234 |
| Sspon.07G0029260 | -2,305 | 0,0235 |
| Sspon.07G0029270 | 5,369 | 0,0235 |
| Sspon.07G0029320 | 1,277 | 0,0235 |

| | | |
|---|---|---|
| Sspon.07G0029500 | -2,764 | 0,0235 |
| Sspon.07G0029550 | 4,244 | 0,0236 |
| Sspon.07G0029630 | -1,163 | 0,0236 |
| Sspon.07G0029660 | 3,829 | 0,0236 |
| Sspon.07G0029670 | 3,292 | 0,0236 |
| Sspon.07G0029680 | -4,814 | 0,0238 |
| Sspon.07G0029690 | -5,309 | 0,0238 |
| Sspon.07G0029750 | 2,417 | 0,0238 |
| Sspon.07G0029830 | -1,804 | 0,0239 |
| Sspon.07G0029850 | -1,579 | 0,0239 |
| Sspon.07G0030000 | -2,966 | 0,0239 |
| Sspon.07G0030030 | 1,786 | 0,0240 |
| Sspon.07G0030060 | -1,952 | 0,0240 |
| Sspon.07G0030100 | -4,133 | 0,0241 |
| Sspon.07G0030130 | 3,766 | 0,0241 |
| Sspon.07G0030150 | -1,788 | 0,0241 |
| Sspon.07G0030220 | -1,153 | 0,0242 |
| Sspon.07G0030240 | 2,652 | 0,0242 |
| Sspon.07G0030440 | -1,731 | 0,0243 |
| Sspon.07G0030580 | 1,802 | 0,0243 |
| Sspon.07G0030590 | -2,133 | 0,0244 |
| Sspon.07G0030680 | 1,443 | 0,0244 |
| Sspon.07G0030790 | 2,335 | 0,0244 |
| Sspon.07G0030890 | 1,031 | 0,0244 |
| Sspon.07G0031110 | -1,418 | 0,0245 |
| Sspon.07G0031220 | -3,580 | 0,0245 |
| Sspon.07G0031250 | -5,246 | 0,0245 |
| Sspon.07G0031470 | 4,249 | 0,0246 |
| Sspon.07G0031490 | -1,076 | 0,0246 |
| Sspon.07G0031590 | 1,323 | 0,0246 |
| Sspon.07G0031600 | -2,747 | 0,0246 |
| Sspon.07G0031650 | 3,228 | 0,0246 |
| Sspon.07G0031660 | -6,013 | 0,0246 |
| Sspon.07G0031750 | 5,138 | 0,0247 |
| Sspon.07G0031780 | 1,053 | 0,0248 |
| Sspon.07G0031800 | 1,064 | 0,0249 |
| Sspon.07G0031870 | 13,982 | 0,0249 |
| Sspon.07G0031930 | 1,927 | 0,0249 |
| Sspon.07G0031940 | -1,835 | 0,0249 |
| Sspon.07G0031950 | 3,134 | 0,0249 |
| Sspon.07G0031960 | -5,676 | 0,0249 |
| Sspon.07G0031980 | -5,638 | 0,0250 |
| Sspon.07G0031990 | 1,145 | 0,0250 |

| | | |
|---|---|---|
| Sspon.07G0032220 | 1,073 | 0,0250 |
| Sspon.07G0032380 | -1,145 | 0,0250 |
| Sspon.07G0032490 | -1,282 | 0,0250 |
| Sspon.07G0032500 | -1,656 | 0,0251 |
| Sspon.07G0032570 | 1,224 | 0,0251 |
| Sspon.07G0032680 | -2,723 | 0,0252 |
| Sspon.07G0032770 | 5,684 | 0,0252 |
| Sspon.07G0032800 | 1,035 | 0,0252 |
| Sspon.07G0032830 | -1,631 | 0,0252 |
| Sspon.07G0032870 | 5,047 | 0,0253 |
| Sspon.07G0032930 | -1,790 | 0,0253 |
| Sspon.07G0032960 | 2,540 | 0,0253 |
| Sspon.07G0033000 | 1,001 | 0,0254 |
| Sspon.07G0033030 | 1,757 | 0,0254 |
| Sspon.07G0033210 | 1,036 | 0,0254 |
| Sspon.07G0033230 | -5,409 | 0,0254 |
| Sspon.07G0033240 | 1,054 | 0,0254 |
| Sspon.07G0033270 | 2,127 | 0,0254 |
| Sspon.07G0033350 | -1,001 | 0,0255 |
| Sspon.07G0033440 | -2,096 | 0,0255 |
| Sspon.07G0033510 | 1,316 | 0,0255 |
| Sspon.07G0033530 | -1,621 | 0,0255 |
| Sspon.07G0033560 | 5,264 | 0,0256 |
| Sspon.07G0033590 | 2,365 | 0,0256 |
| Sspon.07G0033620 | 6,129 | 0,0256 |
| Sspon.07G0033660 | -4,331 | 0,0256 |
| Sspon.07G0033740 | 6,641 | 0,0256 |
| Sspon.07G0033820 | -5,418 | 0,0256 |
| Sspon.07G0033870 | 1,100 | 0,0257 |
| Sspon.07G0033880 | -1,149 | 0,0257 |
| Sspon.07G0033980 | -4,642 | 0,0258 |
| Sspon.07G0034020 | -1,431 | 0,0258 |
| Sspon.07G0034030 | 2,441 | 0,0258 |
| Sspon.07G0034050 | -3,659 | 0,0258 |
| Sspon.07G0034150 | -4,779 | 0,0258 |
| Sspon.07G0034190 | -5,441 | 0,0259 |
| Sspon.07G0034200 | 1,024 | 0,0260 |
| Sspon.07G0034270 | -3,563 | 0,0260 |
| Sspon.07G0034300 | 2,242 | 0,0260 |
| Sspon.07G0034330 | -1,726 | 0,0260 |
| Sspon.07G0034440 | -4,724 | 0,0261 |
| Sspon.07G0034530 | -2,571 | 0,0261 |
| Sspon.07G0034650 | -2,731 | 0,0261 |

| | | |
|---|---|---|
| Sspon.07G0034670 | 1,383 | 0,0261 |
| Sspon.07G0034690 | 3,034 | 0,0262 |
| Sspon.07G0034740 | -4,531 | 0,0262 |
| Sspon.07G0034790 | -1,593 | 0,0262 |
| Sspon.07G0035020 | 1,104 | 0,0262 |
| Sspon.07G0035090 | 3,953 | 0,0263 |
| Sspon.07G0035130 | 1,141 | 0,0264 |
| Sspon.07G0035150 | -2,187 | 0,0264 |
| Sspon.07G0035200 | -5,443 | 0,0264 |
| Sspon.07G0035320 | 1,157 | 0,0265 |
| Sspon.07G0035340 | 1,381 | 0,0265 |
| Sspon.07G0035650 | -3,761 | 0,0265 |
| Sspon.07G0035750 | -3,554 | 0,0266 |
| Sspon.07G0035770 | -1,040 | 0,0266 |
| Sspon.07G0035810 | 3,018 | 0,0266 |
| Sspon.07G0035820 | 1,104 | 0,0266 |
| Sspon.07G0035860 | 2,272 | 0,0266 |
| Sspon.07G0035900 | -1,513 | 0,0267 |
| Sspon.07G0035980 | 14,222 | 0,0267 |
| Sspon.07G0036150 | 3,091 | 0,0267 |
| Sspon.07G0036250 | -2,492 | 0,0267 |
| Sspon.07G0036290 | 1,701 | 0,0268 |
| Sspon.07G0036310 | -2,284 | 0,0269 |
| Sspon.07G0036440 | 3,660 | 0,0269 |
| Sspon.07G0036660 | 1,045 | 0,0269 |
| Sspon.07G0037060 | 1,188 | 0,0270 |
| Sspon.07G0037110 | -4,047 | 0,0272 |
| Sspon.07G0037120 | 1,739 | 0,0273 |
| Sspon.07G0037150 | -5,579 | 0,0273 |
| Sspon.07G0037160 | -3,217 | 0,0273 |
| Sspon.07G0037380 | -5,327 | 0,0274 |
| Sspon.07G0037400 | 2,239 | 0,0274 |
| Sspon.07G0037530 | -3,808 | 0,0274 |
| Sspon.07G0037650 | 1,440 | 0,0274 |
| Sspon.07G0037710 | -3,242 | 0,0274 |
| Sspon.07G0037770 | 3,056 | 0,0274 |
| Sspon.07G0037910 | 7,181 | 0,0274 |
| Sspon.07G0038040 | 2,272 | 0,0275 |
| Sspon.07G0038130 | -2,942 | 0,0275 |
| Sspon.07G0038140 | 2,781 | 0,0275 |
| Sspon.07G0038400 | 4,645 | 0,0275 |
| Sspon.07G0038460 | 6,616 | 0,0275 |
| Sspon.07G0038540 | 4,245 | 0,0276 |

| | | |
|---|---|---|
| Sspon.07G0038550 | 1,489 | 0,0276 |
| Sspon.07G0038560 | 1,418 | 0,0277 |
| Sspon.07G0038590 | 1,590 | 0,0278 |
| Sspon.07G0038720 | 1,857 | 0,0279 |
| Sspon.08G0000210 | 1,075 | 0,0279 |
| Sspon.08G0000240 | 5,153 | 0,0279 |
| Sspon.08G0000330 | 3,969 | 0,0280 |
| Sspon.08G0000370 | 2,219 | 0,0280 |
| Sspon.08G0000380 | -1,655 | 0,0280 |
| Sspon.08G0000540 | 3,081 | 0,0280 |
| Sspon.08G0000550 | 1,056 | 0,0281 |
| Sspon.08G0000580 | 1,283 | 0,0281 |
| Sspon.08G0000810 | 1,793 | 0,0282 |
| Sspon.08G0000860 | -4,054 | 0,0282 |
| Sspon.08G0000870 | -4,759 | 0,0282 |
| Sspon.08G0000930 | 1,260 | 0,0283 |
| Sspon.08G0000960 | 4,197 | 0,0283 |
| Sspon.08G0001060 | 1,035 | 0,0284 |
| Sspon.08G0001270 | -4,411 | 0,0284 |
| Sspon.08G0001290 | -2,239 | 0,0285 |
| Sspon.08G0001680 | 4,236 | 0,0285 |
| Sspon.08G0001750 | 3,096 | 0,0285 |
| Sspon.08G0001800 | 4,810 | 0,0285 |
| Sspon.08G0001900 | 1,327 | 0,0285 |
| Sspon.08G0001930 | 6,355 | 0,0285 |
| Sspon.08G0002150 | 2,991 | 0,0285 |
| Sspon.08G0002170 | -5,085 | 0,0285 |
| Sspon.08G0002230 | 2,502 | 0,0286 |
| Sspon.08G0002240 | -1,113 | 0,0286 |
| Sspon.08G0002280 | 2,054 | 0,0286 |
| Sspon.08G0002290 | -1,848 | 0,0286 |
| Sspon.08G0002300 | 7,370 | 0,0287 |
| Sspon.08G0002350 | 1,585 | 0,0287 |
| Sspon.08G0002380 | -4,979 | 0,0288 |
| Sspon.08G0002400 | 1,313 | 0,0289 |
| Sspon.08G0002450 | 1,092 | 0,0289 |
| Sspon.08G0002580 | -4,452 | 0,0289 |
| Sspon.08G0002610 | 3,370 | 0,0290 |
| Sspon.08G0002650 | 3,031 | 0,0290 |
| Sspon.08G0002660 | -1,020 | 0,0290 |
| Sspon.08G0002720 | -1,556 | 0,0291 |
| Sspon.08G0002890 | -4,837 | 0,0291 |
| Sspon.08G0002950 | -2,088 | 0,0292 |

| | | |
|---|---|---|
| Sspon.08G0003030 | 1,905 | 0,0292 |
| Sspon.08G0003130 | -2,558 | 0,0292 |
| Sspon.08G0003200 | 1,632 | 0,0293 |
| Sspon.08G0003210 | -2,532 | 0,0293 |
| Sspon.08G0003260 | -1,119 | 0,0293 |
| Sspon.08G0003320 | -3,769 | 0,0294 |
| Sspon.08G0003390 | -1,602 | 0,0294 |
| Sspon.08G0003480 | -4,491 | 0,0295 |
| Sspon.08G0003490 | 3,119 | 0,0296 |
| Sspon.08G0003500 | -3,911 | 0,0297 |
| Sspon.08G0003690 | -5,873 | 0,0297 |
| Sspon.08G0003750 | -2,666 | 0,0298 |
| Sspon.08G0003780 | -4,226 | 0,0298 |
| Sspon.08G0003790 | 3,639 | 0,0298 |
| Sspon.08G0003890 | 5,980 | 0,0298 |
| Sspon.08G0003980 | -1,565 | 0,0299 |
| Sspon.08G0004020 | -1,605 | 0,0299 |
| Sspon.08G0004040 | -4,464 | 0,0299 |
| Sspon.08G0004050 | -2,373 | 0,0300 |
| Sspon.08G0004100 | -1,679 | 0,0300 |
| Sspon.08G0004140 | 8,383 | 0,0300 |
| Sspon.08G0004150 | 6,915 | 0,0300 |
| Sspon.08G0004160 | 1,831 | 0,0301 |
| Sspon.08G0004260 | -2,491 | 0,0301 |
| Sspon.08G0004360 | 1,014 | 0,0301 |
| Sspon.08G0004370 | -5,261 | 0,0301 |
| Sspon.08G0004390 | 1,411 | 0,0301 |
| Sspon.08G0004570 | -1,033 | 0,0301 |
| Sspon.08G0004600 | 1,316 | 0,0301 |
| Sspon.08G0004760 | -2,469 | 0,0302 |
| Sspon.08G0004780 | -3,692 | 0,0302 |
| Sspon.08G0004820 | -1,108 | 0,0302 |
| Sspon.08G0004840 | 4,591 | 0,0302 |
| Sspon.08G0004880 | 5,410 | 0,0302 |
| Sspon.08G0004930 | -3,262 | 0,0302 |
| Sspon.08G0004950 | -5,127 | 0,0303 |
| Sspon.08G0005000 | 1,640 | 0,0304 |
| Sspon.08G0005010 | 6,130 | 0,0304 |
| Sspon.08G0005030 | 2,628 | 0,0304 |
| Sspon.08G0005110 | -1,808 | 0,0304 |
| Sspon.08G0005130 | 2,187 | 0,0304 |
| Sspon.08G0005140 | 1,451 | 0,0305 |
| Sspon.08G0005220 | 3,603 | 0,0306 |

| | | |
|---|---|---|
| Sspon.08G0005250 | -1,299 | 0,0306 |
| Sspon.08G0005330 | -1,108 | 0,0307 |
| Sspon.08G0005420 | -1,047 | 0,0307 |
| Sspon.08G0005450 | 5,403 | 0,0307 |
| Sspon.08G0005470 | -1,633 | 0,0307 |
| Sspon.08G0005550 | -4,440 | 0,0307 |
| Sspon.08G0005600 | 4,874 | 0,0308 |
| Sspon.08G0005620 | -1,610 | 0,0308 |
| Sspon.08G0005630 | 5,651 | 0,0308 |
| Sspon.08G0005690 | 3,902 | 0,0308 |
| Sspon.08G0005700 | 1,240 | 0,0309 |
| Sspon.08G0005750 | 6,910 | 0,0309 |
| Sspon.08G0005830 | 3,615 | 0,0310 |
| Sspon.08G0005860 | 6,552 | 0,0310 |
| Sspon.08G0005960 | 2,567 | 0,0310 |
| Sspon.08G0006210 | 1,301 | 0,0311 |
| Sspon.08G0006250 | 2,319 | 0,0311 |
| Sspon.08G0006260 | 4,165 | 0,0311 |
| Sspon.08G0006500 | 1,949 | 0,0311 |
| Sspon.08G0006570 | -5,263 | 0,0311 |
| Sspon.08G0006630 | 1,311 | 0,0311 |
| Sspon.08G0006640 | 1,608 | 0,0311 |
| Sspon.08G0006650 | -3,966 | 0,0311 |
| Sspon.08G0006660 | 3,379 | 0,0312 |
| Sspon.08G0006720 | -3,338 | 0,0312 |
| Sspon.08G0006810 | 6,599 | 0,0312 |
| Sspon.08G0006830 | -3,440 | 0,0312 |
| Sspon.08G0006850 | 1,098 | 0,0313 |
| Sspon.08G0006920 | 1,291 | 0,0314 |
| Sspon.08G0006950 | 6,968 | 0,0314 |
| Sspon.08G0006970 | 3,385 | 0,0314 |
| Sspon.08G0007010 | 2,996 | 0,0316 |
| Sspon.08G0007080 | -1,096 | 0,0317 |
| Sspon.08G0007130 | -2,234 | 0,0317 |
| Sspon.08G0007140 | -4,185 | 0,0317 |
| Sspon.08G0007150 | -1,805 | 0,0317 |
| Sspon.08G0007210 | 2,094 | 0,0317 |
| Sspon.08G0007270 | -2,604 | 0,0317 |
| Sspon.08G0007350 | 2,110 | 0,0317 |
| Sspon.08G0007360 | -5,871 | 0,0317 |
| Sspon.08G0007450 | -1,039 | 0,0318 |
| Sspon.08G0007480 | -3,416 | 0,0318 |
| Sspon.08G0007560 | 5,469 | 0,0318 |

| | | |
|---|---|---|
| Sspon.08G0007590 | 6,208 | 0,0319 |
| Sspon.08G0007730 | 1,171 | 0,0319 |
| Sspon.08G0007790 | -4,318 | 0,0322 |
| Sspon.08G0007840 | 2,927 | 0,0323 |
| Sspon.08G0007860 | 2,261 | 0,0323 |
| Sspon.08G0007890 | 1,638 | 0,0325 |
| Sspon.08G0008020 | 2,064 | 0,0325 |
| Sspon.08G0008080 | -2,067 | 0,0326 |
| Sspon.08G0008190 | -3,031 | 0,0326 |
| Sspon.08G0008200 | -1,370 | 0,0327 |
| Sspon.08G0008340 | -4,600 | 0,0327 |
| Sspon.08G0008350 | 6,036 | 0,0328 |
| Sspon.08G0008370 | -5,697 | 0,0328 |
| Sspon.08G0008390 | 2,151 | 0,0328 |
| Sspon.08G0008510 | 2,455 | 0,0329 |
| Sspon.08G0008520 | 4,888 | 0,0329 |
| Sspon.08G0008560 | 1,007 | 0,0329 |
| Sspon.08G0008570 | 2,183 | 0,0330 |
| Sspon.08G0008580 | 1,230 | 0,0330 |
| Sspon.08G0008620 | -3,384 | 0,0331 |
| Sspon.08G0008640 | -1,185 | 0,0331 |
| Sspon.08G0008690 | -1,522 | 0,0331 |
| Sspon.08G0008730 | 1,961 | 0,0331 |
| Sspon.08G0008930 | 1,301 | 0,0332 |
| Sspon.08G0008950 | 1,189 | 0,0332 |
| Sspon.08G0008960 | 2,421 | 0,0332 |
| Sspon.08G0009040 | 3,502 | 0,0332 |
| Sspon.08G0009060 | 6,306 | 0,0333 |
| Sspon.08G0009110 | 1,417 | 0,0333 |
| Sspon.08G0009250 | 1,607 | 0,0333 |
| Sspon.08G0009260 | 2,833 | 0,0333 |
| Sspon.08G0009320 | -2,280 | 0,0333 |
| Sspon.08G0009350 | 1,421 | 0,0334 |
| Sspon.08G0009410 | -1,226 | 0,0334 |
| Sspon.08G0009430 | 5,846 | 0,0334 |
| Sspon.08G0009450 | 1,769 | 0,0335 |
| Sspon.08G0009590 | 4,024 | 0,0335 |
| Sspon.08G0009620 | 1,283 | 0,0335 |
| Sspon.08G0009650 | 3,667 | 0,0335 |
| Sspon.08G0009680 | 1,031 | 0,0335 |
| Sspon.08G0009760 | 1,349 | 0,0335 |
| Sspon.08G0009790 | 5,815 | 0,0335 |
| Sspon.08G0009920 | -5,250 | 0,0336 |

| | | |
|---|---|---|
| Sspon.08G0009930 | 1,610 | 0,0337 |
| Sspon.08G0009940 | 1,899 | 0,0338 |
| Sspon.08G0010020 | 4,252 | 0,0338 |
| Sspon.08G0010040 | 6,287 | 0,0338 |
| Sspon.08G0010130 | -2,139 | 0,0338 |
| Sspon.08G0010170 | 6,281 | 0,0338 |
| Sspon.08G0010210 | -2,008 | 0,0338 |
| Sspon.08G0010270 | 5,162 | 0,0339 |
| Sspon.08G0010430 | -2,720 | 0,0339 |
| Sspon.08G0010440 | -4,324 | 0,0339 |
| Sspon.08G0010460 | -1,481 | 0,0341 |
| Sspon.08G0010570 | -1,605 | 0,0341 |
| Sspon.08G0010830 | -2,584 | 0,0342 |
| Sspon.08G0010900 | -1,576 | 0,0343 |
| Sspon.08G0011070 | -4,420 | 0,0343 |
| Sspon.08G0011110 | 1,286 | 0,0343 |
| Sspon.08G0011150 | 1,010 | 0,0344 |
| Sspon.08G0011160 | -3,910 | 0,0344 |
| Sspon.08G0011190 | 2,095 | 0,0344 |
| Sspon.08G0011210 | 3,009 | 0,0345 |
| Sspon.08G0011400 | 1,154 | 0,0345 |
| Sspon.08G0011410 | -3,739 | 0,0345 |
| Sspon.08G0011480 | -2,786 | 0,0346 |
| Sspon.08G0011490 | 1,348 | 0,0346 |
| Sspon.08G0011700 | -1,999 | 0,0346 |
| Sspon.08G0011710 | 2,638 | 0,0346 |
| Sspon.08G0011720 | -1,741 | 0,0347 |
| Sspon.08G0011750 | 1,914 | 0,0347 |
| Sspon.08G0011760 | 2,660 | 0,0347 |
| Sspon.08G0011770 | 2,738 | 0,0347 |
| Sspon.08G0011810 | 1,511 | 0,0348 |
| Sspon.08G0012130 | 4,232 | 0,0348 |
| Sspon.08G0012140 | -5,970 | 0,0349 |
| Sspon.08G0012170 | 1,146 | 0,0349 |
| Sspon.08G0012330 | 1,610 | 0,0349 |
| Sspon.08G0012340 | 3,692 | 0,0350 |
| Sspon.08G0012410 | -2,248 | 0,0350 |
| Sspon.08G0012500 | 2,094 | 0,0351 |
| Sspon.08G0012600 | -1,386 | 0,0352 |
| Sspon.08G0012780 | -1,096 | 0,0352 |
| Sspon.08G0012790 | 1,067 | 0,0353 |
| Sspon.08G0012830 | -2,084 | 0,0353 |
| Sspon.08G0012920 | 5,714 | 0,0354 |

| | | |
|---|---|---|
| Sspon.08G0012930 | -1,665 | 0,0354 |
| Sspon.08G0013020 | 1,277 | 0,0355 |
| Sspon.08G0013090 | 4,214 | 0,0355 |
| Sspon.08G0013150 | -1,044 | 0,0355 |
| Sspon.08G0013160 | -3,561 | 0,0355 |
| Sspon.08G0013210 | 1,711 | 0,0356 |
| Sspon.08G0013220 | -4,332 | 0,0356 |
| Sspon.08G0013270 | -1,837 | 0,0356 |
| Sspon.08G0013320 | -4,537 | 0,0358 |
| Sspon.08G0013330 | -3,879 | 0,0358 |
| Sspon.08G0013400 | 3,515 | 0,0358 |
| Sspon.08G0013420 | -2,435 | 0,0360 |
| Sspon.08G0013440 | 1,158 | 0,0360 |
| Sspon.08G0013450 | -3,313 | 0,0360 |
| Sspon.08G0013460 | 2,886 | 0,0363 |
| Sspon.08G0013500 | 3,131 | 0,0363 |
| Sspon.08G0013590 | -4,519 | 0,0364 |
| Sspon.08G0013690 | 3,456 | 0,0364 |
| Sspon.08G0013720 | -1,925 | 0,0366 |
| Sspon.08G0013810 | -1,241 | 0,0367 |
| Sspon.08G0013960 | -2,999 | 0,0367 |
| Sspon.08G0013970 | -3,873 | 0,0367 |
| Sspon.08G0014160 | 1,266 | 0,0367 |
| Sspon.08G0014290 | 4,052 | 0,0368 |
| Sspon.08G0014350 | 1,611 | 0,0368 |
| Sspon.08G0014470 | 2,764 | 0,0369 |
| Sspon.08G0014530 | 5,203 | 0,0369 |
| Sspon.08G0014650 | 1,876 | 0,0369 |
| Sspon.08G0014670 | 2,704 | 0,0369 |
| Sspon.08G0014710 | -2,014 | 0,0369 |
| Sspon.08G0014720 | -1,500 | 0,0370 |
| Sspon.08G0014740 | -4,296 | 0,0370 |
| Sspon.08G0014920 | 2,092 | 0,0370 |
| Sspon.08G0015090 | -1,024 | 0,0370 |
| Sspon.08G0015360 | -1,764 | 0,0371 |
| Sspon.08G0015380 | -1,559 | 0,0371 |
| Sspon.08G0015430 | 7,054 | 0,0372 |
| Sspon.08G0015550 | -2,689 | 0,0372 |
| Sspon.08G0015560 | 3,452 | 0,0373 |
| Sspon.08G0015590 | 2,808 | 0,0373 |
| Sspon.08G0015600 | 1,902 | 0,0374 |
| Sspon.08G0015660 | -1,202 | 0,0374 |
| Sspon.08G0015710 | 4,866 | 0,0375 |

| | | |
|---|---|---|
| Sspon.08G0015730 | -2,226 | 0,0375 |
| Sspon.08G0015750 | -1,957 | 0,0375 |
| Sspon.08G0015770 | -4,051 | 0,0375 |
| Sspon.08G0016030 | -2,799 | 0,0375 |
| Sspon.08G0016150 | -3,323 | 0,0376 |
| Sspon.08G0016170 | 6,691 | 0,0376 |
| Sspon.08G0016210 | -1,782 | 0,0376 |
| Sspon.08G0016220 | -1,608 | 0,0376 |
| Sspon.08G0016260 | -3,527 | 0,0377 |
| Sspon.08G0016300 | 4,784 | 0,0377 |
| Sspon.08G0016310 | 1,096 | 0,0377 |
| Sspon.08G0016330 | -3,300 | 0,0377 |
| Sspon.08G0016370 | 1,161 | 0,0377 |
| Sspon.08G0016380 | 1,993 | 0,0378 |
| Sspon.08G0016400 | 5,119 | 0,0378 |
| Sspon.08G0016420 | -1,693 | 0,0378 |
| Sspon.08G0016550 | -3,568 | 0,0378 |
| Sspon.08G0016560 | -1,145 | 0,0379 |
| Sspon.08G0016590 | 1,072 | 0,0379 |
| Sspon.08G0016680 | 2,149 | 0,0380 |
| Sspon.08G0016730 | 2,061 | 0,0380 |
| Sspon.08G0016770 | 1,036 | 0,0380 |
| Sspon.08G0016790 | -2,392 | 0,0381 |
| Sspon.08G0016840 | 1,682 | 0,0381 |
| Sspon.08G0016850 | -1,231 | 0,0382 |
| Sspon.08G0016890 | 1,110 | 0,0383 |
| Sspon.08G0016930 | -3,744 | 0,0383 |
| Sspon.08G0017010 | -1,402 | 0,0383 |
| Sspon.08G0017110 | -1,521 | 0,0384 |
| Sspon.08G0017190 | -1,000 | 0,0384 |
| Sspon.08G0017220 | -1,489 | 0,0384 |
| Sspon.08G0017290 | 1,156 | 0,0384 |
| Sspon.08G0017340 | 2,742 | 0,0385 |
| Sspon.08G0017630 | -1,696 | 0,0385 |
| Sspon.08G0017680 | 1,209 | 0,0386 |
| Sspon.08G0017740 | 3,720 | 0,0387 |
| Sspon.08G0017810 | -1,717 | 0,0388 |
| Sspon.08G0017920 | -4,791 | 0,0389 |
| Sspon.08G0017970 | 1,964 | 0,0389 |
| Sspon.08G0018010 | 4,781 | 0,0389 |
| Sspon.08G0018100 | 1,214 | 0,0389 |
| Sspon.08G0018110 | -3,543 | 0,0390 |
| Sspon.08G0018170 | -5,374 | 0,0391 |

| | | |
|---|---|---|
| Sspon.08G0018350 | -3,741 | 0,0392 |
| Sspon.08G0018410 | 1,046 | 0,0392 |
| Sspon.08G0018500 | 3,191 | 0,0392 |
| Sspon.08G0018530 | -1,539 | 0,0392 |
| Sspon.08G0018570 | -1,590 | 0,0393 |
| Sspon.08G0018600 | 1,241 | 0,0393 |
| Sspon.08G0018620 | -1,089 | 0,0395 |
| Sspon.08G0018670 | 4,042 | 0,0398 |
| Sspon.08G0018700 | -1,712 | 0,0398 |
| Sspon.08G0018760 | 3,351 | 0,0399 |
| Sspon.08G0018900 | -1,461 | 0,0400 |
| Sspon.08G0018920 | -2,739 | 0,0400 |
| Sspon.08G0018940 | 2,075 | 0,0400 |
| Sspon.08G0018980 | -1,015 | 0,0401 |
| Sspon.08G0019040 | 2,113 | 0,0402 |
| Sspon.08G0019090 | 2,032 | 0,0403 |
| Sspon.08G0019140 | -1,270 | 0,0403 |
| Sspon.08G0019180 | 5,159 | 0,0403 |
| Sspon.08G0019220 | -2,581 | 0,0403 |
| Sspon.08G0019230 | -2,082 | 0,0404 |
| Sspon.08G0019260 | -3,410 | 0,0404 |
| Sspon.08G0019310 | -5,261 | 0,0404 |
| Sspon.08G0019360 | -4,216 | 0,0404 |
| Sspon.08G0019370 | 1,857 | 0,0404 |
| Sspon.08G0019380 | 1,687 | 0,0405 |
| Sspon.08G0019440 | 1,534 | 0,0405 |
| Sspon.08G0019450 | -1,136 | 0,0405 |
| Sspon.08G0019590 | 2,991 | 0,0405 |
| Sspon.08G0019660 | -4,813 | 0,0405 |
| Sspon.08G0019690 | 2,750 | 0,0406 |
| Sspon.08G0019750 | 2,545 | 0,0406 |
| Sspon.08G0019770 | -4,186 | 0,0406 |
| Sspon.08G0019940 | -1,964 | 0,0407 |
| Sspon.08G0020050 | 7,412 | 0,0407 |
| Sspon.08G0020070 | 1,913 | 0,0408 |
| Sspon.08G0020200 | -3,023 | 0,0408 |
| Sspon.08G0020250 | -5,017 | 0,0409 |
| Sspon.08G0020490 | -3,485 | 0,0409 |
| Sspon.08G0020510 | 6,216 | 0,0410 |
| Sspon.08G0020640 | 2,062 | 0,0410 |
| Sspon.08G0020790 | 1,494 | 0,0410 |
| Sspon.08G0020800 | -3,926 | 0,0410 |
| Sspon.08G0020830 | 1,966 | 0,0410 |

| | | |
|---|---|---|
| Sspon.08G0020840 | 4,875 | 0,0411 |
| Sspon.08G0020940 | -1,644 | 0,0412 |
| Sspon.08G0020990 | -1,419 | 0,0412 |
| Sspon.08G0021010 | 1,863 | 0,0412 |
| Sspon.08G0021020 | 4,050 | 0,0413 |
| Sspon.08G0021080 | 4,690 | 0,0414 |
| Sspon.08G0021090 | -1,671 | 0,0415 |
| Sspon.08G0021220 | -3,797 | 0,0416 |
| Sspon.08G0021250 | 7,464 | 0,0416 |
| Sspon.08G0021490 | 2,366 | 0,0417 |
| Sspon.08G0021590 | 1,058 | 0,0419 |
| Sspon.08G0021600 | 4,617 | 0,0419 |
| Sspon.08G0021630 | 2,128 | 0,0420 |
| Sspon.08G0021640 | 1,328 | 0,0421 |
| Sspon.08G0021660 | 2,279 | 0,0423 |
| Sspon.08G0021730 | 3,336 | 0,0423 |
| Sspon.08G0021780 | -3,103 | 0,0423 |
| Sspon.08G0021810 | -3,681 | 0,0423 |
| Sspon.08G0021820 | 1,277 | 0,0423 |
| Sspon.08G0021840 | -2,716 | 0,0424 |
| Sspon.08G0021950 | 4,444 | 0,0425 |
| Sspon.08G0022010 | -1,889 | 0,0425 |
| Sspon.08G0022270 | 1,704 | 0,0425 |
| Sspon.08G0022290 | -4,970 | 0,0426 |
| Sspon.08G0022300 | 3,305 | 0,0426 |
| Sspon.08G0022410 | -1,600 | 0,0427 |
| Sspon.08G0022440 | 4,506 | 0,0428 |
| Sspon.08G0022490 | 2,271 | 0,0429 |
| Sspon.08G0022550 | 1,926 | 0,0430 |
| Sspon.08G0022620 | 4,129 | 0,0431 |
| Sspon.08G0022650 | -1,068 | 0,0432 |
| Sspon.08G0022770 | 2,213 | 0,0432 |
| Sspon.08G0022800 | -4,007 | 0,0432 |
| Sspon.08G0022930 | -3,348 | 0,0432 |
| Sspon.08G0022950 | -1,651 | 0,0432 |
| Sspon.08G0022960 | -1,690 | 0,0434 |
| Sspon.08G0022990 | -1,697 | 0,0435 |
| Sspon.08G0023020 | -1,698 | 0,0436 |
| Sspon.08G0023030 | -3,214 | 0,0436 |
| Sspon.08G0023050 | -4,908 | 0,0436 |
| Sspon.08G0023060 | -3,701 | 0,0438 |
| Sspon.08G0023110 | 2,658 | 0,0438 |
| Sspon.08G0023170 | 4,467 | 0,0438 |

| | | |
|---|---|---|
| Sspon.08G0023230 | -3,427 | 0,0439 |
| Sspon.08G0023390 | 2,699 | 0,0440 |
| Sspon.08G0023410 | 2,399 | 0,0440 |
| Sspon.08G0023600 | -4,403 | 0,0442 |
| Sspon.08G0023730 | -1,352 | 0,0442 |
| Sspon.08G0023780 | -3,988 | 0,0442 |
| Sspon.08G0023800 | -1,511 | 0,0442 |
| Sspon.08G0023890 | 5,885 | 0,0442 |
| Sspon.08G0024020 | 4,601 | 0,0442 |
| Sspon.08G0024140 | 2,527 | 0,0443 |
| Sspon.08G0024160 | 1,805 | 0,0443 |
| Sspon.08G0024280 | 3,449 | 0,0444 |
| Sspon.08G0024290 | 1,947 | 0,0446 |
| Sspon.08G0024320 | 1,539 | 0,0446 |
| Sspon.08G0024370 | -3,721 | 0,0447 |
| Sspon.08G0024400 | -1,352 | 0,0447 |
| Sspon.08G0024420 | 2,691 | 0,0448 |
| Sspon.08G0024540 | 1,240 | 0,0448 |
| Sspon.08G0024630 | -4,024 | 0,0448 |
| Sspon.08G0024650 | -2,217 | 0,0449 |
| Sspon.08G0024660 | -3,869 | 0,0449 |
| Sspon.08G0024680 | -4,369 | 0,0449 |
| Sspon.08G0024720 | -1,159 | 0,0449 |
| Sspon.08G0024780 | 6,210 | 0,0450 |
| Sspon.08G0024790 | -3,808 | 0,0450 |
| Sspon.08G0024900 | -1,417 | 0,0451 |
| Sspon.08G0024940 | 3,940 | 0,0452 |
| Sspon.08G0024990 | -2,343 | 0,0452 |
| Sspon.08G0025120 | 2,434 | 0,0452 |
| Sspon.08G0025150 | 1,652 | 0,0453 |
| Sspon.08G0025170 | 1,720 | 0,0453 |
| Sspon.08G0025210 | -2,986 | 0,0454 |
| Sspon.08G0025220 | 2,169 | 0,0454 |
| Sspon.08G0025260 | 2,368 | 0,0454 |
| Sspon.08G0025300 | -2,140 | 0,0455 |
| Sspon.08G0025360 | 1,055 | 0,0457 |
| Sspon.08G0025400 | -1,636 | 0,0457 |
| Sspon.08G0025460 | -3,256 | 0,0457 |
| Sspon.08G0025600 | 2,871 | 0,0458 |
| Sspon.08G0025680 | -3,841 | 0,0458 |
| Sspon.08G0025800 | -4,274 | 0,0459 |
| Sspon.08G0025910 | -2,987 | 0,0459 |
| Sspon.08G0026020 | 2,833 | 0,0459 |

| | | |
|---|---|---|
| Sspon.08G0026030 | -1,494 | 0,0460 |
| Sspon.08G0026110 | 1,493 | 0,0460 |
| Sspon.08G0026220 | -4,726 | 0,0460 |
| Sspon.08G0026310 | -1,430 | 0,0460 |
| Sspon.08G0026320 | -1,860 | 0,0461 |
| Sspon.08G0026330 | 1,052 | 0,0462 |
| Sspon.08G0026340 | -5,412 | 0,0462 |
| Sspon.08G0026400 | -2,185 | 0,0462 |
| Sspon.08G0026420 | 1,650 | 0,0463 |
| Sspon.08G0026460 | 1,387 | 0,0464 |
| Sspon.08G0026530 | 1,866 | 0,0464 |
| Sspon.08G0026600 | -2,889 | 0,0464 |
| Sspon.08G0026640 | 4,321 | 0,0465 |
| Sspon.08G0026660 | -1,834 | 0,0465 |
| Sspon.08G0026670 | 1,575 | 0,0466 |
| Sspon.08G0026890 | 2,531 | 0,0466 |
| Sspon.08G0026980 | -1,906 | 0,0467 |
| Sspon.08G0027030 | -2,948 | 0,0470 |
| Sspon.08G0027110 | 1,290 | 0,0470 |
| Sspon.08G0027200 | 1,040 | 0,0470 |
| Sspon.08G0027260 | -1,105 | 0,0470 |
| Sspon.08G0027270 | -3,962 | 0,0471 |
| Sspon.08G0027310 | 4,716 | 0,0473 |
| Sspon.08G0027330 | -4,486 | 0,0474 |
| Sspon.08G0027470 | -2,722 | 0,0474 |
| Sspon.08G0027480 | 1,787 | 0,0475 |
| Sspon.08G0027510 | 2,498 | 0,0475 |
| Sspon.08G0027530 | -1,075 | 0,0475 |
| Sspon.08G0027540 | 4,409 | 0,0475 |
| Sspon.08G0027650 | -1,174 | 0,0477 |
| Sspon.08G0027680 | 2,126 | 0,0477 |
| Sspon.08G0027750 | 3,910 | 0,0478 |
| Sspon.08G0027820 | 4,856 | 0,0478 |
| Sspon.08G0027850 | 2,242 | 0,0479 |
| Sspon.08G0027890 | 1,019 | 0,0480 |
| Sspon.08G0027900 | 2,711 | 0,0480 |
| Sspon.08G0027910 | -1,580 | 0,0480 |
| Sspon.08G0028190 | 7,449 | 0,0481 |
| Sspon.08G0028330 | -3,409 | 0,0481 |
| Sspon.08G0028340 | 1,730 | 0,0483 |
| Sspon.08G0028440 | -3,818 | 0,0483 |
| Sspon.08G0028630 | -1,163 | 0,0484 |
| Sspon.08G0028790 | 4,507 | 0,0484 |

| | | |
|---|---|---|
| Sspon.08G0028950 | 1,927 | 0,0484 |
| Sspon.08G0028980 | -3,417 | 0,0484 |
| Sspon.08G0029100 | 1,630 | 0,0484 |
| Sspon.08G0029230 | 1,295 | 0,0485 |
| Sspon.08G0029240 | -5,357 | 0,0485 |
| Sspon.08G0029320 | 1,203 | 0,0486 |
| Sspon.08G0029470 | -2,903 | 0,0486 |
| Sspon.08G0029490 | 1,176 | 0,0487 |
| Sspon.08G0029550 | -3,394 | 0,0489 |
| Sspon.08G0029560 | 4,792 | 0,0489 |
| Sspon.08G0029590 | -3,137 | 0,0489 |
| Sspon.08G0029740 | 3,374 | 0,0489 |
| Sspon.08G0029760 | -2,247 | 0,0490 |
| Sspon.08G0029810 | 3,312 | 0,0490 |
| Sspon.08G0029900 | 9,745 | 0,0491 |
| Sspon.08G0029930 | -3,354 | 0,0491 |
| Sspon.08G0029990 | 1,098 | 0,0493 |
| Sspon.08G0030050 | 5,938 | 0,0493 |
| Sspon.08G0030140 | -4,347 | 0,0493 |
| Sspon.08G0030320 | 4,788 | 0,0493 |
| Sspon.08G0030480 | -2,109 | 0,0493 |
| Sspon.08G0030500 | -3,813 | 0,0494 |
| Sspon.08G0030510 | -5,150 | 0,0495 |
| Sspon.08G0030590 | -2,941 | 0,0496 |
| Sspon.08G0030720 | -3,296 | 0,0499 |
| Sspon.08G0030830 | -2,620 | 0,0499 |
| Sspon.08G0030860 | -3,379 | 0,0499 |
| Sspon.08G0030870 | 2,287 | 0,0499 |
| Sspon.08G0030880 | 6,175 | 0,0500 |
